# A widely distributed metalloenzyme class enables gut microbial metabolism of host- and diet-derived catechols

**DOI:** 10.1101/725358

**Authors:** Vayu Maini Rekdal, Paola Nol Bernardino, Michael U. Luescher, Sina Kiamehr, Peter J. Turnbaugh, Elizabeth N. Bess, Emily P. Balskus

## Abstract

Catechol dehydroxylation is a central chemical transformation in the gut microbial metabolism of plant- and host-derived small molecules. However, the molecular basis for this transformation and its distribution among gut microorganisms are poorly understood. Here, we characterize a molybdenum-dependent enzyme from the prevalent human gut bacterium *Eggerthella lenta* that specifically dehydroxylates catecholamine neurotransmitters available in the human gut. Our findings suggest that this activity enables *E. lenta* to use dopamine as an electron acceptor under anaerobic conditions. In addition to characterizing catecholamine dehydroxylation, we identify candidate molybdenum-dependent enzymes that dehydroxylate additional host-and plant-derived small molecules. These gut bacterial catechol dehydroxylases are specific in their substrate scope and transcriptional regulation and belong to a distinct group of largely uncharacterized molybdenum-dependent enzymes that likely mediate both primary and secondary metabolism in multiple environments. Finally, we observe catechol dehydroxylation in the gut microbiotas of diverse mammals, suggesting that this chemistry is present in habitats beyond the human gut. Altogether, our data reveal the molecular basis of catechol dehydroxylation among gut bacteria and suggest that the chemical strategies that mediate metabolism and interactions in the human gut are relevant to a broad range of species and habitats.

## Introduction

The human gastrointestinal tract is one of the densest microbial habitats on Earth. Possessing 150-fold more genes than the human genome, the trillions of organisms that make up this community (the human gut microbiota) harbor metabolic capabilities that expand the range of chemistry taking place in the body (*1–3*). Microbial metabolism affects host nutrition and health by breaking down otherwise inaccessible carbohydrates, biosynthesizing essential vitamins, and transforming endogenous and exogenous small molecules into bioactive metabolites (*4*). Gut microbial activities can also vary significantly between individuals, affecting the toxicity and efficacy of drugs (*5–9*), susceptibility to infection (*10–11*), and host metabolism (*12–13*). To decipher the biological roles of gut microbial metabolism, it is critical that we uncover the enzymes responsible for prominent transformations. This will not only increase the information gained from microbiome sequencing data but may also illuminate strategies for manipulating and studying microbial functions. Yet, the vast majority of gut microbial metabolic reactions have not yet been linked to specific enzymes.

A prominent but poorly understood gut microbial activity is the dehydroxylation of catechols (1,2-dihydroxylated aromatic rings), a structural motif commonly found in a diverse range of compounds that includes dietary phytochemicals, host neurotransmitters, clinically used drugs, and microbial siderophores (*14–16*) (Fig. 1A). Discovered over six decades ago, catechol dehydroxylation is a uniquely microbial reaction that selectively replaces the *para* hydroxyl group of the catechol with a hydrogen atom (*17*) (Fig. 1A). This reaction is particularly challenging due to the stability of the aromatic ring system. Prominent substrates for microbial dehydroxylation include the drug fostamatinib (*18*), the catecholamine neurotransmitters norepinephrine and dopamine (*19, 20*), and the phytochemicals ellagic acid (found in nuts and berries), caffeic acid (a universal lignin precursor in plants), and catechin (present in chocolate and tea) (*21–23*) (Fig. 1B). Dehydroxylation alters the bioactivity of the catechol compound (*24, 25*) and produces metabolites that act both locally in the gut and systemically to influence human health and disease (*18, 25–30*). However, the gut microbial enzymes responsible for catechol dehydroxylation have remained largely unknown.

**Figure 1.**
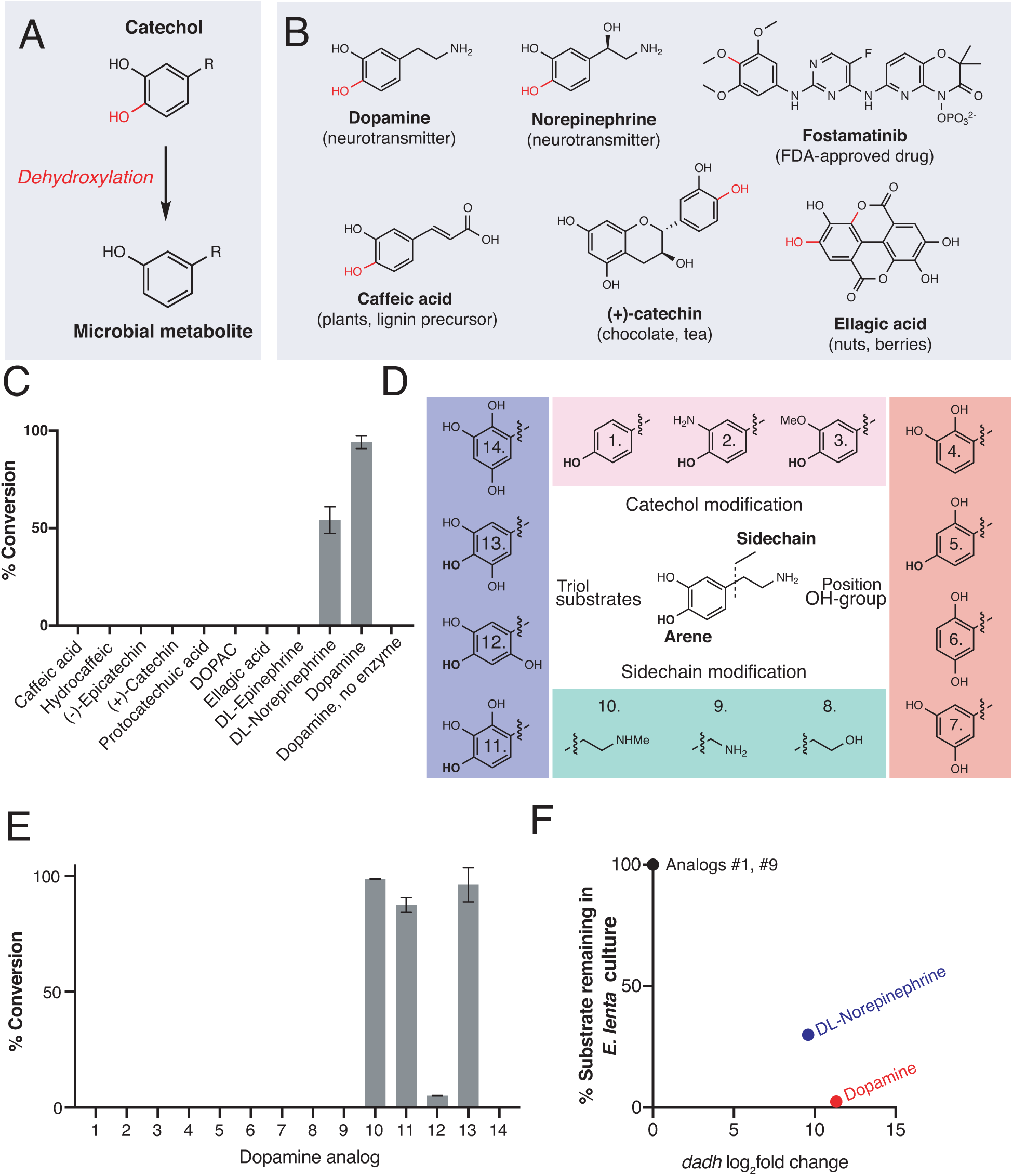
An enzyme from the prevalent human gut Actinobacterium *Eggerthella lenta* specifically metabolizes catecholamines that are available in the gut. A) The catechol structural motif is dehydroxylated by the gut microbiota. B) Examples of catechols known to be dehydroxylated by gut microbes. Red indicates the carbon-oxygen bond that is broken or the hydroxyl group that is removed in the dehydroxylation reaction. C) Activity of natively purified Dadh towards a panel of physiologically relevant catechol substrates. Enzyme (0.1 µM) was incubated with substrate (500 µM) for 22 hours at room temperature, followed by analysis using LC-MS. Bars represent the mean ± the standard error (SEM) of three biological replicates. See table S1 for the full chemical structures. D) Dopamine analogs evaluated in this study. E) Activity of natively purified Dadh towards dopamine analogs in C). Enzyme (0.1 µM) was incubated with substrate (500 µM) for 22 hours at room temperature, followed by analysis using LC-MS. Bars represent the mean ± the SEM of three biological replicates. See table S2 for the full chemical structures. F) Transcriptional induction and whole-cell dehydroxylation activity of *E. lenta* A2 in response to dopamine and a subset of dopamine analogs (500 µM each). Transcriptional induction was assessed using RNA-seq, with the fold induction shown on the x-axis (log_2_foldchange>4, FDR<0.01). To assess whole-cell metabolism, *E. lenta* was grown anaerobically for 48 hours in BHI medium with 500 µM of each substrate, and the culture supernatant was analyzed for dehydroxylated metabolites using a colorimetric assay or LC-MS. RNA-sequencing data represent the log2fold change of triplicate experiments. The metabolism data represent the mean ± the SEM of three biological replicates. Error bars are not visible because they are smaller than the datapoint.

We recently reported the discovery of a catechol dehydroxylating enzyme from the prevalent human gut Actinobacterium *Eggerthella lenta*. This enzyme participates in an interspecies gut microbial pathway that degrades the Parkinson’s disease medication L-dopa by catalyzing the regioselective *p*-dehydroxylation of dopamine to *m*-tyramine (*29*). To identify the enzyme, we grew *E. lenta* strain A2 with and without dopamine and used RNA sequencing (RNA-seq) to find genes induced by dopamine. Only 15 genes were significantly upregulated in the presence of dopamine, including a putative molybdenum-dependent enzyme that was induced >2500 fold. Hypothesizing this gene encoded the dopamine dehydroxylase, we purified the enzyme from *E. lenta* and confirmed its activity *in vitro*. Dopamine dehydroxylase (Dadh) is predicted to bind bis-molybdopterin guanine nucleotide (bis-MGD), a complex metallocofactor that contains a catalytically essential molybdenum atom (*31*). Our previous work illuminated a role for Dadh in dopamine metabolism by pure strains and complex communities. Here, we sought to explore the substrate scope of Dadh and its broader role in catechol dehydroxylation by the gut microbiota.

## Results

### A molybdenum-dependent enzyme from *Eggerthella lenta* specifically metabolizes catecholamines that are available in the gut

Because the human gut microbiota metabolizes a range of catecholic compounds (Fig. 1B), we first investigated whether the recently discovered Dadh possessed promiscuous dehydroxylase activity. We evaluated the reactivity of natively purified *E. lenta* A2 Dadh towards a panel of established or potential host- and diet-derived catechol substrates (table S1 and fig. S1). This enzyme displayed a narrow substrate scope, metabolizing only dopamine and the structurally related neurotransmitter norepinephrine, which differ only by the presence of a benzylic hydroxyl group (Fig. 1B). To identify the elements necessary for substrate recognition by Dadh, we profiled its activity towards synthetic and commercially available dopamine analogs (**1–14**) (Fig. 1D, fig. S1, and table S2). We found that Dadh tolerated only minor modifications to the dopamine scaffold, including a single *N*-methylation and the presence of additional hydroxyl groups on the aromatic ring (Fig. 1F). The catechol moiety was absolutely necessary for activity, and dehydroxylation required that at least one hydroxyl group to be in the *para* position relative to the aminoethyl substituent. These data demonstrated that Dadh is specific to the catecholamine scaffold.

This result prompted us to explore whether the transcriptional regulation of Dadh displayed similar specificity. Thus, we cultured *E. lenta* A2 in the presence of a subset of the dopamine analogs that we had tested in the previous experiment, measured dehydroxylation using liquid chromatography-mass spectometry (LC-MS), and profiled the global transcriptome using RNA-seq. Consistent with the biochemical data, we found that the regulation of *dadh* was also specific for the catecholamine scaffold (Fig. 1F, table S3). While the catecholamines dopamine and norepinephrine induced *dadh* expression and were dehydroxylated by *E. lenta*, analogs lacking the catechol (analog **1** in Fig. 1D) or having a shorter side chain (analog **9** in Fig. 1D) did not induce a transcriptional or metabolic response (Fig. 1F, table S3) (*29*). Together with our biochemical results, these transcriptional data suggest that Dadh may have evolved for the purpose of catecholamine neurotransmitter metabolism in *E. lenta*. We propose that dopamine is an endogenous substrate of this enzyme, because it was the best substrate both *in vitro* and *in vivo*, induced the highest levels of expression in *E. lenta*, and is produced at substantial levels by the human gastrointestinal tract (*32*).

In addition to uncovering a preference for the catecholamine scaffold, the substrate scope of Dadh *in vitro* reveal potential mechanistic distinctions between this enzyme and the only other biochemically characterized reductive aromatic dehydroxylase, 4-hydroxybenzoyl CoA reductase (4-HCBR) (*35*). 4-HCBR is a distinct molybdenum dependent-enzyme containing a monomeric molybdopterin co-factor that uses a Birch reduction-like mechanism to remove a single aromatic hydroxyl group from 4-hydroxybenzoyl CoA. While 4-HCBR requires an electron-withdrawing thioester group to stabilize radical anion intermediates (*33*), Dadh does not require an electron-withdrawing substituent and can tolerate additional electron-donating hydroxyl groups (Fig. 1D and E, analogs **11-13**). We preliminarily propose a mechanism for Dadh in which the dopamine *p*-hydroxyl group coordinates to the molybdenum center. This could be followed by tautomerization of the *m*-hydroxyl group to a ketone with protonation of the adjacent carbon atom. Oxygen atom transfer to molybdenum could be accompanied by rearomatization, providing the dehydroxylated product (fig. S2). Our proposal is consistent with the postulated mechanisms of other oxygen transfer reactions catalyzed by bis-MGD enzymes (*34, 35*).

### Dopamine promotes gut bacterial growth by serving as an alternative electron acceptor

The specificity of Dadh for dopamine suggested this metabolic activity might have an important physiological role in *E. lenta*. We noted the chemical parallels between catechol dehydroxylation and reductive dehalogenation, a metabolic process in which halogenated aromatics serve as alternative electron acceptors in certain environmental bacteria (*35*). This insight inspired the hypothesis that dopamine dehydroxylation could serve a similar role in anaerobic respiration for gut bacteria. While we observed no growth benefit when *E. lenta* was grown in complex BHI medium containing dopamine (fig. S3), we found that including dopamine in a minimal medium lacking electron acceptors (basal medium) increased the endpoint optical density of *E. lenta* cultures (Fig. 2A). This growth-promoting effect was only observed in dopamine-metabolizing *E. lenta* strains, as non-metabolizing strains that express an apparently inactive enzyme (*29*) did not gain a growth advantage (Fig. 2A, fig. S4, and table S4). The effect of dopamine on *E. lenta* contrasts with recent studies of digoxin, a drug that is reduced by *E. lenta* without impacting growth in the same medium (*6*).

**Figure 2.**
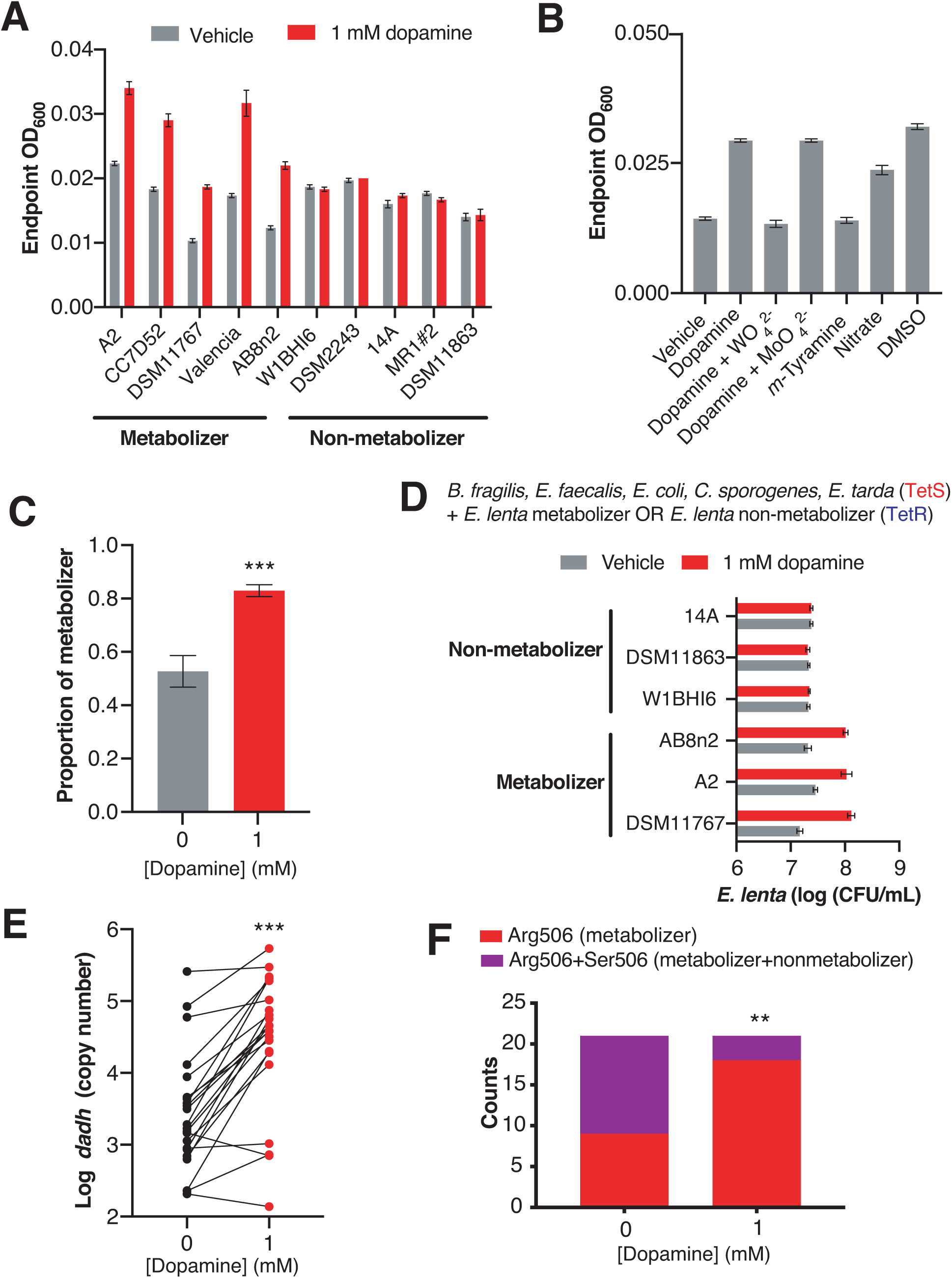
Dopamine increases gut bacterial growth by serving as an alternative electron acceptor. A) Growth of dopamine metabolizing and non-metabolizing *E. lenta* strains in minimal medium limited in electron acceptors (basal medium) containing 10 mM acetate. Strains were grown anaerobically for 48-72 hours at 37 °C before growth was assessed. Bars represent the mean ± the SEM of three biological replicates. B) Tungstate inhibits of growth and dopamine metabolism by *E. lenta* A2 in basal medium containing 10 mM acetate. *E. lenta* was grown anaerobically for 48 hours at 37 °C. Dopamine, *m*-tyramine, and nitrate were added to a final concentration of 1 mM, while DMSO was added to a final concentration of 14 mM at the time of inoculation. Tungstate (WO_4_^2–^) and molybdate (MoO_4_^2–^) were added at a final concentration of 0.5 mM. Bars represent the mean ± the SEM of three biological replicates. C) Competition of dopamine metabolizing (Valencia) and non-metabolizing (W1BHI6) *E. lenta* strains in basal medium containing 10 mM acetate. Strains were grown together for 72 hours at 37 °C and were then plated. Antibiotic resistance was used to determine strain identity. Bars represent the mean ± the SEM of six biological replicates. (***P<0.001, two-tailed unpaired t-test). D) Growth of defined gut bacterial consortia containing dopamine metabolizing and non-metabolizing *E. lenta* strains in basal medium containing 10 mM acetate. Tetracycline resistant (TetR) *E. lenta* strains were grown with tetracycline sensitive (TetS) gut isolates for 48 hours at 37 °C. Plating on BHI containing tetracycline allowed enumeration of *E. lenta*. Bars represent the mean ± the SEM of three biological replicates. E) Abundance of *dadh* in complex human gut communities cultured *ex vivo*. Samples from unrelated individuals (n = 24) were grown for 72 hours at 37 °C in basal medium containing 10 mM acetate with or without dopamine and qPCR was used to assess abundance of *dadh*. Two individuals were excluded from this analysis as they did not demonstrate quantitative metabolism of dopamine after incubation. Each point represents a different individual. Lines connect data from the same individual between the two conditions. (***P<0.001, two-tailed unpaired t-test). F) Counts of *dadh* variants in the presence and absence of dopamine. The same gDNA used in E) was used to amplify full-length *dadh* and determine the SNP status at position 506 using Sanger sequencing. As in panel E, two individuals were removed prior to analysis as they did not demonstrate quantitative metabolism of dopamine after incubation. In addition, one individual was not included in this analysis due to failure of obtaining high quality sequencing data. (**P<0.01, Fisher’s exact test).

We further investigated the relationship between dopamine and bacterial growth in the metabolizing strain *E. lenta* A2. The growth increase observed in response to dopamine was dose-dependent (fig. S5), mirrored the effects of the known electron acceptors DMSO and nitrate (*6, 36*), and did not derive from the product of dopamine dehydroxylation, *m*-tyramine (Fig. 2B). Additionally, the growth benefit was directly tied to dopamine dehydroxylation. Inclusion of tungstate in the growth medium, which inactivates the big-MGD cofactor of Dadh, blocked metabolism and inhibited the growth increase. In contrast, inclusion of molybdate in the growth medium did not impact growth or metabolism (Fig. 2B and fig. S6). Molybdate and tungstate alone did not impact *E. lenta* A2 growth in the basal medium (fig. S7). Taken together, these results indicate that active metabolism of dopamine provides a growth advantage to *E. lenta*, likely by serving as an alternative electron acceptor.

We next examined whether dopamine could promote *E. lenta* growth in microbial communities. First, we competed dopamine metabolizing and non-metabolizing *E. lenta* strains in minimal medium. *E. lenta* is genetically intractable, preventing the use of engineered plasmids encoding defined fluorescence or antibiotic resistance as markers of strain identity. Instead, we took advantage of intrinsic differences in tetracyline (Tet) resistance to differentiate the closely related strains in pairwise competitions (*37*). Inclusion of dopamine in growth medium significantly increased the proportion of the metabolizer relatively to the non-metabolizer in this competition experiment (p<0.001, two-tailed unpaired t-test) (Fig. 2C and fig. S8). This was driven by the growth increase of the metabolizer rather than a decrease in the non-metabolizer (Fig. 2C and fig. S8).

Next, we explored the impact of dopamine on Tet-resistant *E. lenta* in the presence of a defined bacterial community representing the major phylogenetic diversity in the human gut (Fig. 2D and table S4) (*38, 39*). We found that including dopamine in the medium boosted the growth of metabolizers by an order of magnitude while non-metabolizing strains did not gain a growth advantage (Fig. 2D). Finally, we evaluated the impact of dopamine on *E. lenta* strains present in complex human gut microbiotas. We cultured fecal samples from 24 unrelated subjects *ex vivo* in the presence and absence of dopamine and used qPCR to assess the abundance of metabolizing *E. lenta* strains and *dadh*. Strikingly, both *dadh* and *E. lenta* significantly increased by an order of magnitude in cultures containing dopamine (p<0.005, two-tailed unpaired t-test) (Fig. 2E) (fig. S9). We also amplified the full length *dadh* gene from these cultures and sequenced the region harboring the SNP that distinguishes metabolizing and non-metabolizing strains (*29*). These assays indicated that the increase in *dadh* abundance in the complex communities was accompanied by a significant shift from a mixture of inactive and active *dadh* variants to a dominance of the metabolizing R506 variant (p<0.01, Fisher’s exact test) (Fig. 2F). Overall, these results show that dopamine dehydroxylation increases the fitness of metabolizing *E. lenta* strains in both defined and complex microbial communities.

### A screen of human gut Actinobacteria uncovers dehydroxylation of host and plant-derived catechols

Having uncovered Dadh’s specialized role in gut bacterial dopamine metabolism, we sought to identify strains and enzymes that could dehydroxylate other catechol substrates. Among human gut bacteria, only *Eggerthella* and closely related members of the Actinobacteria phylum have been reported to perform catechol dehydroxylation. For example, *Eggerthella* metabolizes dopamine and (+)-catechin (*40*), while related *Gordonibacter* species dehydroxylate ellagic acid (*41*) and a lignan derivative in the multi-step biosynthesis of the anti-cancer metabolite enterodiol (*42*). These reports suggest that Actinobacteria could be a promising starting point to identify new dehydroxylating strains and enzymes. Thus, we assembled a library of related gut Actinobacteria (*37*) and screened these strains individually (n=3 replicates) for metabolism of a range of compounds relevant in the human gut, including plant- and host-derived small molecules, bacterial siderophores, and FDA-approved catecholic drugs (*14–16*) (table S5) (Fig. 3A). We initially used a colorimetric assay that detects catechols to assess metabolism, which allowed us to rapidly screen for potential catechol depletion across the collection of 25 strains and 19 compounds of interest.

**Figure 3.**
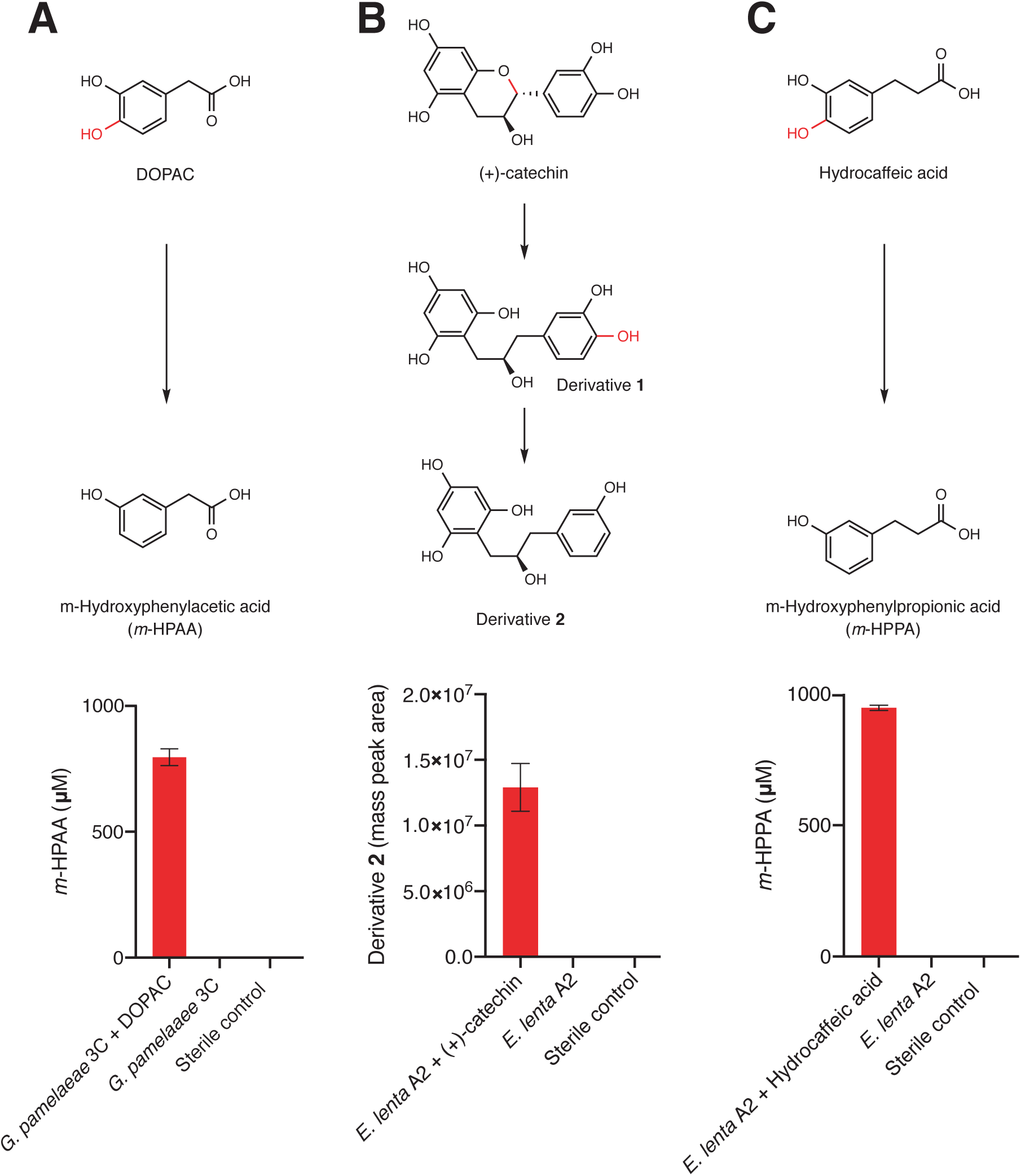
Dehydroxylation of DOPAC, (+)-catechin, and hydrocaffeic acid by *Gordonibacter pamelaeae* 3C and *Eggerthella lenta* A2. A-C) Pathways for metabolism of (A) DOPAC, (B) (+)- catechin, (C) and hydrocaffeic acid by human gut Actinobacteria. While DOPAC and hydrocaffeic acid are dehydroxylated directly, (+)-catechin metabolism proceeds by initial benzyl ether reduction followed by dehydroxylation of the catecholic derivative. A) Metabolism of DOPAC by *G. pamelaeae* 3C. This strain was grown in BHI medium with and without 1 mM DOPAC for 48 hours at 37 °C. Metabolism was assessed using LC-MS/MS. Bars represent the mean ± the SEM concentration of the metabolite *m*-hydroxyphenylacetic acid (*m-*HPAA) resulting from direct DOPAC dehydroxylation (three biological replicates). B) (+)-catechin metabolism by *E. lenta* A2. This strain was grown in BHI medium with and without 1 mM (+)-catechin for 48 hours at 37 °C. Metabolism was assessed using high resolution LC-MS. Bars represent the mean ± the SEM mass peak area of the Extracted Ion Chromatogram (EIC) for the dehydroxylated derivative **2** shown in A) (three biological replicates). Due to the absence of a pure standard, integrated peak area of the high-resolution mass is displayed. C) Metabolism of hydrocaffeic acid by *E. lenta* A2. This strain was grown in BHI medium with and without 1 mM hydrocaffeic acid for 48 hours at 37 °C. Metabolism was assessed using LC-MS/MS. Bars represent the mean ± the SEM concentration of *m-*hydroxyphenylpropionic acid (*m-*HPPA) resulting from the direct dehydroxylation of hydrocaffeic acid (three biological replicates).

We observed complete depletion of several host and diet-derived catechols in this initial screen (fig. S10). We chose to focus on the dehydroxylation of hydrocaffeic acid, (+)-catechin, and DOPAC for further characterization, repeating the incubations with these compounds and using LC-MS/MS to confirm the production of dehydroxylated metabolites. This analysis confirmed that both DOPAC and hydrocaffeic acid are directly dehydroxylated by members of this library, while (+)-catechin undergoes benzyl ether reduction followed by dehydroxylation to give the ring-opened derivative **2**, as has been observed previously (*40*) (Fig. 3 and table S6). While (+)-catechin metabolism has been previously linked to *Eggerthella* (*40*), the dehydroxylation of DOPAC and hydrocaffeic acid has only been previously observed by complex gut microbiota communities (*17, 21*). The variability in these activities across closely related gut bacterial strains suggests that different enzymes might dehydroxylate different catechols.

### Gut Actinobacteria dehydroxylate individual catechols using distinct enzymes

We next sought to determine the molecular basis of the dehydroxylation reactions examined above. To test the hypothesis that dehydroxylation resulted from specific rather than promiscuous enzymes, we first established that dehydroxylation is an inducible activity in *Gordonibacter* and *Eggerthella* strains (fig. S11). This allowed us to use cell lysates as a proxy for transcriptional induction and a means of examining dehydroxylase activity. We grew *E. lenta* A2 in the presence of (+)-catechin, hydrocaffeic acid, and dopamine, and grew *G. pamelaeae* 3C in the presence of DOPAC. We then screened each anaerobic lysate for its activity towards all of these substrates. Consistent with our prediction, each lysate quantitively dehydroxylated only the catechol substrate with which the strain had been grown. While the *E. lenta* lysates did not display any promiscuity (Fig. 4A), cell lysate from *G. pamelaeae* grown in the presence of DOPAC displayed low activity (<40%) towards hydrocaffeic acid, which structurally resembles DOPAC (Fig. 4B). Overall, these results suggest that different catechol substrates induce the expression of distinct dehydroxylase enzymes that are specific in their biochemical activity and transcriptional regulation. We expected that these enzymes would be molybdenum-dependent like Dadh, as these are the only types of enzymes known to be capable of aromatic dehydroxylation (*29, 31, 33–34*).

**Figure 4.**
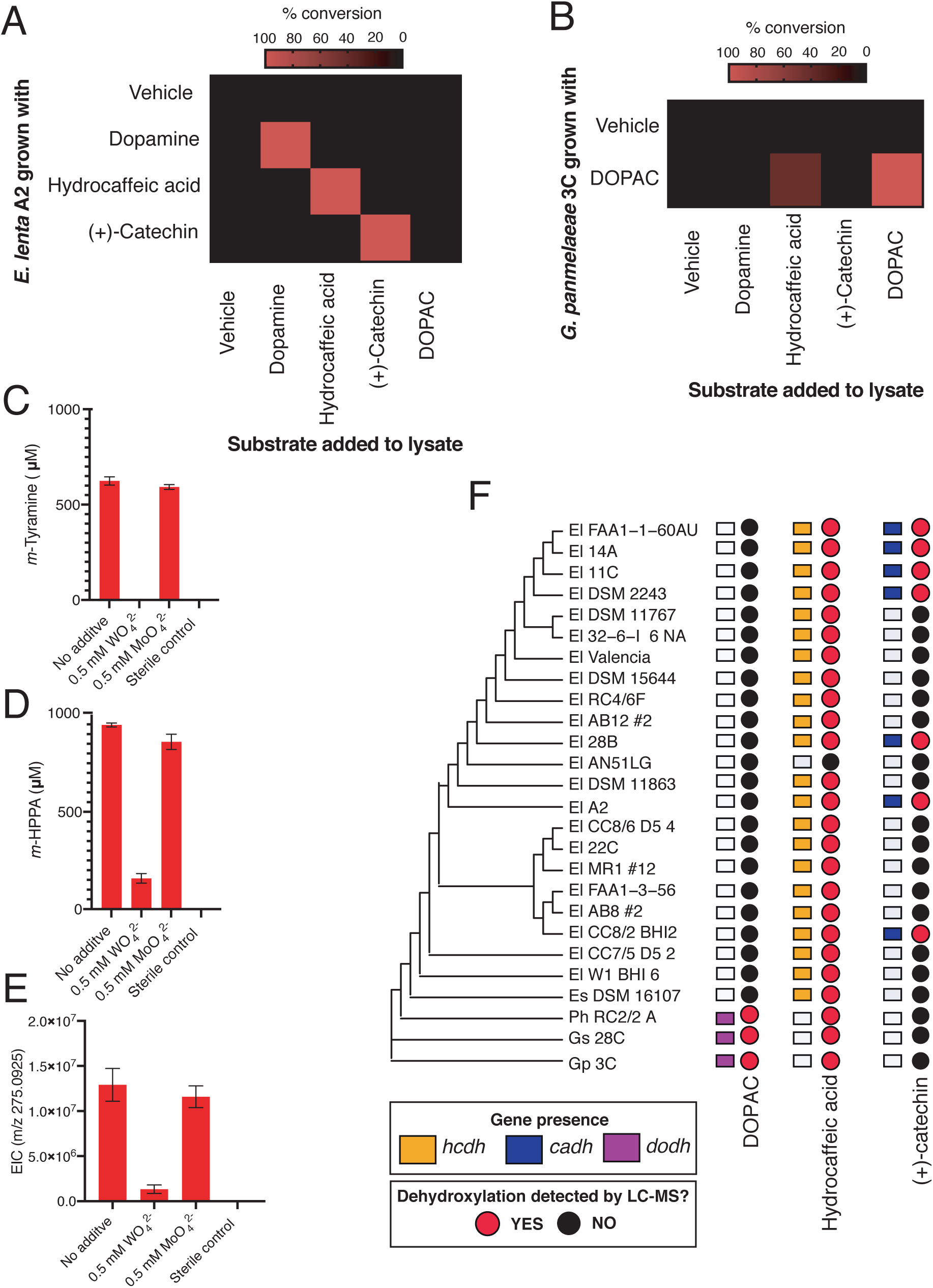
Gut Actinobacteria dehydroxylate individual catechols using distinct enzymes. A) Specificity of dehydroxylase regulation and activity in *E. lenta* A2. *E. lenta* A2 was grown anaerobically in BHI medium containing 1 % arginine and 10 mM formate. 0.5 mM of catechol was added to induce dehydroxylase expression, followed by anaerobic lysis and enzyme assays. Crude lysates were exposed to different substrates (500 µM) and reactions were allowed to proceed anaerobically for 20 hours. Assays mixtures were analyzed using LC-MS. Heat map represents the mean of three biological replicates. B) Specificity of DOPAC dehydroxylase regulation and activity in *G. pamelaeae* A2. *G. pamelaeae* 3C was grown anaerobically in BHI medium containing 10 mM formate. 0.5 mM of catechol was added to induce dehydroxylase expression, followed by anaerobic lysis and enzyme assays. Crude lysates were exposed to different substrates (500 µM) and reactions were allowed to proceed anaerobically for 20 hours. Assays mixtures were analyzed using LC-MS. Heat map represents the mean of three biological replicates C-E) Culturing in the presence of tungstate reveals the molybdenum dependence of catechol dehydroxylation by *E. lenta* A2. The strain was grown anaerobically with 1 mM dopamine (C), hydrocaffeic acid (D), or (+)-catechin (E) and with 0.5 mM tungstate (WO_4_^2−^) or molybdate (MoO_4_^2−^) for 48 hours 37 °C. Metabolites in culture supernatants were analyzed by LC-MS. Bars represent the mean ± the SEM of three biological replicates. F) Distribution of putative catechol dehydroxylases and their associated metabolic activities across the gut Actinobacterial library used in our study. The three represents the phylogeny of gut Actinobacterial strains adapted from (*2*). El = *Eggerthella lenta*, Es = *Eggerthella sinesis*, Ph = *Paraggerthella*, Gs = *Gordonibacter* sp., Gp = *Gordonibacter pamelaeae*. Squares represent gene presence/absence of select dehydroxylases across gut Actinobacterial strains (90% coverage, 75% amino acid identity cutoff, e-value=0). To assess metabolism of catechol substrates by gut Actinobacterial strains, individual strains were grown in triplicate in the presence of a single catechol substrate for 48 hours at 37 °C in BHI medium. Metabolism was assessed using LC-MS/MS. A red dot indicates that the mass peak area of the dehydroxylated product was detected in cultures from this strain, while a black dot indicates lack of metabolism.

To identify the molecular basis of (+)-catechin and hydrocaffeic acid dehydroxylation in *E. lenta* A2, we turned to RNA-seq, an approach that we successfully used previously to identify the dopamine dehydroxylase (*29*). We grew *E. lenta* A2 to early exponential phase and then added each catechol substrate, harvesting the cells after 1.5 hours of induction. Hydrocaffeic acid and (+)-catechin each upregulated a number of genes (25 and 41, respectively), including two predicted molybdenum-dependent enzymes (tables S7 and S8). While one of these predicted molybdenum-dependent enzymes was among the highest upregulated genes in response to each substrate (450-fold induced in response to catechin, >2000-fold with hydrocaffeic acid), the other enzyme was only 3-fold induced relative to the vehicle. Thus, we propose that the highest upregulated molybdenum-dependent enzyme in each dataset is the most reasonable candidate dehydroxylase. The enzyme upregulated in response to hydrocaffeic acid (Elenta-A2_02815) shared 35.3% amino acid identity with Dadh, while the enzyme upregulated in response to (+)-catechin shared 50.9% amino acid identity with Dadh (tables S7-S9).

To evaluate the involvement of a molybdenum enzyme in each dehydroxylation reaction, we cultured *E. lenta* in the presence of tungstate, which inactivates the metallocofactor (*29, 44*). As with dopamine dehydroxylation, tungstate inhibited dehydroxylation of (+)-catechin and hydrocaffeic acid by *E. lenta* A2 without inhibiting growth in the rich BHI medium, suggesting these activities are indeed molybdenum dependent (Fig 4C-E and fig. S12). Tungstate did not inhibit benzyl ether reduction of (+)-catechin, indicating this step is performed by a distinct enzyme (fig. S13). Finally, we found that the distribution of the genes encoding these the putative hydrocaffeic acid and (+)-catechin dehydroxylating enzymes across closely related *Eggerthella* strains correlated with the ability of these organisms to metabolize each substrate (Fig. 4F). For example, all *Eggerthella* strains except AN5LG harbored the putative hydrocaffeic acid dehydroxylase and could dehydroxylate this substrate. Similarly, carriage of the putative catechin dehydroxylase correlated with (+)-catechin metabolism. Taken together, these data strongly suggest these distinct molybdenum-dependent enzymes dehydroxylate hydrocaffeic acid and (+)- catechin.

We next sought to identify the enzyme responsible for dehydroxylation of DOPAC in *G. pamelaeae* 3C. We added DOPAC to *G. pamelaeae* 3C cultures at mid-exponential phase and harvested cells after 3 hours of induction when the cultures had reached early stationary phase. In this experimental setup, *G. pamelaeae* 3C upregulated 99 different genes, including four distinct molybdenum-dependent enzyme-encoding genes (table S10). One of these genes (C1877_13905) was among the highest upregulated genes across the dataset (>1700-fold induced). To further explore the association between this gene and DOPAC dehydroxylation, we repeated the RNA-seq experiment, growing *G. pamelaeae* 3C in the presence of DOPAC from the time of inoculation and then harvesting cells in mid-exponential phase as soon we could detect metabolism (12 hours of total growth), In this experiment, the same molybdenum-dependent enzyme-encoding gene (C1877_13905) that we observed highly upregulated in our first experiment (table S11) was among the highest upregulated genes. The only two other molybdenum-dependent enzymes induced in this experiment were expressed at order magnitude lower levels (2-fold induced), making the highest upregulated molybdenum-dependent enzyme observed across both datasets a reasonable candidate DOPAC dehydroxylase.

This assignment is also supported by comparative genomics. First, the presence of the gene encoding this candidate molybdenum enzyme correlated with DOPAC dehydroxylation among members of our gut Actinobacterial library (Fig. 4F). Consistent with our lysate assays, those organisms harboring the putative DOPAC dehydroxylase also had activity towards hydrocaffeic acid, which could explain the pattern of hydrocaffeic acid metabolism across the gut Actinobacterial library (Fig. 4F). Finally, Dadh was the most closely related biochemically characterized homolog to the candidate dehydroxylase (20% amino acid ID), and the putative DOPAC dehydroxylase is also similar (45% amino acid ID) to catechol lignan dehydroxylase (Cldh), an enzyme recently implicated in the dehydroxylation of the lignan dmSECO in *Gordonibacter* (*42*) (table S9). Though this functional assignment awaits biochemical confirmation, we propose that the highest upregulated enzyme across our two independent datasets is the DOPAC dehydroxylase. Interestingly, unlike with the *Eggerthella* dehydroxylases, tungstate did not inhibit dehydroxylation of DOPAC by *G. pamelaeae* (fig. S14). This may be explained if the dehydroxylating enzyme can use both molybdenum and tungsten for catalysis, as is seen in certain closely-related enzymes (*43*).

Despite being expected to perform the same type of chemical reaction, the putative dehydroxylases from *E. lenta* and *G. pamelaeae* differ from each other in sequence identity, genomic context, and predicted subunit composition (table S9 and fig. S15). The dopamine, catechin and hydrocaffeic acid dehydroxylases from *E. lenta* (*dadh*, *cadh* and *hcdh*, respectively) are likely membrane-bound complexes as they co-localize with genes encoding an electron shuttling 4Fe-4S ferredoxin and a putative membrane anchor (fig. S15) (*44, 45*). These enzymes all carry a Twin-Arginine-Translocation (TAT) signal sequence, suggesting they are exported from the cytoplasm. In contrast, the *G. pamelaeae* enzymes are smaller than the *E. lenta* enzymes and co-localize with a gene predicted to encode a small electron shuttling 4Fe-4S protein, suggesting the potential involvement of a protein complex that is not membrane anchored (fig. S15). The *G. pamelaeae* dehydroxylases are also encoded adjacent to members of the Major Facilitator Superfamily, transporters that may import or export the catechol substrates or dehydroxylated metabolites (fig. S15). Together, these data support a model in which catechol dehydroxylation is performed by distinct subtypes of molybdenum-dependent enzymes.

### Catechol dehydroxylases are distinct from other molybdenum-dependent enzymes and are widely distributed across sequenced microbes

The broader differences in subunit composition and genomic context between the *E. lenta* and *G. pamelaeae* dehydroxylases made us curious about their relationship to other characterized molybdenum-dependent enzymes. Surprisingly, these catechol dehydroxylases differ from the only other biochemically characterized aromatic dehydroxylase, 4-HCBR; whereas 4-HCBR belongs to the xanthine oxidase family of molybdenum-dependent enzymes, the catechol dehydroxylases belong to the bis-MGD family of molybdenum-dependent enzymes, suggesting independent evolutionary origins (*31, 33–34*). Further phylogenetic analysis revealed that catechol dehydroxylases form a unique clade within the bis-MGD enzyme family, clustering away from the pyrogallol hydroxytransferase (Pht), the only other bis-MGD enzyme known to modify the aromatic ring of a substrate (*46*) (Fig. 5A and table S13). The catechol dehydroxylases are instead most closely related to acetylene hydratase, an enzyme that adds water to acetylene to provide a carbon source for the marine Proteobacterium *Pelobacter acetylenicus* (Fig. 6A) (*34, 47–48*). Consistent with this phylogenetic analysis, a sequence similarity network (SSN) analysis using sequences of bis-MGD enzymes revealed distinct clusters of catechol dehydroxylases, further suggesting these enzymes are functionally distinct from known family members (fig. S16). The clustering of the dehydroxylases on the SSN did not reflect the phylogeny of the organisms because sequences from both *Eggerthella* and *Gordonibacter* were widely distributed across clusters containing distinct, biochemically characterized enzymes (fig. S16). In addition, we found that the two dehydroxylase clusters contained sequences from organisms other than *Eggerthella* and *Gordonibacter* (fig. S16). Based on these data, we propose that catechol dehydroxylases belong to a distinct group of molybdenum-dependent enzymes.

**Figure 5.**
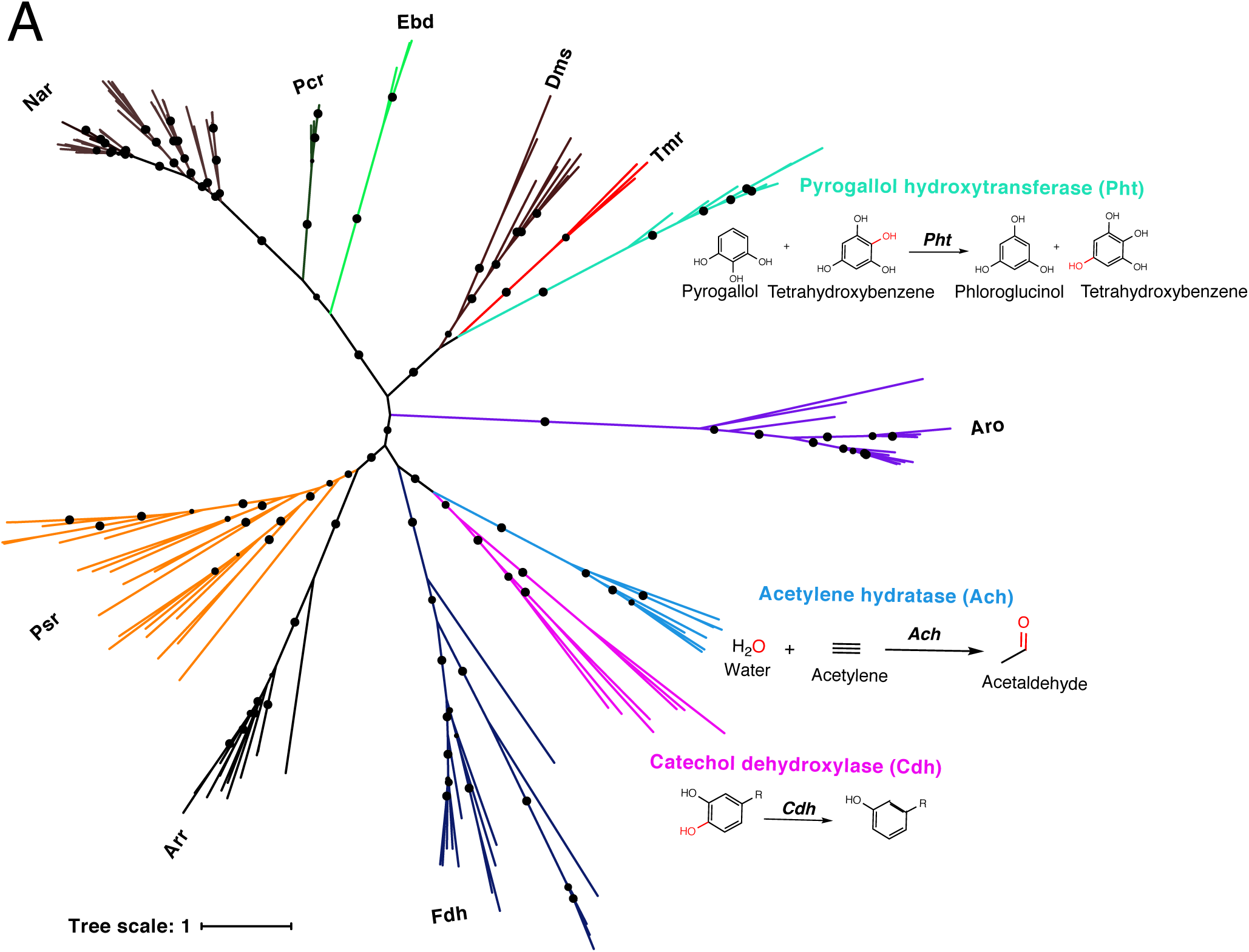
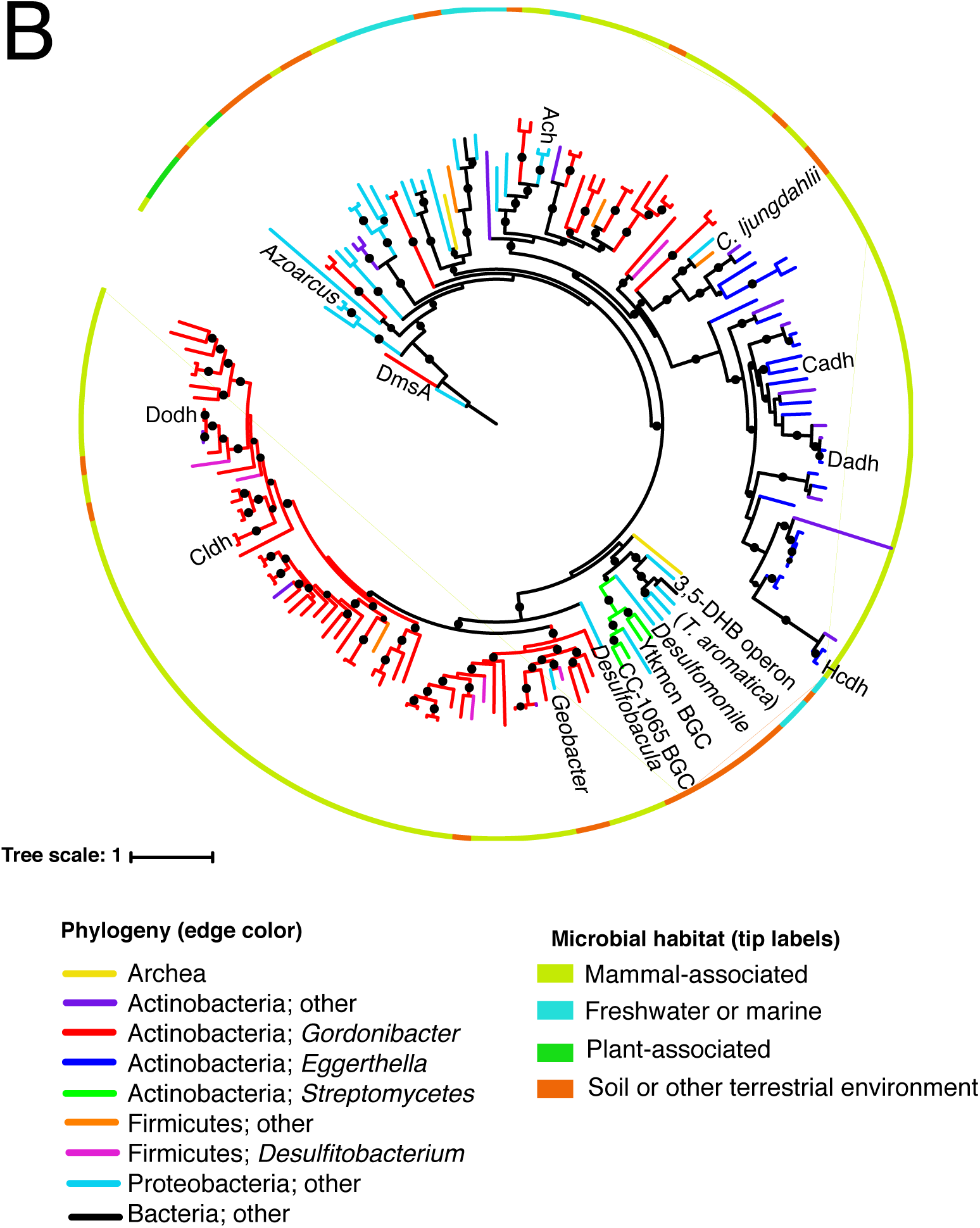
Catechol dehydroxylases are distinct from other molybdenum-dependent enzymes and are widely distributed across sequenced microbes. Phylogenetic analysis of newly discovered catechol dehydroxylases reveals a unique evolutionary origin and relationship to acetylene hydratase. Psr = polysulfide reductase; Arr = arsenate reductase; Fdh = formate dehydrogenase; Pht = phloroglucinol transhydroxylase; Dms = DMSO reductase; Tmr = TMAO reductase; Aro = arsenite oxidase; Ebd = ethylbenzene dehydrogenase; Pcr = perchlorate reductase; Nar = nitrate reductase. The maximum likelihood tree was constructed using sequences from (*48*) as well as additional family members and reproduced the known phylogeny of this enzyme family. Black circles on branches indicate bootstrap values greater than 0.7. B) Maximum likelihood phylogenetic tree for catechol dehydroxylase homologs identified by querying 26 gut Actinobacterial genomes (*37*) and the NCBI nucleotide collection for Dadh and Cldh (see methods for details). The color of the lines indicates the phylogeny of the organism harboring the homolog. The color of the border indicates the primary habitat from which the organism was originally isolated. Names at the end of the branches indicate either enzyme names, characterized gene clusters that encode the enzyme, or the name of the genera or species harboring the enzyme. BGC = biosynthetic gene cluster; Ytkmcn = yatakemycin; Ach = acetylene hydratase from *P. acetylenicu*s; DmsA = biochemically characterized DMSO reductase from *E. coli*, which was used as an outgroup to root the tree. The examples mentioned in the figure are described in the main text. Black circles on branches indicate bootstrap values greater than 0.7.

**Figure 6.**
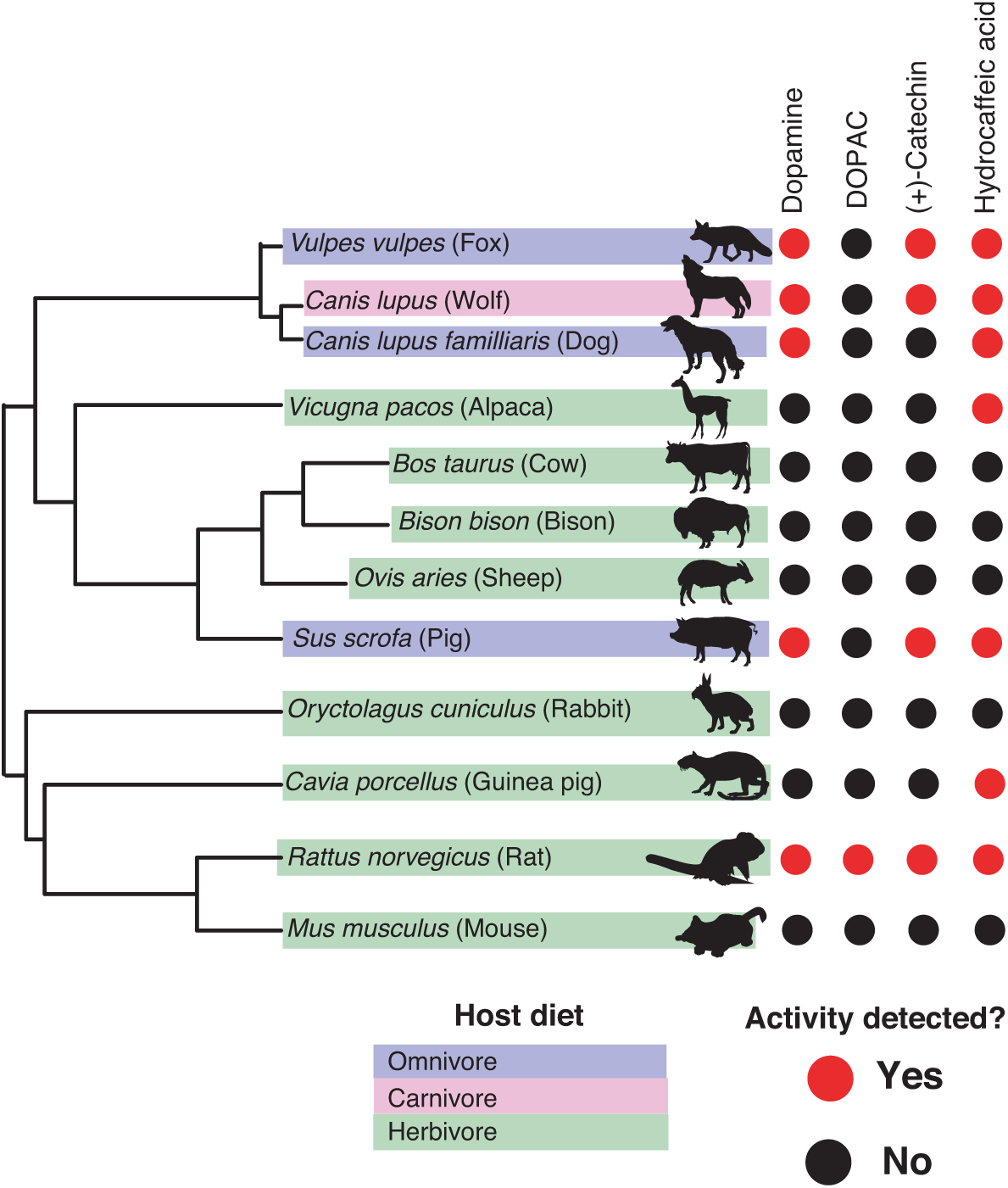
Gut microbiotas of mammals representing distinct diets and phylogenetic origins can dehydroxylate catechols. Catechol dehydroxylation of dopamine, DOPAC, (+)-catechin, and hydrocaffeic acid by gut microbiota samples from mammals spanning distinct diets and phylogenetic groups. Gut communities from 12 different mammals and 3 individuals per animal were cultured anaerobically for 96 hours in basal medium with 0.5 mM catechol at 37 °C. The results summarize animals and individuals where the known dehydroxylation pathways examined in human gut Actinobacteria took place, as assessed by LC-MS/MS (fig. S15). Red indicates that metabolism took place in at least one of the individuals, and black indicates lack of metabolism, as assessed by the detection of the dehydroxylated metabolite using LC-MS/MS. The phylogenetic tree was created using the aptg plugin in R and missing branches were added manually based on mammalian phylogeny. The icons were adapted under a Creative Commons license (https://creativecommons.org/licenses/by/3.0/) at phylopic (http://phylopic.org), including Alpaca logo (made my Steven Traver), Bison (Lukasiniho). Cow (Steven Traver), Dog (Tracy A Heath), Fox (Anthony Caravaggi), Guinea pig (Zimices), Mouse (Madeleine Price Ball), Pig (Steven Traver), Rabbit (Steven Traver), Rabbit (Steven Traver), Rat (Rebecca Groom), Sheep (Zimices), and Wolf (Tracy A Heath).

To assess whether this enzyme group harbors greater diversity than the enzymes described in this study, we queried the NCBI nucleotide database and our collection of Actinobacterial genomes for homologs of the *Eggerthella* and *Gordonibacter* catechol dehydroxylases. Phylogenetic analyses of the enzyme sequences revealed a large diversity of putative dehydroxylases, including numerous uncharacterized sequences encoded in both *Gordonibacter* and *Eggerthella* genomes (Fig. 5B). This highlights that catechol dehydroxylases likely have diversified within these closely related gut Actinobacteria and that many substrate-enzyme pairs remain to be discovered. Our phylogenetic analysis also revealed that catechol dehydroxylases are not restricted to human-associated Actinobacteria and appear to be part of a larger group of bis-MGD enzymes present in diverse bacteria and even Archaea (Fig. 5B). These organisms come from mammal-associated, plant-associated, soil, and aquatic habitats. Notable organisms encoding putative dehydroxylases include soil-dwelling Streptomycetes (*49, 50*), the industrially important anaerobe *Clostridium ljungdahlii* (*51*), and a large number of anaerobic bacterial genera known for their ability degrade aromatic compounds, including *Azoarcus, Thauera, Desulfobacula, Geobacter, Desulfumonile*, and *Desulfitobacterium* (Fig. 5B) (*52–59*). Some sequences are shared between gut and environmental microbes, raising questions about the evolutionary processes driving this diversity (Fig. 5B).

As the vast majority of dehydroxylase homologs remain uncharacterized, it is difficult to assign the biochemical activities of the major clades and define the characteristic features of these enzymes. However, we are confident that at least some portion of the sequences captured in this analysis are true catechol dehydroxylases. First, we found that representatives sequences from across our phylogenetic tree are more closely related to acetylene hydratase and the *Gordonibacter* and *Eggerthella* dehydroxylases than to any other member of the bis-MGD enzyme family, indicating shared evolutionary origins (table S14 and figs. S17 and S18). Moreover, we noticed that recent genetic studies have implicated some homologs from environmental bacteria in catechol dehydroxylation. For instance, two catechol dehydroxylase homologs from *Streptomyces* are present in biosynthetic gene clusters that produce the potent anti-tumor compounds yatakemycin and CC-1065 (*49, 50*) (Fig 5B). Gene knock-out and complementation studies revealed that the dehydroxylase homolog is essential for CC-1065 production and likely catalyzes reductive dehydroxylation of a late-stage biosynthetic intermediate (*49*). Another homolog is present in the 3,5-dihydroxybenzoate (3,5-DHB) degradation operon within the anaerobic soil Proteobacterium *Thaeura aromatica* (Fig. 5B). Strains lacking this enzyme exhibit impaired growth on 3,5-DHB as a sole carbon source, suggesting a possible role for this enzyme in metabolizing the one of the two catecholic intermediates involved in this pathway (*58, 60–61*). Based on this analysis, we conclude that the catechol dehydroxylases harbor vast uncharacterized diversity that contributes to both primary and secondary metabolic pathways in habitats beyond the human gut.

### Catechol dehydroxylase reactivity is conserved across the gut microbiotas of mammals representing distinct diets and phylogenetic origins

Our phylogenetic analysis suggested that catechol dehydroxylase activity is present in a range of habitats and phyla, making us curious whether we could detect this metabolism in microbial communities beyond the human gut. As a first step, we explored whether this reactivity was present in gut microbiotas of non-human mammals. We assembled a panel of gut microbiota samples from 12 different mammals representing diverse phylogenetic origins and diets, with 3 individuals per mammal (*62–63*) (Fig. 7 and fig. S19). We cultured these gut communities anaerobically *ex vivo*, assessed metabolism using a colorimetric assay, and confirmed potential hits using LC-MS/MS (fig. S15). We observed catechol dehydroxylation by at least one of the three individuals for each species across both diet and phylogeny (Fig. 6). Hydrocaffeic acid dehydroxylation occurred in >50% of species, while dopamine and (+)-catechin metabolism were observed in 5/12 and 4/12 animals, respectively (Fig. 6). Other substrates were metabolized by fewer mammals, including DOPAC, which was only metabolized by the rat gut microbiota. Interestingly, the rat samples had activity towards all compounds tested. While a larger sample size is required to reach clear conclusions about possible links between metabolism of specific catechols and individual mammal gut microbiota, our results clearly demonstrate that catechol dehydroxylation is found in distantly related mammals that have large differences in gut microbial species composition and gene content (*62–64*). Therefore, we conclude that that catechol dehydroxylation is present in a variety of different microbial habitats.

## Discussion

For many decades the human gut microbiota has been known to dehydroxylate catechols, but the molecular basis of this enigmatic transformation has remained largely unknown. In this study, we characterized the specificity of the activity and regulation of an enzyme that metabolizes dopamine (Dadh). We then used this knowledge to identify candidate enzymes that dehydroxylate additional host-and plant-derived small molecules. Together, the catechol dehydroxylases represent a new group of molybdenum-dependent enzymes that is present in diverse microbial phyla and environments. Our studies of Dadh revealed a high specificity for catecholamines, supporting the hypothesis that the physiological role of this enzyme is to enable neurotransmitter metabolism by *E. lenta*. This idea is also consistent with recent observations of gut bacteria using specific neurotransmitters for growth (*66*). To our knowledge, Dadh is the first enzyme from a human gut symbiont demonstrated to metabolize catecholamines. However, interactions between catecholamines and intestinal pathogens are well-characterized and have long been known as key players in virulence and infection (*67–68*). Whereas pathogenic organisms such as *Y. enterocolitica*, *E. coli* and *S. enterica* require the intact catechol group of dopamine and norepinephrine to sequester iron and boost growth (*67–70*), *E. lenta* dehydroxylates the catechol and likely uses it as an electron acceptor. Thus, Dadh might represent a novel strategy by which gut bacteria take advantage of catecholamines present in the gastrointestinal tract (*32, 71*) Understanding the interplay between pathogenic and commensal interactions with catecholamines is an intriguing avenue for further research.

In addition to characterizing Dadh, we discover candidate dehydroxylases that metabolize (+)-catechin, hydrocaffeic acid, and DOPAC. Further biochemical studies are important for validating the activities of these enzymes, but our preliminary data support a working model in which catechol dehydroxylation is performed by enzymes that are specialized for individual substrates. We identified large numbers of uncharacterized dehydroxylases in *Eggerthella* and *Gordonibacter* (Fig. 5B), hinting at an expansion of this group of enzymes among human gut Actinobacteria. While it remains to be seen whether these uncharacterized enzymes are also specific for distinct substrates, this type of diversification of closely related enzymes indicates a potentially important role for catechol dehydroxylation in the human gut microbiota. Expansion of enzyme families within specific clades of gut microbes is well-characterized in the context of polysaccharide metabolism. For example, human gut Bacteroides isolates harbor hundreds of polysaccharide utilization loci but upregulate only a subset of genes in response to distinct substrates (*72–75*). This transcriptional regulation and biochemical specificity enables utilization of host- or plant-derived carbon sources depending on their availability (*76–78*). The diversity of catechol dehydroxylases might have evolved in a similar manner, providing a biochemical arsenal that enables Actinobacteria to use a range of different electron acceptors whose availability depends on the diet or physiology of the host. Identifying the functions of uncharacterized catechol dehydroxylases could shed light the adaptation of gut organisms to small molecules produced and ingested by the host.

In addition to uncovering the diversity of dehydroxylases among human gut bacteria, our study illustrates the chemical strategies used to enable microbial survival and interactions in the human gut may be relevant to a broad range of species and habitats. While mammalian gut microbiomes have previously been compared in terms of gene content and species composition (*63–65*), our study provides functional evidence for conservation of specific gut microbial metabolic pathways across distinct hosts. While this hints at a potentially important role of catechol dehydroxylation among mammalian gut communities, the distribution of putative dehydroxylases among environmental microbes suggests the relevance of this chemistry likely extends to many additional microbial habitats. The shared metabolic capabilities between gut and environmental microbes reinforces findings from recent studies of gut microbial enzymes. For example, gut microbial carbohydrate-degrading enzymes and glycyl radical enzymes that play important roles in degrading diet-derived polysaccharides, amino acids, and osmolytes in the human gut can also be found among environmental isolates (*74, 79–81*). Enzyme discovery in the human gut microbiota therefore not only has implications for improving human health and disease, but also for discovering novel catalytic functions and metabolic pathways broadly relevant to microbial life. Our study now sets the stage for further investigations of the chemical mechanisms and biological consequences of catechol dehydroxylation in the human body and beyond.

Finally, our findings provide a framework for linking metabolic transformations performed by complex gut microbial communities to individual strains, genes, and enzymes. Our broad exploration of a class of metabolic transformations contrasts with the more common focus on metabolism of individual drugs or dietary compounds (*29, 6, 79–83*). This functional group-focused approach may greatly increase the efficiency with which we can link metabolic activities to microbial genes and enzymes. We envision that related experimental workflows could find broad utility in the discovery of gut microbial enzymes catalyzing other widespread, biologically significant reactions, including reductive metabolism of additional functional groups that are prevalent in diverse molecules encountered by the gut microbiota (*1*).

### General materials and methods

The following chemicals were used in this study: tetracycline (Sigma Aldrich, catalog# 87128-25G), *p*-tyramine (Sigma Aldrich, catalog# T2879-1G), DL-3,4-Dihydroxymandelic acid (Carbo Synth, catalog# FD22118), protocatechuic Acid (Millipore Sigma, catalog# 37580-25G-F), DL norepinephrine (Millipore Sigma, catalog# A7256-1G), L-norepinephrine (Matrix Scientific, catalog# 037592-500MG) L-epinephrine (Alfa Aesar, catalog# L04911.06), DL-epinephrine (Sigma Aldrich, catalog# E4642-5G), 3,4-dihydroxyphenylacetic acid (Millipore Sigma, catalog# 850217-1G), 3,4-dihydroxyhydrocinnamic acid (hydrocaffeic acid) (Millipore Sigma, catalog# 102601-10G), caffeic acid (Millipore Sigma, catalog# C0625-2G), (+)-catechin hydrate (Millimore Sigma, catalog# C1251-5G), (+/–)-catechin hydrate (Millipore Sigma, catalog# C1788-500MG), (−)-Epicatechin (Millipore Sigma, catalog# E1753-1G), L-(−)-a- Methyldopa (Chemcruz, catalog# sc-203092), ellagic acid (Millipore Sigma, catalog# E2250-1G), 2,3-dihydroxybenzoic acid (Millipore Sigma, catalog# 126209-5G), *R*-(−)-apomorphine hydrochloride hemihydrate (Sigma Aldrich, catalog# A4393-100MG), hydroxytyrosol (Ava Chem Scientific, catalog# 2528), enterobactin (generous gift from Prof. Elizabeth Nolan, MIT), fenoldopam mesylate (Sigma Aldrich, catalog# SML0198-10MG), 5-hydroxydopamine (Sigma Aldrich, catalog# 151564-100G), 6-hydroxydopamine (Sigma Aldrich, catalog #H4381-100MG), 3-methoxytyramine (Sigma Aldrich, catalog# M4251-100MG), 3,4-dihydroxybenzylamine (Sigma Aldrich, catalog# 858781-250MG), *N*-methyldopamine (Santa Cruz Biotechnology, catalog# sc-358430A), 4-(2-aminoethyl)benzene-1,3-diol (Enamine, catalog # EN300-65185), *m*-tyramine (Chemcruz, catalog# sc-255257), 3-hydroxyphenylacetic acid (Sigma Aldrich, catalog# H49901-5G), 3-hydroxyphenylpropionic acid (Toronto Research Chemicals, catalog# H940090), L-dopa (Oakwood Chemical, catalog# 358380-25g), dopamine (Sigma-Aldrich, catalog# PHR1090-1G, or Millipore Sigma, catalog# H8502-25G), *m*-tyramine (Santa Cruz Biotechnology, catalog# sc-255257), carbidopa (Sigma-Aldrich, catalog# PHR1655-1G), L-arginine (Sigma-Aldrich, catalog# A5006-100G), sodium molybdate (Sigma-Aldrich, catalog # 243655-100G), sodium tungstate (72069-25G), SIGMAFAST protease inhibitor tablets (Sigma-Aldrich, catalog#: S8830), benzyl viologen (Sigma-Aldrich, catalog# 271845-250mg), methyl viologen (Sigma-Aldrich, catalog# 856177-1g), diquat (Sigma-Aldrich, catalog# 45422-250mg), sodium dithionite (Sigma-Aldrich, catalog# 157953-5G). LC-MS grade acetonitrile and methanol for LC-MS analyses were purchased from Honeywell Burdick & Jackson or Sigma-Aldrich. Brain Heart Infusion (BHI) broth was purchased from Beckton Dickinson (catalog# 211060) or from VWR (catalog# 95021-488).

All bacterial culturing work was performed in an anaerobic chamber (Coy Laboratory Products) under an atmosphere of 10% hydrogen, 10% carbon dioxide, and nitrogen as the balance, unless otherwise noted. Hungate tubes were used for anaerobic culturing unless otherwise noted (Chemglass, catalog# CLS-4209-01). All lysate work and biochemical experiments were performed in an anaerobic chamber (Coy Laboratory Products) situated in a cold room at 4 °C under an atmosphere of 10% hydrogen and nitrogen as the balance. Gut Actinobacterial strains were grown on BHI containing 1% arginine (w/v) to obtain isolated colonies for culturing.

All genomic DNA (gDNA) was extracted from bacterial cultures using the DNeasy UltraClean Microbial Kit (Qiagen, catalog # 12224-50) according to the manufacturer’s protocol.

The remaining methods and data can be found in the supplementary materials.

## Acknowledgments

We acknowledge the Broad Institute Microbial Omics Core (MOC) for assistance with RNA sequencing analysis and experimental design, Dr. Rachel Carmody, Dr. Richard Losick, and Dr. Rachelle Gaudet (all at Harvard University) for helpful discussions. We thank Dr. Liz Ortiz (UC Irvine) for critical experimental feedback. We thank Dr. Elizabeth Nolan (MIT) for providing enterobactin. We thank Dr. Rachel Carmody and Dr. Aspen Reese for kindly providing the mammalian fecal samples.

## Author contributions

V.M.R, E.P.B, P.J.T, and E.N.B conceived of the project. V.M.R purified and characterized the dopamine dehydroxylase, performed all growth assays with catechols, performed RNA-sequencing experiments and analysis, performed lysate assays and cell resuspensions, performed LC-MS/MS assays and analysis, and performed phylogenetic analysis. P.N.B performed the systematic screen of gut Actinobacterial catechol metabolism. S.K performed culturing of mammalian fecal samples with catechol compounds. M.U.L designed and synthesized dopamine analogs for testing with the dopamine dehydroxylase. V.M.R, P.N.B, E.N.B, and M.U.L, provided critical feedback on experiments. V.M.R., P.N.B, E.N.B, P.J.T., and E.P.B wrote the manuscript.

## Data and materials availability

RNA-Seq data has been deposited into the Sequence Read Archive available by way of BioProject PRJNA557713 for *Eggerthella lenta* A2 and PRJNA557714 for *Gordonibacter pamelaeae* 3C.

