## Supplementary Materials for "A widely distributed metalloenzyme class enables gut microbial metabolism of host- and diet-derived catechols"

**Part 1:** Materials, methods, references, and data for all experiments except for synthesis and characterization of dopamine analogs (pages 2-70)

**Part 2:** Materials, methods, references and characterization data for synthesis of dopamine analogs (pages 71-131)

**Part 1**: Materials, methods, references, and data for all experiments except for synthesis and characterization of dopamine

**LC-MS methods**

Method A: Samples were analyzed using an Agilent technologies 6410 Triple Quad LC/MS and a Dikma Technologies Inspire Phenyl column (4.6 × 150 mm, 5 µm; catalog #81801). The flow rate was 0.5 mL $min^{-1}$ using 0.1% formic acid in water as mobile phase A and 0.1% formic acid in acetonitrile as mobile phase B. The column temperature was maintained at room temperature. The following gradient was applied: 0-2 min: 0% B isocratic, 2-9 min: 0-10% B, 9-11 min: 10-95% B, 11-15 min: 95% B isocratic, 15-18 min: 95-0% B, 18-21 min: 0% B isocratic. For mass spectrometry, the source temperature was 300 °C, and the masses of dopamine (precursor ion *m/z* = 154.3, daughter ion *m/z* = 137.3), and tyramine (precursor ion *m/z* = 138.3, daughter ion *m/z* = 121.3) were monitored at a collision energy of 15 mV and fragmentor setting of 135 in positive MRM mode.

Method B: Samples were analyzed using an Agilent technologies 6410 Triple Quad LC/MS and a Thermo Scientific Acclaim Polar Advantage II column (3 µM, 120A, 2.1*150 mm, product #: 063187). The flow rate was 0.2 mL $min^{-1}$ using 0.1% formic acid in water as mobile phase A and methanol as mobile phase B. The following gradient was applied: 0-4 min: 50% B isocratic, 4-7 min: 50-99%, 7-9 min: 99-50%, 9-13 min: 50% B isocratic. For mass spectrometry, the source temperature was 300 °C, and the masses of trihydroxydopamine (precursor ion *m/z* = 170.3, daughter ion *m/z* = 153.3), dopamine (precursor ion *m*/z = 154.3, daughter ion *m/z* = 137.3), phenylethylamine (precursor ion *m/z* = 122.3, daughter ion *m/z* = 105.2), and tyramine (precursor ion *m/z* = 138.3, daughter ion *m/z* = 121.3) were monitored at a collision energy of 15 mV and fragmentor setting of 135 in positive MRM mode.

Method C: Samples were analyzed using an Agilent technologies 6530 Accurate-Mass Q-TOF LC/MS and a Dikma Technologies Inspire Phenyl column (4.6 × 150 mm, 5 µm; catalog #81801). The flow rate was 0.4 mL $min^{-1}$ using 0.1% formic acid in water as mobile phase A and 0.1% formic acid in acetonitrile as mobile phase B. The column temperature was maintained at room temperature. The following gradient was applied: 0-2 min: 5% B isocratic, 2-25 min: 0-95% B, 25-30 min: 95% B isocratic, 30-40 min: 95-5% B. For the MS detection, the ESI mass spectra data were recorded in positive mode for a mass range of *m/z* 50 to 3000. A mass window of ± 0.005 Da was used to extract the ion of [M+H].

Method D: Samples were analyzed using an Agilent technologies 6530 Accurate-Mass Q-TOF LC/MS and a Dikma Technologies Inspire Phenyl column (4.6 × 150 mm, 5 µm; catalog #81801). The flow rate was 0.4 mL $min^{-1}$ using 0.1% formic acid in water as mobile phase A and 0.1% formic acid in acetonitrile as mobile phase B. The column temperature was maintained at room temperature. The following gradient was applied: 0-2 min: 5% B isocratic, 2-25 min: 0-95% B, 25-30 min: 95% B isocratic, 30-40 min: 95-5% B. For the MS detection, the ESI mass spectra data were recorded in negative mode for a mass range of *m/z* 50 to 3000. A mass window of ± 0.005 Da was used to extract the ion of [M+H].

Method E: Samples were analyzed using an Agilent technologies 6410 Triple Quad LC/MS and a Thermo Scientific Acclaim Polar Advantage II column (3 µM, 120A, 2.1*150 mm, product

#: 063187). The flow rate was 0.2 mL $min^{-1}$ using 0.1% formic acid in water as mobile phase A and methanol as mobile phase B. The following gradient was applied: 0-4 min: 50% B isocratic, 4-7 min: 50-99%, 7-9 min: 99-50%, 9-13 min: 50% B isocratic. For mass spectrometry, the source temperature was 300 °C, and the masses of catechin (precursor ion *m/z* = 289.2, daughter ion *m/z* = 109.1), benzyl ether reduced catechin (precursor ion *m/z* = 291.2, daughter ion *m/z* = 123.1), benzyl ether reduced, dehydroxylated catechin (precursor ion *m/z* = 275.2, daughter ion *m/z* = 107.1) were monitored at a collision energy of 15 mV and fragmentor setting of 135 in negative MRM mode.

Method F: Samples were analyzed using an Agilent technologies 6410 Triple Quad LC/MS and a Thermo Scientific Acclaim Polar Advantage II column (3 µM, 120A, 2.1*150 mm, product #: 063187). The flow rate was 0.2 mL $min^{-1}$ using 0.1% formic acid in water as mobile phase A and methanol as mobile phase B. The following gradient was applied: 0-4 min: 50% B isocratic, 4-7 min: 50-99%, 7-9 min: 99-50%, 9-13 min: 50% B isocratic. For mass spectrometry, the source temperature was 300 °C, and the masses of hydrocaffeic acid (precursor ion *m/z* = 181.2, daughter ion *m/z* = 137.2), hydroxyphenylpropionic acid (precursor ion *m/z* = 165.1, daughter ion *m/z* = 121.2), DOPAC (precursor ion m/z = 167.2, daughter ion m/z = 123.2), and hydroxyphenylacetic acid (precursor ion *m/z* = 151.3, daughter ion *m/z* = 107.3) were monitored at a collision energy of 15 mV and fragmentor setting of 135 in negative MRM mode.

Method G: Samples were analyzed using an Agilent technologies 6410 Triple Quad LC/MS and a Thermo Scientific Acclaim polar advantage II column (3 µM, 120A, 2.1*150 mm, product #: 063187). The flow rate was 0.2 mL $min^{-1}$ using 0.1% formic acid in water as mobile phase A and methanol as mobile phase B. The following gradient was applied: 0-4 min: 50% B isocratic, 4-7 min: 50-99%, 7-9 min: 99-50%, 9-13 min: 50% B isocratic. For mass spectrometry, the source temperature was 275 °C, and the masses of norepinephrine (precursor ion *m/z* = 170.1, daughter ion *m/z* = 152.1) and octopamine (precursor ion *m/z* = 154.2, daughter ion *m/z* = 136.1) were monitored at a collision energy of 5 mV and fragmentor setting of 135 in positive MRM mode.

Method H: Samples were analyzed using an Agilent technologies 6410 Triple Quad LC/MS and a Thermo Scientific Acclaim Polar Advantage II column (3 µM, 120A, 2.1*150 mm, product #: 063187). The flow rate was 0.2 mL $min^{-1}$ using 0.1% formic acid in water as mobile phase A and methanol as mobile phase B. The following gradient was applied: 0-4 min: 50% B isocratic, 4-7 min: 50-99%, 7-9 min: 99-50%, 9-13 min: 50% B isocratic. For mass spectrometry, the source temperature was 275 °C, and the masses of caffeic acid (precursor ion *m/z* = 179.2, daughter ion *m/z* = 135.2) and coumaric acid (precursor ion *m/z* = 163.3, daughter ion *m/z* =119.2) were monitored at a collision energy of 5 mV and fragmentor setting of 135 in negative MRM mode.

Method I: Samples were analyzed using an Agilent technologies 6410 Triple Quad LC/MS and a Thermo Scientific Acclaim Polar Advantage II column (3 µM, 120A, 2.1*150 mm, product #: 063187). The flow rate was 0.2 mL $min^{-1}$ using 0.1% formic acid in water as mobile phase A and methanol as mobile phase B. The following gradient was applied: 0-4 min: 50% B isocratic, 4-7 min: 50-99%, 7-9 min: 99-50%, 9-13 min: 50% B isocratic. For mass spectrometry, the source temperature was 300 °C, and the masses of dihydroxybenzoic acid (precursor ion *m/z* = 153.1, daughter ion *m/z* = 137.1) and hydroxybenzoic acid (precursor ion *m/z* = 137.1, daughter ion *m/z* = 93.2) were monitored at a collision energy of 15 mV and fragmentor setting of 135 in negative MRM mode.

Method J: Samples were analyzed using an Agilent technologies 6410 Triple Quad LC/MS and a Thermo Scientific Acclaim Polar Advantage II column (3 µM, 120A, 2.1*150 mm, product #: 063187). The flow rate was 0.2 mL $min^{-1}$ using 0.1% formic acid in water as mobile phase A and methanol as mobile phase B. The following gradient was applied: 0-4 min: 50% B isocratic, 4-7 min: 50-99%, 7-9 min: 99-50%, 9-13 min: 50% B isocratic. For mass spectrometry, the source temperature was 300 °C, and the masses of norepinephrine (precursor ion *m/z* = 184.1, daughter ion *m/z* = 166.1) and dehydroxynorepinephrine (precursor ion *m/z* = 168.1, daughter ion *m/z* = 150.1) were monitored at a collision energy of 15 mV and fragmentor setting of 135 in positive MRM mode.

Method K: Samples were analyzed using an Agilent technologies 6410 Triple Quad LC/MS and a Thermo Scientific Acclaim Polar Advantage II column (3 µM, 120A, 2.1*150 mm, product #: 063187). The flow rate was 0.2 mL $min^{-1}$ using 0.1% formic acid in water as mobile phase A and methanol as mobile phase B. The following gradient was applied: 0-4 min: 50% B isocratic, 4-7 min: 50-99%, 7-9 min: 99-50%, 9-13 min: 50% B isocratic. For mass spectrometry, the source temperature was 300 °C, and the masses of dihydroxybenzylamine (precursor ion *m/z* = 140.3, daughter ion *m/z* = 123.2) and hydroxybenzylamine (precursor ion *m/z* = 124.3, daughter ion *m/z* = 107.2) were monitored at a collision energy of 15 mV and fragmentor setting of 135 in positive MRM mode.

Method L: Samples were analyzed using an Agilent technologies 6410 Triple Quad LC/MS and a Thermo Scientific Acclaim Polar Advantage II column (3 µM, 120A, 2.1*150 mm, product #: 063187). The flow rate was 0.2 mL $min^{-1}$ using 0.1% formic acid in water as mobile phase A and methanol as mobile phase B. The following gradient was applied: 0-4 min: 50% B isocratic, 4-7 min: 50-99%, 7-9 min: 99-50%, 9-13 min: 50% B isocratic. For mass spectrometry, the source temperature was 300 °C, and the masses of 3-aminotyramine (precursor ion *m/z* = 153.3, daughter ion *m/z* = 136.2) and 3-aminophenylethylamine (precursor ion *m/z* = 137.3, daughter ion *m/z* = 120.2) were monitored at a collision energy of 15 mV and fragmentor setting of 135 in positive MRM mode.

Method M: Samples were analyzed using an Agilent technologies 6410 Triple Quad LC/MS and a Thermo Scientific Acclaim Polar Advantage II column (3 µM, 120A, 2.1*150 mm, product #: 063187). The flow rate was 0.2 mL $min^{-1}$ using 0.1% formic acid in water as mobile phase A and methanol as mobile phase B. The following gradient was applied: 0-4 min: 50% B isocratic, 4-7 min: 50-99%, 7-9 min: 99-50%, 9-13 min: 50% B isocratic. For mass spectrometry, the source temperature was 300 °C, and the masses 3-methoxytyramine (precursor ion *m/z* = 151.1, daughter ion *m/z* = 91.1) and 3-methoxyphenylethylamine (precursor ion *m/z* = 135.1, daughter ion *m/z* = 75.1) were monitored at a collision energy of 15 mV and fragmentor setting of 135 in positive MRM mode.

Method N: Samples were analyzed using an Agilent technologies 6410 Triple Quad LC/MS and a Thermo Scientific Acclaim Polar Advantage II column (3 µM, 120A, 2.1*150 mm, product #: 063187). The flow rate was 0.2 mL $min^{-1}$ using 0.1% formic acid in water as mobile phase A and methanol as mobile phase B. The following gradient was applied: 0-4 min: 50% B isocratic, 4-7 min: 50-99%, 7-9 min: 99-50%, 9-13 min: 50% B isocratic. For mass spectrometry, the source temperature was 300 °C, and the masses 3-hydroxytyrosol (precursor ion $m/z$ = 153.2, daughter ion $m/z$ = 123.1) and tyrosol (precursor ion $m/z$ = 137.2, daughter ion $m/z$ = 107.1) were monitored at a collision energy of 15 mV and fragmentor setting of 135 in negative MRM mode.

**Colorimetric assay for catechol detection**

The colorimetric assay for dopamine dehydroxylation was based on the Arnow test (*1*). Briefly, 50 µL of 0.5 M aqueous HCl was added to 50 µL of culture supernatant. After mixing, 50 µL of an aqueous solution containing both sodium molybdate and sodium nitrite (0.1 g/mL each) was added, which produced a yellow color. Finally, 50 µL of 1 M aqueous NaOH was added followed by pipetting up and down to mix. This allowed the characteristic pink color to develop. Absorbance was measured at 500 nm immediately using a Synergy HTX Multi-Mode Microplate Reader (BioTek) or SPECTROstar Nano (BMG LABTECH).

**Anaerobic activity-based purification of *E. lenta* A2 dopamine dehydroxylase**

Protein purification: Experiments were performed as described previously (*9*), with minor modifications. All procedures were carried out under strictly anaerobic conditions at 4 °C. Procedures outside the anaerobic chamber were performed in tightly sealed containers to prevent oxygen contamination. First, *E. lenta* A2 starter cultures were inoculated from single colonies into liquid BHI medium and were grown for 30 hours. Starter cultures were diluted 1:100 into 5 L of BHI medium containing 1% arginine and 10 mM formate and grown anaerobically at 37 °C for 16 hours. Dopamine was added as a solid to a final concentration of 0.5 mM in the cultures. Cells were pelleted in 5 separate 1 L bottles by centrifugation (6000 rpm, 15 mins), and each pellet was resuspended in 20 mL of 20 mM Tris pH 8 containing 4 mg/mL SIGMAFAST protease inhibitor cocktail. Resuspended cells were then lysed using two rounds of sonication in an anaerobic chamber (Branson Sonifier 450, 2 min total, 10 sec on, 40 sec off, 25% amplitude). The lysates were then clarified by centrifugation (10800 rpm, 15 mins), and the soluble fractions were subjected to two rounds of ammonium sulfate precipitation. During the precipitation, three different tubes each containing 40 mL total clarified lysate were precipitated in parallel. Solid ammonium sulfate was first dissolved in these clarified lysates to a final concentration of 30% (w/v), and lysates were left for 1 hour and 20 minutes followed by centrifugation to pellet the precipitates (4000 rpm, 15 mins). The supernatant was saved, and the pellet was discarded. The supernatant was mixed with additional solid ammonium sulfate to achieve a final concentration of 40% (w/v) and left for 1 hour and 20 minutes. Following centrifugation (4000 rpm, 15 mins) and removal of supernatant, each pellet containing the precipitated proteins was re-dissolved in 20 mL 20 mM Tris pH 8 containing 0.5 M ammonium sulfate. The re-dissolved pellets were combined and centrifuged to remove particulates (10800 rpm, 15 mins). The resulting 60 mL solution was injected onto an FPLC (Bio-Rad BioLogic DuoFlow System equipped with GE Life Sciences DynaLoop90) for hydrophobic interaction chromatography (HIC) using 5 x 1mL HiTrap Phenyl HP columns (GE Life Sciences, catalog# 17135101). Fractions were eluted with a gradient of 0.5 M to 0 M ammonium sulfate (in 20 mM Tris pH 8) at a flow rate of 1 mL/min and were tested for activity using the assay described below. The majority of the dopamine dehydroxylase activity eluted around 0.05 M-0.1 M ammonium sulfate. Active fractions displaying >50% conversion of dopamine were combined and injected onto the FPLC system described above for anion exchange chromatography using a UNO Q1 column (Bio-Rad, catalog# 720-0001) at a flow-rate of 1 mL/min. Fractions were eluted using a gradient of 0 to 1 M NaCl in 20 mM Tris pH 8 and were tested for activity. The majority of the dopamine dehydroxylase activity eluted around 250 mM NaCl. Active fractions were combined and concentrated 20-fold using a spin concentrator with a 5 kDa cutoff (4000 rpm centrifugation speed). 250 µL of the concentrate was injected onto FPLC for size exclusion chromatography using an Enrich 24 mL column (Enrich SEC 650, 10*300 column, Bio-Rad, catalog# 780-1650). Fractions were eluted over a 26 mL volume run isocratically in 20 mM Tris pH 8 containing 250 mM NaCl and were subjected to activity assays. Active fractions were then combined and used for enzyme assays and were run on SDS-PAGE to assess the presence of protein. Absorbance at 280 nm was used to determine the protein concentration, using a predicted extinction coefficient of 317735 $M^{-1}$ $cm^{-1}$ for the dopamine dehydroxylase. Activity assays during protein purification: 50 µL aliquots of fractions from FPLC runs were mixed, in the following order, with 1 µL electron donors (final concentration 1 mM each of methyl viologen, 1 mM diquat dibromide, 1 mM benzyl viologen, all dissolved in water), 2 µL sodium dithionite (2 mM final concentration, dissolved in water), and 1 µL substrate (500 µM final concentration, dissolved in water). The assay mixtures were left at room temperature in an anaerobic chamber for 12–14 hours to allow dopamine dehydroxylation to proceed, followed by assessment of activity using the colorimetric assay for catechol detection. Due to the inability of Dadh to survive freeze-thawing even in the presence of glycerol, the natively purified enzyme was always immediately used for enzyme assays.

**Assays of the *E. lenta* A2 dopamine dehydroxylase substrate scope.**

Active fractions from the size exclusion chromatography described above were combined and then diluted in 20 mM Tris pH 8 containing 250 mM NaCl to a final enzyme concentration of 0.1 µM. The enzyme mixture was transferred the wells of a 96 well plate, for a final volume of 50 µL in each well (VWR, catalog# 82006-636). 1 µL of substrate (in water, or 50:50 water:DMF for caffeic acid and catechin substrates) was then added at a final concentration of 500 µM. Following this, 1 µL of a solution containing electron donors (final concentration 1 mM each of methyl viologen, 1 mM diquat dibromide, 1 mM benzyl viologen, all dissolved in water) and 2 µL of sodium dithionite (2 mM final concentration, dissolved in water) were added. The resulting solution was mixed by pipetting and the 96-well plate was then sealed tightly with an aluminum seal. The enzyme assay mixtures were left at room temperature in an anaerobic chamber for 22 hours to allow dehydroxylation to proceed. The enzyme reaction mixtures were quenched by bringing the samples out of the anaerobic chamber and freezing at –20 ºC. These mixtures were then diluted 1:10 with LC-MS grade methanol and analyzed by LC-MS/MS. For the screen with physiologically relevant catechol substrates, samples containing caffeic acid were analyzed using Method H, hydrocaffeic acid and DOPAC were analyzed using Method F, catechin was analyzed using Method E, protocathecuic acid was analyzed using Method I, epinephrine was analyzed using Method J, norepinephrine was analyzed using Method G, and ellagic acid was analyzed using Method D. For the screen with dopamine analogs, all monohydroxylated, dihydroxylated, and trihydroxylated phenylethylamine analogs were analyzed using method B, *N*-methyldopamine was analyzed using Method C, methoxytyramine was analyzed using Method M, dihydroxybenzylamine was analyzed using Method K, hydroxytyrosol was analyzed using Method N, and aminotyramine was analyzed using Method L.

**Metabolism of dopamine analogs by *E. lenta* A2 cells**

Cells were cultured in 96-well plates and all experiments were performed anaerobically. The strains screened for dopamine dehydroxylation have been previously described (*2,3*). *E. lenta* A2 was inoculated from a single colony into 10 mL of BHI liquid medium and grown for 48 hours at 37 ºC to provide turbid starter cultures. These were diluted 1:10 in triplicate into 200 µL of fresh BHI medium containing 500 µM substrate (*p*-tyramine, dopamine, 3,4-dihydroxybenzylamine, or DL-norepinephrine). These cultures were grown for 48 hours at 37 ºC. Cultures were harvested by centrifugation at 4000 rpm for 10 minutes, and the supernatants were diluted 1:10 with LC-MS grade methanol. Samples containing dopamine or *p*-tyramine were analyzed using Method B, norepinephrine was analyzed using Method G, dihydroxybenzylamine was analyzed using Method K.

**RNA-sequencing experiments with *E. lenta* A2**

We repeated the setup previously used in the RNA-sequencing experiment with dopamine (*9*). Turbid 48-hour starter cultures of *E. lenta* in BHI medium were inoculated 1:100 into 5 mL of BHI medium containing 1% arginine and 10 mM formate, and cultures were grown at 37 °C anaerobically. When the cultures reached $OD_{600}$=0.200, hydrocaffeic acid, (+)-catechin, *p*-tyramine, 3,4-dihydroxybenzylamine, DL-norepinephrine, or *N*-methyldopamine were added at final concentrations of 500 µM to triplicate cultures. All compounds except for (+)-catechin were dissolved in water; (+)-catechin was dissolved in DMF. Control cultures contained vehicle (water or DMF). Cultures were harvested when they reached $OD_{600}$=0.500. They were centrifuged for 15 minutes at 4000 rpm, and cell pellets were re-suspended in 500 µL Trizol reagent (ThermoFisher, catalog#: 15596026). Total RNA was isolated by first bead beating to lyse cells and then using the Zymo Research Direct-Zol RNA MiniPrep Plus kit (Catalog # R2070) according to the manufacturer's protocol. Illumina cDNA libraries were generated using a modified version of the RNAtag-Seq protocol (*4*). Briefly, 500 ng of total RNA was fragmented, depleted of genomic DNA, and dephosphorylated prior to its ligation to DNA adapters carrying 5'-AN8-3' barcodes with a 5' phosphate and a 3' blocking group. Barcoded RNAs were pooled and depleted of rRNA using the RiboZero rRNA depletion kit (Epicentre). These pools of barcoded RNAs were converted to Illumina cDNA libraries in 3 main steps: (i) reverse transcription of the RNA using a primer designed to the constant region of the barcoded adaptor; (ii) addition of a second adapter on the 3' end of the cDNA during reverse transcription using SmartScribe RT (Clonetech) as described (*4*); (iii) PCR amplification using primers that target the constant regions of the 3' and 5' ligated adaptors and contain the full sequence of the Illumina sequencing adaptors. cDNA libraries were sequenced on Illumina HiSeq 2500. For the analysis of RNAtag-Seq data, reads from each sample in the pool were identified based on their associated barcode using custom scripts, and up to 1 mismatch in the barcode was allowed with the caveat that it did not enable assignment to more than one barcode. Barcode sequences were removed from the first read as were terminal G's from the second read that may have been added by SMARTScribe during template switching. Reads were aligned to the *Eggerthella lenta* A2 genome using BWA (*6*) and read counts were assigned to genes and other genomic features using custom scripts. Differential expression analysis was conducted with DESeq2 (*7*) and/or edgeR (*8*).

**RNA-sequencing experiments with *G. pamelaeae* 3C**

Method 1 (compound added at mid-exponential phase): Turbid 48-hour starter cultures of *G. pamelaeae* 3C grown in BHI medium were inoculated 1:100 into triplicate Hungate tubes containing 20 mL BHI medium with 10 mM formate. When cultures reached $OD_{600}$=0.110, DOPAC (0.5 mM final) or vehicle (water) was added to the cultures. The cultures were then grown at 37 °C anaerobically and harvested when they reached $OD_{600}$=0.185. They were centrifuged, and cell pellets were re-suspended in 500 µL Trizol reagent (ThermoFisher, catalog#: 15596026).

Method 2 (compound added at the beginning of growth): Turbid 48-hour starter cultures of *G. pamelaeae* 3C grown in BHI medium were inoculated 1:100 into triplicate hungate tubes containing 20 mL BHI with 10 mM formate and DOPAC (0.5 mM final) or vehicle (water). These cultures were then left to grow at 37 °C anaerobically. When cultures reached $OD_{600}$=0.110, they were harvested. They were centrifuged, and cell pellets were re-suspended in 500 µL Trizol reagent (ThermoFisher, catalog#: 15596026).

RNA extraction and sequencing: this was performed using the exactly same setup as described above, except the reads were aligned to the genome of *Gordonibacter pamelaeae* 3C.

**Growth of *E. lenta* A2 in BHI with and without dopamine**

Cells were cultured in Hungate tubes and all experiments were performed anaerobically. *E. lenta* A2 was inoculated from a single colony into 10 mL BHI liquid medium and grown for 48 hours at 37 ºC to provide turbid starter cultures. These were diluted 1:100 in triplicate into 5 mL BHI medium containing either 0.5 mM dopamine or vehicle. Growth was assessed by measuring the optical density at 600 nm using a Genesys 20 spectrophotometer (Thermo Scientific).

**Preparation of basal medium lacking electron acceptors**

The medium was prepared as described previously, with minor modifications (*9*). A 100-fold stock solution of salts was first prepared by dissolving 100 g NaCl, 50 g $MgCl_2 \cdot 6H_2O$, 20 g $KH_2PO_4$, 30 g $NH_4Cl$, 30 g KCl, 1.5 g $CaCl_2$ x $2H_2O$ in 1 L of water. Then, 10 mL of this solution was added to 1 L of water containing 1 g yeast extract (Beckton Dickinson #288260), 1 g tryptone (Beckton Dickinson #21175), and 0.25 mL of 0.1% resazurin (dissolved in MilliQ water). This medium was autoclaved. Following autoclaving, the medium was left to cool for 15 minutes in an atmosphere of air (outside the anaerobic chamber). After cooling, the following components were added using sterile technique: 10 mL of ATCC Trace element mix (ATCC, catalog# MD-TMS), 10 mL of Vitamin Supplement (ATCC, catalog# MD-VS), solid $NaHCO_3$ (SIGMA, 2.52 g, to give 30 mM) and solid L-cysteine HCl (SIGMA, 63 mg, to give 0.4 mM). The medium had a final pH of 7.2-7.3. The medium was then sparged with nitrogen gas (for how long) and was brought into the anaerobic chamber to equilibrate for at least 30 hours prior to use. In all experiments utilizing the basal medium, except for those experiments performed with *Gordonibacter pamelaeae* 3C or the screen for catechol metabolism by mammalian gut microbiota samples, sodium acetate was added at a final concentration of 10 mM at the time of bacterial inoculation. In experiments performed with *Gordonibacter pamelaeae* 3C, sodium formate was added at a final concentration of 10 mM. In the *ex vivo* experiments with the mammalian gut microbiota, neither acetate nor formate were added to the basal medium.

**Growth of single *E. lenta* strains in basal medium**

Cells were cultured in hungate tubes and all experiments were performed anaerobically. *E. lenta* strains were inoculated from single colonies into 10 mL of BHI liquid medium and grown for 48-72 hours at 37 ºC to provide turbid starter cultures. These were diluted 1:100 in triplicate into 5 mL of basal medium containing 10 mM acetate and either 1 mM dopamine (in water) or vehicle (water). If applicable. molybdate (0.5 mM), tungstate (0.5 mM), DMSO (14 mM), or nitrate (1 mM) were added at the time of inoculation. Cultures were grown anaerobically for 36-72 hours at 37 ºC. Endpoint growth was assessed by measuring the optical density at 600 nm using a Genesys 20 spectrophotometer (Thermo Scientific). Catechol dehydroxylation was assessed at the end of growth in culture supernatants using the colorimetric method.

**Competition of *E. lenta* strains in basal medium**

Cells were cultured in hungate tubes and all experiments were performed anaerobically. *E. lenta* strains W1BHI6 (Tet resistant non-metabolizer, and Valencia (Tet sensitive metabolizer) were inoculated from single colonies into individual tubes containing 10 mL of BHI liquid medium and grown for 48 hours at 37 ºC to provide turbid starter cultures. For the competition experiment, 50 µL of each starter culture of the two competing strains was combined in triplicate in 5 mL of basal medium containing 10 mM acetate and either 1 mM dopamine or vehicle (water). Following inoculation, cultures were grown anaerobically for 72 hours at 37 ºC. At the end of the incubation, growth of *E. lenta* was assessed. Cultures were serially diluted in PBS under anaerobic conditions, and 8 µL of each serial dilution ($10^{-1}$ through $10^{-7}$) was plated onto BHI plates containing 1% arginine (w/v) with and without 10 µg/mL Tetracycline using a spot plating method. Plates were grown at 37 ºC for 72 hours following by counting of colonies. To calculate the proportion of metabolizer in the W1BHI6/Valencia competition experiment, we selected a dilution where distinct colonies were clearly visible ($10^{-4}$-$10^{-5}$) and counted the number of colonies growing on the BHI 1% arginine Tetracycline plates (W1BHI6) as well as the colonies growing on the BHI 1% arginine plates (Both Valencia and W1BHI6). To get the number of metabolizer (Valencia) colonies, we subtracted the number of Tetraycline resistant colonies from the colonies on the no Tetracycline plate.

**Growth of *E. lenta* strains in the presence of a defined community**

Cells were cultured in hungate tubes and all experiments were performed anaerobically. *E. lenta* strains, as well as *Enterococcus faecalis* OGR1F, *Escherichia coli* MG1655, *Bacteroides fragilis* ATCC 25285, *Clostridium sporogenes* ATCC 15579, *Edwarsiella tarda* ATCC 23685, were inoculated from single colonies into individual tubes containing 10 mL of BHI liquid medium and grown for 48-72 hours at 37 ºC to provide turbid starter cultures. Growth was assessed by measuring the optical density at 600 nm using a Genesys 20 spectrophotometer (Thermo Scientifc). These starter cultures were then diluted to a final $OD_{600}$ of 0.100 in BHI medium anaerobically. The defined community was created by combining equal volumes of all strains (after diluting each culture to $OD_{600}$=0.100) except for *E. lenta*. The community was then inoculated 1:100 in triplicate into 5 mL basal medium containing 10 mM acetate and either 1 mM dopamine or vehicle. *E. lenta* strains were then added by diluting the *E. lenta* starter cultures (normalized to $OD_{600}$ = 0.100) 1:50 into the tubes containing the defined community. Cultures were then grown anaerobically for 72 hours at 37 ºC. At the end of the incubation, growth of *E. lenta* was assessed. Cultures were serially diluted in PBS under anaerobic conditions, and 8 µL of each serial dilution ($10^{-1}$ through $10^{-7}$) was plated onto BHI plates containing 1% arginine (w/v) and 10 µg/mL Tetracycline (spot plating method). Plates were grown at 37 ºC for 72 hours following by counting of colonies.

**Human fecal samples used in this study**

The human fecal samples used in this study have been previously described (*9*). To prepare them for culturing, all samples were resuspended anaerobically in anaerobic PBS at a final concentration of 0.1 g/mL. The mixture was vortexed to produce a homogenous slurry and was then left for 30 minutes to let particulates settle. Aliquots of the supernatant were dissolved 50:50 with 40% glycerol and flash-frozen in liquid nitrogen, creating slurries that were used for anaerobic culturing of human fecal samples. Slurries were stored at –80 ºC and were thawed anaerobically at room temperature at the time of use.

**Growth of human fecal samples in basal medium with dopamine**

Fecal slurries from n=24 unrelated humans were diluted 1:100 into two different hungate tubes containing 5 mL of basal medium with 10 mM acetate and either 1 mM dopamine or vehicle (water). These fecal microbiota cultures were grown anaerobically for 72 hours at 37 ºC. Metabolism was then assessed in culture supernatants using the colorimetric method. In addition, cultures were spun down and the total community gDNA was extracted from the entire 5 mL of culture for downstream PCR and qPCR assays as detailed below.

**qPCR assays for *E. lenta* and *dadh* abundance in human fecal samples grown in basal medium with and without dopamine**

Assays were performed as previously described (*9*). gDNA was extracted from the culture pellets generated in the experiments described above ("Growth of fecal samples in basal medium with dopamine") using the DNeasy UltraClean Microbial Kit. The extracted DNA from each culture was used for qPCR assays containing 10 µL of iTaq Universal SYBRgreen Supermix (Bio-rad, catalog 3: 1725121), 7 µL of water, and 10 µM each of forward and reverse primers. PCR was performed on a CFX96 Thermocycler (Bio-Rad), using the following program: initial denaturation at 95 ºC for 5 minutes 34 cycles of 95 ºC for 1 min, 60 ºC for 1 min, 72 ºC for 1 min. The program ended with a final extension at 34 ºC for 5 mins. The primers used were: 16S primers for *E. lenta* (*10*): CAGCAGGGAAGAAATTCGAC and TTGAGCCCTCGGATTAGAGA; primers for dopamine dehydroxylase: GAGATCTGGTCCACCGTCAT and AGTGGAAGTACACCGGGATG (*9*).

**Amplification of full-length *dadh* and sequencing of the SNP at position 506 from human fecal samples grown in basal medium with and without dopamine**

gDNA was extracted from the culture pellets generated in the experiments described above ("Growth of fecal samples in basal medium with dopamine") using the DNeasy UltraClean Microbial Kit. The extracted DNA from each culture was used for PCR assays containing 10 µL of Phusion High-Fidelity PCR Master mix with HF buffer (NEB, catalog# M0531L), 7 µL of water, and 10 µM each of forward and reverse primers. The primers used to amplify the full-length dopamine dehydroxylase from these samples were ATGGGTAACCTGACCATG and TTACTCCCTCCCTTCGTA. PCR was performed on a C1000 Touch Thermocycler (Bio-Rad), using the following program: initial denaturation at 98 ºC for 30 s, 34 cycles of 98 ºC for 10 s, 61 ºC for 15 s, 72 ºC for 2.5 mins. The program ended with a final extension at 72 ºC for 5 mins. Amplicons were purified using the Illustra GFX PCR DNA and Gel Band Purification Kit (GE Healthcare, catalog# 28-9034-70) and were sequenced using Sanger sequencing (Eton Biosciences) for the region containing the SNP at position 506 using primers GGGGTGTCCATGTTGCCGGT and ACCGGCTACGGCAACGGC. Sequence chromatograms were analyzed in Ape Plasmid Editor (version 2.0.47), and the single nucleotide polymorphism (SNP) at position 506 was called by visual inspection compared to results obtained from control cultures of *E. lenta* strains.

**Screen of gut Actinobacteria for metabolism of catechols**

This procedure was performed in an anaerobic chamber (Coy Laboratory Products, atmospheric conditions: 20% $CO_2$, 2-2.5% $H_2$, and the balance $N_2$)—equilibrating media and consumables to the atmosphere prior to use—until centrifugation, which was performed using a benchtop centrifuge. The 96-well plates used in this experiment were purchased from VWR (catalog# 10861-562). Into the wells of flat-bottom 96-well plates, 100 µL of BHI medium supplemented with L-cysteine-HCl (0.05%, w/v), *L*-arginine (1%, w/v), and sodium formate (10 mM) (referred to here as BHI++) were aliquoted. Seed cultures were prepared by inoculating wells, in triplicate, with Actinobacterial strains that were cultured on BHI++ agar plates. Additional wells served as sterile controls. Plates were sealed with tape and incubated at 37 °C for 12 to 18 hours to afford dense cultures. Next, 99 µL of BHI++ medium containing 500 µM of compound were aliquoted into the wells of a 96-well plate. To these wells, 1 µL of dense seed culture (or sterile control) was added. Plates were sealed and incubated at 37 °C for 24 or 48 hours. Plates were then centrifuged at 2000 rpm for 10 min at 4 °C, and the supernatant was aspirated and transferred to a fresh 96-well plate. An aliquot (35 µL) of supernatant was then immediately screened via the catechol colorimetric assay (described above in "Colorimetric assay for catechol detection"). Absorbance was immediately measured at 500 nm using a plate reader (Spectrostar Nano, BMG LABTECH). A standard curve (2-fold serial dilutions, 1000-15.6 µM in BHI++) was simultaneously prepared, developed, and analyzed using the conditions listed above. The catechol concentrations in bacterial cultures were normalized to the sterile control. To confirm metabolism of (+)-catechin, DOPAC, and hydrocaffeic acid, the incubations were repeated following the same procedure with minor modifications. Strains were grown in BHI for 48 hours anaerobically at 37 °C. Cultures were harvested by centrifugation and were then analyzed by LC-MS. To prepare samples for LC-MS, 20 µL of the culture supernatant was diluted 1:10 with 180 µL of methanol, followed by centrifugation at 4000 rpm for 10 minutes to pellet particulates, salts, and proteins. 50 µL of the resulting supernatant was then transferred to a 96-well plate and 5 µL of the supernatant was injected onto the instrument using Method E for catechin and Method F for hydrocaffeic acid and DOPAC. The following stock solutions were used in the screens: dihydroxymandelic acid (50 mM in water), dopamine (50 mM in water), protocatechuic acid (50 mM in ethanol), L-dopa (50 mM in 0.5 M HCl), norepinephrine (50 mM in 0.5 M HCl), epinephrine (50 mM in 0.5 M HCl), DOPAC (50 mM in 0.5 M HCl), Hydrocaffeic acid (50 mM in ethanol), caffeic acid (50 mM in ethanol), (+)-catechin (50 mM in ethanol), (+/–)-catechin (50 mM in ethanol), (–)-epicatechin (50 mM in DMSO), methyldopa (solid dissolved directly into the media at 0.5 mM final concentration), carbidopa (solid dissolved directly into the media at 0.5 mM final concentration), dihydroxybenzoic acid (50 mM in methanol). Hydroxytyrosol (50 mM in water), enterobactin (10 mM in DMSO), apomorphine (50 mM in DMSO).

**Confirmation of dehydroxylation of (+)-catechin and hydrocaffeic acid by *E. lenta* A2**

Cells were cultured in hungate tubes and all experiments were performed anaerobically. *E. lenta* A2 was inoculated from a single colony into 10 mL BHI liquid medium and grown for 48 hours at 37 ºC to provide turbid starter cultures. These were diluted 1:100 in triplicate into 5 mL of BHI medium containing either 0.5 mM hydrocaffeic acid (in water), 0.5 mM (+)-catechin (in

DMF), or vehicle (water or DMF). After 48 hours of anaerobic growth at 37 ºC, cultures were harvested by centrifugation and were then analyzed by LC-MS. To prepare samples for LC-MS, 20 µL of the culture supernatant was diluted 1:10 with 180 µL of methanol, followed by centrifugation at 4000 rpm for 10 minutes to pellet particulates, salts, and proteins. 50 µL of the resulting supernatant was then transferred to a 96-well plate and 5 µL of the supernatant was injected onto the instrument using Method E for catechin and Method F for hydrocaffeic acid.

**Confirmation of dehydroxylation of DOPAC by *Gordonibacter pamelaeae* 3C**

Cells were cultured in hungate tubes and all experiments were performed anaerobically. *G. pamelaeae* 3C was inoculated from a single colony into 10 mL of BHI liquid medium and grown for 48 hours at 37 ºC to provide turbid starter cultures. These were diluted 1:100 in triplicate into 5 mL of BHI medium containing 10 mM formate and 0.5 mM DOPAC or vehicle (water). After 72 hours of anaerobic growth at 37 ºC, the cultures were harvested by centrifugation and were then analyzed by LC-MS. To prepare samples for LC-MS, 20 µL of the culture supernatant was diluted 1:10 with 180 µL of methanol, followed by centrifugation at 4000 rpm for 10 minutes to pellet particulates, salts, and proteins. 50 µL of the resulting supernatant was then transferred to the LC-MS 96-well plate and 5 µL of the supernatant was injected onto the instrument using Method F for DOPAC.

**PBS resuspension assays for inducibility and oxygen sensitivity of catechol dehydroxylases**

Assays with *E. lenta* A2: Cells were cultured in hungate tubes and all experiments were performed anaerobically. *E. lenta* A2 was inoculated from a single colony into 10 mL of BHI liquid medium and grown for 48 hours at 37 ºC to provide turbid starter cultures. These were diluted 1:100 in triplicate into 10 mL of BHI medium containing 1% arginine and 10 mM formate and either 0.5 mM dopamine (in water), 0.5 mM hydrocaffeic acid (in water), 0.5 mM (+)-catechin (in DMF), or vehicle (water or DMF). After 18 hours of anaerobic growth at 37 ºC, cultures had reached an $OD_{600}$ of 0.700 and were harvested by centrifugation (4000 rpm, 15 minutes). The bacterial pellets were resuspended anaerobically in 10 mL of pre-reduced PBS to wash the cells, followed by an additional round of centrifugation to pellet the washed cells (4000 rpm, 15 minutes). The cells were then resuspended in 5 mL of pre-reduced PBS. 0.1 mL aliquots of this resuspension was transferred to Eppendorf tubes containing either vehicle or 0.5 mM catechol substrate. The samples were vortexed briefly and incubated anaerobically at room temperature for 20 hours to allow for metabolism to proceed. To assess the impact of oxygen on the metabolism of catechols, 0.1 mL of the PBS resuspension in an Eppendorf tube was brought outside the anaerobic chamber, followed by addition of 0.5 mM substrate in the presence of atmospheric oxygen. The samples were vortexed briefly and were incubated at room temperature for 20 hours. Samples were then analyzed by LC-MS. To prepare samples for LC-MS, 20 µL of the culture supernatant was diluted 1:10 with 180 µL of methanol, followed by centrifugation at 4000 rpm for 10 minutes to pellet particulates, salts, and proteins. 50 µL of the resulting supernatant was then transferred to the LC-MS 96-well plate and 5 µL of the supernatant was injected onto the instrument using Method B for dopamine, Method E for catechin and Method F for hydrocaffeic acid. Assays with *Gordonibacter pamelaeae* 3C: Cells were cultured in hungate tubes and all experiments were performed anaerobically. *G. pamelaeae* 3C was inoculated from a single colony into 10 mL of BHI liquid medium and grown for 48 hours at 37 ºC to provide turbid starter cultures. These were diluted 1:100 in triplicate into 10 mL of BHI medium containing 10 mM formate and either 0.5 mM DOPAC or vehicle. After 18 hours of anaerobic growth at 37 ºC, cultures had reached an $OD_{600}$ of 0.180 and were harvested by centrifugation (4000 rpm, 15 minutes). The bacterial pellets were resuspended anaerobically in 10 mL of pre-reduced PBS to wash the cells, followed by an additional round of centrifugation to pellet the washed cells. The cells were then resuspended in 5 mL of pre-reduced PBS. 0.1 mL aliquots of this resuspension were transferred to Eppendorf tubes containing either vehicle or 0.5 mM catechol substrate. The samples were vortexed briefly and incubated anaerobically at room temperature for 20 hours to allow for metabolism to proceed. To assess the impact of oxygen on DOPAC metabolism, 0.1 mL of the PBS resuspension in an Eppendorf tube was brought outside the anaerobic chamber, followed by addition of 0.5 mM substrate in the presence of atmospheric oxygen. The samples were vortexed briefly and were incubated at room temperature for 20 hours. Samples were then analyzed by LC-MS. To prepare samples for LC-MS, 20 µL of the culture supernatant was diluted 1:10 with 180 µL of methanol, followed by centrifugation at 4000 rpm for 10 minutes to pellet particulates, salts, and proteins. 50 µL of the resulting supernatant was then transferred to the LC-MS 96-well plate and 5 µL of the supernatant was injected onto the instrument using Method F.

**Effect of tungstate on growth and catechol dehydroxylation by *E. lenta* A2 and *G. pamelaeae* 3C**

Assays with *E. lenta* A2: Starter cultures of *E. lenta* A2 were grown over 48 hours in 10 mL of BHI medium and then inoculated 1:100 into 200 µL of BHI medium containing either 500 µM dopamine (in water), 500 µM (+)-catechin (in DMF), or 500 µM hydrocaffeic acid (in water), and either sodium tungstate (0.5 mM, in water), sodium molybdate (0.5 mM, in water), or vehicle (water or DMF). Cultures were grown for 48 hours anaerobically at 37 °C and were harvested by centrifugation. Supernatants were dissolved 1:10 in LC-MS grade methanol and analyzed using LC-MS/MS Methods B, E, or F described above. Experiments were performed anaerobically, and cultures were grown in 96-well plates (VWR, catalog# 29442-054). Assays with *Gordonibacter pamelaeae* 3C: Starter cultures of *G. pamelaeae* 3C were grown over 48 hours in 10 mL of BHI medium and then inoculated 1:100 into 200 µL of BHI medium containing 500 µM DOPAC and either sodium tungstate (0.5 mM, in water), sodium molybdate (0.5 mM, in water), or vehicle (water). Cultures were grown for 48 hours anaerobically at 37 °C and were harvested by centrifugation. Supernatants were dissolved 1:10 in LC-MS grade methanol and analyzed using LC-MS/MS Method F described above. Experiments were performed anaerobically, and cultures were grown in 96-well plates (VWR, catalog# 29442-054).

**Lysate assays for transcriptional and biochemical specificity of dehydroxylases from *G. pamelaeae* 3C and *E. lenta* A2**

Assays in *E. lenta* A2: Bacterial cultures were grown in hungate tubes. All bacterial growth and lysate experiments were performed in an anaerobic chamber. Lysis and sample processing took place in an anaerobic chamber kept at 4 ºC. *E. lenta* A2 was inoculated from a single colony into 10 mL of BHI liquid medium and grown for 48 hours at 37 ºC to provide turbid starter cultures. These were diluted 1:100 in triplicate into 50 mL of BHI medium containing 1% arginine and 10 mM formate and either 1 mM dopamine, 1 mM hydrocaffeic acid, 1 mM (+)-catechin, or vehicle. After 18 hours of anaerobic growth at 37 ºC, cultures had reached $OD_{600}$ of 0.700 and were harvested by centrifugation. The bacterial pellets were resuspended anaerobically in 10 mL of cold, pre-reduced PBS to wash the cells, followed by an additional round of centrifugation to pellet the washed cells. The washed cells from each culture were then transferred to an Eppendorf tube and resuspended in 1.4 mL of lysis buffer (20 mM Tris pH 8 containing 4 mg/mL SIGMAFAST protease inhibitor cocktail). The cells were lysed using sonication in an anaerobic chamber. 50 µL of this lysate was transferred in triplicate to a 96 well plate (VWR, catalog# 82006-636). 1 µL of substrate was then added to each of the replicates at a final concentration of 0.5 mM. These samples were incubated anaerobically at room temperature for 28 hours to allow for metabolism to proceed. Samples were then analyzed by LC-MS. To prepare samples for LC-MS, 20 µL of the culture supernatant was diluted 1:10 with 180 µL of methanol, followed by centrifugation at 4000 rpm for 10 minutes to pellet particulates, salts, and proteins. 50 µL of the resulting supernatant was then transferred to the LC-MS 96-well plate and 5 µL of the supernatant was injected onto the instrument using method B for dopamine, method E for catechin and method F for hydrocaffeic acid. Assays in *Gordonibacter pamelaeae* 3C: Bacterial cultures were grown in hungate tubes. All bacterial growth and lysate experiments were performed in an anaerobic chamber. Lysis and sample processing took place in an anaerobic chamber kept at 4 ºC. *G. pamelaeae* 3C was inoculated from a single colony into 10 mL of BHI liquid medium and grown for 48 hours at 37 ºC to provide turbid starter cultures. These were diluted 1:100 in triplicate into 50 mL of BHI medium containing 10 mM formate and 1 mM DOPAC or vehicle. After 18 hours of anaerobic growth at 37 ºC, cultures had reached $OD_{600}$ of 0.180 and were harvested by centrifugation. The bacterial pellets were resuspended anaerobically in 10 mL of cold, pre-reduced PBS to wash the cells, followed by an additional round of centrifugation to pellet the washed cells. The washed cells from each culture were then transferred to an Eppendorf tube and resuspended in 1.4 mL of lysis buffer (20 mM Tris pH 8 containing 4 mg/mL SIGMAFAST protease inhibitor cocktail). The cells were lysed using sonication in an anaerobic chamber. 50 µL of this lysate was transferred in triplicate to a 96 well plate (VWR, catalog# 82006-636). 1 µL substrate was then added to each of the replicates, for a final concentration of 0.5 mM. These samples were left anaerobically at room temperature for 28 hours to allow for metabolism to proceed. Samples were then analyzed by LC-MS. To prepare samples for LC-MS, 20 µL of the culture supernatant was diluted 1:10 with 180 µL of methanol, followed by centrifugation at 4000 rpm for 10 minutes to pellet particulates, salts, and proteins. 50 µL of the resulting supernatant was then transferred to the LC-MS 96-well plate and 5 µL of the supernatant was injected onto the instrument using LC-MS/MS Method F described above.

**Comparative genomics among human gut Actinobacteria**

To characterize the distribution of Cadh, Hcdh, Dodh among our gut Actinobacterial strain library, we performed a tBLASTn search. We queried the genomes for Cadh, Hcdh, Dodh and used 90% coverage, 75% amino acid identity, and e-value=0 as the cutoff for assessing the presence of each dehydroxylase.

**Phylogenetic analysis of relationship between catechol dehydroxylases and other characterized members of the bis-molybdopterin guanine dinucleotide enzyme family**

For phylogenetic analysis of the bis-MGD enzymes, we gathered sequences that have been previously used to study the evolution of bis-MGD enzymes (*14*). However, we also added sequences to capture additional diversity of biochemically characterized bis-MGD enzymes that were not included in the original tree described in (*14*). In particular, we performed a pBLAST search in Uniprot using perchlorate reductase (Uniprot ID# PCRA_DECAR), ethylbenzene dehydrogenase (Uniprot ID# Q5NZV2_AROAE), acetylene hydratase (Uniprot ID# AHY_PELAE), and pyrogallol transhydroxylase (Uniprot ID# PGTL_PELAC) as the queries, and collected sequences with 85-90% amino acid ID. In addition, we added the sequences of Dadh, Hcdh, Cadh from *E. lenta* A2, and Dodh and Cadh from G. *pamelaeae* 3C. The sequences were combined with those reported in (*14*) and were aligned in Geneious (version 11) using MUSCLE. We subsequently used FastTree (standard settings, 20 rate categories of sites) to create a maximum likelihood tree. The tree files were uploaded to the Interactive Tree of Life web server (https://itol.embl.de/) to annotate the trees (*15*).

**Construction of sequence similarity network of the bis-molybdopterin guanine dinucleotide enzyme family.**

A SSN was generated using the EFI-EST tool (http://efi.igb.illinois.edu/efi-est/) on July 15 2017 (*16*). In particular, we generated an SSN of the molybdopterin dinucleotide binding domain enzyme superfamily (PF01568), including sequences between 700 and 1400 amino acids in length and using an initial alignment score of e-150. Nodes represented sequences with 75% amino acid identity. The SSN was imported into Cytoscope v 3.2.1 and visualized with the 'Organic layout' setting. The alignment score cutoff was increased to e-167, until enzymes with known functions separated into putatively isofunctional clusters.

**Phylogenetic analysis of catechol dehydroxylases encoded by gut Actinobacteria and environmental isolates**

To identify additional diversity beyond the newly identified putative dehydroxylases from this study, we created a database containing putative homologs from a collection of 26 previously sequenced Actinobacterial genomes (*3*), as well as from genomes publicly available through NCBI. First, the *Eggerthella lenta* A2 dopamine dehydroxylase (Dadh) protein sequence was used as the query sequence for a tBLASTn search of 26 previously sequenced Actinobacterial genomes (*2,3*) (April 23, 2019). The genomes were loaded in Geneious (version 11) and hits with an amino acid ID of >30% and e-value of e-34 were considered potential dehydroxylase hits and were saved. This cutoff was chosen because sequences captured within this window were more closely related to the acetylene hydratase and Dadh than to any other biochemically characterized moco enzyme, as assessed by percent amino acid identity. In addition, we used the representative *Gordonibacter* enzyme Cldh as a separate query to identify the more distantly related, smaller enzymes from *Gordonibacter* that were not detected when using the large, multi-subunit Dadh as the query. Specifically, we used tBLASTn to search the 26 Actinobacterial genomes for the *Gordonibacter pamelaeae* 3C Cldh protein sequence. Hits from *Paraeggerthella hongongensis, Gordonibacter pamelaeae* 3C and *Gordonibacter sp.* 28C, the only organisms containing these smaller Cldh-like enzymes in our collection, were saved. Again, amino acid ID of >30% and e-value of e-34 were considered potential hits because sequences captured within this window were more closely related to the acetylene hydratase and Dadh than to any other biochemically characterized moco enzyme, as assessed by percent amino acid identity. The hits from our searches with Cldh and Dadh were combined into a preliminary database in Geneious. To expand the sequence diversity within this database, we used Cldh and Dadh as queries for two separate tBLASTn searches in NCBI (nucleotide collection). To ensure that we captured diversity beyond human gut microbes, we excluded *Gordonibacter* and *Eggerthella* as organisms in the tBLASTn searches for Cldh and Dadh queries, respectively. For the two searches, sequences of >29% amino acid ID and e-value of e-55 were considered potential dehydroxylase hits. This was a more conservative cutoff than we used with human Actinobacteria and was selected based on the observation that the pBLAST alignment of Dadh and Cldh has an e value of e-45 and 29% amino acid ID. The sequences retrieved from NCBI were added to the database already containing the hits from searches of the 26 Actinobacterial genomes. In addition, we added the biochemically characterized *E. coli* bis-MGD enzyme DMSO reductase (DmsA, Uniprot ID#P18775) to this database as the outgroup. This sequence was also used as the root of the tree. For phylogenetic analysis of these sequences, we first aligned sequences in Geneious using MUSCLE and removed sequences that were 95% identical to each other (considered duplicates). After deleting these duplicate sequences, we re-aligned the sequences using MUSCLE (standard settings) and subsequently used FastTree (standard settings, 20 rate categories of sites) to create a maximum likelihood tree. The tree files were uploaded to the Interactive Tree of Life web server (https://itol.embl.de/) to annotate the trees (*15*).

**Phylogenetic analysis of relationship between representative dehydroxylase homologs (from figure S17 and table S14) and other characterized members of the bis-molybdopterin guanine dinucleotide enzyme family**

Once we had constructed the two trees described above (Fig 5 in the main text) and uncovered dehydroxylase homologs in gut and environmental bacteria, we wanted to explore the phylogenetic relationship between these enzymes and the broader bis-molybdopterin guanine dinucleotide (bis-MGD) enzyme family. To do this, we added the representative sequences from fig. S17 and table S14 to the sequence database already described in *"Phylogenetic analysis of relationship between catechol dehydroxylases and other characterized members of the bis-molybdopterin guanine dinucleotide enzyme family"* here in methods. Using MUSCLE, we aligned the newly added sequences with the sequences represented on the tree (Fig 6 in main text). We then used FastTree (standard settings, 20 rate categories of sites) in Geneious (version 11) to generate the tree seen in fig. S24. The tree files were uploaded to the Interactive Tree of Life web server (https://itol.embl.de/) to annotate the trees (*15*).

**Mammalian fecal samples used in this study**

The collection of fecal samples from mammals (n=12 different species, n=3 individuals per species) has been previously described (*12*). To prepare these samples for culturing, all samples were resuspended anaerobically in pre-reduced PBS at a final concentration of 0.1 g/mL. The mixture was vortexed to produce a homogenous slurry and was then left for 30 minutes to let particulates settle. Aliquots of the supernatant were dissolved 50:50 with 40% glycerol in water and flash-frozen in liquid nitrogen, creating slurries. These slurries were stored at _80 ºC and were defrosted anaerobically at room temperature at the time of use.

**Screen for catechol dehydroxylation by mammalian gut microbiota samples**

Mammalian fecal slurries were prepared as described above and were defrosted by incubation at room temperature at the time of use. 20 µL of each slurry was then combined with 980 µL of basal medium containing 500 µM each of dopamine, (+) catechin, DOPAC, or hydrocaffeic acid. Each individual sample was grown in one well, with the n=3 individual samples for each animal serving as the biological replicates. Control wells contained compound but no bacteria. Samples were grown anaerobically at 37 ºC for 96 hours in a 96-well plate (Agilent Technology, catalog# A696001000). Following growth, we first assessed the total microbial growth by measuring the $OD_{600}$ in a plate reader (BioTek Synergy HTX). Cultures were then harvested by centrifugation and 50 µL of supernatant was transferred to a new 96-well plate, at which time the catechol colorimetric assay was used to assess total dehydroxylation by the complex microbial community. Samples that had potential catechol depletion as assessed by the colorimetric assay were then further analyzed by LC-MS. To prepare samples for LC-MS, 20 µL of the culture supernatant was diluted 1:10 with 180 µL of methanol, followed by centrifugation at 4000 rpm for 10 minutes to pellet particulates, salts, and proteins. 50 µL of the resulting supernatant was then transferred to the LC-MS 96-well plate and 5 µL of the supernatant was injected onto the instrument using Method A for dopamine, Method E for catechin, and Method F for DOPAC and hydrocaffeic acid.

**Construction of a mammalian phylogenetic tree**

The mammalian phylogenetic tree was generated using the Automatic Phylogenetic Tree Generator (aptg, version 0.1.0) script in R (version 3.5.1). Mammals not part of the aptg database were added manually to the tree using additional information about the mammalian phylogeny as a reference (*12*). The icons seen in Fig. 4 in the main text were adapted under a Creative Commons license (https://creativecommons.org/licenses/by/3.0/) at phylopic (http://phylopic.org), including Alpaca logo (made by Steven Traver), Bison (Lukasiniho). Cow (Steven Traver), Dog (Tracy A Heath), Fox (Anthony Caravaggi), Guinea pig (Zimices), Mouse (Madeleine Price Ball), Pig (Steven Traver), Rabbit (Steven Traver), Rabbit (Steven Traver), Rat (Rebecca Groom), Sheep (Zimices), and Wolf (Tracy A. Heath).

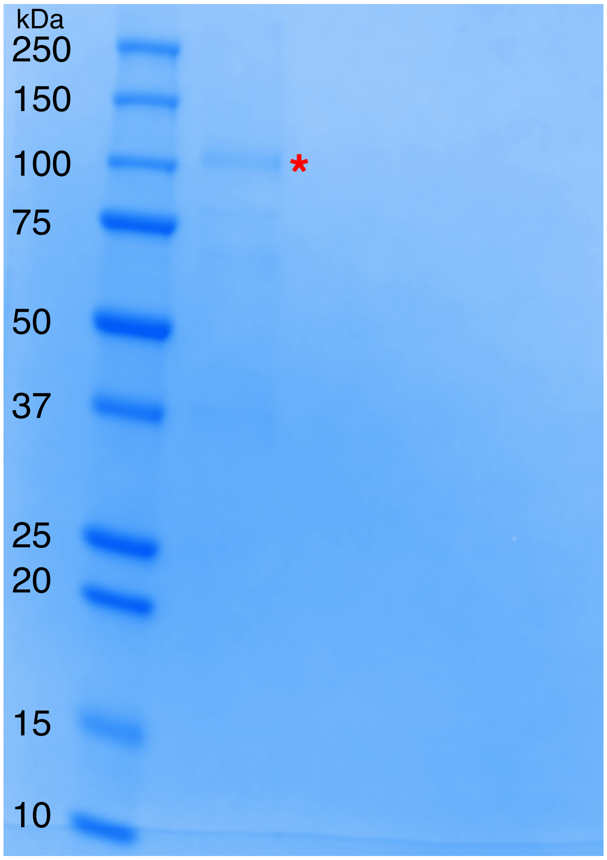

**Figure S1. SDS-PAGE of natively purified dopamine dehydroxylase from *E. lenta* A2.** Ladder is the Precision Plus Protein™ All Blue Standards (first lane from the left), while the subsequent lane represents the combined dopamine-dehydroxylating fractions from the size exclusion column, the last chromatography step of the activity-based purification from *E. lenta* A2. The dopamine dehydroxylase (115 kDa predicted size) is highlighted with the red asterisk.

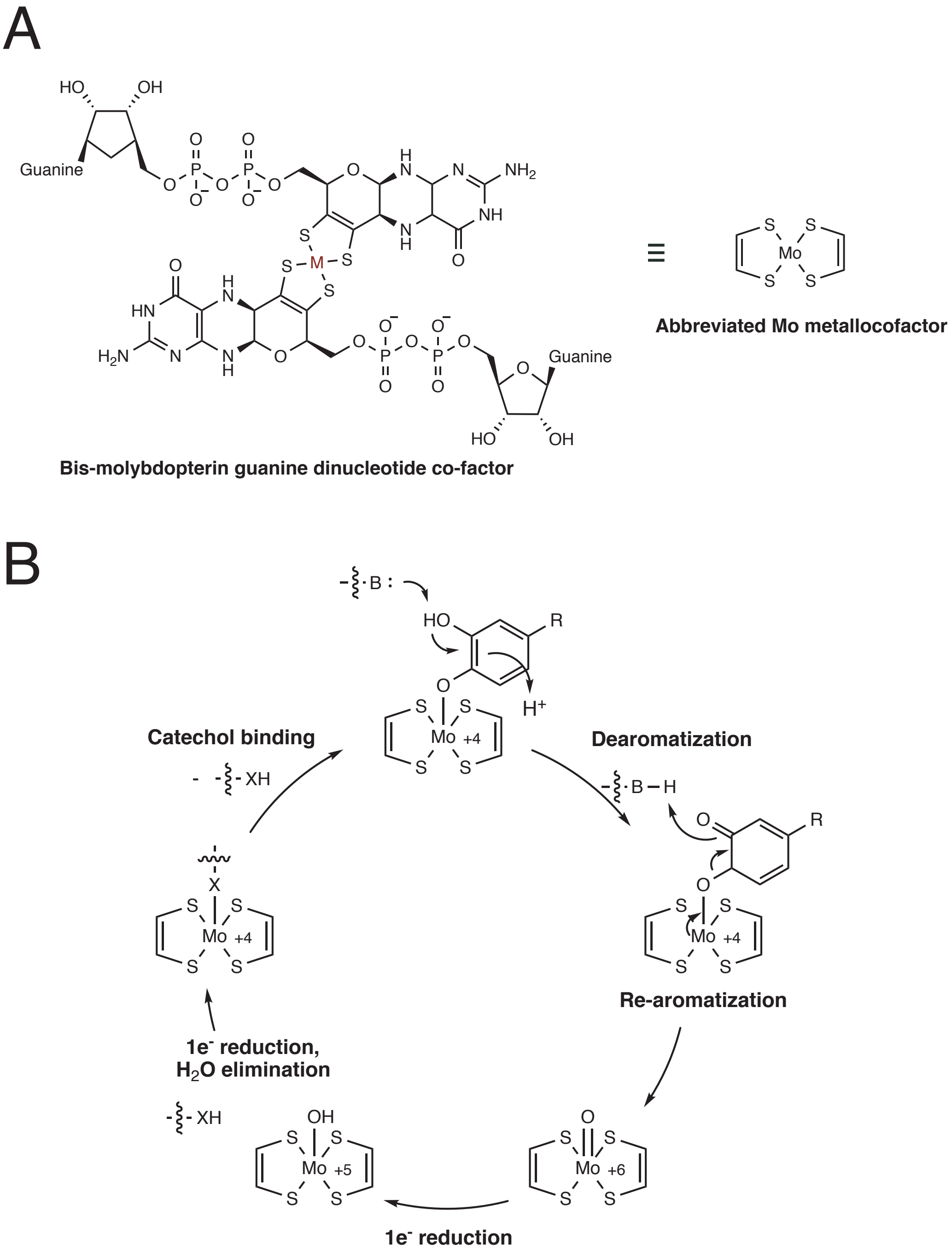

**Figure S2. Proposed mechanism for dopamine dehydroxylation by *E. lenta* A2 dopamine dehydroxylase.** A) structure of the bis-molybdopterin guanine dinucleotide co-factor that coordinates the catalytically essential molybdenum atom. M = metal. This metal can be either molybdenum (Mo) or tungsten (W). The metal center is predicted to be molybdenum in the dopamine dehydroxylase. B) Proposed mechanism for dopamine dehydroxylation involves de-aromatization and C–O bond cleavage driven by re-aromatization. The molybdenum atom is coordinated by an active site amino acid residue (here indicated as X, likely Aspartate or Cysteine), which is displaced upon catechol binding.

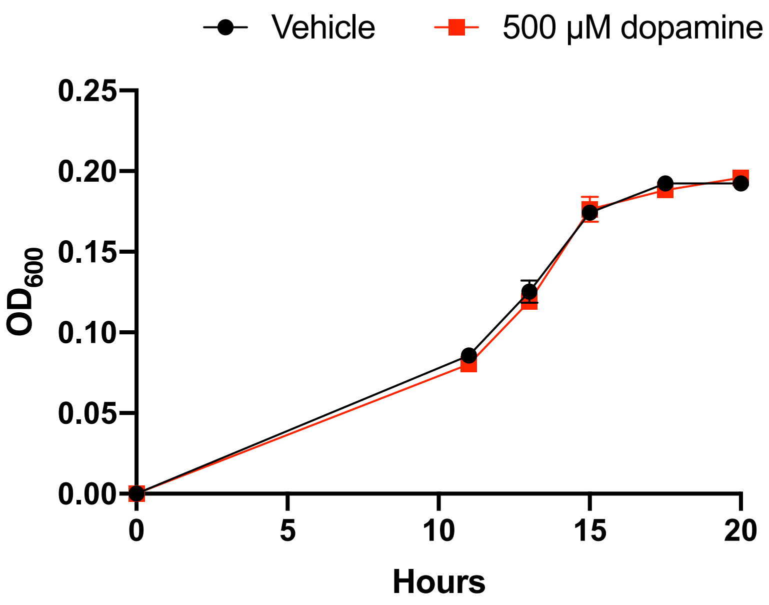

**Figure S3. Growth of *E. lenta* A2 in BHI medium with and without dopamine.** *E. lenta* A2 was grown anaerobically in BHI medium at 37 °C with and without dopamine. The data shown are the mean ± the SEM (n=3 replicate growth experiments).

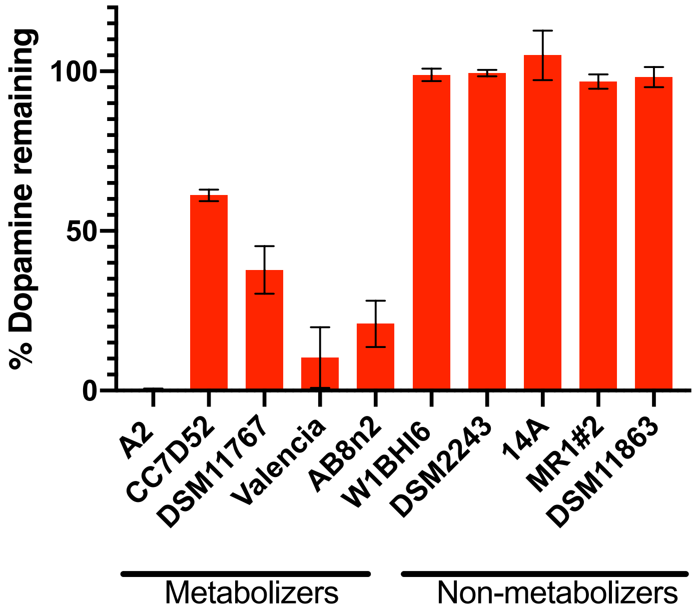

**Figure S4. Dehydroxylation of dopamine by *E. lenta* strains grown in in basal medium.** Strains were grown anaerobically for 48-72 hours at 37 ºC in basal medium containing 10 mM acetate before metabolism was assessed using the catechol colorimetric assay. Bars represent mean ± the SEM (n=3 biological replicates).

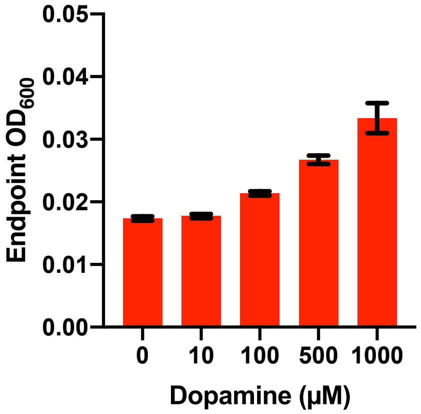

**Figure S5. Dose dependence of dopamine-promoted growth of *E. lenta* A2 in basal medium.** *E. lenta* was grown anaerobically in basal medium containing 10 mM acetate at 37 °C with varying concentrations of dopamine for 48 hours. Bars represent the mean ± the SEM (n=3 biological replicates).

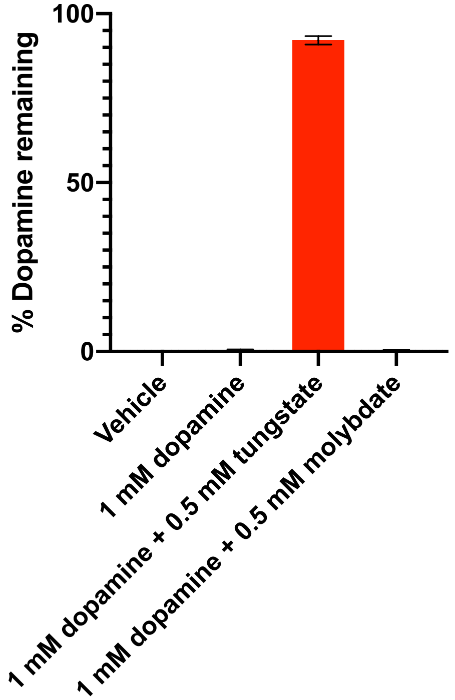

**Figure S6. Dehydroxylation of dopamine by *E. lenta* A2 in the presence of tungstate and molybdate in basal medium.** *E. lenta* was grown anaerobically in basal medium containing 10 mM acetate at 37 °C and 1 mM dopamine and either tungstate or molybdate (0.5 mM each) for 48 hours. Metabolism was assessed using the catechol colorimetric assay. Bars represent the mean ± the SEM of three biological replicates.

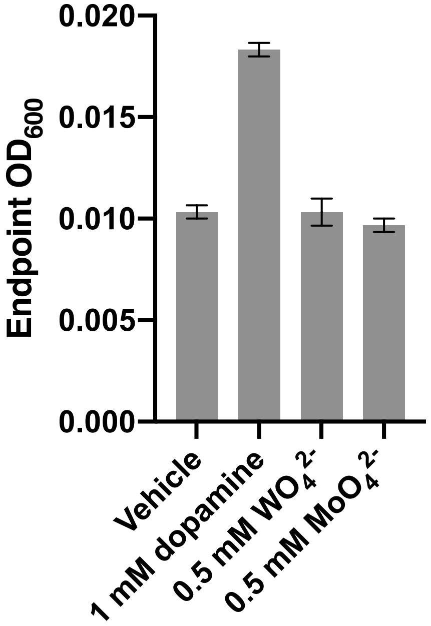

**Figure S7. Impact of tungstate and molybdate on *E. lenta* A2 growth in basal medium.** *E. lenta* was grown anaerobically in basal medium containing 10 mM acetate at 37 °C with either 1 mM dopamine, tungstate, or molybdate (0.5 mM each) for 36 hours before growth was assessed. Bars represent the mean ± the SEM of three biological replicates.

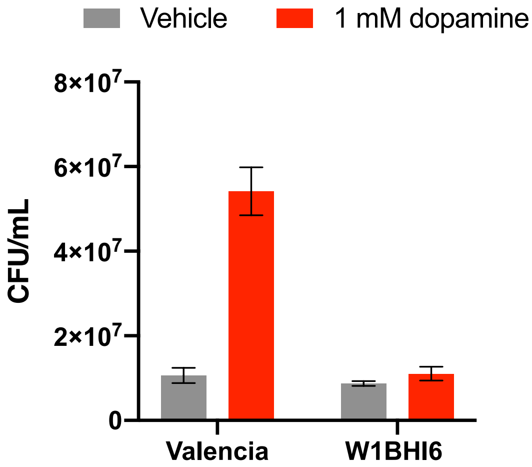

**Figure S8. CFU counts of *E. lenta* strains W1BHI6 and Valencia co-cultured in basal medium with and without dopamine.** Strains were grown together in basal medium containing 10 mM acetate for 72 hours at 37 ºC and were then plated. Antibiotic resistance was used to determine strain identity. Bars represent the mean ± the SEM of six biological replicates

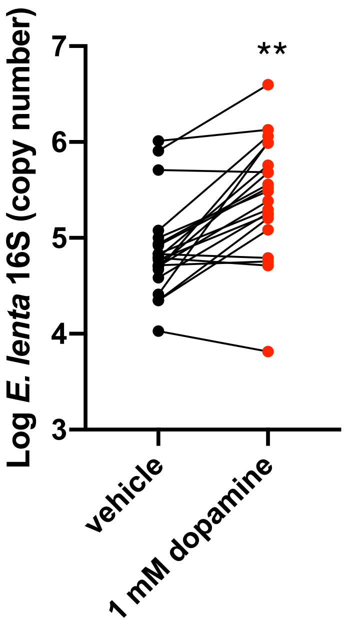

**Figure S9. *E. lenta* qPCR abundance in human fecal samples cultured with and without dopamine *ex vivo*.** Samples from unrelated individuals (n = 24) were grown for 72 hours at 37 ºC in basal medium containing 10 mM acetate with or without dopamine. qPCR was used to assess abundance of *E. lenta*. Two individuals were excluded from this analysis as they did not demonstrate quantitative metabolism of dopamine after incubation. Each point represents a different individual. Lines connect data from the same individual between the two conditions. (**$P<0.005$, two-tailed unpaired t-test).

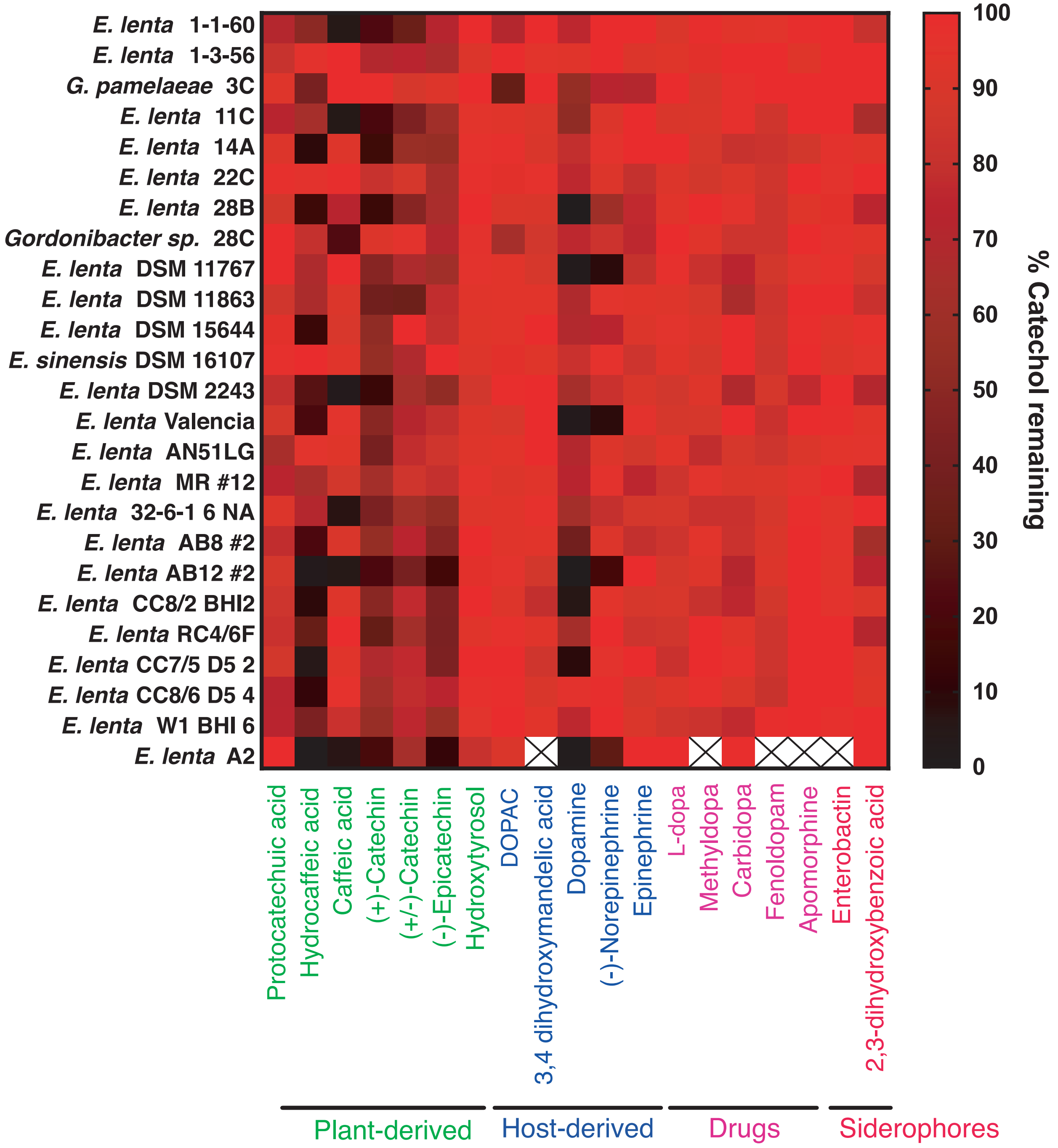

**Figure S10. Colorimetric screen for catechol dehydroxylation by human gut Actinobacteria.** Individual strains were grown in the presence of a single catechol substrate for 24-48 hours at 37 ºC in BHI medium containing 10 mM formate and 1 % arginine (w/v). Metabolism was assessed using a colorimetric assay that detects the catechol functional group. Data from bacterial incubations were normalized to the sterile control. Data represent the mean of three biological replicates. An X means that the strain was not screened for metabolism of the specific compound.

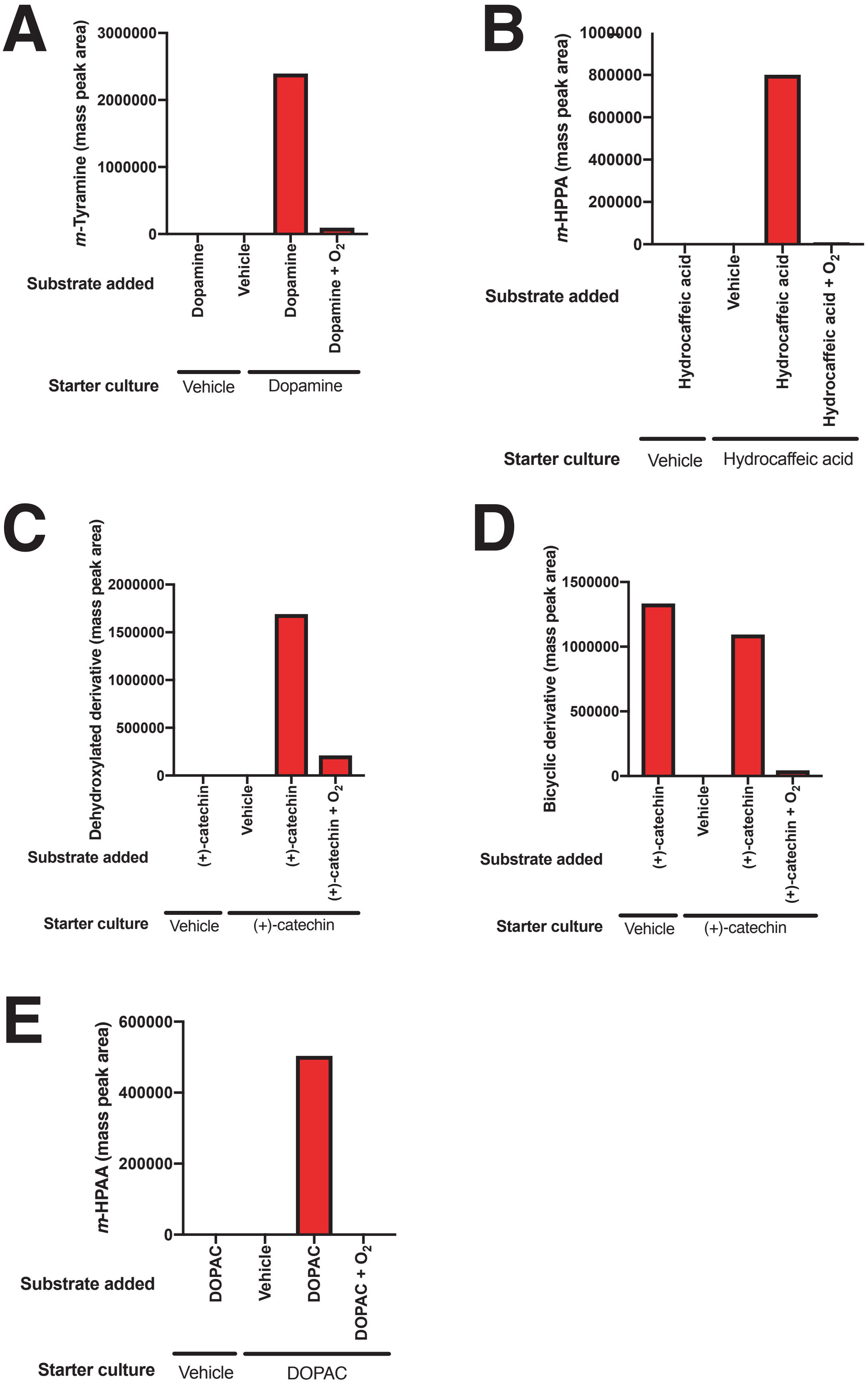
A
m-Tyramine (mass peak area)
3000000
2000000
1000000
0
Substrate added
Dopamine
Vehicle
Dopamine
Dopamine + O2
Starter culture
Vehicle
Dopamine
B
m-HPPA (mass peak area)
1000000
800000
600000
400000
200000
0
Substrate added
Hydrocaffeic acid
Vehicle
Hydrocaffeic acid
Hydrocaffeic acid + O2
Starter culture
Vehicle
Hydrocaffeic acid
C
Dehydroxylated derivative (mass peak area)
2000000
1500000
1000000
500000
0
Substrate added
(+)-catechin
Vehicle
(+)-catechin
(+)-catechin + O2
Starter culture
Vehicle
(+)-catechin
D
Bicyclic derivative (mass peak area)
1500000
1000000
500000
0
Substrate added
(+)-catechin
Vehicle
(+)-catechin
(+)-catechin + O2
Starter culture
Vehicle
(+)-catechin
E
m-HPAA (mass peak area)
600000
400000
200000
0
Substrate added
DOPAC
Vehicle
DOPAC
DOPAC + O2
Starter culture
Vehicle
DOPAC

**Figure S11. Inducibility and oxygen sensitivity of catechol dehydroxylation by whole cell suspensions of *E. lenta* A2 and *G. pamelaeae* 3C.** A) Inducibility and oxygen sensitivity of dopamine dehydroxylase activity in *E. lenta* A2. *E. lenta* A2 was grown anaerobically in BHI medium containing 1 % arginine and 10 mM formate. 0.5 mM of dopamine was added to induce dehydroxylase expression, followed by pelleting of cells and resuspension in PBS. Whole cell suspensions were exposed to dopamine (500 µM) or vehicle (water), and reactions were allowed to proceed anaerobically or aerobically for 20 hours. Assays mixtures were analyzed using LC-MS. Bar graph displays the mass peak area for a single sample. B) Inducibility and oxygen sensitivity of hydrocaffeic acid dehydroxylase activity in *E. lenta* A2. *E. lenta* A2 was grown anaerobically in BHI medium containing 1 % arginine and 10 mM formate. 0.5 mM of hydrocaffeic acid was added to induce dehydroxylase expression, followed by pelleting of cells and resuspension in PBS. Whole cell suspensions were exposed to hydrocaffeic acid (500 µM) or vehicle (water) and reactions were allowed to proceed anaerobically or aerobically for 20 hours. Assays mixtures were analyzed using LC-MS. Bar graph displays the mass peak area of *m*-hydroxyphenylpropionic acid (*m*-HPPA) for a single sample. Data shown are representative of at least two independent experiments. C) Inducibility and oxygen sensitivity of catechin dehydroxylase activity in *E. lenta* A2. *E. lenta* A2 was grown anaerobically in BHI medium containing 1% arginine and 10 mM formate. 0.5 mM of (+)-catechin was added to induce dehydroxylase expression, followed by pelleting of cells and resuspension in PBS. Whole cell suspensions were exposed to (+)-catechin (500 µM) or vehicle (DMF) and reactions were allowed to proceed anaerobically or aerobically for 20 hours. Assays mixtures were analyzed using LC-MS. Bar graph displays the mass peak area of the dehydroxylated catechin derivative in a single sample. D) Inducibility and oxygen sensitivity of (+)-catechin benzyl ether reduction activity in *E. lenta* A2. *E. lenta* A2 was grown anaerobically in BHI medium containing 1 % arginine and 10 mM formate. 0.5 mM of (+)-catechin was added to induce dehydroxylase expression, followed by pelleting of cells and resuspension in PBS. Whole cell suspensions were exposed to (+)-catechin (500 µM) or vehicle (DMF) and reactions were allowed to proceed anaerobically or aerobically for 20 hours. Assays mixtures were analyzed using LC-MS. Bar graph displays the mass peak area of the benzyl ether reduced catechin derivative in a single sample. Benzyl ether reduction was constitutive but oxygen sensitive in *E. lenta* A2. E) Inducibility and oxygen sensitivity of DOPAC acid dehydroxylase activity *G. pamelaeae* 3C. *G. pamelaeae* 3C was grown anaerobically in BHI medium containing 10 mM formate. 0.5 mM of DOPAC was added to induce dehydroxylase expression, followed by pelleting of cells and resuspension in PBS. Whole cell suspensions were exposed to DOPAC (500 µM) or vehicle (water) and reactions were allowed to proceed anaerobically or aerobically for 20 hours. Assays mixtures were analyzed using LC-MS. Bar graph displays the mass peak area *m*-hydroxyphenylacetic acid (*m*-HPAA) for a single sample.

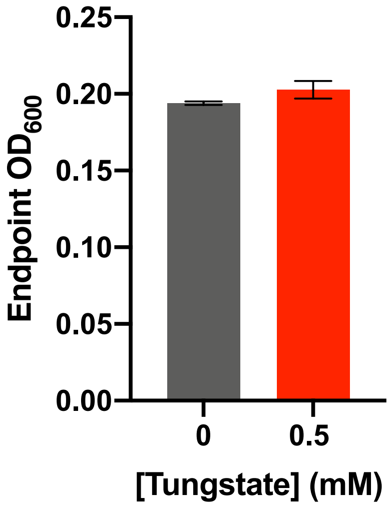

**Figure S12. Impact of tungstate on *E. lenta* A2 growth in BHI medium.** *E. lenta* was grown anaerobically in BHI at 37 °C with 0.5 mM tungstate for 48 hours before growth was assessed. Bars represent the mean ± the SEM of three biological replicates. At 0.5 mM, the concentration at which tungstate inhibits catechol dehydroxylation by *E. lenta*, there is no effect on growth.

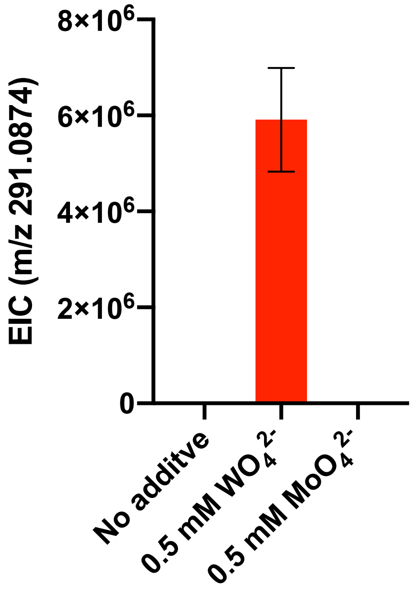

**Figure S13. Impact of tungstate on *E. lenta* A2 benzyl ether reduction of (+)-catechin in BHI medium.** *E. lenta* was grown anaerobically in BHI at 37 °C with (+)-catechin and either molybdate ($MoO_4^{2-}$) or tungstate ($WO_4^{2-}$) (0.5 mM each) for 48 hours. Culture supernatants were analyzed by LC-MS/MS. Bars represent the mean ± the SEM mass peak area of the benzyl ether reduced catechin derivative (three biological replicates). Treatment of cultures with tungstate blocks dehydroxylation, thus leading to build-up of the benzyl ether reduced catechin derivative (m/z 291.0874).

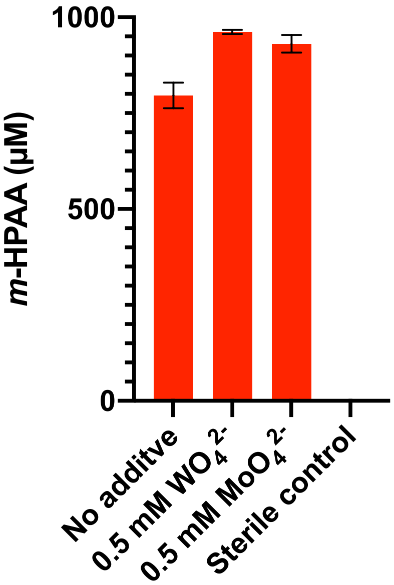

**Figure S14. Impact of tungstate on DOPAC dehydroxylation by *G. pamelaeae* 3C grown in BHI medium.** *G. pamelaeae* 3C was grown anaerobically with 0.5 mM DOPAC and either tungstate ($WO_4^{2-}$) or molybdate ($MoO_4^{2-}$) (0.5 mM each) for 48 hours 37 ºC. The concentration of *m*-HPAA, the product of direct DOPAC dehydroxylation, in culture supernatants was analyzed by LC-MS. Bars represent the mean ± the SEM of three biological replicates.

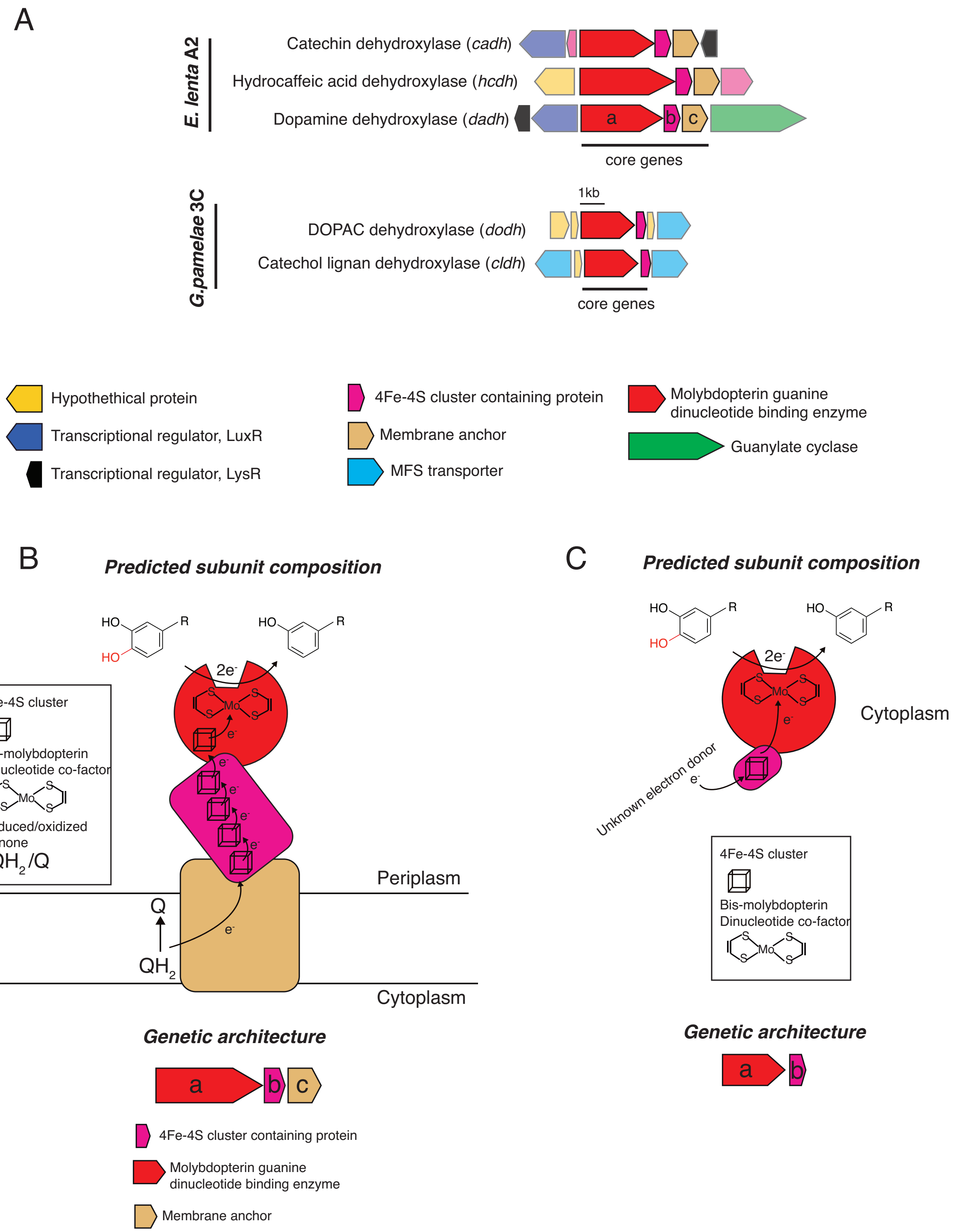

**Figure S15. Genomic contexts and predicted subunit composition of *Gordonibacter* and *E. lenta* dehydroxylases.** A) Genomic contexts of putative catechol dehydroxylases. Opaque ORFs encode core genes (ranging from a to c). Transparent ORFs encode genes not expected to directly participate in dehydroxylation. The *E. lenta* enzymes (Dadh, Hcdh, Cadh) are predicted to share similar subunit architectures. The molybdenum-dependent enzyme (a, red) associates with an electron shuttling 4Fe-4S ferredoxin (b, purple) and a membrane anchor (c, beige). In contrast, the *G. pamelaeae* predicted molybdenum-dependent enzymes are composed of a catalytic molybdenum subunit (a, red) and a small associated ferredoxin (b, purple). B) Model for *E.lenta* dehydroxylases. The predicted catalytic molybdenum-containing subunit (a, red) is co-localized with a predicted 4Fe-4S cluster-containing protein (b, purple) and a predicted membrane anchor (c, gold) in the genome. The molybdenum-containing enzyme is also predicted to contain an N-terminal 4Fe-4S cluster. Interestingly, the molybdenum-containing enzyme carries a Twin-Arginine-Translocation (TAT) signal sequence, suggesting it is exported. There, it could form a membrane-anchored complex that receives electrons from quinone oxidation in the membrane. Electrons from quinone oxidation could move through the iron sulfur clusters of the various subunits to ultimately reduce the molybdenum cofactor that performs the 2e- reduction of the catechol substrate. C) Model for *Gordonibacter* dehydroxylases. The predicted catalytic molybdenum-containing subunit (a, red) is co-localized with a 4Fe-4S cluster-containing protein (b, purple) in the genome. These enzymes are not predicted to be translocated across the periplasm as they lack the TAT signal sequence. The molybdenum cofactor-containing subunit also does not encode for a canonical 4Fe-4S cluster. In contrast to the *E. lenta* dehydroxylases, the *Gordonibacter* dehydroxylases are predicted to be free-standing as there is no membrane anchor encoded nearby. Electrons from could derive from an as-yet-unknown donor and be transferred to the iron sulfur cluster of the predicted 4Fe-4S cluster-containing protein to reduce the molybdenum cofactor that performs the 2e- reduction of the catechol substrate.

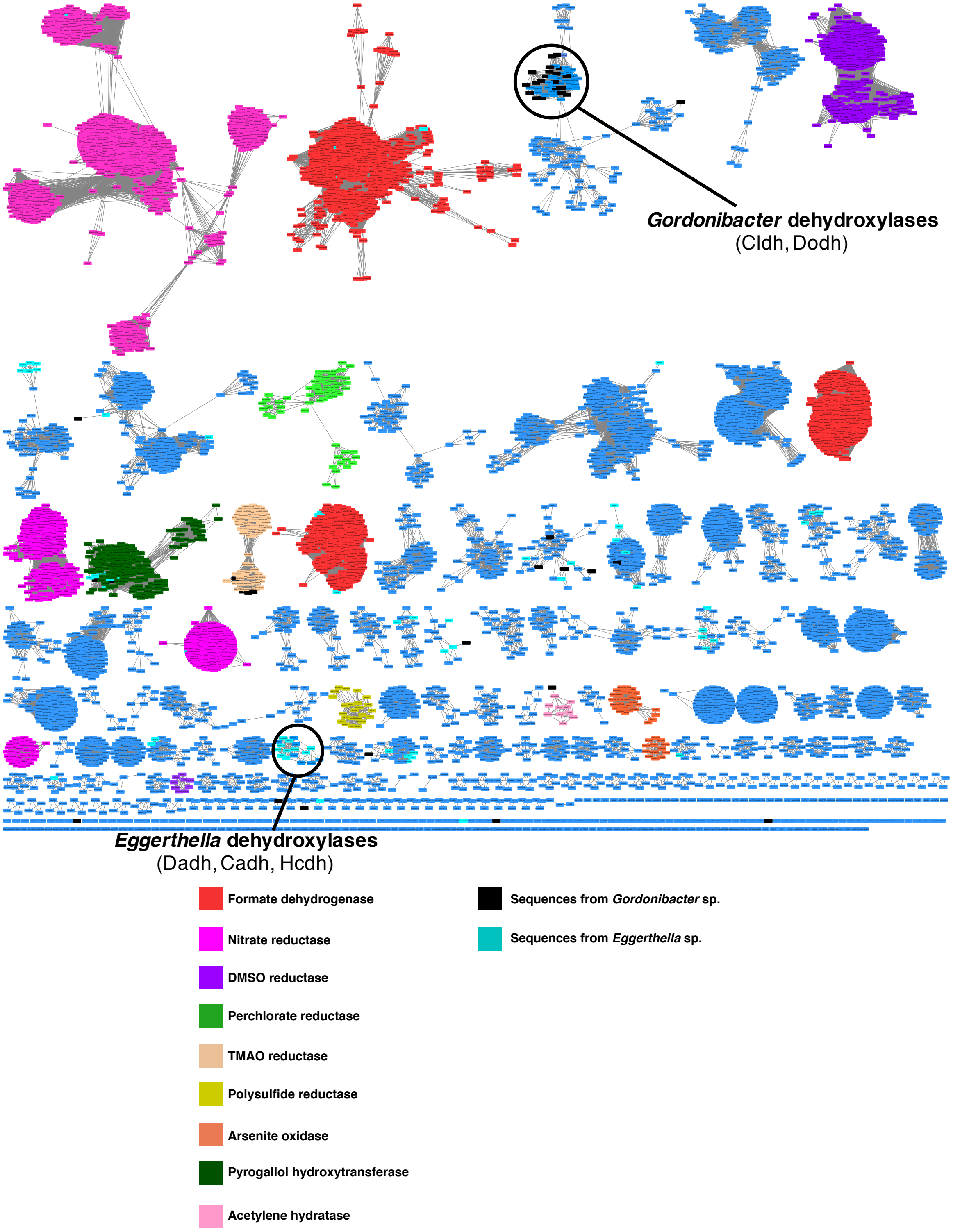

**Figure S16. Sequence similarity network of the bis-MGD enzyme family reveals that the *Gordonibacter* and *Eggerthella* dehydroxylases belong to distinct, uncharacterized isofunctional clusters.** The SSN was constructed using molybdopterin dinucleotide binding domain enzyme superfamily (PF01568) from Uniprot. Nodes represent proteins with 75%

sequence identity. SSN displayed with an e-value threshold of $10^{-167}$. All nodes that co-clustered with characterized enzymes are shown in the same color, denoting putative isofunctional activity. Turquois nodes represent sequences belonging to *Eggerthella* sp.*,* while black nodes represent sequences belonging to *Gordonibacter* sp. The clusters containing the *Eggerthella* and *Gordonibacter* catechol dehydroxylases form unique clusters separate from sequences of other biochemically characterized enzymes and away from other *Gordonibacter* and *Eggerthella* enzymes.

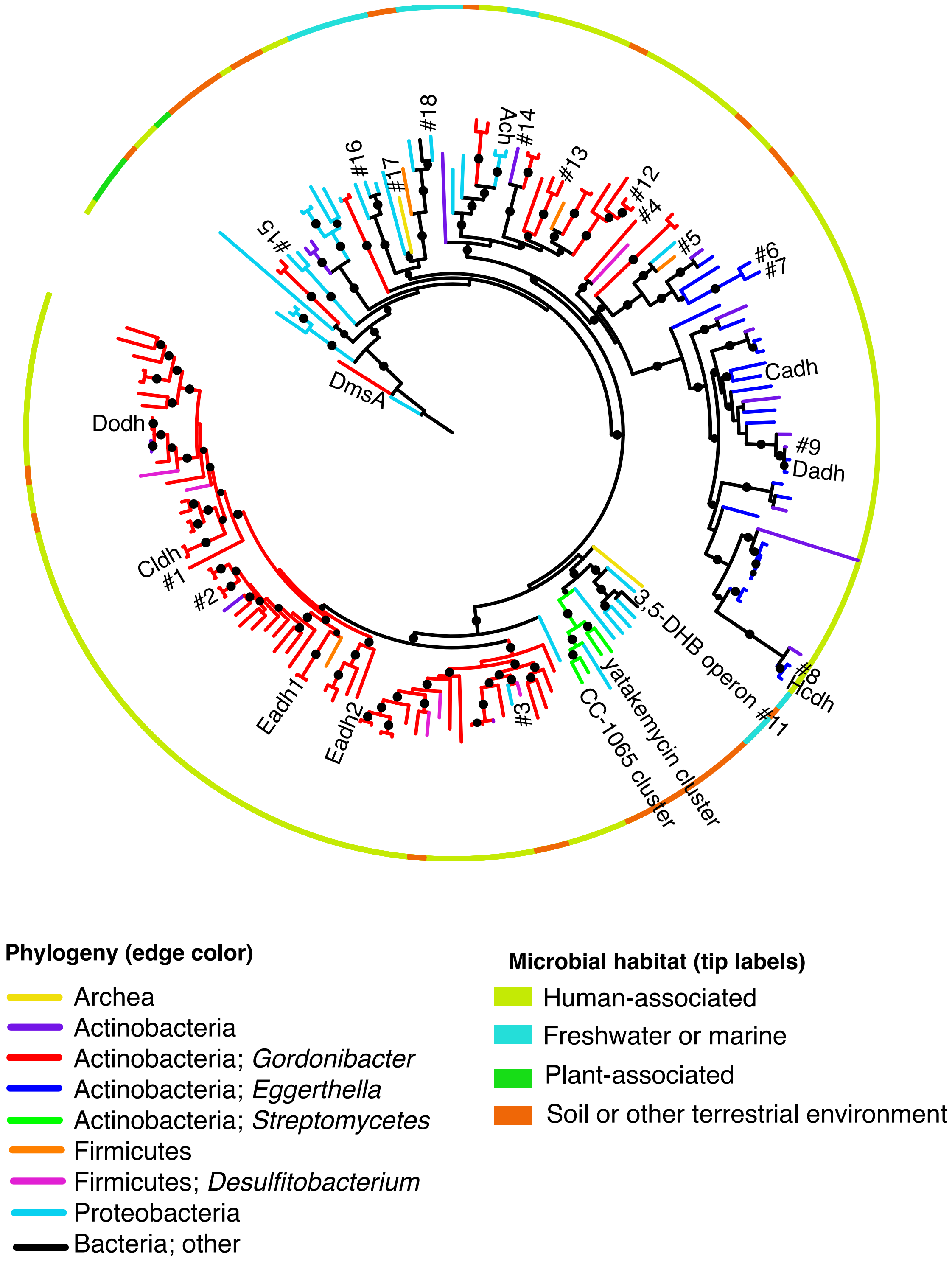

**Figure S17. Phylogenetic analysis of representative catechol dehydroxylase homologs highlighting sequences included in further analyses.** This phylogenetic tree is the same as in Figure 5 in the main text. The numbers indicate homologs that were selected as representative sequences across the tree for further phylogenetic analysis. The accession numbers of these sequences can be found in table S14.

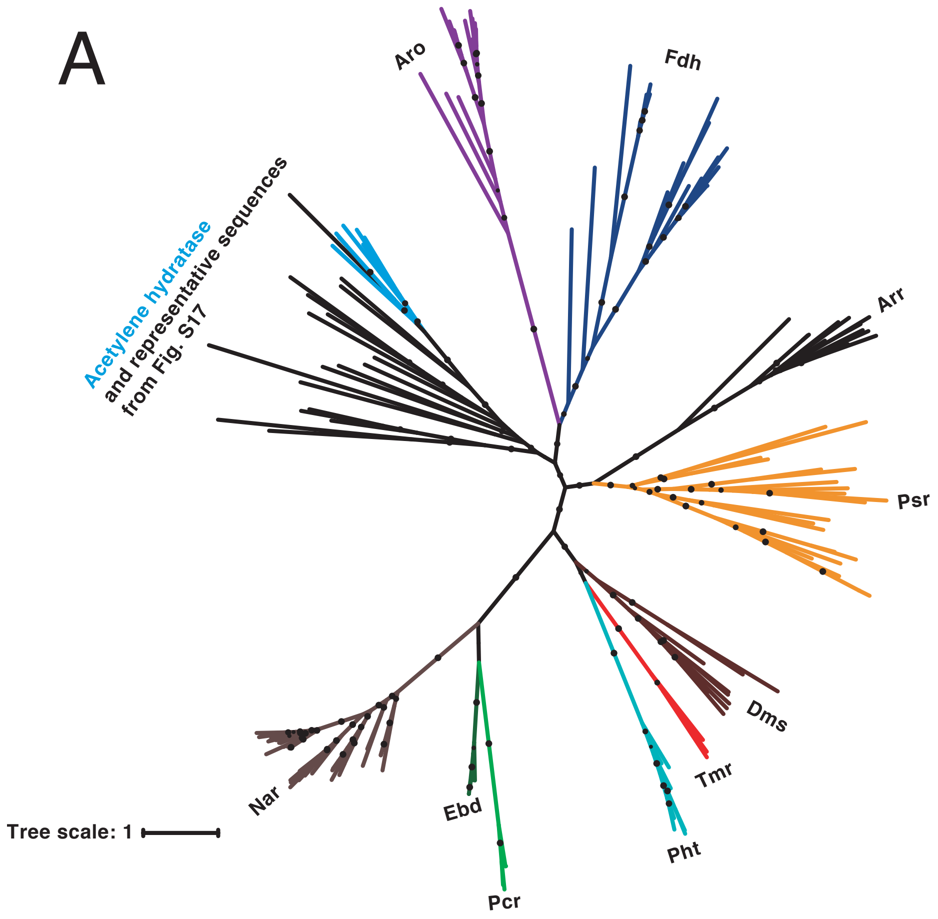

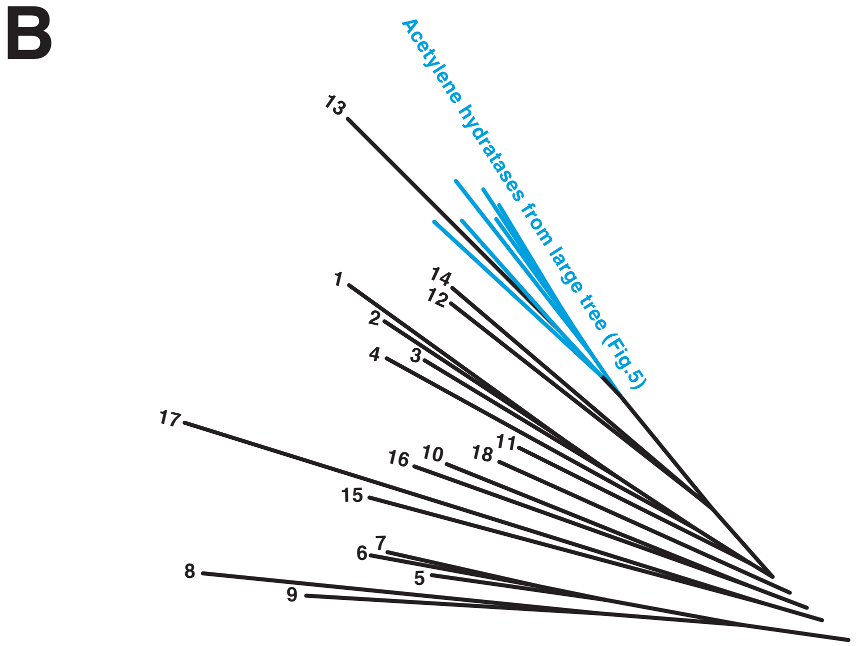

**Figure S18. Phylogenetic analysis of putative catechol dehydroxylase sequences from diverse bacterial phyla reveals a relationship with the biochemically characterized molybdenum-dependent enzyme acetylene hydratase.** A) The representative sequences from fig. S17 were added to the tree containing biochemically characterized members of the bis-MGD enzyme family (see main text Fig. 5). Psr = polysulfide reductase; Arr = arsenate reductase; Fdh = formate dehydrogenase; Pht = phloroglucinol transhydroxylase; Dms = DMSO reductase; Tmr = TMAO reductase; Aro = arsenite oxidase; Ebd = ethylbenzene dehydrogenase; Pcr = perchlorate reductase; Nar = nitrate reductase. Black circles on branches indicate bootstrap values greater than 0.7. This analysis revealed that all of these representative catechol dehydroxylase homologs are more closely related to each other and the acetylene hydratase sequences than they are to any other biochemically characterized bis-MGD enzyme family member. Acetylene hydratases are shown in blue and the representative sequences selected from fig.S17 are shown in black. B) Enlarged view of the clade containing the representative sequences and acetylene hydratases (main text Fig. 5).

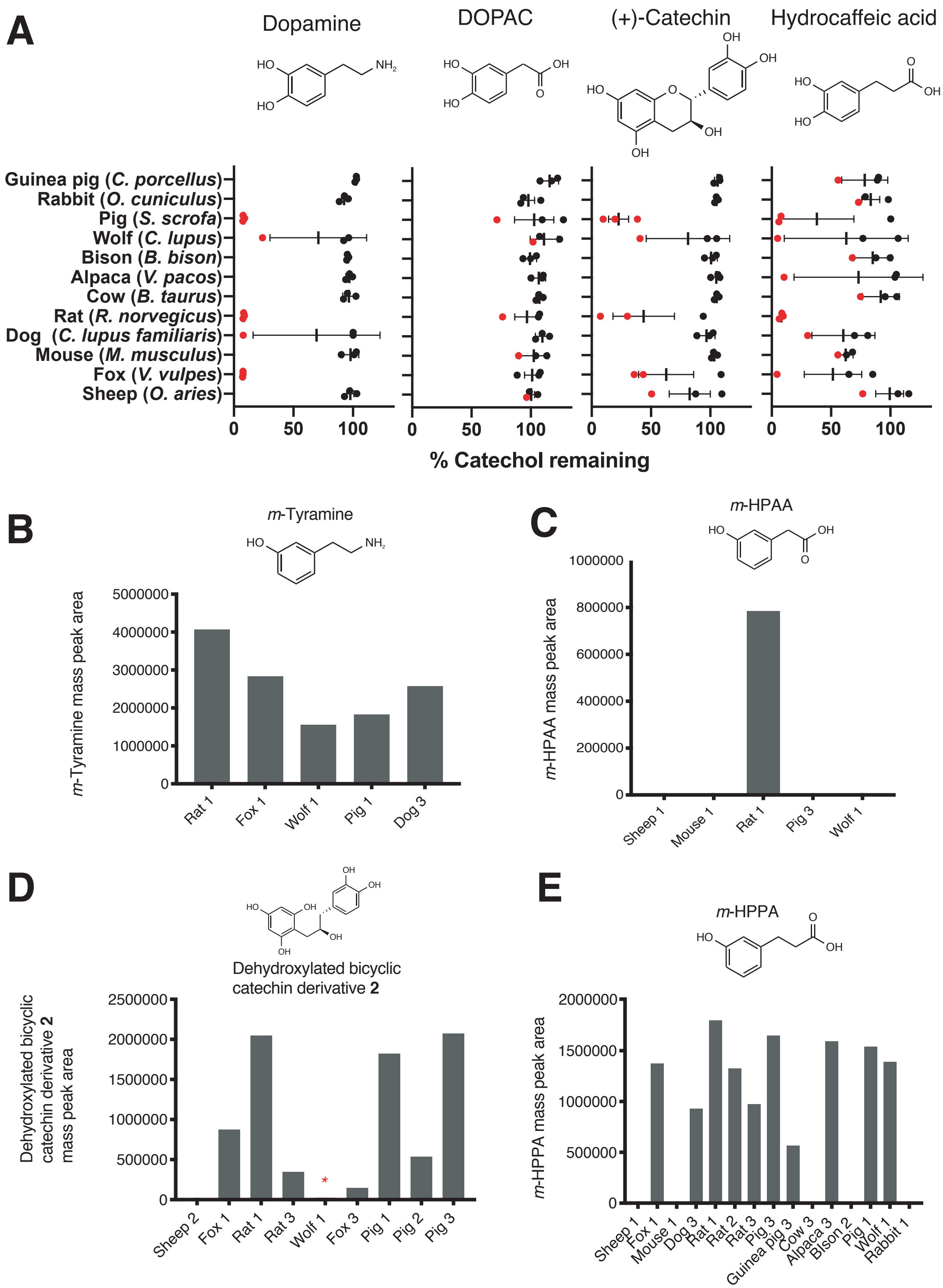

**Figure S19. Screen for catechol dehydroxylation by gut microbiota samples from diverse mammals.** A) Dehydroxylation of dopamine, DOPAC, (+)-catechin, and hydrocaffeic acid by gut microbiota samples from mammals spanning distinct diets and phylogenetic groups. Gut communities from n = 12 different mammals and n = 3 individuals per animal were cultured anaerobically for 96 hours in basal medium with 0.5 mM catechol at 37 ºC. Metabolism was then assessed by the colorimetric method for catechol detection. Data were normalized to the sterile control. Each dot represents a different individual for each animal, and the red color indicates samples that were selected for further LC-MS/MS analysis. Bars display the mean and standard deviation. B) *m*-Tyramine mass peak area in select mammalian gut microbiota samples incubated with dopamine and selected for LC-MS/MS analysis. Samples that displayed catechol depletion in A) performed dehydroxylation of dopamine. C) *m*-HPAA mass peak area in select mammalian gut microbiota samples incubated with DOPAC and selected for LC-MS/MS analysis. Only the rat microbiota (individual 1) had activity towards DOPAC. D) Dehydroxylated catechin derivative mass peak area in select mammalian gut microbiota samples incubated with (+)-catechin and selected for LC-MS/MS analysis. All samples selected for further analysis except the sheep microbiota displayed the full two-step conversion of (+)-catechin into the dehydroxylated derivative. The red asterisk indicates that the wolf microbiota had activity based on LC-MS/MS even though the mass peak area was lower than what is clearly visible with the current scale of the Y-axis. E) C) *m*-HPPA mass peak area in select mammalian gut microbiota samples incubated with hydrocaffeic acid and selected for LC-MS/MS analysis.

| Name | Structure | Natural origin | Catalog number |
|---|---|---|---|
| Caffeic acid | 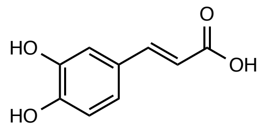 | Plant | Millipore Sigma, catalog# C0625-2G |
| Hydrocaffeic acid | 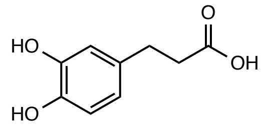 | Plant | Millipore Sigma, catalog# 102601-10G |
| (-)-epicatechin | 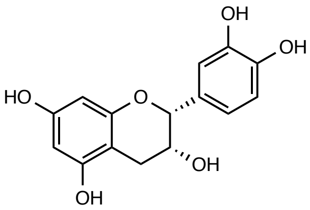 | Plant | Millipore Sigma, catalog# E1753-1G |
| (+)-catechin | 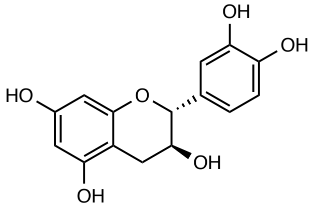 | Plant | Millimore Sigma, catalog# C1251-5G |
| 3,4-dihydroxybenzoic acid | 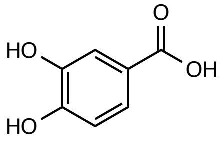 | Plant | Millipore Sigma, catalog# 37580-25G-F |
| 3,4-dihydroxyphenylacetic acid | 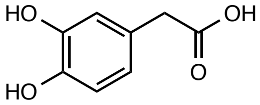 | Host | Millipore Sigma, catalog# 850217-1G |

| | | | |
|---|---|---|---|
| Ellagic acid | 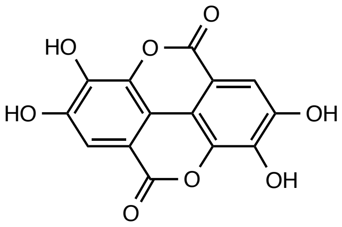 | Plant | Millipore Sigma, catalog# E2250-1G |
| DL-epinephrine | 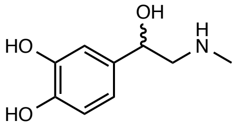 | Host | Sigma Aldrich, catalog# E4642-5G |
| DL-norepinephrine | 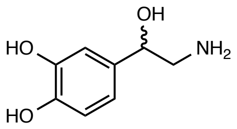 | Host | Millipore Sigma, catalog# A7256-1G |
| Dopamine | 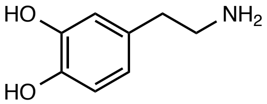 | Host | Sigma-Aldrich, catalog# PHR1090-1G |

**Table S1.**
Physiologically relevant catechol substrates used in assays with natively purified dopamine dehydroxylase from *E. lenta* A2.

| **Analog#** | **Name** | **Structure** | **Catalog number** |
|---|---|---|---|
| | Dopamine | 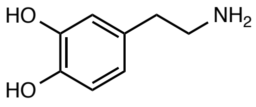 | Sigma-Aldrich, catalog# PHR1090-1G |
| **1** | *p*-tyramine | 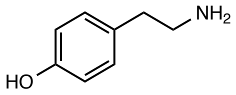 | Sigma Aldrich, catalog# T2879-1G |
| **2** | 3-aminotyramine | 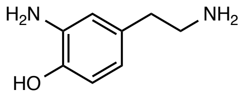 | Synthesized in our laboratory |

| | | | |
|---|---|---|---|
| **3** | 3-methoxytyramine | 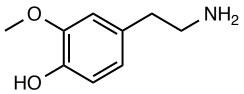 | Sigma Aldrich, catalog# M4251-100MG |
| **4** | 2,3-dihydroxyphenylethylamine | 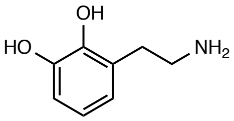 | Synthesized in our laboratory |
| **5** | 4,6-dihydroxyphenylethylamine | 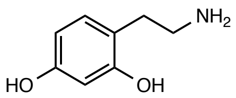 | Enamine, catalog # EN300-65185 |
| **6** | 1,4-dihydroxyphenylethylamine | 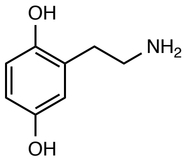 | Synthesized in our laboratory |
| **7** | 3,5-dihydroxyphenylethylamine | 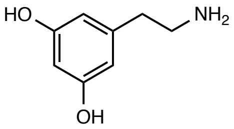 | Synthesized in our laboratory |
| **8** | Hydroxytyrosol | 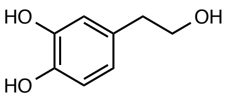 | Synthesized in our laboratory |

| | | | |
|---|---|---|---|
| **9** | 3,4-dihydroxybenzylamine | 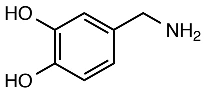 | Sigma Aldrich, catalog# 858781-250MG |
| **10** | *N*-methyldopamine | 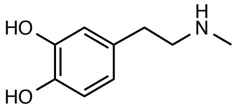 | Santa Cruz, catalog# sc-358430A |
| **11** | 2-hydroxydopamine | 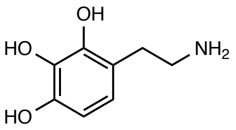 | Synthesized in our laboratory |
| **12** | 6-hydroxydopamine | 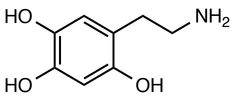 | Sigma Aldrich, catalog #H4381-100MG |
| **13** | 5-hydroxydopamine | 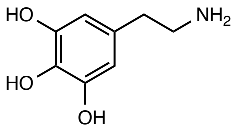 | Sigma Aldrich, catalog# 151564-100G |
| **14** | 2,3,5-trihydroxyphenylethylamine | 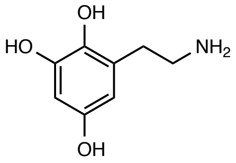 | Synthesized in our laboratory |

**Table S2.**
Commercially available and synthesized dopamine analogs used in assays with natively purified dopamine dehydroxylase from *E. lenta* A2.

| Locus tag | Annotation | log2FoldChange | p-value (FDR<0.1) |
|---|---|---|---|
| Elenta-A2_00671 | fumarate_reductase/succinate_dehydrogenase | 11.41 | 1.08E-25 |
| Elenta-A2_00535 | molydopterin_dinucleotide-binding_region (*dadh*) | 9.60 | 6.64E-13 |
| Elenta-A2_00533 | hypothetical_protein | 7.14 | 5.34E-13 |
| Elenta-A2_00999 | hypothetical_protein | 2.20 | 2.37E-13 |
| Elenta-A2_01000 | hypothetical_protein | 2.19 | 1.36E-15 |
| Elenta-A2_00997 | DBCCT_transporter | 2.15 | 3.73E-10 |
| Elenta-A2_00998 | Creatininase | 2.01 | 5.47E-06 |
| Elenta-A2_00996 | General_substrate_transporter | 1.61 | 8.79E-07 |
| Elenta-A2_01462 | hypothetical_protein | 1.43 | 3.08E-09 |
| Elenta-A2_01463 | 4Fe-4S_ferredoxin_iron-sulfur_binding_domain_protein | 1.43 | 4.98E-07 |
| Elenta-A2_01840 | Integrase_core_domain_protein | 1.42 | 1.04E-07 |
| Elenta-A2_01464 | molydopterin_dinucleotide-binding_region | 1.35 | 6.36E-16 |
| Elenta-A2_02451 | DDEAD/DEAH_box_helicase_domain_protein | 1.25 | 1.18E-25 |
| Elenta-A2_01910 | molydopterin_dinucleotide-binding_region | -1.12 | 1.90E-19 |
| Elenta-A2_01559 | membrane_bound_O-acyl_transferase | -1.17 | 2.43E-07 |
| Elenta-A2_00170 | flavocytochrome_c | -1.79 | 3.16E-10 |

**Table S3.**
Differentially expressed genes upon exposure of *Eggerthella lenta* A2 to 0.5 mM DL-norepinephrine relative to vehicle (>|2|-fold difference, FDR<0.1). The dopamine dehydroxylase (*dadh*) in *E. lenta* A2 is highlighted in red.

| **Bacterial strains used in this study** |
|---|
| *Eggerthella lenta* 1-1-60 |
| *Eggerthella lenta* 11C |
| *Eggerthella lenta* 1-3-56 |
| *Eggerthella lenta* 14A |
| *Eggerthella lena* 22C |
| *Eggerthella lenta* 28B |
| *Eggerthella lenta* 32-6-I 6 NA |
| *Eggerthella lenta* AB12 #2 |
| *Eggerthella lenta* AB8 #2 |
| *Eggerthella lenta* AN51LG |
| *Eggerthella lenta* CC7/5 D5 2 |
| *Eggerthella lenta* CC8/2 BHI2 |
| *Eggerthella lenta* CC8/6 D5 4 |
| *Eggerthella lenta* DSM 11767 |
| *Eggerthella lenta* DSM 11863 |
| *Eggerthella lenta* DSM 15644 |

| |
|---|
| *Eggerthella lenta* DSM2243 |
| *Eggerthella lenta* MR #12 |
| *Eggerthella lenta* RC4/6F |
| *Eggerthella lenta* Valencia |
| *Eggerthella lenta* W1 BHI 6 |
| *Eggerthella sinesis* DSM 16107 |
| *Eggerthella lenta* A2 |
| *Gordonibacter pamelaeae* 28C |
| *Gordonibacter pamelaeae* 3C |
| *Enterococcus faecalis* OG1RF |
| *Eschericia coli* MG1655 |
| *Bacteroides fragilis* ATCC 25285 |
| *Clostridium sporogenes* ATCC 15579 |
| *Edwarsiella tarda* ATCC 23685 |
| *Paraeggerthella hongkongensis* RC2/2A |

**Table S4.**
Bacterial strains used in this study

| Name | Structure | Natural origin | Catalog number |
|---|---|---|---|
| Protocatechuic acid | 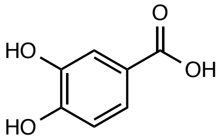  | Plant | Millipore Sigma, catalog# 37580-25G-F |
| Hydrocaffeic acid | 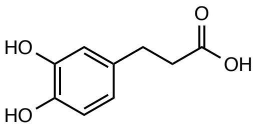  | Plant | Millipore Sigma, catalog# 102601-10G |

| | | | |
|---|---|---|---|
| Caffeic acid | 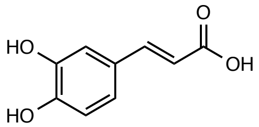 | Plant | Millipore Sigma, catalog# C0625-2G |
| (+)-catechin | 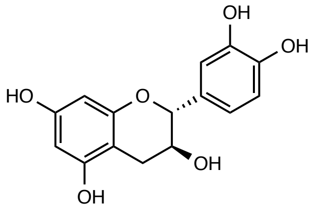 | Plant | Millimore Sigma, catalog# C1251-5G |
| (+/-)-catechin | 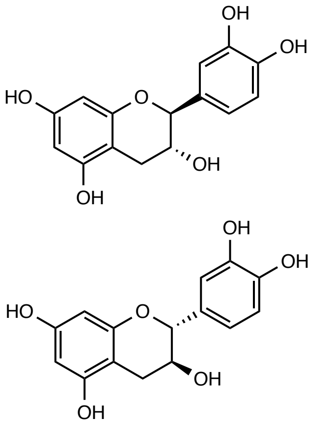 | Plant | Millipore Sigma, catalog# C1788-500MG |
| (-)-epicatechin | 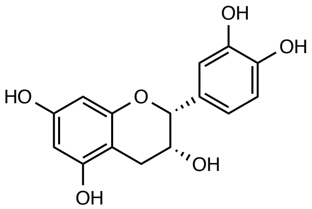 | Plant | Millipore Sigma, catalog# E1753-1G |
| Hydroxytyrosol | 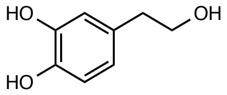 | Plant | Ava Chem Scientific, catalog# 2528 |

| | | | |
|---|---|---|---|
| DOPAC | 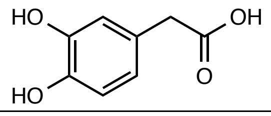 | Host | Millipore Sigma, catalog# 850217-1G |
| DL-3,4-DHMA | 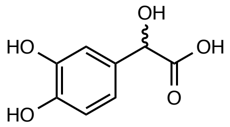 | Host | Carbo Synth, catalog# FD22118 |
| Dopamine | 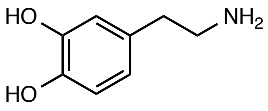 | Host | Sigma-Aldrich, catalog# H8502-25G |
| L-norepinephrine | 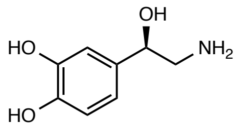 | Host | Matrix Scientific, catalog# 037592-500MG |
| L-epinephrine | 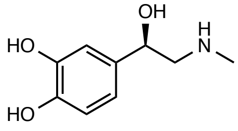 | Host | Alfa Aesar, catalog# L04911.06 |
| L-dopa | 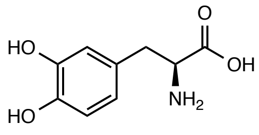 | FDA-approved drug | Oakwood Chemical, catalog# 358380-25g |

| | | | |
|---|---|---|---|
| Methyldopa | 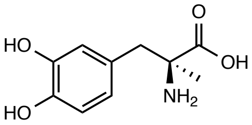 | FDA-approved drug | Chemcruz, catalog# sc-203092 |
| Carbidopa | 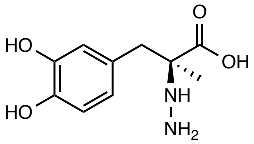 | FDA-approved drug | Sigma-Aldrich, catalog# PHR1655-1G |
| Fenoldopam | 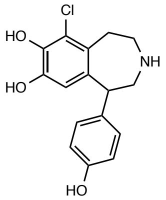 | FDA-approved drug | Sigma Aldrich, catalog# SML0198-10MG |
| Apomorphine | 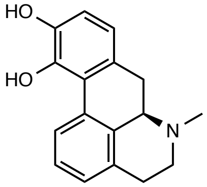 | FDA-approved drug | Sigma Aldrich, catalog# A4393-100MG |
| Enterobactin | 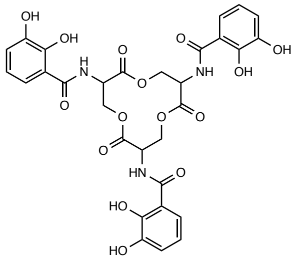 | Microbial siderophore | Gift from professorProf. Liz Elizabeth Nolan, MIT |

| | | | |
|---|---|---|---|
| 2,3-DHBA | 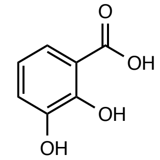 | Microbial siderophore | Millipore Sigma, catalog# 126209-5G |

**Table S5.**
Catechol substrates used in screen of gut Actinobacteria for catechol metabolism. 3,4-DHMA stands for 3,4-dihydroxymandelic acid. 2,3-DHBA stands for 2,3-dihydroxybenzoic acid.

**Dehydroxylated metabolite detected? (1=yes, 0=no)**

| | (+)-catechin | Hydrocaffeic acid | DOPAC |
|---|---|---|---|
| *Eggerthella lenta* 1-1-60 | 1 | 1 | 0 |
| *Eggerthella lenta* 11C | 1 | 1 | 0 |
| *Eggerthella lenta* 1-3-56 | 0 | 1 | 0 |
| *Eggerthella lenta* 14A | 1 | 1 | 0 |
| *Eggerthella lena* 22C | 0 | 1 | 0 |
| *Eggerthella lenta* 28B | 1 | 1 | 0 |
| *Eggerthella lenta* 32-6-I 6 NA | 0 | 1 | 0 |
| *Eggerthella lenta* AB12 #2 | 0 | 1 | 0 |
| *Eggerthella lenta* AB8 #2 | 0 | 1 | 0 |
| *Eggerthella lenta* AN51LG | 0 | 0 | 0 |
| *Eggerthella lenta* CC7/5 D5 2 | 0 | 1 | 0 |
| *Eggerthella lenta* CC8/2 BHI2 | 1 | 1 | 0 |
| *Eggerthella lenta* CC8/6 D5 4 | 0 | 1 | 0 |
| *Eggerthella lenta* DSM 11767 | 0 | 1 | 0 |
| *Eggerthella lenta* DSM 11863 | 0 | 1 | 0 |
| *Eggerthella lenta* DSM 15644 | 0 | 1 | 0 |
| *Eggerthella lenta* DSM2243 | 0 | 1 | 0 |
| *Eggerthella lenta* MR #12 | 0 | 1 | 0 |
| *Eggerthella lenta* RC4/6F | 0 | 1 | 0 |
| *Eggerthella lenta* Valencia | 0 | 1 | 0 |
| *Eggerthella lenta* W1 BHI 6 | 0 | 1 | 0 |
| *Eggerthella sinesis* DSM 16107 | 0 | 1 | 0 |
| *Eggerthella lenta* A2 | 1 | 1 | 0 |
| *Paraeggerthella hongkongensis* RC2/2A | 0 | 1 | 1 |
| *Gordonibacter pamelaeae* 28C | 0 | 1 | 1 |
| *Gordonibacter pamelaeae* 3C | 0 | 1 | 1 |
| *Sterile control* | 0 | 0 | 0 |

**Table S6.**
Confirmation of catechol metabolism by gut Actinobacterial library. We re-cultured strains with DOPAC, (+)-catechin, and hydrocaffeic acid to follow up on our colorimetric screen. *Paraeggerthella* RC2/2A was not included in the original screen but was included in the second round. Individual strains were grown in the presence of a single catechol substrate (n=3 replicates) for 48 hours at 37 ºC in BHI medium. Metabolism was assessed using LC-MS/MS. The table indicates whether the dehydroxylated metabolite was detected by LC-MS/MS.

| Locus tag | Annotation | log2FoldChange | p-value (FDR<0.01) |
|---|---|---|---|
| Elenta-A2_02813 | Hypothetical protein (*hcdh* membrane anchor) | 11.21 | 4.7E-62 |
| Elenta-A2_02815 | Molybdopterin dinucleotide binding region (*hcdh*) | 11.14 | 4.7E-163 |
| Elenta-A2_02814 | 4Fe-4S ferredoxin (*hcdh* 4Fe4S partner) | 10.31 | 8.0E-53 |
| Elenta-A2_02810 | 4Fe-4S ferredoxin | 8.31 | 6.8E-18 |
| Elenta-A2_02816 | Hypothetical protein | 7.75 | 3.1E-12 |
| Elenta-A2_02811 | 4Fe-4S ferredoxin | 6.83 | 6.7E-17 |
| Elenta-A2_02820 | Hypothetical protein | 6.37 | 5.3E-47 |
| Elenta-A2_02819 | Cytoplasmic chaperone TorD family protein | 6.29 | 2.7E-52 |
| Elenta-A2_02817 | Hypothetical protein | 5.80 | 5.1E-40 |
| Elenta-A2_02822 | Transcriptional regulator, LuxR family | 3.80 | 5.6E-14 |
| Elenta-A2_02821 | hypothetical_protein | 2.70 | 2.95E-83 |
| Elenta-A2_02818 | Hypothetical protein | 2.55 | 0.00013 |
| Elenta-A2_01460 | formate_dehydrogenase_family_accessory_protein | 2.12 | 1.32E-05 |
| Elenta-A2_01461 | formate_dehydrogenase_accessory_protein | 1.67 | 8.90E-07 |
| Elenta-A2_01459 | peptidase_C60_sortase_A_and_B | 1.52 | 0.004867669 |
| Elenta-A2_01464 | molydopterin_dinucleotide-binding_region | 1.50 | 3.06E-48 |
| Elenta-A2_01463 | 4Fe-4S_ferredoxin | 1.44 | 2.12E-23 |
| Elenta-A2_01465 | molybdopterin_oxidoreductase_Fe4S4_region | 1.39 | 1.76E-14 |
| Elenta-A2_01462 | hypothetical_protein | 1.22 | 1.24E-11 |
| Elenta-A2_02489 | ABC_transporter_related | 1.10 | 7.47E-06 |
| Elenta-A2_01331 | hypothetical_protein | 1.08 | 0.000121478 |
| Elenta-A2_00815 | hypothetical_protein | -1.08 | 1.49E-05 |
| Elenta-A2_01910 | molydopterin_dinucleotide-binding_region | -1.26 | 7.51E-17 |
| Elenta-A2_01911 | 4Fe-4S_ferredoxin | -1.36 | 5.47E-16 |

**Table S7.**

Differentially expressed genes upon exposure of *Eggerthella lenta* A2 to 0.5 mM hydrocaffeic acid relative to vehicle (>|2|-fold difference, FDR<0.1). *E. lenta* A2 was grown with 500 µM hydrocaffeic acid or a vehicle control in BHI medium containing 1% (w/v) arginine and 10 mM formate. The catalytic subunit of the putative hydrocaffeic acid dehydroxylase (*hcdh*), as well as its predicted 4Fe-4S and membrane anchor partners, are highlighted in red.

| Locus tag | Annotation | log2FoldChange | p-value (FDR<0.1) |
|---|---|---|---|
| Elenta-A2_00233 | fumarate_reductase/succinate_dehydrogenase | 14.38 | 3.22E-38 |
| Elenta-A2_00691 | fumarate_reductase/succinate_dehydrogenase | 11.32 | 2.09E-32 |
| Elenta-A2_00578 | 4Fe-4S_ferredoxin_iron-sulfur_binding_domain_protein (*cadh* 4Fe4S partner) | 9.09 | 1.24E-24 |
| Elenta-A2_00577 | molydopterin_dinucleotide-binding_region (*cadh*) | 8.85 | 2.50E-123 |
| Elenta-A2_00579 | hypothetical_protein (*cadh* membrane anchor) | 8.00 | 5.40E-60 |

| | | | |
|---|---|---|---|
| Elenta-A2_01215 | fumarate_reductase/succinate_dehydrogenase | 7.55 | 0 |
| Elenta-A2_00587 | hypothetical_protein | 3.99 | 6.36E-09 |
| Elenta-A2_01625 | NADH:flavin_oxidoreductase/NADH_oxidase | 3.99 | 8.47E-141 |
| Elenta-A2_01615 | Enoyl-CoA_hydratase/isomerase | 3.95 | 5.16E-178 |
| Elenta-A2_01619 | hypothetical_protein | 3.91 | 2.90E-66 |
| Elenta-A2_01621 | Electron_transfer_flavoprotein_alpha/beta-_subunit | 3.87 | 1.60E-20 |
| Elenta-A2_02817 | hypothetical_protein | 3.86 | 5.40E-12 |
| Elenta-A2_01620 | Electron transfer flavoprotein alpha subunit | 3.84 | 5.01E-23 |
| Elenta-A2_01613 | DL-carnitine dehydratase/bile acid inducible protein F | 3.82 | 1.66E-168 |
| Elenta-A2_01624 | acyl-CoA_dehydrogenase_domain_protein | 3.79 | 2.85E-217 |
| Elenta-A2_01614 | NADPH dependent FMN reductase | 3.72 | 6.55E-23 |
| Elenta-A2_01622 | acyl-CoA_dehydrogenase_domain_protein | 3.71 | 7.25E-25 |
| Elenta-A2_01609 | hypothetical_protein | 3.22 | 2.83E-43 |
| Elenta-A2_01618 | hydrolase | 3.15 | 3.25E-60 |
| Elenta-A2_01608 | transcriptional regulator2C LysR family | 2.98 | 3.05E-71 |
| Elenta-A2_01616 | AMP-dependent_synthetase_and_ligase | 2.97 | 1.06E-20 |
| Elenta-A2_01610 | transcriptional_regulator2C_LysR_family | 2.80 | 3.39E-21 |
| Elenta-A2_01546 | thioesterase_superfamily_protein | 2.71 | 8.10E-08 |
| Elenta-A2_01617 | beta-lactamase_domain_protein | 2.55 | 1.09E-40 |
| Elenta-A2_01493 | fumarate_reductase/succinate_dehydrogenase | 2.54 | 3.05E-33 |
| Elenta-A2_01611 | major_facilitator_superfamily_MFS_1 | 2.40 | 6.45E-60 |
| Elenta-A2_00576 | transcriptional_regulator2C_LuxR_family | 2.16 | 7.78E-19 |
| Elenta-A2_00581 | molybdopterin_dinucleotide-binding_region | 1.62 | 1.75E-10 |
| Elenta-A2_01602 | acyl-CoA_dehydrogenase_domain_protein | 1.62 | 2.17E-21 |
| Elenta-A2_02346 | argininosuccinate_synthase | 1.43 | 2.86E-24 |
| Elenta-A2_00138 | ABC_transporter_related | 1.41 | 8.05E-13 |
| Elenta-A2_02622 | chaperonin_GroEL | 1.30 | 4.58E-60 |
| Elenta-A2_00137 | hypothetical_protein | 1.29 | 1.67E-08 |
| Elenta-A2_00139 | transcriptional_regulator2C_TetR_famil | 1.27 | 1.80E-08 |
| Elenta-A2_01820 | phospho-N-acetylmuramoyl-pentapeptide-transferase | 1.23 | 1.99E-24 |
| Elenta-A2_01129 | MATE_efflux_family_protein | 1.21 | 1.10E-19 |
| Elenta-A2_02587 | hypothetical_protein | 1.19 | 1.09E-34 |
| lenta-A2_00167 | cell_wall/surface_repeat_protein | 1.19 | 3.32E-13 |
| Elenta-A2_00256 | Rhodanese_domain_protei | 1.17 | 7.25E-22 |
| Elenta-A2_02345 | argininosuccinate_lyase | 1.08 | 1.01E-10 |
| Elenta-A2_01456 | chaperonin_Cpn10 | 1.08 | 7.89E-25 |
| Elenta-A2_01456 | aspartate_carbamoyltransferase | 1.06 | 8.19E-10 |
| Elenta-A2_01326 | hypothetical_protein | -1.00 | 1.52E-13 |
| Elenta-A2_02456 | hypothetical_protein | -1.01 | 1.47E-24 |

| | | | |
|---|---|---|---|
| Elenta-A2_00896 | hypothetical_protein | -1.02 | 5.90E-12 |
| Elenta-A2_02832 | hypothetical_protein | -1.02 | 5.13E-30 |
| Elenta-A2_01974 | hypothetical_protein | -1.12 | 5.15E-24 |
| Elenta-A2_02146 | hypothetical_protein | -1.12 | 4.71E-33 |
| Elenta-A2_00730 | polar_amino_acid_ABC_transporter | -1.13 | 8.82E-09 |
| Elenta-A2_01591 | hypothetical_protein | -1.13 | 2.10E-13 |
| Elenta-A2_01359 | 4Fe-4S_ferredoxin_iron-sulfur_binding_domain_protein | -1.14 | 1.43E-08 |
| Elenta-A2_01382 | General_secretory_system_II_protein_E_domain_protein | -1.17 | 1.43E-07 |
| Elenta-A2_01347 | Endoribonuclease_L-PSP | -1.21 | 1.91E-31 |
| Elenta-A2_02130 | extracellular_solute-binding_protein | -1.21 | 2.28E-20 |
| Elenta-A2_02855 | chaperone_protein_DnaK | -1.22 | 8.03E-17 |
| Elenta-A2_00895 | tRNA_pseudouridine_synthase_A | -1.25 | 1.14E-24 |
| Elenta-A2_01051 | extracellular_solute-binding_protein | -1.26 | 3.04E-36 |
| Elenta-A2_00971 | ABC_transporter_related | -1.26 | 5.43E-38 |
| Elenta-A2_01601 | Aldehyde_Dehydrogenase | -1.26 | 2.44E-15 |
| Elenta-A2_00527 | anaerobic_dimethyl_sulfoxide_reductase | -1.34 | 1.02E-07 |
| Elenta-A2_00974 | polar_amino_acid_ABC_transporte | -1.39 | 3.48E-22 |
| Elenta-A2_02132 | hypothetical_protein | -1.43 | 6.39E-18 |
| Elenta-A2_02336 | hypothetical_protein | -1.44 | 7.43E-31 |
| Elenta-A2_02706 | membrane_protein_of_unknown_function | -1.44 | 4.33E-33 |
| Elenta-A2_02665 | two_component_transcriptional_regulator2C | -1.45 | 5.14E-37 |
| Elenta-A2_0097 | extracellular_solute-binding_protein | -1.46 | 9.23E-30 |
| Elenta-A2_01226 | ATPase2C_P-type | -1.52 | 7.75E-18 |
| Elenta-A2_00973 | polar_amino_acid_ABC_transporter | -1.54 | 1.91E-30 |
| Elenta-A2_00729 | extracellular_solute-binding_protein | -1.60 | 5.32E-16 |
| Elenta-A2_00458 | heat_shock_protein_Hsp20 | -1.70 | 1.35E-13 |
| Elenta-A2_01911 | 4Fe-4S_ferredoxin_iron-sulfur_binding_domain_protein | -1.71 | 2.48E-17 |
| Elenta-A2_01480 | 22C5-didehydrogluconate_reductase | -1.79 | 8.27E-34 |
| Elenta-A2_01940 | Mg2_transporter_protein | -1.85 | 8.42E-35 |
| Elenta-A2_02609 | lipase_class_3 | -1.90 | 6.13E-27 |
| Elenta-A2_01586 | membrane_protein_of_uknown_functio | -1.91 | 4.52E-11 |
| Elenta-A2_00505 | hypothetical_protein | -1.99 | 2.65E-26 |
| Elenta-A2_01113 | hypothetical_protein | -2.02 | 1.42E-09 |
| Elenta-A2_01910 | molydopterin_dinucleotide-binding_region | -2.09 | 1.29E-56 |
| Elenta-A2_01464 | molydopterin_dinucleotide-binding_region | -2.11 | 6.01E-18 |
| Elenta-A2_00508 | Polysulphide_reductase_NrfD | -2.15 | 9.73E-25 |
| Elenta-A2_01462 | hypothetical_protein | -2.15 | 2.94E-10 |
| Elenta-A2_00506 | molybdopterin_oxidoreductase | -2.28 | 6.51E-37 |
| Elenta-A2_00676 | Radical_SAM_domain_protein | -2.56 | 3.98E-15 |

| | | | |
|---|---|---|---|
| Elenta-A2_02199 | acterioferritin | -2.60 | 4.10E-11 |
| Elenta-A2_00507 | 4Fe-4S_ferredoxin_iron-sulfur_binding_domain_protein | -2.89 | 1.77E-11 |

**Table S8.**
Differentially expressed genes upon exposure of *Eggerthella lenta* A2 to 0.5 mM (+)-catechin relative to vehicle (>|2|-fold difference, FDR<0.1). *E. lenta* A2 was grown with 500 μM (+)-catechin or a vehicle control in BHI medium containing 1% (w/v) arginine and 10 mM formate. The catalytic subunit of the putative catechin dehydroxylase (*cadh*), as well as its predicted 4Fe-4S and membrane anchor partners, are highlighted in red.

| | Ach | Cldh | Dodh | Dadh | Cadh | Hcdh |
|---|---|---|---|---|---|---|
| **Ach** | | 26 | 23.9 | 19.3 | 20.7 | 15.7 |
| **Cldh** | 25.8 | | 45.2 | 22 | 23.2 | 17 |
| **Dodh** | 23.9 | 45 | | 20.2 | 22 | 15.7 |
| **Dadh** | 19.3 | 22 | 20.2 | | 50.9 | 35.3 |
| **Cadh** | 20.7 | 23 | 22 | 50.9 | | 39 |
| **Hcdh** | 15.7 | 17 | 15.7 | 35.3 | 39 | |

**Table S9**
Percent amino acid identity between the putative dehydroxylases from *G. pamelaeae* 3C and *E. lenta* A2. Ach stands for acetylene hydratase from *P. acetylenicus*. This protein is the closest biochemically characterized homolog to the newly identified catechol dehydroxylases. Colors represent % identity, with identity going from low (blue) to high (red).

| Locus tag | Annotation | log2FoldChange | p-value (FDR<0.1) |
|---|---|---|---|
| C1877_00905 | Radical_SAM_protein | 11.21 | 5.39E-80 |
| C1877_13890 | MFS_transporter | 11.16 | 0 |
| C1877_00920 | nitrate_ABC_transporter_substrate-binding_protein | 10.96 | 1.32E-49 |
| C1877_13905 | Dehydrogenase (*dodh* ) | 10.81 | 0 |
| C1877_13900 | Oxidoreductase (*dodh* 4Fe-4S partner) | 10.63 | 0 |
| C1877_00910 | ABC_transporter_permease | 10.28 | 1.31E-104 |
| C1877_13910 | Hypothetical_protein | 10.27 | 1.64E-171 |
| C1877_13895 | Hypothetical_protein | 10.10 | 0 |
| C1877_00915 | ABC_transporter_ATP-binding_protein | 9.66 | 1.72E-116 |
| C1877_00925 | BC_transporter_permease | 9.25 | 7.05E-46 |
| C1877_07205 | GTPase_G3E_family | 8.99 | 1.57E-271 |
| C1877_13865 | rimI:ribosomal-protein-alanine_N-acetyltransferase | 8.92 | 2.22E-252 |
| C1877_13885 | hypothetical_protein | 8.90 | 6.02E-288 |
| C1877_07570 | hypothetical_protein | 8.78 | 1.07E-254 |

| | | | |
|---|---|---|---|
| C1877_04445 | sodium:solute_symporter | 8.58 | 0 |
| C1877_07565 | dehydrogenase | 8.32 | 5.42E-93 |
| C1877_13930 | hypothetical_protein | 8.28 | 4.42E-32 |
| C1877_13870 | hypothetical_protein | 8.04 | 1.77E-70 |
| C1877_07200 | DUF3641_domain-containing_protein | 8.00 | 0 |
| C1877_13875 | hypothetical_protein | 7.85 | 0 |
| C1877_07185 | ABC_transporter_substrate-binding_protein | 7.85 | 6.85E-131 |
| C1877_07195 | ABC_transporter_permease | 7.74 | 1.21E-235 |
| C1877_07180 | DUF1847_domain-containing_protein | 7.59 | 3.20E-64 |
| C1877_07580 | MFS_transporter | 7.55 | 0 |
| C1877_07210 | methyltransferase | 7.34 | 4.27E-300 |
| C1877_07190 | ABC_transporter_ATP-binding_protein | 7.23 | 7.82E-100 |
| C1877_13880 | hypothetical_protein | 7.04 | 1.69E-91 |
| C1877_09915 | radical_SAM_protein | 6.85 | 0 |
| C1877_00930 | ABC_transporter_ATP-binding_protein | 6.44 | 1.27E-103 |
| C1877_09910 | hypothetical_protein | 6.36 | 0 |
| C1877_04725 | hypothetical_protein | 5.95 | 0 |
| C1877_00940 | hypothetical_protein | 5.89 | 1.85E-133 |
| C1877_13835 | DUF3641_domain-containing_protein | 5.56 | 0 |
| C1877_13840: | 4-carboxymuconolactone_decarboxylase | 5.32 | 0 |
| C1877_08360 | ABC_transporter_ATP-binding_protein | 5.24 | 1.80E-24 |
| C1877_07315 | hypothetical_protein | 5.22 | 1.59E-220 |
| C1877_13860 | hypothetical_protein | 5.14 | 0 |
| C1877_07560 | oxidoreductase | 5.03 | 4.60E-210 |
| C1877_13785 | 4Fe-4S_ferredoxin | 4.89 | 2.47E-20 |
| C1877_02200 | ATP-binding_protein | 4.81 | 0.020782133 |
| C1877_08370 | 16S_rRNA_Adenine_N6-dimethyltransferase_RsmA | 4.80 | 7.32E-09 |
| C1877_08365 | ABC_transporter_ATP-binding_protein | 4.78 | 3.06E-14 |
| C1877_13785 | 4Fe-4S_ferredoxin | 4.78 | 4.68E-300 |
| C1877_13790 | dehydrogenase | 4.61 | 0 |
| C1877_13780 | hypothetical_protein | 4.53 | 2.22E-238 |
| C1877_08355 | energy-coupling_factor_transporter | 4.25 | 4.16E-21 |
| C1877_04450 | hypothetical_protein | 4.02 | 1.11E-292 |
| C1877_01125 | phenylacetate--CoA_ligase | 3.97 | 2.44E-97 |
| C1877_05190 | hypothetical_protein | 3.93 | 0.036362059 |
| C1877_03100 | hypothetical_protein | 3.82 | 5.89E-68 |
| C1877_14220 | hypothetical_protein | 3.74 | 0.043112489 |
| C1877_12815 | arylamine_N-acetyltransferase | 3.69 | 0 |
| C1877_14165 | conjugal_transfer_protein_TraC | 3.65 | 0.045993369 |

| | | | |
|---|---|---|---|
| C1877_00945 | hypothetical_protein | 3.60 | 0.001453692 |
| C1877_13920 | LuxR_family_transcriptional_regulator | 3.42 | 9.77E-108 |
| C1877_13850 | hypothetical_protein | 3.41 | 1.34E-201 |
| C1877_13915 | hypothetical_protein | 3.39 | 0 |
| C1877_13925 | hypothetical_protein | 3.18 | 2.86E-187 |
| C1877_01120 | hypothetical_protein | 3.01 | 1.50E-263 |
| C1877_13775 | hypothetical_protein | 3.00 | 1.80E-127 |
| C1877_13770 | dehydrogenase | 2.94 | 0 |
| C1877_07550 | hypothetical_protein | 2.92 | 3.78E-119 |
| C1877_13970 | amidohydrolase | 2.74 | 6.24E-178 |
| C1877_13765 | DUF2064_domain-containing_protein | 2.53 | 9.85E-74 |
| C1877_03335 | TrpB-like_pyridoxal_phosphate-dependent_enzyme | 2.52 | 7.14E-11 |
| C1877_05595 | hypothetical_protein | 2.42 | 0.070104052 |
| C1877_09905 | ABC_transporter_ATP-binding_protein | 2.36 | 4.47E-128 |
| C1877_07555 | hypothetical_protein | 2.17 | 1.47E-56 |
| C1877_06120 | hypothetical_protein | 2.10 | 0.025206455 |
| C1877_07530 | hypothetical_protein | 2.05 | 5.92E-64 |
| C1877_13755 | 4Fe-4S-binding_protei | 2.00 | 2.31E-93 |
| C1877_07585 | helix-turn-helix_transcriptional_regulator | 1.96 | 3.32E-60 |
| C1877_09900 | hypothetical_protein | 1.95 | 7.02E-82 |
| C1877_13760 | radical_SAM_protein | 1.93 | 1.44E-163 |
| C1877_13750 | hypothetical_protein | 1.88 | 2.68E-59 |
| C1877_13230 | formate_dehydrogenase | 1.87 | 0.003539835 |
| C1877_00935 | hypothetical_protein | 1.79 | 4.10E-22 |
| C1877_04455 | endoglucanase | 1.74 | 3.14E-05 |
| C1877_13855 | hypothetical_protein | 1.73 | 1.73E-73 |
| C1877_00900 | hypothetical_protein | 1.72 | 0.010146811 |
| C1877_07535 | hypothetical_protein | 1.71 | 7.39E-45 |
| C1877_07295 | MFS_transporter | 1.66 | 0.01714762 |
| C1877_14955 | hypothetical_protein | 1.63 | 8.11E-66 |
| C1877_07310: | hypothetical_protein | 1.59 | 4.20E-07 |
| C1877_02570 | MFS_transporter | 1.53 | 1.11E-34 |
| C1877_14780: | site-2_protease_family_protein | 1.50 | 0.000440981 |
| C1877_02990 | DUF488_domain-containing_protein: | 1.33 | 0.04199253 |
| C1877_07300 | helix-turn-helix_transcriptional_regulator | 1.28 | 7.21E-08 |
| C1877_04550: | hypothetical_protein | 1.25 | 1.64E-16 |
| C1877_02800 | MFS_transporter | 1.22 | 0.045193782 |
| C1877_12325 | AraC_family_transcriptional_regulator | 1.13 | 0.027997941 |
| C1877_03030 | DUF4405_domain-containing_protein | 1.09 | 5.62E-16 |

| | | | |
|---|---|---|---|
| C1877_00895 | MFS_transporter | 1.09 | 4.56E-05 |
| C1877_07575 | hypothetical_protein | 1.08 | 2.59E-27 |
| C1877_09415 | GNAT_family_N-acetyltransferase | 1.08 | 9.38E-18 |
| C1877_03295: | hypothetical_protein | 1.08 | 0.000322218 |
| C1877_03035 | nickel-dependent_hydrogenase_large_subunit | 1.03 | 8.67E-19 |
| C1877_09205 | amino_acid_ABC_transporter_permease | 1.01 | 0.026568145 |
| C1877_01130 | LysR_family_transcriptional_regulator | 1.01 | 1.73E-14 |
| C1877_14775 | 3-oxoacyl-ACP_reductase | 1.00 | 0.004320782 |
| C1877_06060 | hypothetical_protein | -1.08 | 7.29E-12 |
| C1877_06730 | polysulfide_reductase | -1.08 | 3.98E-06 |
| C1877_06720 | hypothetical_protein | -1.09 | 7.77E-27 |
| C1877_10640 | hypothetical_protein | -1.13 | 4.05E-17 |
| C1877_10720 | LysR_family_transcriptional_regulator | -1.16 | 4.49E-22 |
| C1877_06070: | hypothetical_protein | -1.17 | 1.93E-10 |
| C1877_10665 | putative_sulfate_exporter | -1.29 | 1.91E-05 |
| C1877_10635 | hypothetical_protein | -1.29 | 2.87E-19 |
| C1877_10715 | nitrate_reductase_gamma_subuni | -1.35 | 1.71E-16 |
| C1877_10690 | porphobilinogen_synthase | -1.38 | 1.14E-10 |
| C1877_10685 | hemL:glutamate-1-semialdehyde-aminomutase | -1.42 | 7.29E-12 |
| C1877_10660 | hypothetical_protein | -1.45 | 2.18E-05 |
| C1877_10605 | sulfurtransferase_TusE | -1.51 | 2.66E-14 |
| C1877_10630 | menaquinol_oxidoreductase | -1.53 | 4.72E-23 |
| C1877_10600 | cobyrinic_acid_-diamide_synthase | -1.56 | 5.62E-16 |
| C1877_10705 | glutamyl-tRNA_reductase | -1.60 | 8.69E-18 |
| C1877_10700 | hydroxymethylbilane_synthase | -1.61 | 3.95E-15 |
| C1877_10695 | cobA:uroporphyrinogen-III_C-methyltransferase | -1.63 | 4.61E-10 |
| C1877_10610 | menaquinol_oxidoreductase | -1.63 | 1.98E-27 |
| C1877_10710 | Fe-S-binding_protein | -1.64 | 1.89E-17 |
| C1877_05025 | dehydrogenase | -1.68 | 1.09E-05 |
| C1877_10625 | Fe-S-binding_protein | -1.68 | 1.46E-26 |
| C1877_10615 | 4Fe-4S_dicluster_domain-containing_protein | -1.71 | 1.62E-31 |
| C1877_10620 | menaquinol_oxidoreductase | -1.77 | 3.21E-26 |
| C1877_07910 | hypothetical_protein | -2.12 | 0.000850235 |
| C1877_06075 | IS1_family_transposase | -2.16 | 0.051755192 |
| C1877_05990 | dehydrogenase | -2.20 | 0.068743035 |
| C1877_06180 | DUF1269_domain-containing_protei | -2.66 | 0.072663541 |
| C1877_02225 | hypothetical_protein | -2.68 | 0.040089976 |
| C1877_05995 | oxidoreductase | -3.40 | 0.062311297 |
| C1877_10895 | MFS_transporter | -3.73 | 0.026568145 |

| | | | |
|---|---|---|---|
| C1877_14300 | hypothetical_protein | -4.12 | 0.076180484 |
| C1877_02470 | transporter | -5.02 | 0.002337819 |

**Table S10.**
Differentially expressed genes upon exposure of *G. pamelaeae* 3C to 0.5 mM DOPAC relative to vehicle (>|2|-fold difference, FDR<0.1) when catechol was added during exponential phase. *G. pamelaeae* 3C was grown with 500 μM DOPAC or a vehicle control in BHI medium containing 10 mM formate. The catalytic subunit of the putative DOPAC dehydroxylase (*dodh*), as well as its predicted 4Fe-4S partner, are highlighted in red.

| Locus tag | Annotation | log2FoldChange | p-value (FDR<0.1) |
|---|---|---|---|
| C1877_00910 | ABC_transporter_permease | 8.05 | 4.65E-114 |
| C1877_00920 | nitrate_ABC_transporter_substrate-binding_protein | 7.78 | 1.13E-160 |
| C1877_13870 | hypothetical_protein | 7.42 | 3.46E-42 |
| C1877_13910 | hypothetical_protein | 7.33 | 3.64E-88 |
| C1877_07205 | GTPase_G3E_family | 7.23 | 1.53E-141 |
| C1877_13890 | MFS_transporter | 7.22 | 0 |
| C1877_13905 | Dehydrogenase (*dodh*) | 7.20 | 0 |
| C1877_00915 | ABC_transporter_ATP-binding_protein | 7.18 | 3.24E-140 |
| C1877_13900 | Oxidoreductase (*dodh* 4Fe-4S partner) | 7.11 | 0 |
| C1877_13895 | hypothetical_protein | 7.07 | 0 |
| C1877_00925 | ABC_transporter_permease | 7.05 | 1.11E-34 |
| C1877_13930 | hypothetical_protein | 6.73 | 7.65E-19 |
| C1877_13875 | hypothetical_protein | 6.56 | 1.51E-189 |
| C1877_00905 | radical_SAM_protein | 6.53 | 1.31E-59 |
| C1877_07190 | ABC_transporter_ATP-binding_protein | 6.44 | 3.33E-68 |
| C1877_13865 | ribosomal-protein-alanine_N-acetyltransferase | 6.36 | 2.98E-44 |
| C1877_07200 | DUF3641_domain-containing_protein | 6.12 | 7.13E-248 |
| C1877_07195 | ABC_transporter_permease | 6.12 | 9.85E-146 |
| C1877_07185 | ABC_transporter_substrate-binding_protein: | 6.02 | 2.87E-155 |
| C1877_07180 | DUF1847_domain-containing_protein | 6.01 | 2.12E-97 |
| C1877_13885 | hypothetical_protein | 5.77 | 9.05E-135 |
| C1877_07210 | methyltransferase | 5.43 | 4.47E-140 |
| C1877_04445 | sodium:solute_symporter | 5.03 | 4.08E-263 |
| C1877_13880 | hypothetical_protein | 4.16 | 1.04E-25 |

| | | | |
|---|---|---|---|
| C1877_00930 | ABC_transporter_ATP-binding_protein | 3.88 | 4.79E-147 |
| C1877_13835 | DUF3641_domain-containing_protei | 3.80 | 2.70E-254 |
| C1877_0731 | hypothetical_protein | 3.70 | 3.07E-100 |
| C1877_00940 | hypothetical_protein | 3.70 | 2.50E-59 |
| C1877_13840 | 4-carboxymuconolactone_decarboxylase | 3.42 | 3.87E-196 |
| C1877_09910 | hypothetical_protein | 3.36 | 3.42E-41 |
| C1877_09915 | hypothetical_protein | 3.27 | 2.53E-244 |
| C1877_09915: | radical_SAM_protein | 2.68 | 6.77E-208 |
| C1877_00935 | hypothetical_protein | 2.16 | 3.97E-38 |
| C1877_04450 | hypothetical_protein | 1.81 | 1.44E-61 |
| C1877_09900: | hypothetical_protein | 1.67 | 2.49E-52 |
| C1877_09905 | ABC_transporter_ATP-binding_protein | 1.60 | 5.09E-57 |
| C1877_13860 | hypothetical_protein | 1.60 | 9.49E-24 |
| C1877_14955 | hypothetical_protein | 1.41 | 2.20E-48 |
| C1877_13775 | hypothetical_protein | 1.32 | 1.58E-25 |
| C1877_13970 | amidohydrolase | 1.29 | 6.59E-41 |
| C1877_13785 | 4Fe-4S_ferredoxin | 1.27 | 4.40E-25 |
| C1877_13790 | dehydrogenase | 1.23 | 1.36E-34 |
| C1877_13780: | hypothetical_protein | 1.21 | 4.91E-18 |
| C1877_13750 | hypothetical_protein | 1.20 | 8.36E-32 |
| C1877_13925 | hypothetical_protein | 1.20 | 3.24E-28 |
| C1877_13770: | dehydrogenase | 1.18 | 3.12E-61 |
| C1877_13915: | hypothetical_protein: | 1.18 | 5.70E-44 |
| C1877_13755 | 4Fe-4S-binding_protein | 1.07 | 5.38E-33 |
| C1877_10245 | hypothetical_protein | -1.14 | 0.000531507 |
| C1877_10635 | hypothetical_protein | -1.33 | 1.41E-19 |

| | | | |
|---|---|---|---|
| C1877_10625 | Fe-S-binding_protein | -1.35 | 2.25E-16 |
| C1877_10640 | hypothetical_protein | -1.37 | 1.64E-23 |
| C1877_10610 | menaquinol_oxidoreductase | -1.40 | 4.17E-19 |
| C1877_10620 | menaquinol_oxidoreductase | -1.42 | 6.01E-15 |
| C1877_10630 | menaquinol_oxidoreductase | -1.45 | 7.21E-20 |
| C1877_10710 | Fe-S-binding_protein | -1.48 | 1.09E-13 |
| C1877_10600 | cobyrinic_acid_diamide_synthase | -1.51 | 2.26E-14 |
| C1877_10615 | 4Fe-4S_dicluster_domain-containing_protein | -1.51 | 1.89E-22 |
| C1877_10705 | glutamyl-tRNA_reductase: | -1.55 | 4.09E-15 |
| C1877_10715 | nitrate_reductase_gamma_subunit | -1.56 | 3.01E-20 |
| C1877_02335 | ferrous_iron_transporter_B | -1.57 | 1.60E-06 |
| C1877_10655 | hypothetical_protein | -1.65 | 1.43E-06 |
| C1877_10605 | sulfurtransferase_TusE | -1.65 | 3.30E-16 |
| C1877_10670 | dsrB:dissimilatory-type_sulfite_reductase | -1.66 | 1.58E-07 |
| C1877_10685 | glutamate-1-semialdehyde-aminomutase | -1.75 | 8.83E-16 |
| C1877_10690 | porphobilinogen_synthase | -1.78 | 4.43E-15 |
| C1877_10665 | putative_sulfate_exporter_family_transporter | -1.79 | 1.14E-09 |
| C1877_10675: | dsrA:dissimilatory-type_sulfite_reductase_alpha_subunit | -1.87 | 9.90E-11 |
| C1877_10660 | hypothetical_protein | -1.92 | 1.63E-08 |
| C1877_10695 | uroporphyrinogen-III_C-methyltransferase | -1.95 | 2.68E-13 |
| C1877_10650 | hypothetical_protein | -1.97 | 1.99E-06 |
| C1877_10700 | hydroxymethylbilane_synthase | -1.98 | 1.92E-19 |
| C1877_10680 | 4Fe-4S_dicluster_domain-containing_protein | -2.04 | 4.64E-09 |

**Table S11.**

Differentially expressed genes upon exposure of *G. pamelaeae* 3C to 0.5 mM DOPAC relative to vehicle (>|2|-fold difference, FDR<0.1) when catechol was added during the beginning of growth. *G. pamelaeae* 3C was grown with 500 μM DOPAC or a vehicle control in BHI medium containing 10 mM formate. Cells were harvested in mid-exponential phase when metabolism appeared. The catalytic subunit of the putative DOPAC dehydroxylase (*dodh*), as well as its predicted 4Fe-4S partner, are highlighted in red.

| Name | Abbreviated name | Organism | UNIPROT ID | Genbank protein ID |
|---|---|---|---|---|
| Dopamine dehydroxylase | *dadh* | *E. lenta* A2 | A0A369NIV7 | RDC23575.1 |
| Hydrocaffeic acid dehydroxylase | *hcdh* | *E. lenta* A2 | A0A369MIX7 | RDC18391.1 |
| Catechin dehydroxylase | *cadh* | *E. lenta* A2 | C8WLG2 | RDC23615.1 |
| Dopac dehydroxylase | *dodh* | *G. pamelae* 3C | A0A369LV65 | RDB62136.1 |
| Catechol lignan dehydroxylase | *cldh* | *G. pamelae* 3C | A0A369M2I8 | RDB65137.1 |

**Table S12.**
Accession numbers (Uniprot and Genbank) of putative *Eggerthella* and *Gordonibacter* dehydroxylases identified in this study.

| **Uniprot accession #** |
|---|
| Q93PD2_9RHOO |
| AAQ01672 |
| AAU11839 |
| AAU11840 |
| Q5NZV2_AROAE |
| AAZ43099 |
| A0YDJ9_9GAMM |
| A7LI70_9RHOO |
| A8ZZM3_DESOH |
| ZP_01288441 |
| ZP_01288668 |
| ZP_01667237 |
| ZP_03045699 |
| ZP_02830247 |
| E9NQE6_9RHOO |
| F7V8S9_CLOSS |
| ABB51928 |
| YP_091657 |
| AHY_PELAE |
| ZP_01034989 |
| PCRA_DECAR |
| AAW37220 |
| A0A096LIK4_9ACTN |
| NP_071207 |
| NP_560168 |
| NP_560307 |
| NP_560860 |
| YP_004130 |
| YP_055228 |
| YP_076161 |
| YP_217061 |
| YP_252569 |
| YP_263899 |
| YP_361367 |
| YP_387178 |
| YP_435153 |
| YP_583300 |
| YP_638431 |

| |
|---|
| YP_682694 |
| YP_741477 |
| YP_743256 |
| YP_840970 |
| YP_916609 |
| YP_920808 |
| YP_931484 |
| YP_964317 |
| YP_001002743 |
| YP_001013239 |
| YP_001020903 |
| YP_001055297 |
| YP_001056256 |
| YP_001056789 |
| YP_001152746 |
| YP_001153186 |
| YP_001157040 |
| YP_001233491 |
| YP_001408699 |
| YP_001409193 |
| YP_001562139 |
| YP_001585636 |
| YP_001685488 |
| YP_001951391 |
| YP_001951406 |
| YP_002016790 |
| YP_002129528 |
| YP_002457721 |
| NP_147849 |
| NP_902213 |
| NP_906381 |
| NP_906934 |
| NP_906980 |
| NP_907333 |
| NP_907591 |
| YP_310854 |
| YP_466957 |
| YP_524035 |
| YP_524325 |

| |
|---|
| YP_571843 |
| YP_871247 |
| YP_001318866 |
| YP_001319191 |
| YP_001800219 |
| YP_001942454 |
| YP_002734116 |
| YP_002796487 |
| YP_002828931 |
| YP_002882810 |
| YP_002949599 |
| YP_003099832 |
| YP_003151500 |
| YP_003161194 |
| YP_003182037 |
| YP_003303433 |
| YP_003303806 |
| YP_003311189 |
| YP_003381996 |
| YP_003393294 |
| YP_003863182 |
| YP_001099944 |
| YP_001423003 |
| YP_001632440 |
| YP_003699297 |
| YP_001753784 |
| A0A0F7JXH0_9GAMM |
| CAM58792 |
| A0A0S8CQF4_9BACT |
| A0A160FRG8_9BURK |
| AAU11841 |
| AAQ19491 |
| ABP63660 |
| NP_213709 |
| NP_951834 |
| YP_001634827 |
| NP_415742 |
| NP_415991 |
| EAT99379 |

| |
|---|
| A0A1C5V9P8_9FIRM |
| A0A1E4LUP6_9BURK |
| A0A1F2WFM2_9ACTN |
| A0A1F8PG51_9CHLR |
| A0A1M3JW53_9BURK |
| A0A1M6E5L2_9FIRM |
| A0A1M6FDE7_9FIRM |
| A0A1L3GSN8_9DELT |
| A0A1W6CT59_9PROT |
| YP_429324 |
| AAR05656 |
| NP_460720 |
| A0A1G6AWF4_EUBOX |
| A0A1Z8A857_9GAMM |
| AIOA_ALCFA |
| AIOA_HERAR |
| PGTL_PELAC |
| A0A2A4UZ40_9GAMM |
| A0A369NIV7 |
| A0A369MIX7 |
| C8WLG2 |
| A0A369LV65 |
| A0A369M2I8 |
| A0A369M6P4 |
| A0A369M3A8 |

**Table S13.**
Accession numbers of bis-MGD enzymes used to generate the phylogenetic tree of the bis-MGD enzyme family (Fig. 5A).

| Representative sequence | Organism | Uniprot accession # |
|---|---|---|
| 1 | *Gordonibacter pamelaeae* | A0A369M3J7 |
| 2 | *Gordonibacter pamelaeae* | A0A369M6U3 |
| 3 | *Desulfitobacterium metallireducens* | W0E998 |
| 4 | *Gordonibacter sp.* | A0A369LVG2 |
| 5 | *Clostridium ljungdahlii* | D8GM28 |
| 6 | *Eggerthella sp.* | F0HRV1 |
| 7 | *Eggerthella lenta* | C8WM38 |
| 8 | *Eggerthella lenta* | A0A369MIX7 |

| | | |
|---|---|---|
| 9 | *Eggerthella lenta* | A0A369NIV7 |
| 10 | *Streptomyces zelensis* | A0A1W6EUS9 |
| 11 | *Thauera aromatica* | A0A088SRF0 |
| 12 | *Gordonibacter pamelaeae* | A0A369LSI8 |
| 13 | *Gordonibacter pamelaeae* | D6E7S9 |
| 14 | *Denitrobacterium detoxificans* | A0A172RZZ3 |
| 15 | *Bradyrhizobium canariense* | A0A1H1VHY4 |
| 16 | *Thiocapsa roseopersicina* | X5FC28 |
| 17 | *Thioflavicoccus mobilis 8321* | L0GVW7 |
| 18 | *Ferroglobus placidus* | D3S147 |

**Table S14.**
Accession numbers (Uniprot) and organismal origin of representative dehydroxylases identified from phylogenetic tree in fig. S17.

**Part 2:** Materials, methods, references, and characterization data for synthesis of dopamine analogs

### General materials and methods

All reactions were performed in dried glassware under an atmosphere of dry $N_2$. Reaction mixtures were stirred magnetically unless otherwise indicated and monitored by thin layer chromatography (TLC) on Merck precoated glass-backed silica gel 60 F-254 0.25 mm plates with visualization by fluorescence quenching at 254 nm. TLC plates were stained using a potassium permanganate solution. Chromatographic purification of products (flash column chromatography) was performed on Silicycle Silica Flash F60 (230–400 Mesh) silica gel using a forced flow of eluent at 0.3–0.5 bar. Concentration of reaction product solutions and chromatography fractions under reduced pressure was performed by rotary evaporation at 35–40 °C at the appropriate pressure and then at rt, ca. 0.1 mmHg (vacuum pump) unless otherwise indicated.

All chemicals were purchased from Acros, Aldrich, Fluka, Merck, ABCR, TCI, Alfa Aesar or Strem and used as such unless stated otherwise. Commercial grade reagents and solvents were used without further purification except as indicated below. Toluene, diethylether ($Et_2O$), tetrahydrofuran (THF) and dichloromethane ($CH_2Cl_2$) were purified by pressure filtration through activated alumina. *N*,*N*-Dimethylformamide (DMF), acetonitrile ($CH_3CN$), and ethanol (EtOH) were used as purchased. Yields given refer to chromatographically purified and spectroscopically pure compounds unless otherwise stated.

Infrared (IR) spectra were recorded on a Bruker ALPHA FT-IR spectrophotometer and reported as wavenumber ($cm^{-1}$) of the absorption maxima for the range between 4000 $cm^{-1}$ and 750 $cm^{-1}$ with only major peaks reported. $^1H$ NMR and $^{13}C$ NMR spectra were recorded on a Varian-Inova-500 500 MHz, 125 MHz spectrometer. $^1H$ NMR chemical shifts are expressed in parts per million (δ) downfield from tetramethylsilane (with the $CHCl_3$ peak at 7.26 ppm, MeOH peak at 3.31, DMSO peak at 2.50, and acetone peak at 2.05 used as a standard). $^{13}C$ NMR chemical shifts are expressed in parts per million (δ) downfield from tetramethylsilane (with the central peak of $CHCl_3$ at 77.16 ppm, MeOH peak 49.00, DMSO peak at 39.52, and acetone peak at 29.84 used as a standard). All $^{13}C$ spectra were measured with complete proton decoupling. NMR coupling constants (J) are reported in Hertz (Hz), and splitting patterns are indicated as follows: br, broad; s, singlet; d, doublet; dd, doublet of doublet; t, triplet; m, multiplet. High-resolution mass spectrometric measurements (HRMS) were performed on an Accurate-Mass 6530 Q-TOF LC/MS (Agilent) using dual electrospray ionization (ESI).

### PREPARATION OF 2-AMINOETHYLBENZENEDIOL/-TRIOL DERIVATIVES

All reactions were carried out with degassed solvents under a positive pressure of nitrogen.

### 2-PHENYLACETIC ACID DERIVATIVES AS STARTING MATERIALS

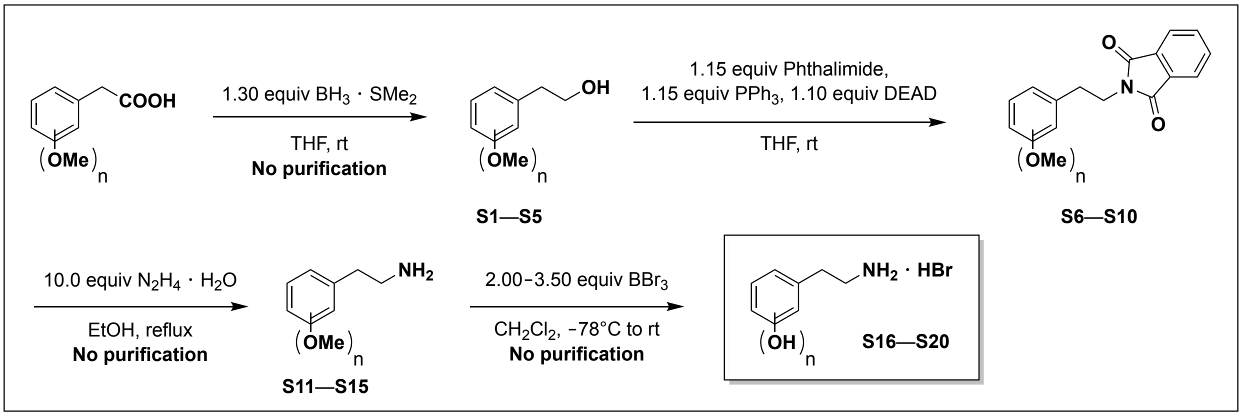

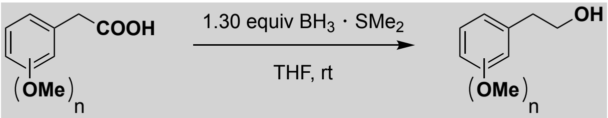

General C'Acid Reduction Procedure: $BH_3 \cdot SMe_2$ (2.0 M in THF; 1.30 equiv) was added dropwise to a solution of commercially available di- / trimethoxyphenylacetic acid (1.00 equiv) in THF (0.25 M) at 0 °C. The resulting mixture was allowed to warm to rt over 3 h and stirring was continued for 14 h while a colorless solid formed. The obtained suspension was cooled to 0 °C and carefully quenched with the dropwise addition of saturated aqueous $NaHCO_3$. The layers were separated, and the aqueous layer was extracted with EtOAc (3 x 50 mL). The combined organic layers were washed with $H_2O$ (2 x 20 mL), brine (2 x 20 mL), dried over anhydrous $Na_2SO_4$, filtered, and concentrated under reduced pressure to yield analytically pure alcohol **S1**–**S5** that was used in the next step without further purification.

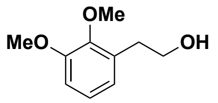

**2-(2,3-Dimethoxyphenyl)ethan-1-ol (S1).** Following the general carboxylic acid reduction procedure using 2-(2,3-dimethoxyphenyl)acetic acid (2.00 g, 10.2 mmol), alcohol **S1** was obtained as a colorless oil (1.85 g, quant.). $^1$H NMR (500 MHz, $CDCl_3$): δ 7.01 (t, *J* = 7.9 Hz, 1H), 6.88 – 6.75 (m, 2H), 3.87 (s, 3H), 3.84 (s, 3H), 3.84 (t, *J* = 6.5 Hz, 2H), 2.91 (t, *J* = 6.5 Hz, 2H), 1.76 (br s, OH); $^{13}$C NMR (125 MHz, $CDCl_3$): δ 152.8, 147.4, 132.6, 124.2, 122.6, 111.0, 63.4, 60.7, 55.7, 33.8. The spectral characteristics were identical to those reported in the current literature, which fails to report the signals for the OMe and one of the $CH_2$ groups.[1]

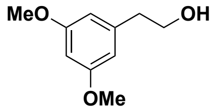

**2-(3,5-Dimethoxyphenyl)ethan-1-ol (S2)**. Following the general carboxylic acid reduction procedure using 2-(3,5-dimethoxyphenyl)acetic acid (1.00 g, 5.10 mmol), alcohol **S2** was obtained as a colorless oil (920 mg, quant.). $^1$H NMR (500 MHz, $CDCl_3$): δ 6.39 (d, *J* = 2.4 Hz, 2H), 6.36 – 6.33 (m, 1H), 3.86 (t, *J* = 6.4 Hz, 2H), 3.79 (s, 6H), 2.82 (t, *J* = 6.4 Hz, 2H), 1.45 (br s, OH). The spectral characteristics were identical to those reported in the current literature.[2]

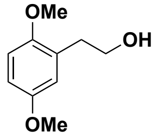

**2-(2,5-Dimethoxyphenyl)ethan-1-ol (S3)**. Following the general carboxylic acid reduction procedure using 2-(2,5-dimethoxyphenyl)acetic acid (2.00 g, 10.2 mmol), alcohol **S3** was obtained as a colorless oil (1.70 g, 92% yield). $^1$H NMR (500 MHz, $CDCl_3$): δ 6.80 (d, *J* = 8.6 Hz, 1H), 6.77 – 6.71 (m, 2H), 3.83 (t, *J* = 6.4 Hz, 2H), 3.79 (s, 3H), 3.76 (s, 3H), 2.88 (t, *J* = 6.4 Hz, 2H), 1.72 (br s, OH). The spectral characteristics were identical to those reported in the current literature.[3]

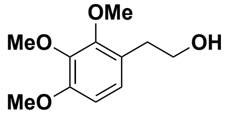

**2-(2,3,4-Trimethoxyphenyl)ethan-1-ol (S4)**. Following the general carboxylic acid reduction procedure using 2-(2,3,4-trimethoxyphenyl)acetic acid (2.00 g, 8.84 mmol), alcohol **S4** was obtained as a colorless oil (1.87 g, quant.). IR (thin film) ν 3348, 2926, 2850, 1769, 1658, 1602, 1498, 1395, 1091, 840 cm$^{-1}$; $^1$H NMR (500 MHz, $CDCl_3$): δ 6.86 (d, *J* = 8.4 Hz, 1H), 6.63 (d, *J* = 8.4 Hz, 1H), 3.90 (s, 3H), 3.87 (s, 3H), 3.84 (s, 3H), 3.80 (t, *J* = 6.4 Hz, 2H), 2.83 (t, *J* = 6.4 Hz, 2H), 1.79 (br s, OH); $^{13}$C NMR (125 MHz, $CDCl_3$): δ 152.6, 152.1, 142.4, 124.7, 124.7, 107.5, 63.5, 61.0, 60.8, 56.1, 33.6; ESI-HRMS calcd for $C_{11}H_{17}O_4$ [M+H] 213.1121, found 213.1128.

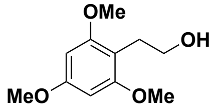

**2-(2,4,6-Trimethoxyphenyl)ethan-1-ol (S5)**. Following the general carboxylic acid reduction procedure using 2-(2,4,6-trimethoxyphenyl)acetic acid (1.00 g, 4.42 mmol), alcohol **S5** was obtained as a colorless oil (940 mg, quant.). $^{1}$H NMR (500 MHz, $CDCl_3$): δ 6.14 (s, 2H), 3.80 (s, 3H), 3.80 (s, 6H), 3.71 (t, *J* = 6.4 Hz, 3H), 2.89 (t, *J* = 6.4 Hz, 2H), 1.96 (br s, OH); $^{13}$C NMR (125 MHz, $CDCl_3$): δ 159.8, 159.2, 107.6, 90.7, 63.1, 55.8, 55.5, 26.2. The spectral characteristics were identical to those reported in the current literature.[4]

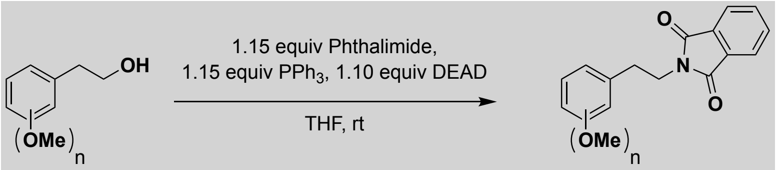

General Mitsunobu Reaction Protocol: Diethyl azodicarboxylate (DEAD; 40% in toluene; 1.10 equiv) was added dropwise over 5–10 min to a solution of $PPh_3$ (1.15 equiv), phthalimide (1.15 equiv) and the corresponding alcohol **S1**–**S5** (1.00 equiv) in THF (0.15 M) at 0 °C. The resulting mixture was allowed to warm to rt over 3 h and stirring was continued for 14 h. The resulting pale-yellow solution was concentrated under reduced pressure and purified by flash column chromatography to afford the desired phthalimide protected amine **S6**–**S10**.

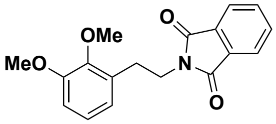

**2-(2,3-Dimethoxyphenethyl)isoindoline-1,3-dione (S6)**. Following the general Mitsunobu reaction protocol, purification by flash column chromatography (hexanes:EtOAc 5:1) afforded phthalimide protected amine **S6** as a colorless solid (2.90 g, 85% yield) using alcohol **S1** (2.00 g, 11.0 mmol) as starting material. $^1$H NMR (500 MHz, $CDCl_3$): δ 7.85 – 7.78 (m, 2H), 7.73 – 7.66 (m, 2H), 6.94 (t, $J$ = 7.9 Hz, 1H), 6.83 – 6.74 (m, 2H), 3.95 – 3.91 (m, 2H), 3.89 (s, 3H), 3.83 (s, 3H), 3.07 – 2.97 (m, 2H); $^{13}$C NMR (125 MHz, $CDCl_3$): δ 168.2, 152.8, 147.7, 133.8, 132.2, 131.9, 123.8, 123.1, 122.3, 111.3, 60.8, 55.7, 38.5, 29.1. The spectral characteristics were identical to those reported in the current literature.[5]

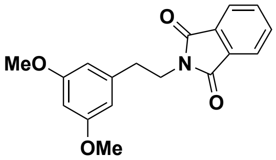

**2-(3,5-Dimethoxyphenethyl)isoindoline-1,3-dione (S7)**. Following the general Mitsunobu reaction protocol, purification by flash column chromatography (hexanes:EtOAc 3:1) afforded phthalimide protected amine **S7** as a colorless solid (1.70 g, 99% yield) using alcohol **S2** (1.00 g, 5.50 mmol) as starting material. $^1$H NMR (500 MHz, $CDCl_3$): δ 7.84 (dd, $J$ = 5.4, 3.1 Hz, 2H), 7.71 (dd, $J$ = 5.4, 3.1 Hz, 2H), 6.45 – 6.38 (m, 2H), 6.32 (t, $J$ = 2.2 Hz, 1H), 3.96 – 3.89 (m, 2H), 3.75 (s, 6H), 2.96 – 2.91 (m, 2H); $^{13}$C NMR (125 MHz, $CDCl_3$): δ 168.3, 161.0, 140.4, 134.0, 132.2, 123.3, 106.8, 99.0, 55.4, 39.2, 35.0; $R_f$ = 0.33 (hexanes:EtOAc 3:1). The spectral characteristics were identical to those reported in the current literature.[5]

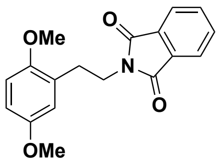

**2-(2,5-Dimethoxyphenethyl)isoindoline-1,3-dione (S8)**. Following the general Mitsunobu reaction protocol, purification by flash column chromatography (hexanes:EtOAc 5:1) afforded phthalimide protected amine **S8** as a colorless solid (3.15 g, 98% yield) using alcohol **S3** (1.85 g, 10.2 mmol) as starting material. $^{1}H$ NMR (500 MHz, $CDCl_3$): δ 7.79 – 7.71 (m, 2H), 7.68 – 7.59 (m, 2H), 6.70 – 6.63 (m, 3H), 3.90 (t, *J* = 7.1 Hz, 2H), 3.66 (s, 3H), 3.62 (s, 3H), 2.93 (t, *J* = 7.1 Hz, 2H); $^{13}C$ NMR (125 MHz, $CDCl_3$): δ 168.1, 153.3, 152.0, 133.7, 132.1, 127.6, 123.0, 116.6, 112.2, 111.1, 55.7, 55.6, 37.8, 29.6; $R_f$ = 0.28 (hexanes:EtOAc 5:1). The spectral characteristics were identical to those reported in the current literature.[5b]

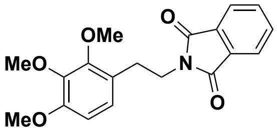

**2-(2,3,4-Trimethoxyphenethyl)isoindoline-1,3-dione (S9)**. Following the general Mitsunobu reaction protocol, purification by flash column chromatography (hexanes:EtOAc 5:1) afforded phthalimide protected amine **S9** as a colorless solid (1.55 g, 96% yield) using alcohol **S4** (1.00 g, 4.71 mmol) as starting material. $^{1}H$ NMR (500 MHz, $CD_3OD$): δ 7.85 – 7.70 (m, 4H), 6.78 (d, *J* = 8.5 Hz, 1H), 6.60 (d, *J* = 8.5 Hz, 1H), 3.86 (t, *J* = 6.8 Hz, 2H), 3.82 (s, 3H), 3.77 (s, 3H), 3.65 (s, 3H), 2.93 – 2.87 (m, 2H); $^{13}C$ NMR (125 MHz, $CD_3OD$): δ 169.7, 154.1, 153.5, 143.4, 135.3, 135.2, 126.0, 124.1, 123.9, 108.7, 61.3, 60.9, 56.4, 39.7, 29.8; $R_f$ = 0.18 (hexanes:EtOAc 3:1). The spectral characteristics were identical to those reported in the current literature.[5]

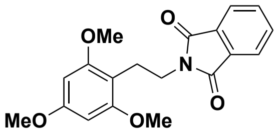

**2-(2,4,6-Trimethoxyphenethyl)isoindoline-1,3-dione (S10)**. Following the general Mitsunobu reaction protocol, purification by flash column chromatography (hexanes:EtOAc 3:1) afforded phthalimide protected amine **S9** as a colorless solid (1.60 g, 99% yield) using alcohol **S5** (1.00 g, 4.71 mmol) as starting material. IR (thin film) ν 3248, 2928, 2850, 1772, 1715, 1634, 1607, 1439, 1205, 1139, 809 $cm^{-1}$; $^{1}H$ NMR (500 MHz, $CDCl_3$): δ 7.84 – 7.73 (m, 2H), 7.71 – 7.62 (m, 2H), 6.00 (s, 2H), 3.85 (t, *J* = 6.3 Hz, 2H), 3.76 (s, 3H), 3.59 (s, 6H), 2.97 (t, *J* = 6.3 Hz, 2H); $^{13}C$ NMR (125 MHz, $CDCl_3$): δ 168.4, 160.0, 159.2, 133.6, 132.5, 122.9, 107.4, 90.2, 55.5, 55.3, 37.6, 21.6; $R_f$ = 0.20 (hexanes:EtOAc 3:1); ESI-HRMS calcd for $C_{18}H_{18}NO_4$ [M+H] 312.1230, found 312.1223.

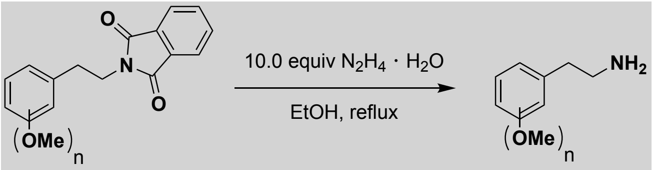

<u>General Amine Phthalimide Deprotection Protocol:</u> Hydrazine monohydrate (10.0 equiv) was added to a suspension of the phthalimide protected amine **S6 – S10** in EtOH (0.15 M). The resulting solution was heated to reflux for 1.5 h while colorless solids crashed out. The resulting suspension was allowed to cool to rt before $H_2O$ (20–50 mL) was added in one portion. Stirring was continued to afford a clear solution that was extracted with EtOAc (3 x 50 mL). The combined organic layers were washed with $H_2O$ (3 x 20 mL), brine (2 x 20 mL), dried over anhydrous

$Na_2SO_4$, filtered and concentrated under reduced pressure to yield analytically pure amine **S11**–**S15** that was used in the next step without further purification.

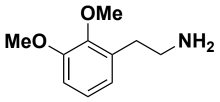

**2-(2,3-Dimethoxyphenyl)ethan-1-amine (S11)**. Following the general phthalimide deprotection protocol using phthalimide protected amine **S6** (1.00 g, 3.21 mmol), primary amine **S11** was obtained as a colorless oil (500 mg, 86% yield). $^1$H NMR (500 MHz, $CD_3OD$): δ 6.99 (dd, *J* = 8.0, 7.7 Hz, 1H), 6.88 (dd, *J* = 8.0, 1.5 Hz, 1H), 6.78 (dd, *J* = 7.7, 1.6 Hz, 1H), 3.84 (s, 3H), 3.79 (s, 3H), 2.88 – 2.80 (m, 2H), 2.80 – 2.73 (m, 2H); $NH_2$-group is not visible; $^{13}$C NMR (125 MHz, $CD_3OD$): δ 154.2, 148.6, 134.3, 125.1, 123.4, 112.1, 61.0, 56.2, 43.5, 34.5. The spectral characteristics were identical to those reported in the current literature.[6]

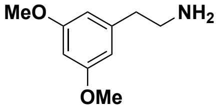

**2-(3,5-Dimethoxyphenyl)ethan-1-amine (S12)**. Following the general phthalimide deprotection protocol using phthalimide protected amine **S7** (400 mg, 1.28 mmol), primary amine **S12** was obtained as a colorless oil (140 mg, 60% yield). $^1$H NMR (500 MHz, $CD_3OD$): δ 6.41 – 6.36 (m, 2H), 6.35 (t, *J* = 2.3 Hz, 1H), 3.76 (s, 6H), 2.92 (t, *J* = 7.3 Hz, 2H), 2.72 (t, *J* = 7.3 Hz, 2H), $NH_2$-group is not visible; $^{13}$C NMR (125 MHz, $CD_3OD$): δ 162.5, 142.6, 107.8, 99.3, 55.7, 43.6, 39.4. The spectral characteristics were identical to those reported in the current literature.[7]

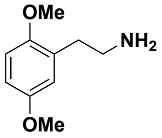

**2-(2,5-Dimethoxyphenyl)ethan-1-amine (S13)**. Following the general phthalimide deprotection protocol using phthalimide protected amine **S8** (500 mg, 1.61 mmol), primary amine **S13** was obtained as a colorless oil (250 mg, 86% yield). $^1$H NMR (500 MHz, $CDCl_3$): δ 6.96 – 6.57 (m, 3H), 3.78 (s, 3H), 3.76 (s, 3H), 3.04 – 2.85 (m, 3H), 2.75 (t, *J* = 6.9 Hz, 2H), 1.90 (br s, $NH_2$); $^{13}$C NMR (125 MHz, $CDCl_3$): δ 153.5, 152.0, 129.2, 117.0, 111.4, 111.3, 55.9, 55.7, 42.1, 34.6. The spectral characteristics were identical to those reported in the current literature.[8]

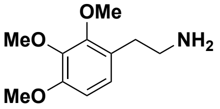

**2-(2,3,4-Trimethoxyphenyl)ethan-1-amine (S14)**. Following the general phthalimide deprotection protocol using phthalimide protected amine **S9** (1.40 g, 4.10 mmol), primary amine **S14** was obtained as a colorless oil (600 mg, 69% yield). $^1$H NMR (500 MHz, $CDCl_3$): δ 6.83 (d, *J* = 8.5 Hz, 1H), 6.61 (d, *J* = 8.5 Hz, 1H), 3.87 (s, 3H), 3.86 (s, 3H), 3.83 (s, 3H), 2.90 (t, *J* = 7.0 Hz, 2H), 2.70 (t, *J* = 7.0 Hz, 2H), 1.51 (br s, $NH_2$); $^{13}$C NMR (125 MHz, $CDCl_3$): δ 152.4, 152.2, 142.5, 125.8, 124.5, 107.3, 61.1, 60.8, 56.1, 43.1, 34.2. The spectral characteristics were identical to those reported in the current literature.[6]

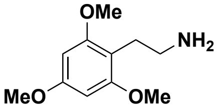

**2-(2,4,6-Trimethoxyphenyl)ethan-1-amine (S15)**. Following the general phthalimide deprotection protocol using phthalimide protected amine **S10** (1.60 g, 4.69 mmol), primary amine **S15** was obtained as a colorless oil (693 mg, 70% yield). IR (thin film) ν 3426, 2938, 2838, 1593, 1498, 1455, 1417, 1204, 1148 $cm^{-1}$; $^1$H NMR (500 MHz, $CD_3OD$): δ 6.20 (s, 2H), 3.79 (s, 6H), 3.79 (s, 3H), 2.79 – 2.71 (m, 4H), $NH_2$-group is not visible; $^{13}$C NMR (125 MHz, $CD_3OD$): δ 161.5, 160.4, 108.1, 91.5, 56.0, 56.0, 55.7, 41.9, 25.8; ESI-HRMS calcd for $C_{11}H_{18}NO_3$ [M+H] 212.1281, found 212.1280.

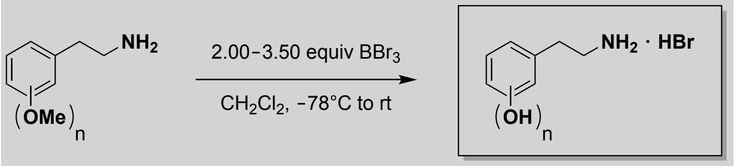

General Phenol Ether Cleavage Protocol: $BBr_3$ (1.0 M in $CH_2Cl_2$; 1.15 equiv for each OMe group) was added dropwise to a solution of the 2-(methoxyphenyl)ethyl amine **S11**–**S15** (1.00 equiv) in $CH_2Cl_2$ (0.025 M) at –78 °C. The resulting mixture was allowed to warm to rt over 3 h. Stirring was continued for 14 h. The resulting suspension was cooled to 0 °C and quenched with the dropwise addition of MeOH (ca. 5 mL). Stirring at rt was continued for 1 h. The resulting suspension was concentrated under reduced pressure to afford a pale-brown oil. The obtained residue was dissolved in a small amount of MeOH and again concentrated under reduced pressure; this step was repeated 3–4 times to remove all of the trimethyl borate side product and obtain analytically pure 2-(2-aminoethyl)benzenediol /-triol derivatives **S16**–**S20** as HBr salts.

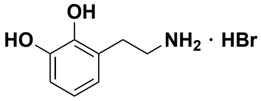

**3-(2-Aminoethyl)benzene-1,2-diol hydrobromide (S16)**. Following the general phenol ether cleavage protocol using phenol ether **S11** (200 mg, 1.10 mmol), catechol amine **S16** was obtained as a brown oil (250 mg, 97% yield). $^1$H NMR (500 MHz, $CD_3OD$): δ 6.73 (dd, $J$ = 6.7, 2.7 Hz, 1H), 6.69 – 6.56 (m, 2H), 3.17 (t, $J$ = 7.4 Hz, 2H), 2.95 (t, $J$ = 7.4 Hz, 2H), OH- and NH-protons are not visible; $^{13}$C NMR (125 MHz, $CD_3OD$): δ 146.0, 144.6, 124.4, 122.3, 120.8, 115.4, 40.8, 29.4. The spectral characteristics were identical to those reported in the current literature.[9]

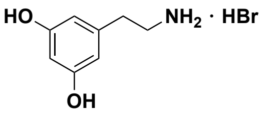

**5-(2-Aminoethyl)benzene-1,3-diol hydrobromide (S17)**. Following the general phenol ether cleavage protocol using phenol ether **S12** (20 mg, 0.110 mmol), resorcinol amine **S17** was obtained as a brown oil (25.0 mg, 97% yield). IR (thin film) ν 3358, 2928, 2853, 1771, 1597, 1495, 1418, 1091, 844 $cm^{-1}$; $^1$H NMR (500 MHz, $CD_3OD$): δ 6.33 – 6.08 (m, 2H), 3.13 (t, $J$ = 7.6 Hz, 2H),

2.80 (t, $J$ = 7.6 Hz, 2H), OH- and NH-protons are not visible; $^{13}$C NMR (125 MHz, $CD_3OD$): δ 160.0, 139.9, 108.1, 102.4, 41.9, 34.5; ESI-HRMS calcd for $C_8H_{12}NO_2$ [M+H] 154.0863, found 154.0860.

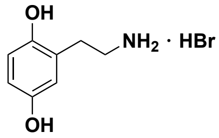

**5-(2-Aminoethyl)benzene-1,4-diol hydrobromide (S18)**. Following the general phenol ether cleavage protocol using phenol ether **S13** (100 mg, 0.552 mmol), diol amine **S18** was obtained as a brown oil (118 mg, 91% yield). IR (thin film) ν 3352, 2927, 2858, 1621, 1505, 1455, 1344, 1202, 1152, 1212 $cm^{-1}$; $^{1}$H NMR (500 MHz, $CD_3OD$): δ 6.66 (d, $J$ = 8.5 Hz, 1H), 6.63 – 6.46 (m, 2H), 3.15 (t, $J$ = 7.2 Hz, 2H), 2.88 (t, $J$ = 7.2 Hz, 2H), OH- and NH-protons are not visible; $^{13}$C NMR (125 MHz, $CD_3OD$): δ 151.1, 149.4, 124.9, 118.2, 116.9, 115.8, 40.9, 29.9; ESI-HRMS calcd for $C_8H_{12}NO_2$ [M+H] 154.0863, found 154.0852.

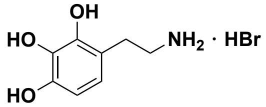

**4-(2-Aminoethyl)benzene-1,2,3-triol hydrobromide (S19)**. Following the general phenol ether cleavage protocol using phenol ether **S14** (200 mg, 0.948 mmol), triol amine **S19** was obtained as a brown oil (240 mg, quant.). IR (thin film) ν 3357, 3222, 2537, 1620, 1484, 1282, 1230, 1182, 1100, 1053, 1016 $cm^{-1}$; $^{1}$H NMR (500 MHz, $CD_3OD$): δ 6.48 (d, $J$ = 8.2 Hz, 1H), 6.32 (d, $J$ = 8.2 Hz, 1H), 3.12 (t, $J$ = 7.3 Hz, 2H), 2.87 (t, $J$ = 7.3 Hz, 2H), OH- and NH-protons are not visible; $^{13}$C NMR (125 MHz, $CD_3OD$): δ 146.6, 145.7, 134.4, 121.4, 115.9, 108.0, 41.2, 29.5; ESI-HRMS calcd for $C_8H_{12}NO_3$ [M+H] 170.0812, found 170.0805.

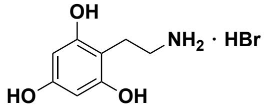

**2-(2-Aminoethyl)benzene-1,3,5-triol hydrobromide (S20)**. Following the general phenol ether cleavage protocol using phenol ether **S15**, triol amine **S20** was obtained, according to MS identification, in low quantities along with brominated species and various methoxybenzene-diols in an inseparable mixture.

Initial attempts to alter reaction temperature or the number of equivalents of $BBr_3$ resulted in low conversion. Heating phenol ether **S15** in the presence of iodo(trimethyl)silane[10] or sodium ethanethiolate[11] afforded mono-deprotected material in a cleaner reaction, but the desired triol amine **S20** was not observed. Due to our inability to access this substrate, we did not evaluate this substrate in any enzyme reactions in our study.

BENZALDEHYDES AS STARTING MATERIALS

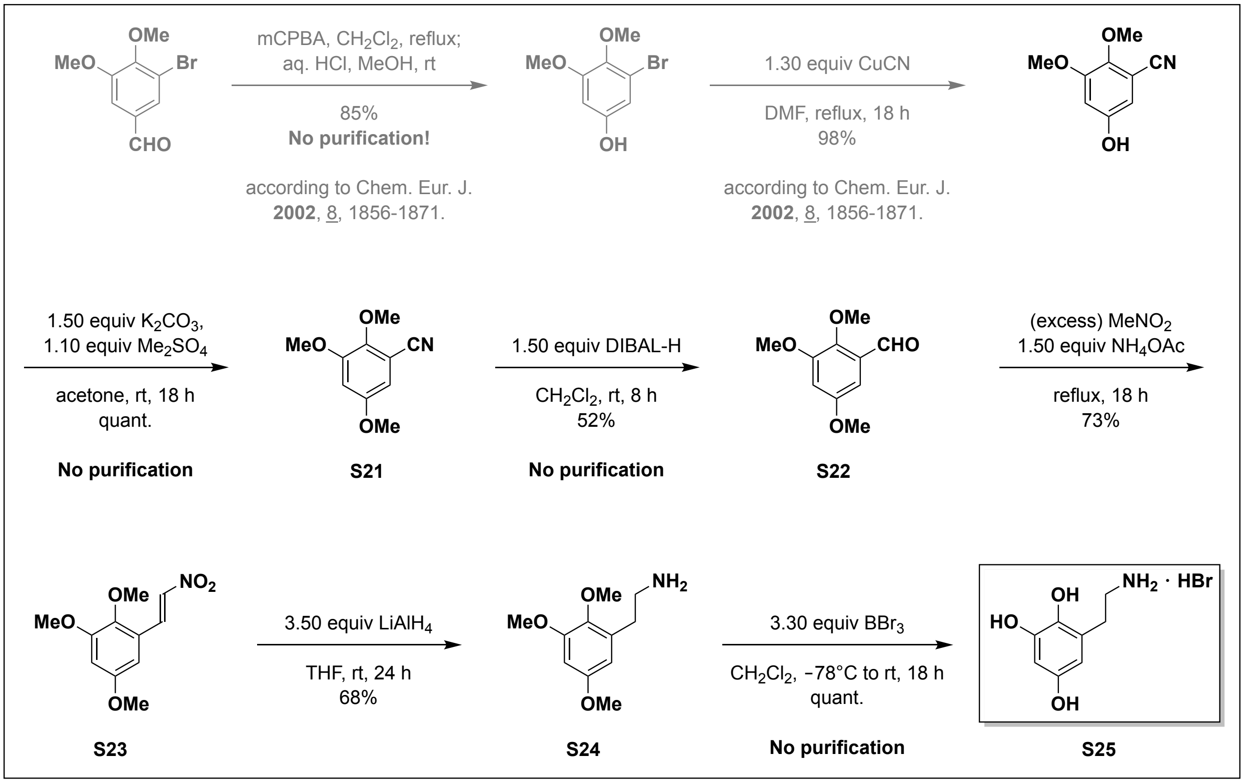

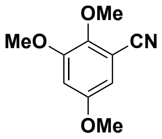

**2,3,5-Trimethoxybenzonitrile (S21)**. $K_2CO_3$ (1.90 g, 13.8 mmol, 1.50 equiv) and dimethyl sulfate (0.960 mL, 1.28 g, 10.1 mmol, 1.10 equiv) were added to a solution of 5-hydroxy-2,3-dimethoxybenzonitrile[12] (1.65 g, 9.21 mmol, 1.00 equiv) in acetone (30 mL) at rt. Stirring was continued for 18 h to afford a pale beige suspension. The solvent was removed under reduced pressure and the resulting crude material was diluted with a mixture of EtOAc–$H_2O$ (1:1; 100 mL). The obtained layers were separated and the aqueous layer was extracted with EtOAc (3 x 20 mL). The combined organic layers were washed with 5% aqueous NaOH (20 mL) and brine (2 x 20 mL), dried over anhydrous $Na_2SO_4$, filtered, and concentrated under reduced pressure to afford analytically pure trimethoxybenzonitrile **S21** (1.78 g, quant.) as a pale beige solid. $^1$H NMR (500 MHz, $CDCl_3$): δ 6.68 (d, $J$ = 2.8 Hz, 1H), 6.56 (d, $J$ = 2.8 Hz, 1H), 3.94 (s, 3H), 3.86 (s, 3H), 3.79 (s, 3H). The spectral characteristics were identical to those reported in the current literature.[13]

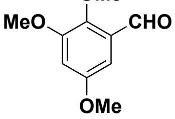

**2,3,5-Trimethoxybenzaldehyde (S22)**. DIBAL-H (1.0 M in $CH_2Cl_2$; 12.4 mL, 12.4 mmol, 1.50 equiv) was added dropwise to a solution of nitrile **S21** (1.60 g, 8.28 mmol, 1.00 equiv) in $CH_2Cl_2$ (33 mL) at 0 °C. The resulting mixture was allowed to warm to rt over 3 h after which stirring was continued for 8 h. The reaction was cooled to 0 °C and HCl (1.0 M in $H_2O$; 10.0 mL) was added dropwise over 10 min. The mixture was allowed to warm to rt and stirring was continued for 2 h. The layers were separated, and the aqueous layer was extracted with $CH_2Cl_2$ (2 x 15 mL). The combined organic layers were washed with $H_2O$ (2 x 15 mL) and brine (20 mL), dried over anhydrous $Na_2SO_4$, filtered, and concentrated under reduced pressure to afford analytically pure benzaldehyde **S22** (845 mg, 52%) as a beige solid. $^1H$ NMR (500 MHz, $CDCl_3$): δ 10.40 (s, 1H), 6.86 (d, *J* = 2.9 Hz, 1H), 6.74 (d, *J* = 2.9 Hz, 1H), 3.93 (s, 3H), 3.89 (s, 3H), 3.82 (s, 3H); $^{13}C$ NMR (125 MHz, $CDCl_3$): δ 190.2, 156.4, 154.4, 148.2, 129.9, 107.7, 99.7, 63.1, 56.4, 56.1. The spectral characteristics were identical to those reported in the current literature.[14]

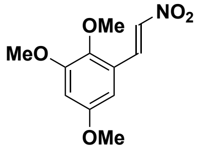

**(*E*)-1,2,5-Trimethoxy-3-(2-nitrovinyl)benzene (S23)**. A mixture of benzaldehyde **S22** (845 mg, 4.31 mmol, 1.00 equiv) and ammonium acetate (500 mg, 6.46 mmol, 1.50 equiv) in nitromethane (40 mL) was heated to reflux for 18 h, after which the reaction was found to be complete according to TLC ($R_f$ = 0.38 starting material; $R_f$ = 0.35 product; hexanes:EtOAc 4:1). The resulting mixture was concentrated under reduced pressure and purified by flash column chromatography (hexanes:EtOAc 6:1) to afford nitrovinyl benzene **S23** (750 mg, 73%) as a yellow solid. IR (thin film) ν 2959, 2846, 1717, 1633, 1601, 1492, 1465, 1332, 1282, 1206, 1176, 1151 $cm^{-1}$; $^1H$ NMR (500 MHz, $CDCl_3$): δ 8.16 (d, *J* = 13.8 Hz, 1H), 7.71 (d, *J* = 13.8 Hz, 1H), 6.61 (d, *J* = 2.6 Hz, 1H), 6.48 (d, *J* = 2.6 Hz, 1H), 3.86 (s, 3H), 3.83 (s, 3H), 3.80 (s, 3H); $^{13}C$ NMR (125 MHz, $CDCl_3$): δ 156.4, 154.1, 144.2, 138.7, 134.8, 124.0, 104.5, 102.9, 61.6, 56.1, 55.9; $R_f$ = 0.35 (hexanes:EtOAc 4:1); ESI-HRMS calcd for $C_{11}H_{14}NO_5$ [M+H] 240.0866, found 240.0871.

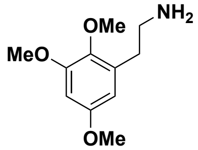

**2-(2,3,5-Trimethoxyphenyl)ethan-1-amine (S24)**. $LiAlH_4$ (2.0 M in THF; 1.83 mL, 3.66 mmol, 3.50 equiv) was added dropwise over 10 min to a solution of nitrovinyl benzene **S23** (250 mg, 1.05 mmol, 1.00 equiv) in THF (6 mL) at 0 °C. The resulting mixture was allowed to warm to rt and stirring was continued for 24 h. The reaction mixture was then cooled to 0 °C and 10% aqueous NaOH (5.0 mL) was added dropwise over 10 min, resulting in an exothermic reaction. Stirring was continued for 1 h, and the resulting suspension was diluted with EtOAc (20 mL) and filtered over a plug of Celite® (EtOAc rinse). The filtrate was dried over anhydrous $Na_2SO_4$, filtered, and concentrated under reduced pressure to afford crude amine **S24**. Purification by flash column chromatography (EtOAc:MeOH 85:15 + 0.1% $Et_3N$) afforded amine **S24** (150 mg, 68%) as a pale yellow oil. IR (thin film) ν 3363, 2937, 2839, 1599, 1492, 1465, 1427, 1380, 1220, 1175, 1150, 1089, 830 $cm^{-1}$; $^1H$ NMR (500 MHz, $CDCl_3$): δ 6.39 (d, *J* = 2.5 Hz, 1H), 6.29 (d, *J* = 2.5 Hz, 1H), 3.84 (s, 3H), 3.77 (s, 3H), 3.76 (s, 3H), 2.97 (t, *J* = 7.0 Hz, 2H), 2.78 (t, *J* = 7.0 Hz, 2H), 2.27 (br s, $NH_2$); $^{13}C$ NMR (125 MHz, $CDCl_3$): δ 156.0, 153.5, 141.5, 133.4, 105.5, 98.5, 60.9, 55.7, 55.6, 42.7, 34.1; $R_f$ = 0.08 (EtOAc:MeOH 85:15); ESI-HRMS calcd for $C_{11}H_{18}NO_3$ [M+H] 212.1281, found 212.1275.

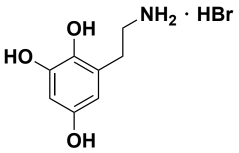

**3-(2-Aminoethyl)benzene-1,2,5-triol hydrobromide (S25)**. $BBr_3$ (1.0 M in $CH_2Cl_2$; 1.41 mL, 1.41 mmol, 3.30 equiv) was added dropwise over 10 min to a solution of phenol ether **S24** (90.0 mg, 0.425 mmol, 1.00 equiv) in $CH_2Cl_2$ (0.033 M) at –78 °C. The resulting mixture was allowed to warm to rt over 3 h. Stirring was continued for 18 h. The resulting suspension was cooled to 0 °C and quenched with the dropwise addition of MeOH (ca. 5 mL). Stirring at rt was continued for 1 h. The resulting solution was concentrated under reduced pressure to afford a pale-brown oil. The obtained residue was dissolved in a small amount of MeOH and again concentrated under reduced pressure; this step was repeated 3–4 times to remove all of the trimethyl borate side product and obtain analytically pure triol amine **S25** as the HBr salt. IR (thin film) ν 3358, 3223, 1604, 1452, 1359, 1291, 1108, 1044 $cm^{-1}$; $^1H$ NMR (500 MHz, $CD_3OD$): δ 6.29 (d, $J$ = 2.6 Hz, 1H), 6.11 (d, $J$ = 2.8 Hz, 1H), 3.15 (t, $J$ = 7.4 Hz, 2H), 2.88 (t, $J$ = 7.3 Hz, 2H), OH- and NH-protons are not visible; $^{13}C$ NMR (125 MHz, $CD_3OD$): δ 151.4, 147.1, 137.7, 125.0, 108.2, 103.3, 41.0, 29.8; ESI-HRMS calcd for $C_8H_{12}NO_3$ [M+H] 170.0812, found 170.0812.

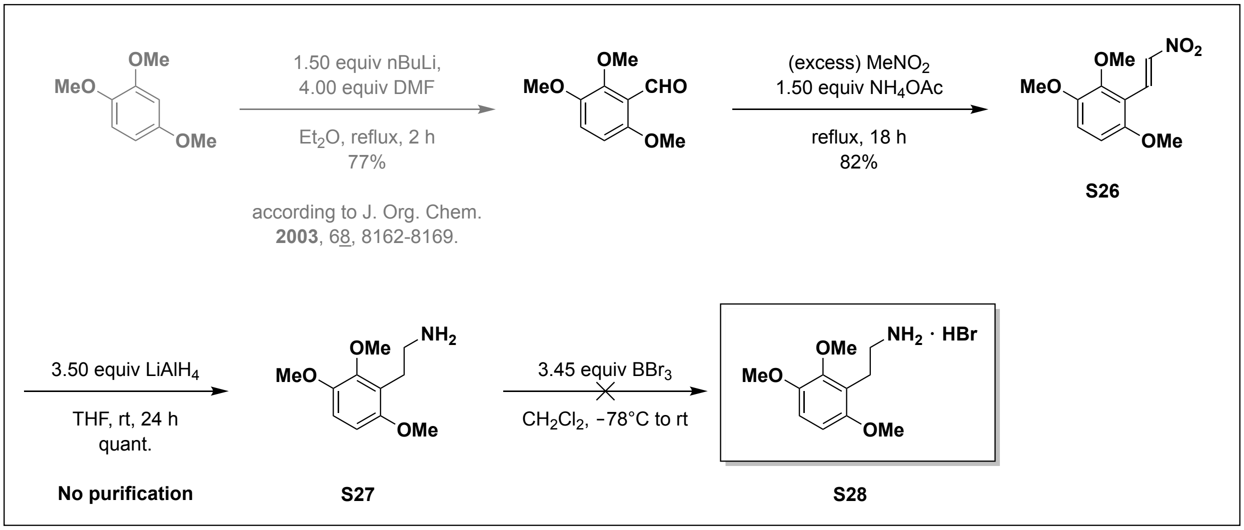

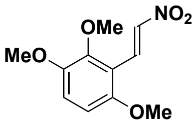

**(*E*)-1,2,4-Trimethoxy-3-(2-nitrovinyl)benzene (S26)**. A mixture of 2,3,6-trimethoxybenzaldehyde[15] (500 mg, 2.55 mmol, 1.00 equiv) and ammonium acetate (295 mg, 3.82 mmol, 1.50 equiv) in nitromethane (23 mL) was heated to reflux for 18 h after which the reaction was found to be according to TLC ($R_f$ = 0.30 starting material; $R_f$ = 0.38 product; hexanes:EtOAc

2:1). The resulting mixture was concentrated under reduced pressure and purified by flash column chromatography (hexanes:EtOAc 5:1) to afford nitrovinyl benzene **S26** (500 mg, 82%) as a yellow solid. IR (thin film) ν 2939, 2854, 1625, 1583, 1507, 1496, 1330, 1284, 1116, 1009 $cm^{-1}$; $^{1}H$ NMR (500 MHz, $CD_3OD$): δ 8.39 (d, *J* = 13.7 Hz, 1H), 8.09 (d, *J* = 13.7 Hz, 1H), 7.17 (d, *J* = 9.2 Hz, 1H), 6.80 (d, *J* = 9.2 Hz, 1H), 3.91 (s, 3H), 3.91 (s, 3H), 3.85 (s, 3H); $^{13}C$ NMR (125 MHz, $CD_3OD$): δ 155.5, 151.6, 148.2, 140.8, 130.7, 118.7, 114.7, 107.1, 61.6, 57.0, 56.6; $R_f$ = 0.38 (hexanes:EtOAc 2:1); ESI-HRMS calcd for $C_{11}H_{14}NO_5$ [M+H] 240.0866, found 240.0860.

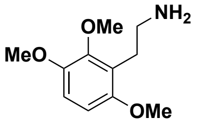

**2-(2,3,6-Trimethoxyphenyl)ethan-1-amine (S27)**. $LiAlH_4$ (2.0 M in THF; 3.66 mL, 7.32 mmol, 3.50 equiv) was added dropwise over 10 min to a solution of nitrovinyl benzene **S26** (200 mg, 2.09 mmol, 1.00 equiv) in THF (12 mL) at 0 °C. The resulting mixture was allowed to warm to rt and stirring was continued for 24 h. The reaction mixture was cooled to 0 °C and 10% aqueous NaOH (10.0 mL) was added dropwise over 10 min, resulting in an exothermic reaction. Stirring was stirred continued for 1 h. The resulting suspension was diluted with EtOAc (20 mL) and filtered over a plug of Celite® (EtOAc rinse). The filtrate was dried over anhydrous $Na_2SO_4$, filtered, and concentrated under reduced pressure to afford analytically pure amine **S27** (438 mg, quant.) as a pale yellow oil. IR (thin film) ν 3363, 2936, 2833, 1648, 1485, 1463, 1253, 1085, 793, 627 $cm^{-1}$; $^{1}H$ NMR (500 MHz, $CDCl_3$): δ 6.73 (d, *J* = 8.8 Hz, 1H), 6.55 (d, *J* = 8.8 Hz, 1H), 3.82 (s, 3H), 3.82 (s, 3H), 3.76 (s, 3H), 2.88 (d, *J* = 6.0 Hz, 2H), 2.83 (d, *J* = 6.0 Hz, 2H), 2.47 (br s, $NH_2$); $^{13}C$ NMR (125 MHz, $CDCl_3$): δ 152.5, 148.4, 147.2, 122.7, 110.2, 105.4, 60.8, 56.2, 55.9, 42.2, 28.5; $R_f$ = 0.10 (EtOAc:MeOH 85:15); ESI-HRMS calcd for $C_{11}H_{18}NO_3$ [M+H] 212.1281, found 212.1273.

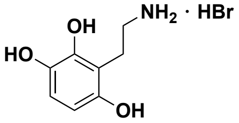

**3-(2-Aminoethyl)benzene-1,2,4-triol hydrobromide (S28)**. Following the general phenol ether cleavage protocol described for the preparation of amine hydrobromide **S25** using phenol ether **S27** as starting material, triol amine **S28** was obtained, according to MS identification, in low quantities along with brominated species. Initial attempts in changing the reaction temperature or the number of equivalents of $BBr_3$ resulted in low conversion and the desired product could not be isolated in pure form. Therefore it was not used in any enzyme assays,

### PREPARATION OF 2-AMINO-4-(2-AMINOETHYL)PHENOL

All reactions were carried out with degassed solvents under a positive pressure of nitrogen.

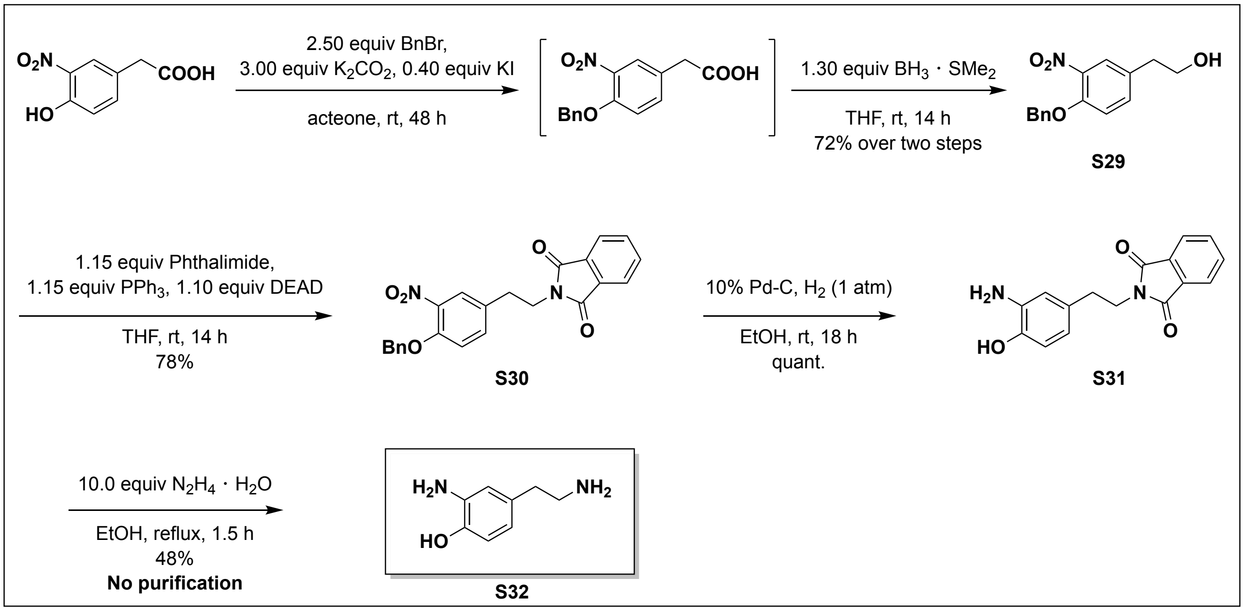

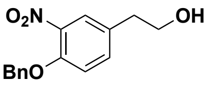

**2-(4-(Benzyloxy)-3-nitrophenyl)ethan-1-ol (S29)**. (Bromomethyl)benzene (3.02 mL, 4.34 g, 25.4 mmol, 2.50 equiv) was added dropwise to a suspension of commercially available 2-(4-hydroxy-3-nitrophenyl)acetic acid (2.00 g, 10.1 mmol, 1.00 equiv), anhydrous potassium carbonate (4.21 g, 30.4 mmol, 3.00 equiv), and anhydrous potassium iodide (674 mg, 4.06 mmol, 0.400 equiv) in acetone (34 mL) at rt. Vigorously stirring was continued for 48 h. The resulting suspension was diluted with a mixture of EtOAc-$H_2O$ (1:1; 100 mL), cooled to 0 °C, and adjusted to pH = 1 using aqueous 1M HCl. This resulted in an exothermic reaction. The layers were separated, and the aqueous layer was extracted with EtOAc (3 x 15 mL). The combined organic layers were washed with $H_2O$ (2 x 15 mL), brine (30 mL), dried over anhydrous $Na_2SO_4$, filtered, and concentrated under reduced pressure to afford crude 2-(4-(benzyloxy)-3-nitrophenyl)acetic acid, which was immediately used in the next step without further purification.

$BH_3$ • $SMe_2$ (2.0 M in THF; 6.57 mL, 13.1 mmol, 1.30 equiv) was added dropwise to a solution of the crude 2-(4-(benzyloxy)-3-nitrophenyl)acetic acid in THF (110 mL) at 0 °C. The resulting mixture was allowed to warm to rt over 3 h and stirring was continued for 14 h. The resulting suspension was cooled to 0 °C and carefully quenched with the dropwise addition of saturated aqueous $NaHCO_3$. The layers were separated, and the aqueous layer was extracted with EtOAc (3 x 50 mL). The combined organic layers were washed with $H_2O$ (2 x 20 mL), brine (2 x 20 mL), dried over anhydrous $Na_2SO_4$, filtered, and concentrated under reduced pressure to afford crude alcohol **S29** as a brown oil. Purification by flash column chromatography (hexanes:EtOAc 1:1) afforded analytically pure alcohol (1.98 g, 72%) as a pale yellow oil. $^1H$ NMR (500 MHz, $CDCl_3$): δ 7.71 (d, $J$ = 2.3 Hz, 1H), 7.46 – 7.29 (m, 6H), 7.05 (d, $J$ = 8.6 Hz, 1H), 5.19 (s, 2H), 3.82 (t, $J$ =

6.5 Hz, 2H), 2.81 (t, $J$ = 6.5 Hz, 2H), 1.90 (br s, OH); $^{13}$C NMR (125 MHz, $CDCl_3$): δ 150.5, 140.0, 135.8, 134.8, 131.8, 128.7, 128.2, 127.0, 125.9, 115.4, 71.3, 63.0, 37.7; $R_f$ = 0.18 (hexanes:EtOAc 1:1). The spectral characteristics were identical to those reported in the current literature.[16]

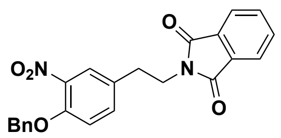

**2-(4-(Benzyloxy)-3-nitrophenethyl)isoindoline-1,3-dione (S30)**. Diethyl azodicarboxylate (DEAD; 40% in toluene; 3.95 mL, 8.05 mmol, 1.10 equiv) was added dropwise over 10 min to a solution of $PPh_3$ (2.21 g, 8.42 mmol, 1.15 equiv), phthalimide (1.24 g, 8.42 mmol, 1.15 equiv) and the alcohol **S29** (1.98 g, 7.32 mmol, 1.00 equiv) in THF (50 mL) at 0 °C. The resulting mixture was allowed to warm to rt over 3 h and stirring was continued for 14 h. The resulting pale-yellow solution was concentrated under reduced pressure and purified by flash column chromatography (hexanes:EtOAc 4:1) to afford the desired phthalimide protected amine **S30** (2.28 g, 78%) as a colorless solid. IR (thin film) ν 2985, 2871, 2783, 1774, 1750, 1640, 1387, 1307, 717 cm$^{-1}$; $^1$H NMR (500 MHz, $CDCl_3$): δ 7.94 – 7.80 (m, 2H), 7.78 – 7.67 (m, 3H), 7.50 – 7.30 (m, 6H), 7.05 (d, $J$ = 8.6 Hz, 1H), 5.20 (s, 2H), 3.91 (t, $J$ = 7.5 Hz, 2H), 2.99 (t, $J$ = 7.5 Hz, 2H); $^{13}$C NMR (125 MHz, $CDCl_3$): δ 168.2, 150.8, 140.1, 135.7, 134.5, 134.2, 132.0, 130.9, 128.8, 128.3, 127.1, 126.0, 123.5, 115.6, 71.3, 38.8, 33.4; $R_f$ = 0.18 (hexanes:EtOAc 1:1); ESI-HRMS calcd for $C_{23}H_{19}N_2O_5$ [M+H] 403.1288, found 403.1266 and $C_{22}H_{19}N_2O_5Na$ [M+Na] 425.1113, found 425.1111.

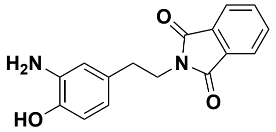

**2-(3-Amino-4-hydroxyphenethyl)isoindoline-1,3-dione (S31)**. A flame dried round-bottomed flask was charged with nitroarene **S30** (100 mg, 0.250 mmol, 1.00 equiv) in a mixture of EtOH-$CH_2Cl_2$ (1:1; 18 mL) at rt. Pd-C (10% on activated charcoal; 15 mg) was added to the clear solution, which was purged with $H_2$ for 15 min with the $H_2$ inlet needle below the solvent surface. The $H_2$ inlet needle was raised above the solvent surface and stirring was continued for 18 h. The resulting black suspension was filtered over a short plug of Celite® ($CH_2Cl_2$ rinse). The filtrate was concentrated under reduced pressure to afford a brown oil that was purified by flash column chromatography (hexanes:EtOAc 1:1) to afford the desired amino alcohol **S31** (70 mg, quant.) as a yellow solid. IR (thin film) ν 3373, 2941, 2824, 1410, 1022, 1005, 822, 760, 617 cm$^{-1}$; $^1$H NMR (500 MHz, DMSO-$d_6$): δ 8.77 (br s, OH), 7.96 – 7.72 (m, 4H), 6.50 (d, $J$ = 7.9 Hz, 1H), 6.46 (d, $J$ = 2.0 Hz, 1H), 6.20 (dd, $J$ = 7.9, 2.1 Hz, 1H), 4.45 (br s, $NH_2$), 3.69 (t, $J$ = 7.6, 2H), 2.67 (t, $J$ = 7.6, 2H); $^{13}$C NMR (125 MHz, DMSO-$d_6$): δ 167.7, 142.5, 136.5, 134.4, 131.6, 128.9, 123.0, 116.3, 114.6, 114.3, 39.5, 33.4; $R_f$ = 0.20 (hexanes:EtOAc 1:1); ESI-HRMS calcd for $C_{16}H_{15}N_2O_3$ [M+H] 283.1077 found 283.1066 and $C_{16}H_{14}N_2O_3Na$ [M+Na] 305.0902, found 305.0880.

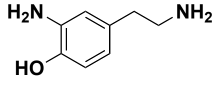

**2-Amino-4-(2-aminoethyl)phenol (S32)**. Hydrazine monohydrate (120 μL, 2.48 mmol, 10.0 equiv) was added to a suspension of the phthalimide protected amine **S31** (70.0 mg, 0.248 mmol,

1.00 equiv) in EtOH (1.5 mL). The resulting solution was heated to reflux for 1.5 h while colorless solids crashed out. The resulting suspension was allowed to cool to rt before $H_2O$ (10 mL) was added in one portion. Stirring was continued to afford a clear solution that was extracted with EtOAc (3 x 5 mL). The combined organic layers were washed with $H_2O$ (3 x 5 mL), brine (3 x 5 mL), dried over anhydrous $Na_2SO_4$, filtered and concentrated under reduced pressure to yield analytically pure amine **S32** (18 mg, 48%). IR (thin film) ν 3384, 2947, 2822, 1580, 1239, 1049, 1021, 837 $cm^{-1}$; $^1H$ NMR (500 MHz, $CD_3OD$): δ 6.66 (d, *J* = 8.0 Hz, 1H), 6.63 (d, *J* = 2.2 Hz, 1H), 6.47 (dd, *J* = 8.0, 2.2 Hz, 1H), 3.05 (t, *J* = 7.5 Hz, 2H), 2.75 (t, *J* = 7.5 Hz, 2H), OH- and NH-protons are not visible; $^{13}C$ NMR (125 MHz, $CD_3OD$): δ 145.2, 136.4, 131.2, 120.2, 117.7, 115.7, 43.6, 37.7; ESI-HRMS calcd for $C_8H_{13}N_2O$ [M+H] 153.1028 found 153.1019.

### PREPARATION OF HYDROXYTYROSOL

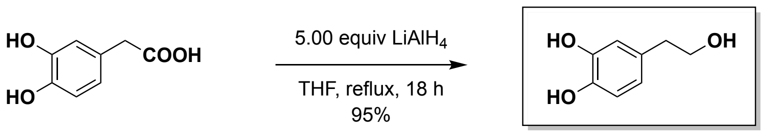

**2-Amino-4-(2-aminoethyl)phenol (S33)**. $LiAlH_4$ (340 mg, 8.92 mmol, 5.00 equiv) was added in small portions to a solution of commercially available 3,4-dihydroxyphenylacetic acid (300 mg, 1.78 mmol, 1.00 equiv) in THF (35 mL) at 0 °C. The suspension was allowed to warm to rt over 30 min before being heated to reflux for 18 h. The resulting mixture was cooled to 0 °C and quenched with the slow addition of aqueous 0.5 M HCl (30 mL). The layers were separated, and the aqueous layer was extracted with EtOAc (3 x 10 mL). The combined organic layers were washed with $H_2O$ (2 x 10 mL), brine (2 x 10 mL), dried over anhydrous $Na_2SO_4$, filtered, and concentrated under reduced pressure to afford an orange oil. Purification by flash column chromatography (hexanes:EtOAc 1:1) afforded triol **S33** (260 mg, 95%) as a pale red oil. $^1H$ NMR (500 MHz, $CD_3OD$): δ 6.74 – 6.60 (m, 2H), 6.58 – 6.48 (m, 1H), 3.67 (t, *J* = 7.2 Hz, 2H), 2.66 (t, *J* = 7.2 Hz, 2H), OH-protons are not visible; $R_f$ = 0.20 (hexanes:EtOAc 1:1). The spectral characteristics were identical to those reported in the current literature.[17]

### ADDITIONAL INFORMATION

- 2-Aminoethylbenzenediol/-triol derivatives (**S16**–**S20**, **S25**) as well as 2-amino-4-(2-aminoethyl)phenol **S32** are sensitive towards oxidation and turn black within hours if stored in the presence of $O_2$. No noticeable change in their composition is observed, according to $^1H$ NMR, when stored in the absence of $O_2$ at 3–4 °C.
- Salt formation of the oxygen sensitive alkylamines (**S16**–**S20**, **S25**, **S32**) leads to more stable compounds as no decomposition was observed when stored in the presence of $O_2$ for several days.
- Upon quenching of the phenol ether cleavage reaction with MeOH, a reaction that contains $BBr_3$, volatile $B(OMe)_3$ is formed as sole side product. This can be removed under reduced pressure to afford pure products. It is important to stir the reaction mixture for approximately 1 h upon the addition of MeOH to allow for the full conversion of $BBr_3$ to $B(OMe)_3$. The obtained residue can be dissolved in additional MeOH and again concentrated under reduced pressure to ensure the complete removal of $B(OMe)_3$.

**[1]H / [13]C spectra**

### [1]H / [13]C SPECTRA

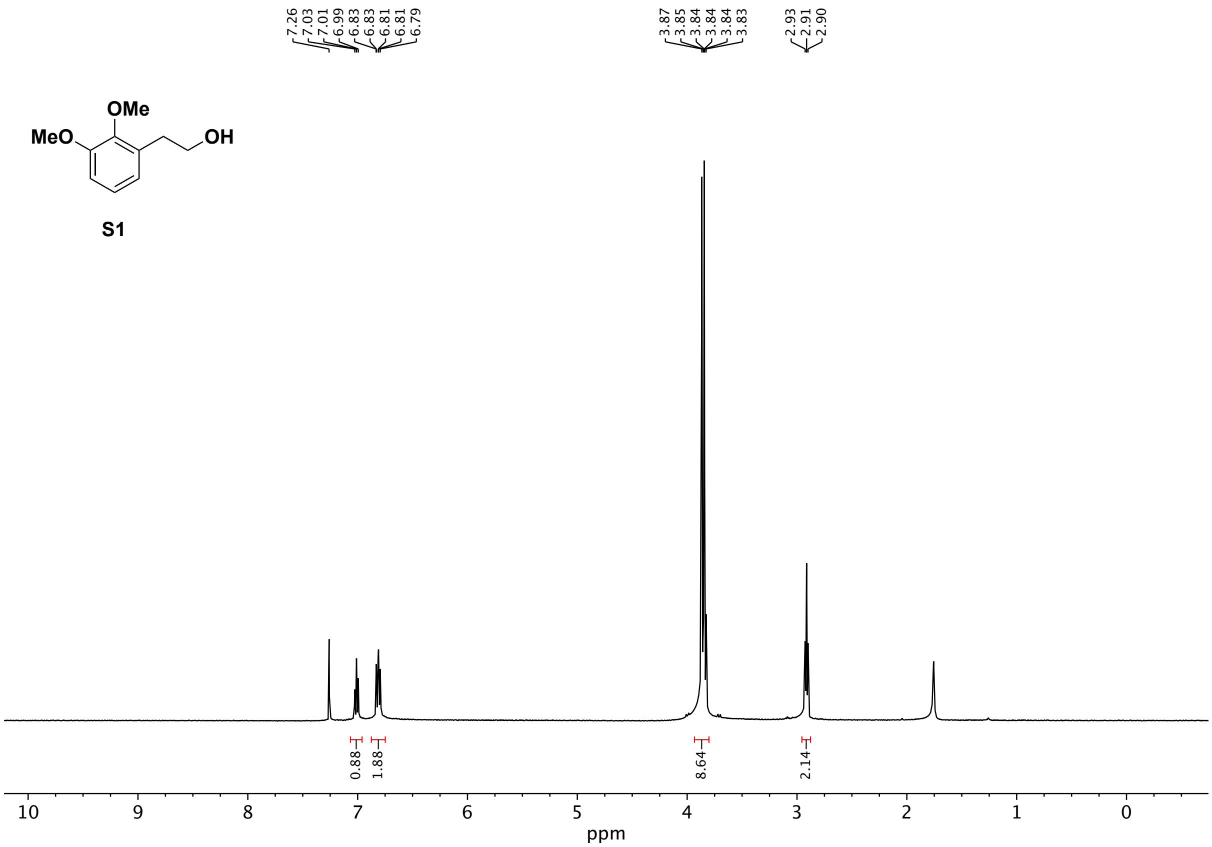

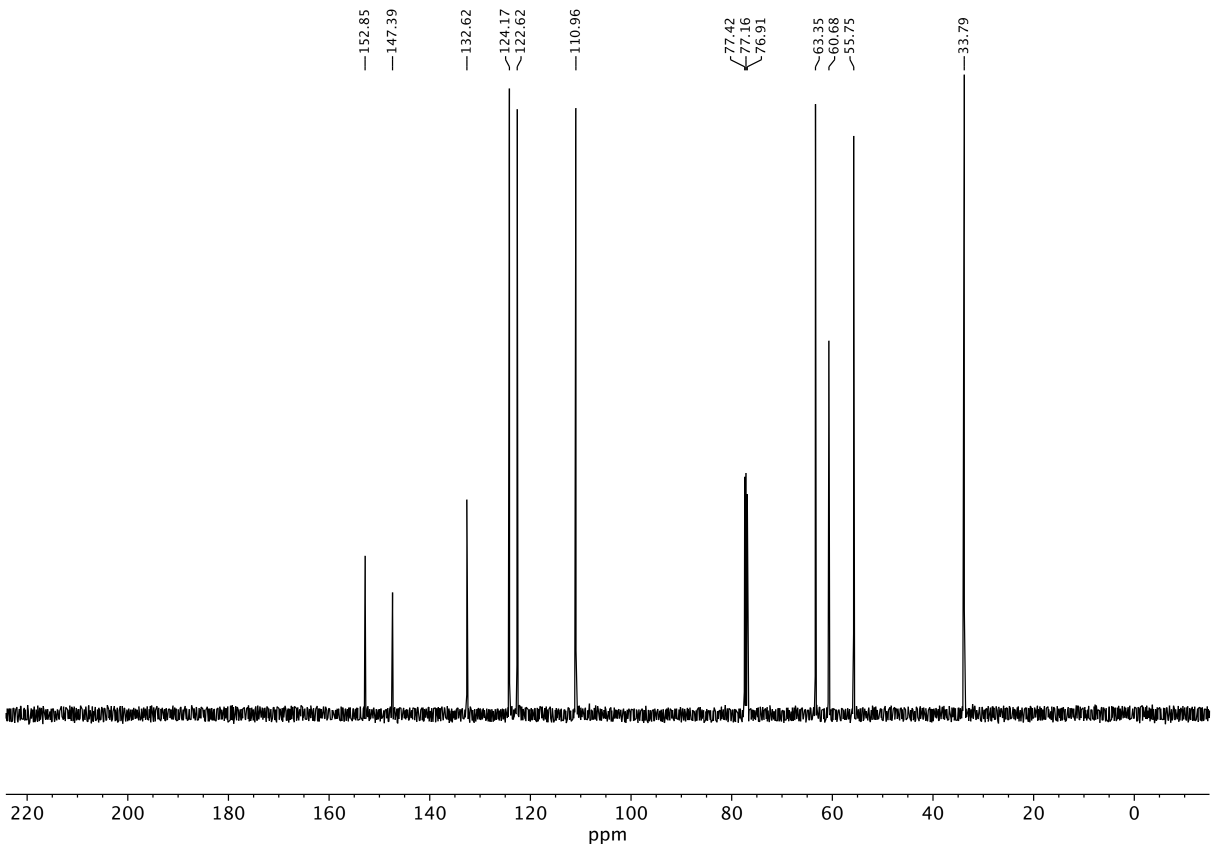
152.85
147.39
132.62
124.17
122.62
110.96
77.42
77.16
76.91
63.35
60.68
55.75
33.79
220
200
180
160
140
120
100
80
60
40
20
0
ppm

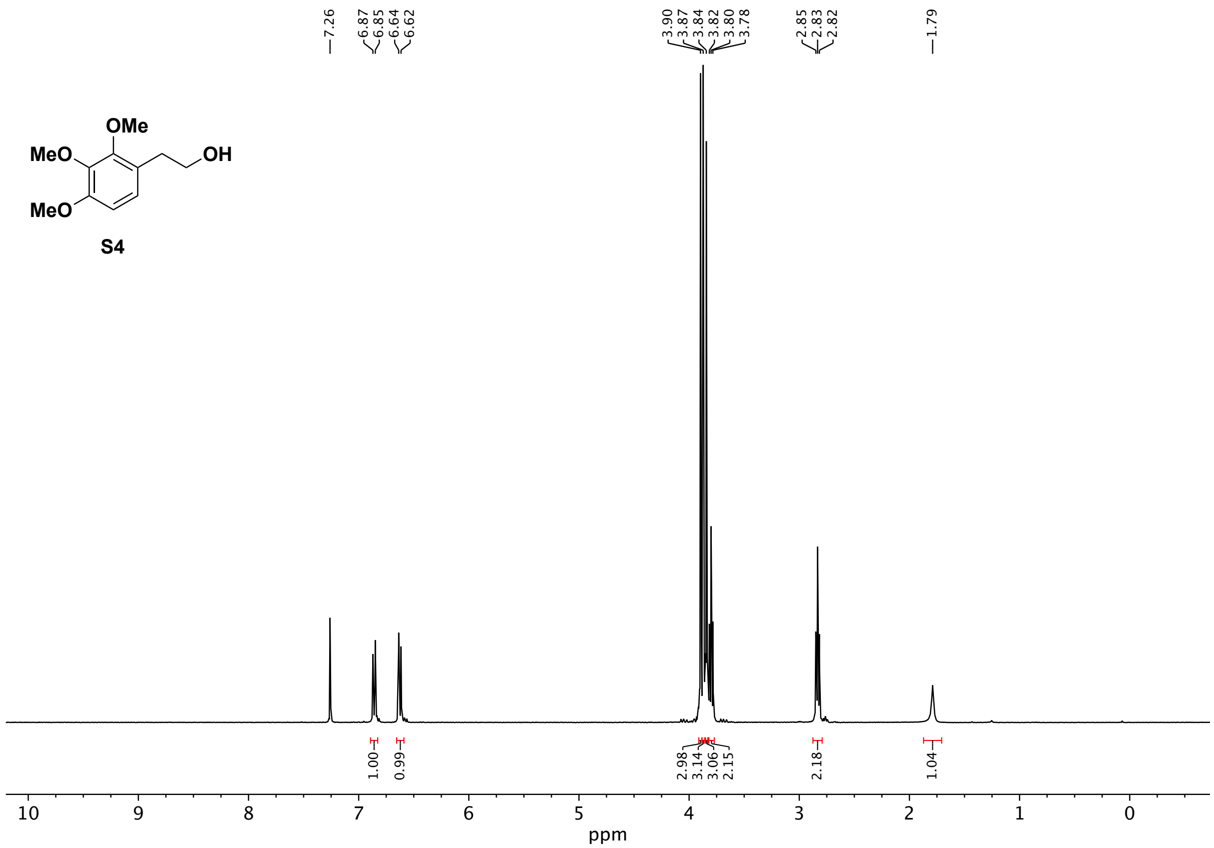

OMe
MeO
OH
MeO
S4
7.26
6.87
6.85
6.64
6.62
3.90
3.87
3.84
3.82
3.80
3.78
2.85
2.83
2.82
1.79
1.00
0.99
2.98
3.14
3.06
2.15
2.18
1.04
10
9
8
7
6
5
4
3
2
1
0
ppm

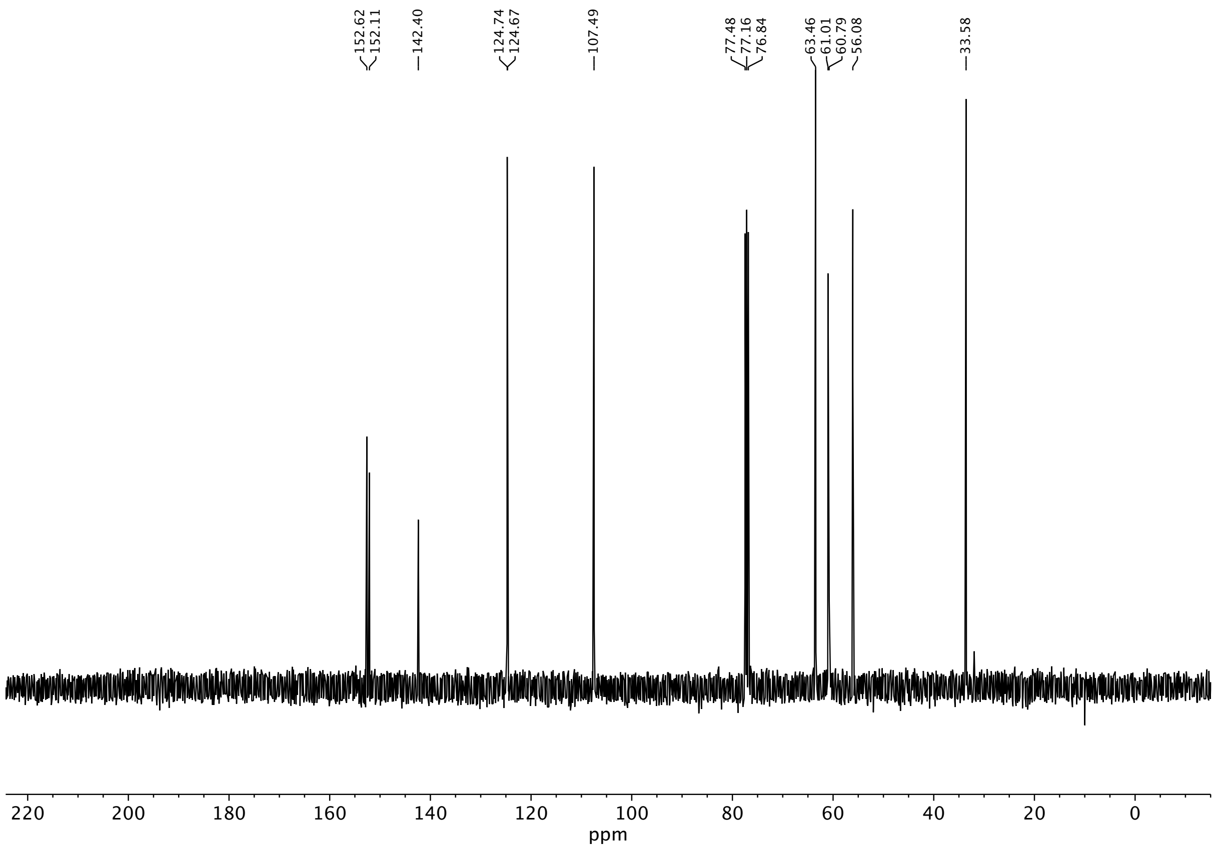
152.62
152.11
142.40
124.74
124.67
107.49
77.48
77.16
76.84
63.46
61.01
60.79
56.08
33.58
220 200 180 160 140 120 100 80 60 40 20 0
ppm

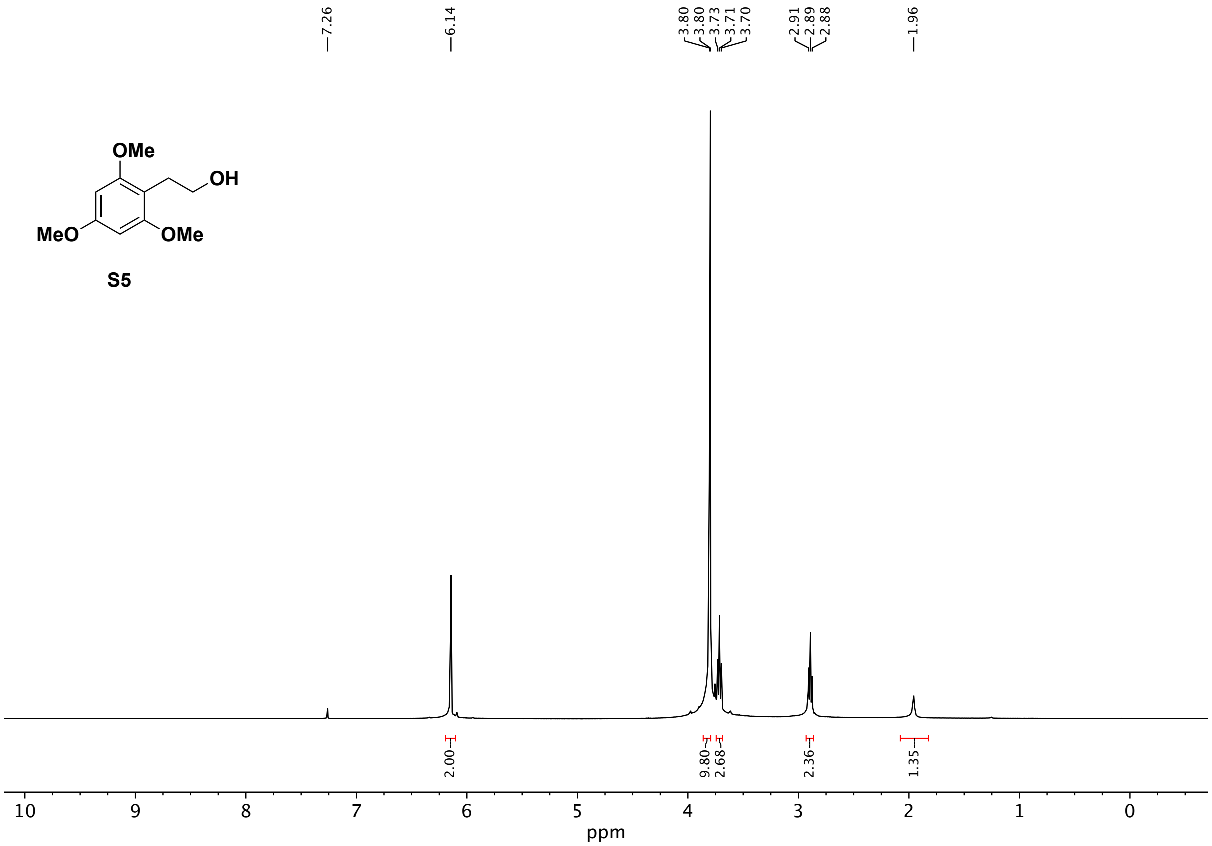
7.26
6.14
3.80
3.80
3.73
3.71
3.70
2.91
2.89
2.88
1.96
OMe
OH
MeO
OMe
S5
2.00
9.80
2.68
2.36
1.35
10
9
8
7
6
5
4
3
2
1
0
ppm

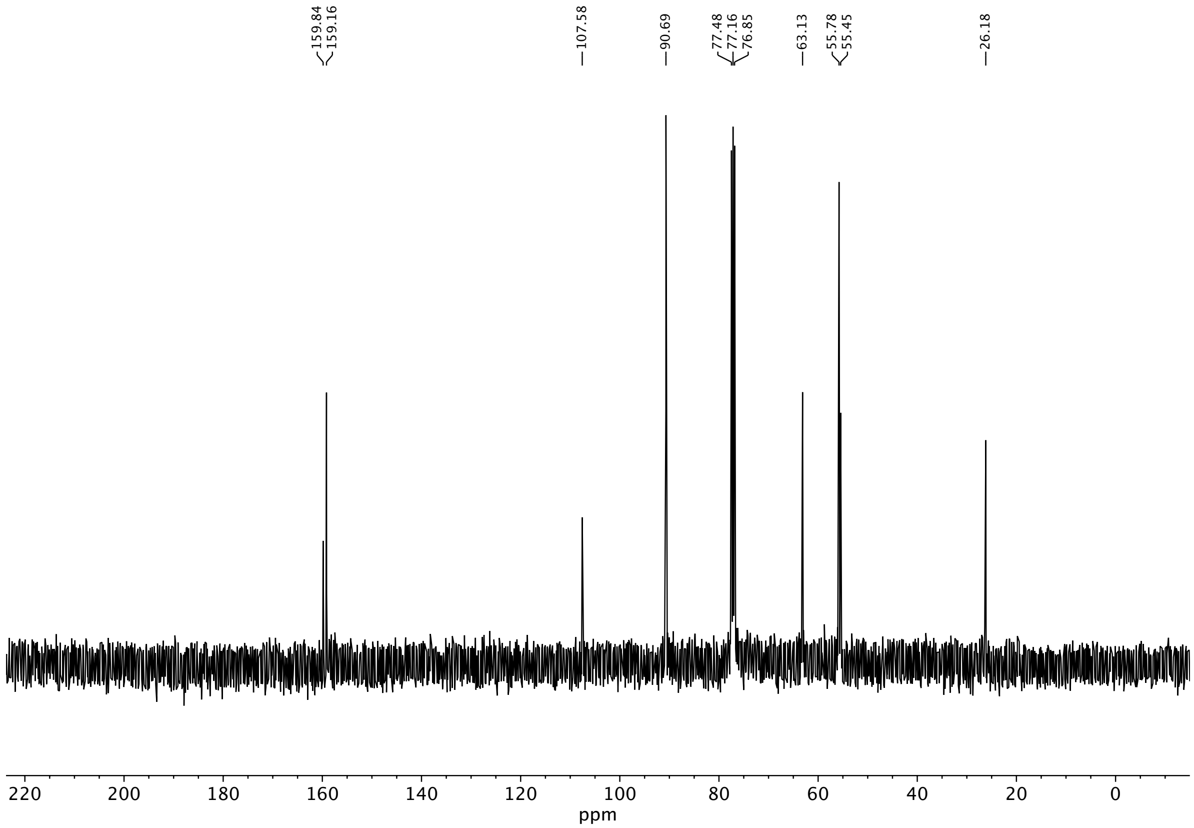
159.84
159.16
107.58
90.69
77.48
77.16
76.85
63.13
55.78
55.45
26.18
220 200 180 160 140 120 100 80 60 40 20 0
ppm

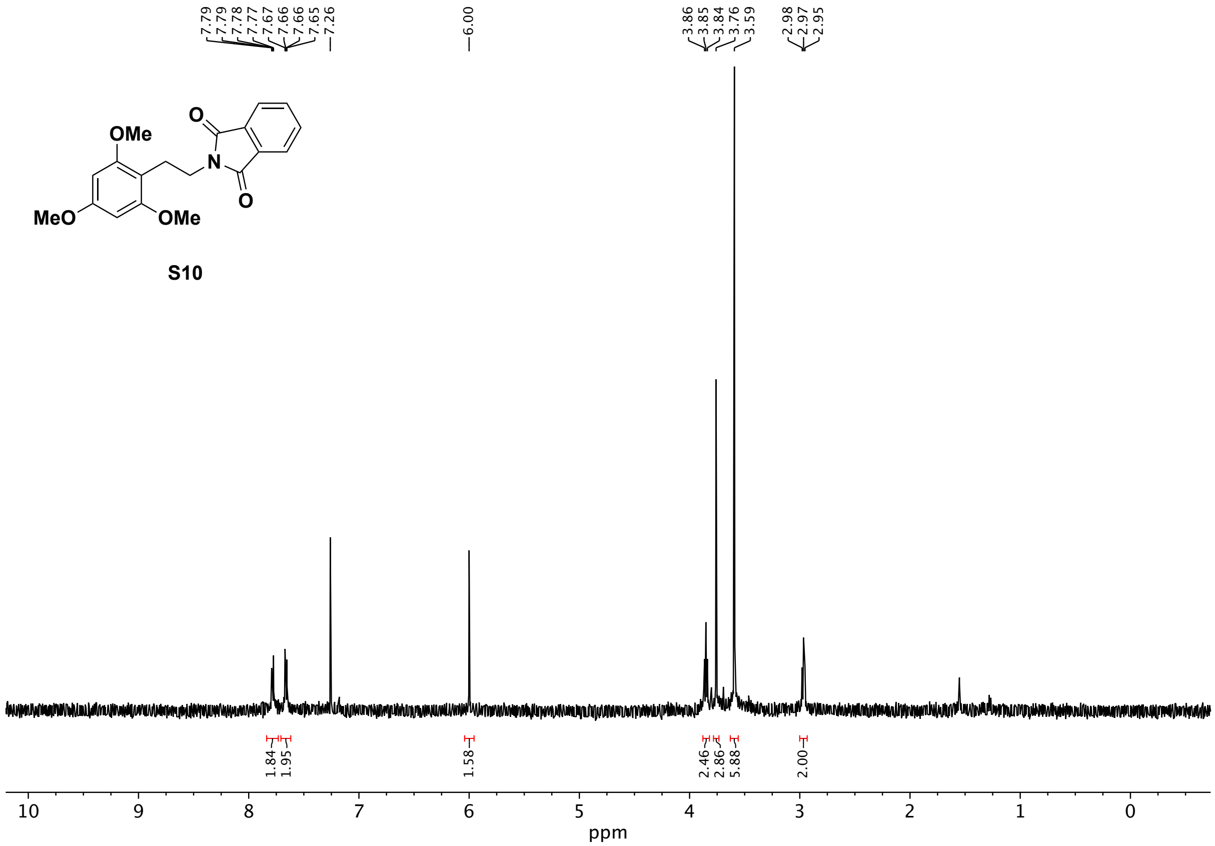
7.79
7.79
7.78
7.77
7.67
7.66
7.66
7.65
7.26
6.00
3.86
3.85
3.84
3.76
3.59
2.98
2.97
2.95
OMe
O
N
O
MeO
OMe
1.84
1.95
1.58
2.46
2.86
5.88
2.00
10
9
8
7
6
5
4
3
2
1
0
ppm

**S10**

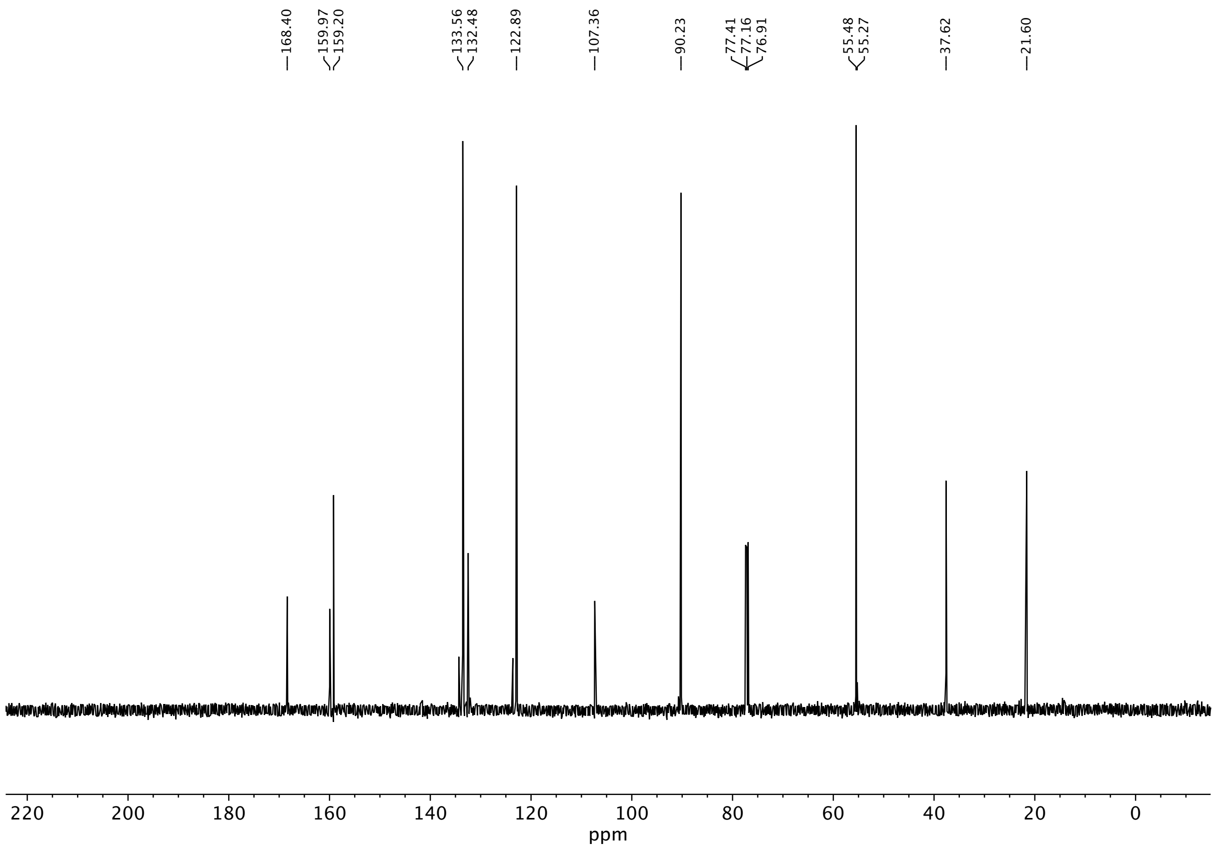
168.40
159.97
159.20
133.56
132.48
122.89
107.36
90.23
77.41
77.16
76.91
55.48
55.27
37.62
21.60
220
200
180
160
140
120
100
80
60
40
20
0
ppm

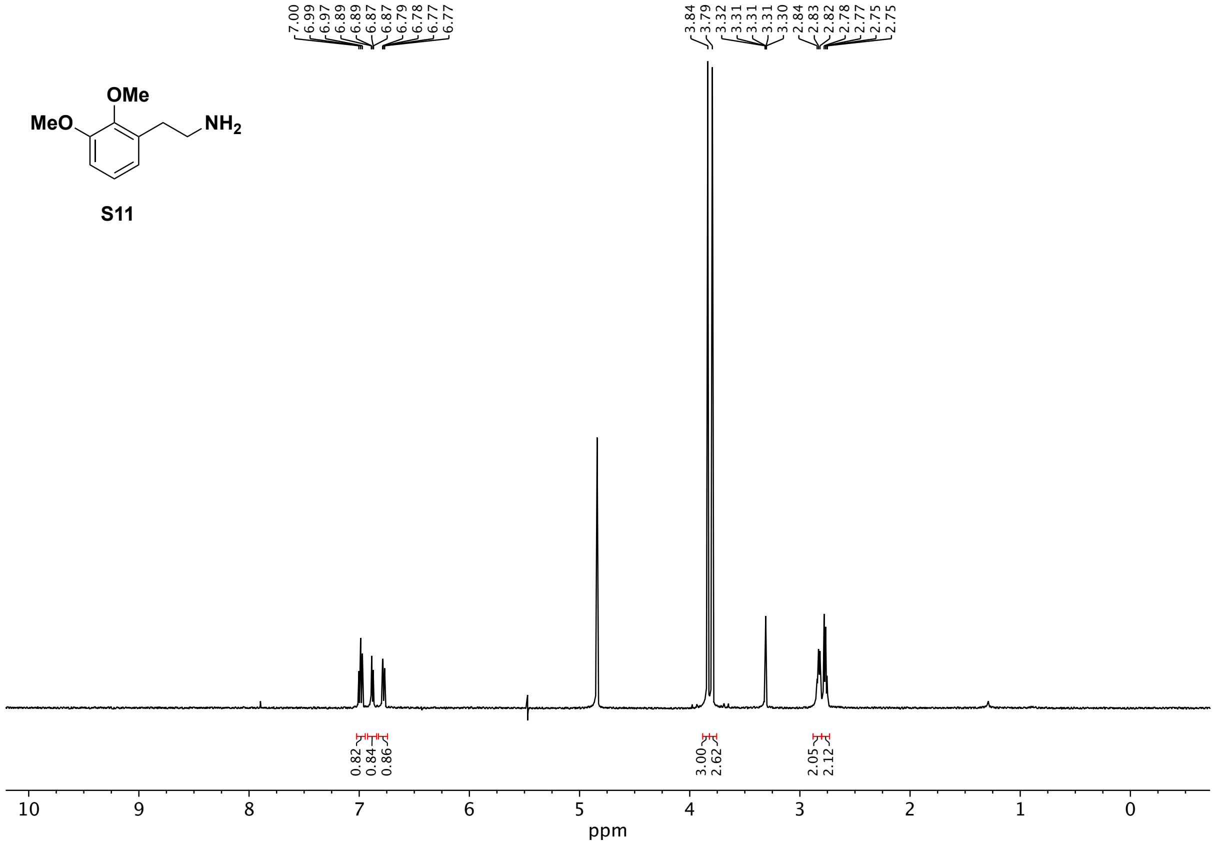
OMe
MeO
NH2
7.00
6.99
6.97
6.89
6.87
6.87
6.79
6.78
6.77
6.77
3.84
3.79
3.32
3.31
3.31
3.31
3.30
2.84
2.83
2.82
2.78
2.77
2.75
2.75
0.82
0.84
0.86
3.00
2.62
2.05
2.12
10
9
8
7
6
5
4
3
2
1
0
ppm

S11

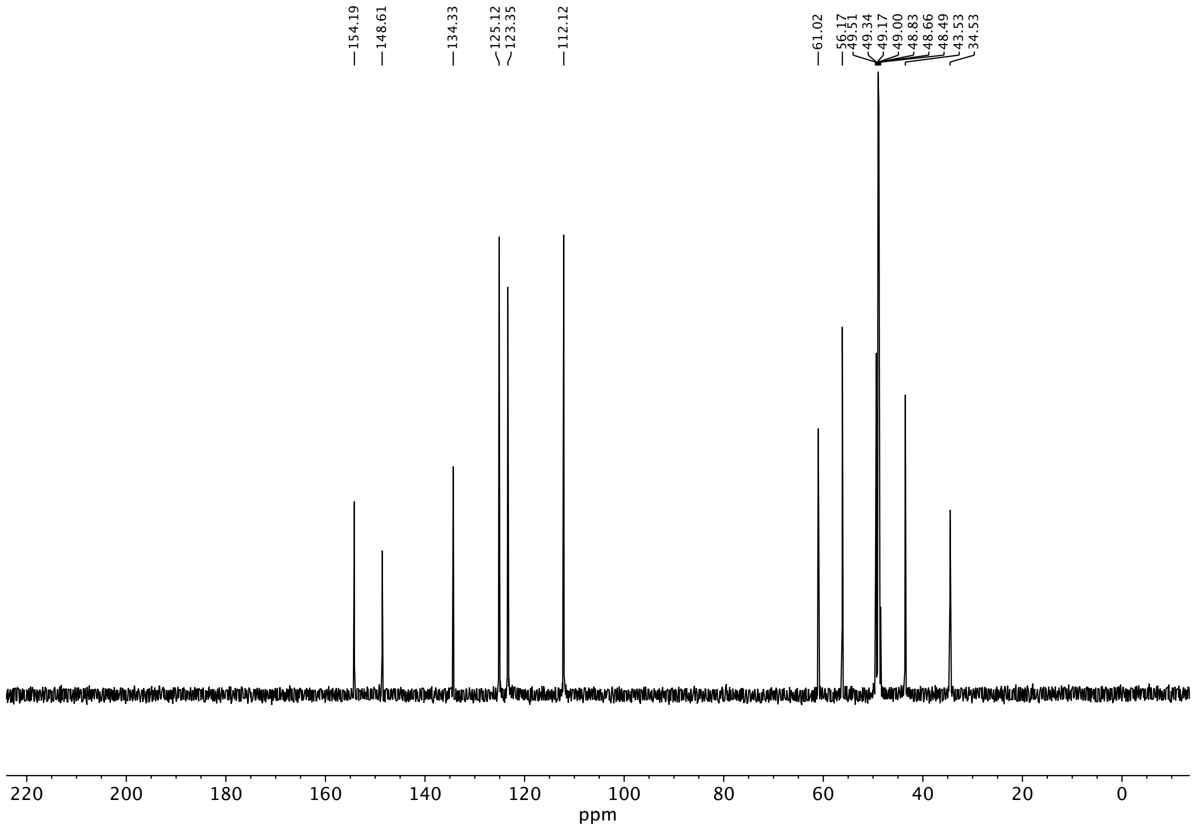
154.19
148.61
134.33
125.12
123.35
112.12
61.02
56.17
49.51
49.34
49.17
49.00
48.83
48.66
48.49
43.53
34.53
220 200 180 160 140 120 100 80 60 40 20 0
ppm

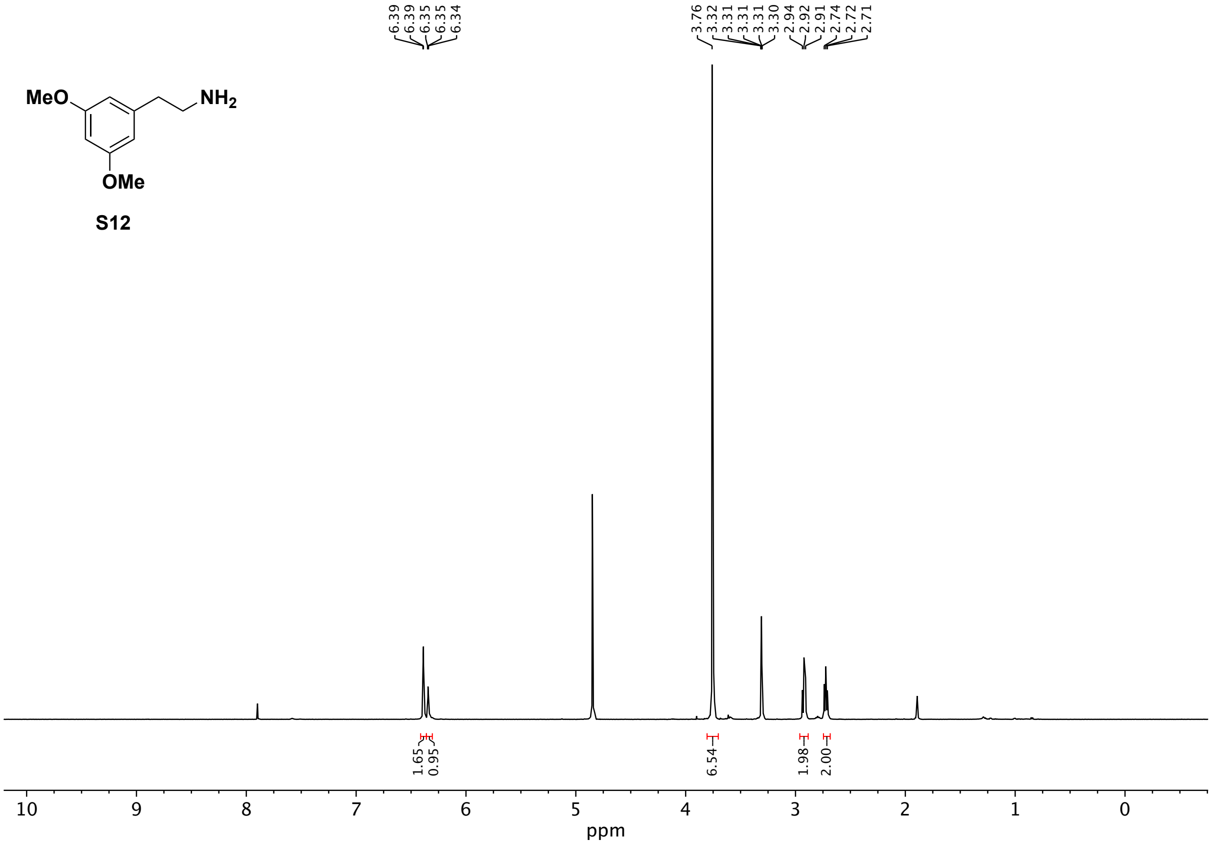
MeO
NH2
OMe
6.39
6.39
6.35
6.35
6.34
3.76
3.32
3.31
3.31
3.31
3.30
2.94
2.92
2.91
2.74
2.72
2.71
1.65
0.95
6.54
1.98
2.00
10
9
8
7
6
5
4
3
2
1
0
ppm

**S12**

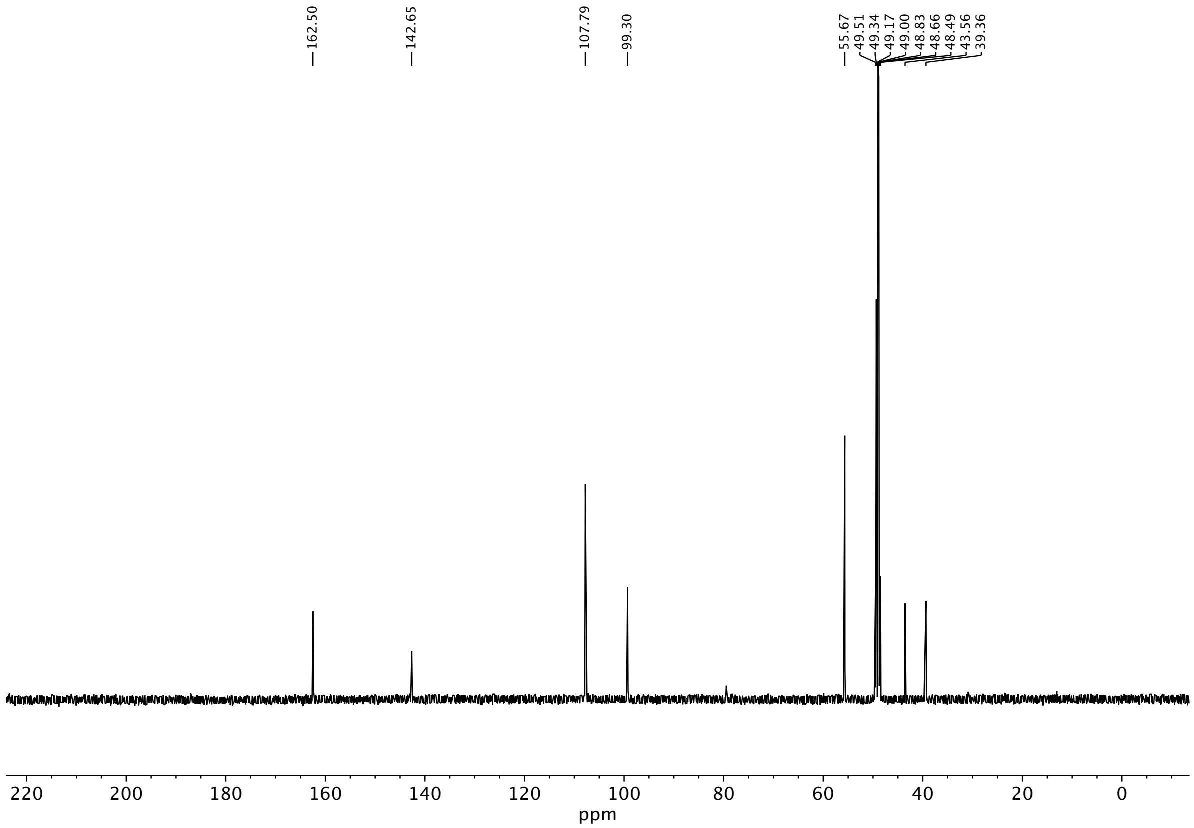
162.50
142.65
107.79
99.30
55.67
49.51
49.34
49.17
49.00
48.83
48.66
48.49
43.56
39.36
220 200 180 160 140 120 100 80 60 40 20 0
ppm

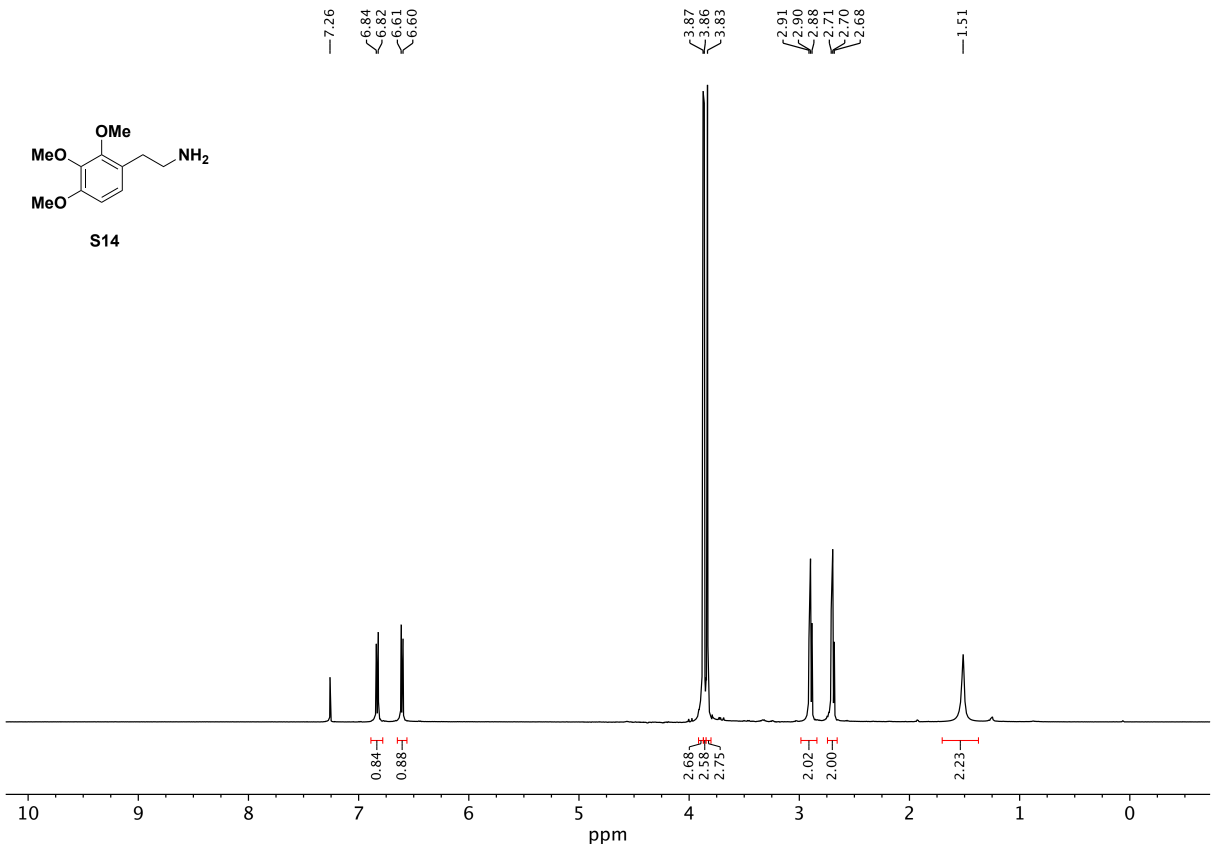
MeO
OMe
NH2
MeO
S14
7.26
6.84
6.82
6.61
6.60
3.87
3.86
3.83
2.91
2.90
2.88
2.71
2.70
2.68
1.51
0.84
0.88
2.68
2.58
2.75
2.02
2.00
2.23
10
9
8
7
6
5
4
3
2
1
0
ppm

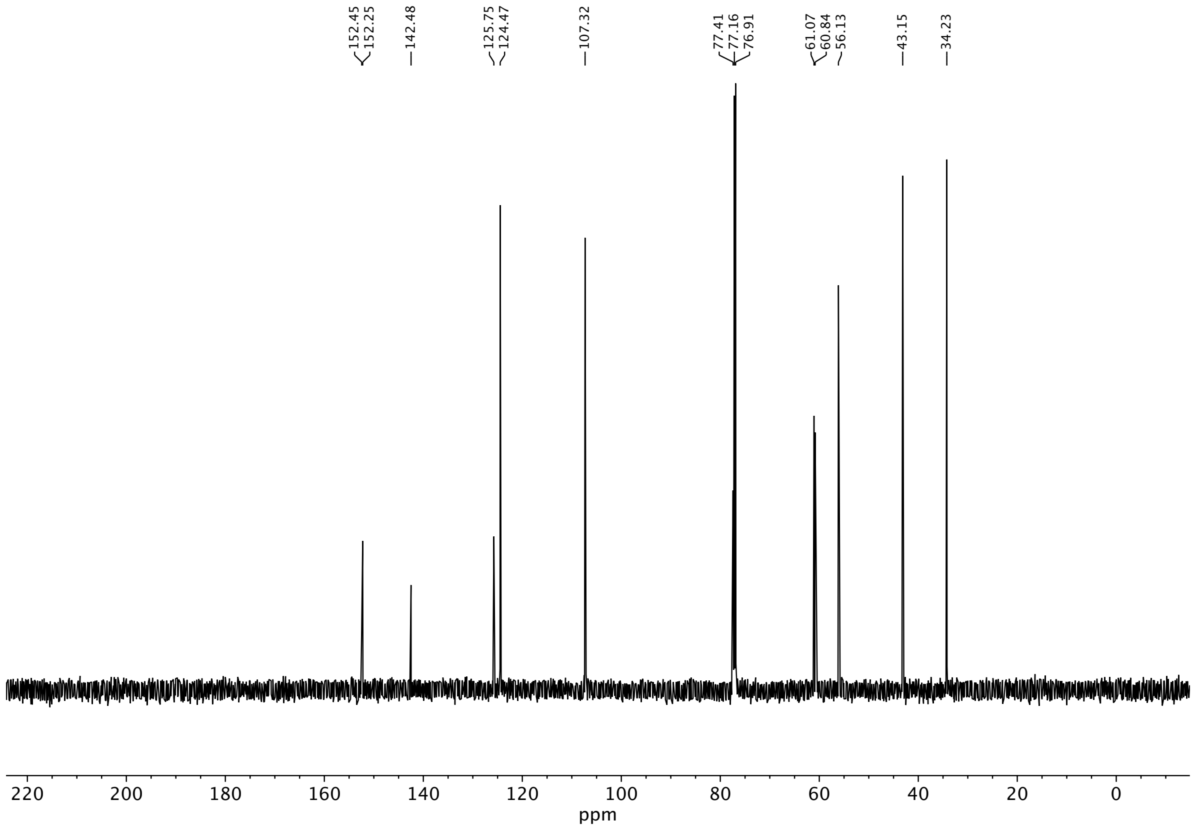

152.45
152.25
142.48
125.75
124.47
107.32
77.41
77.16
76.91
61.07
60.84
56.13
43.15
34.23
220
200
180
160
140
120
100
80
60
40
20
0
ppm

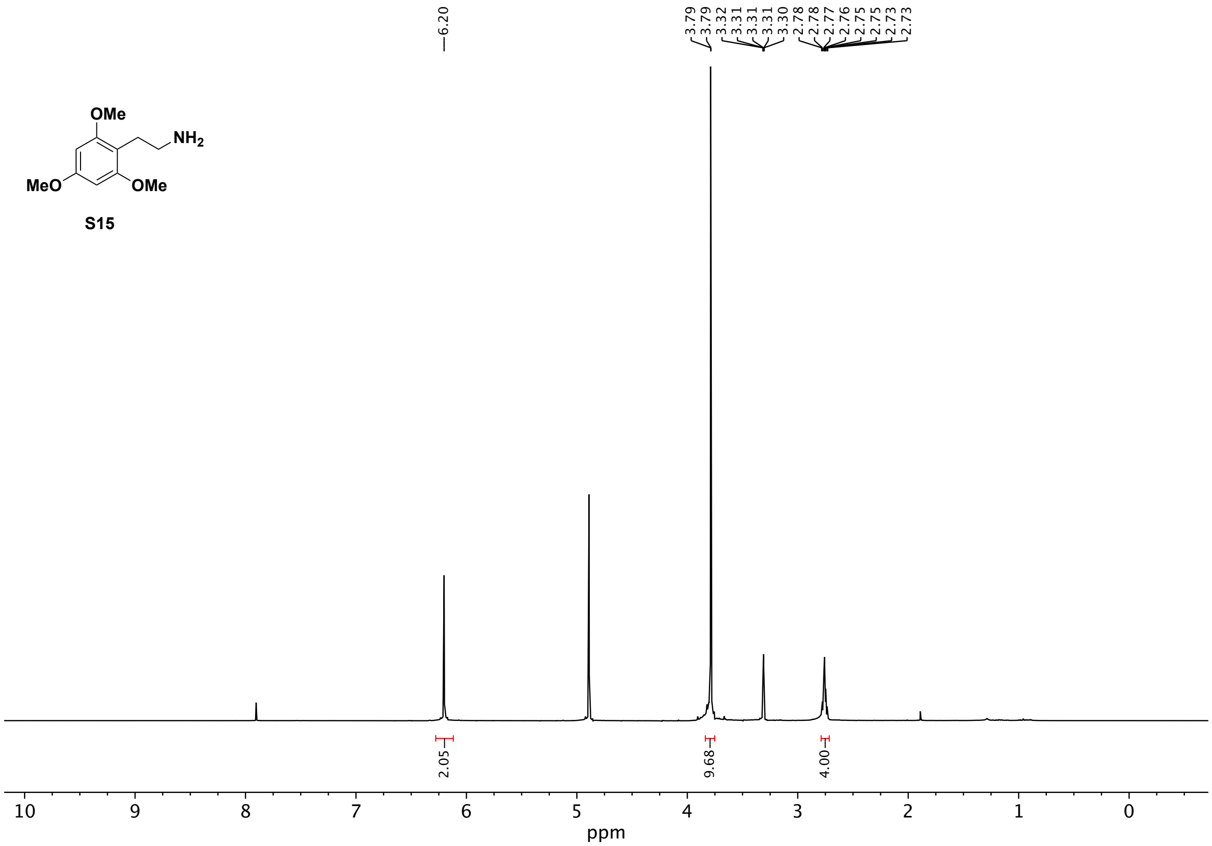
OMe
NH2
MeO
OMe
S15
6.20
3.79
3.79
3.32
3.31
3.31
3.31
3.30
2.78
2.78
2.77
2.76
2.75
2.75
2.73
2.73
2.05
9.68
4.00
10 9 8 7 6 5 4 3 2 1 0
ppm

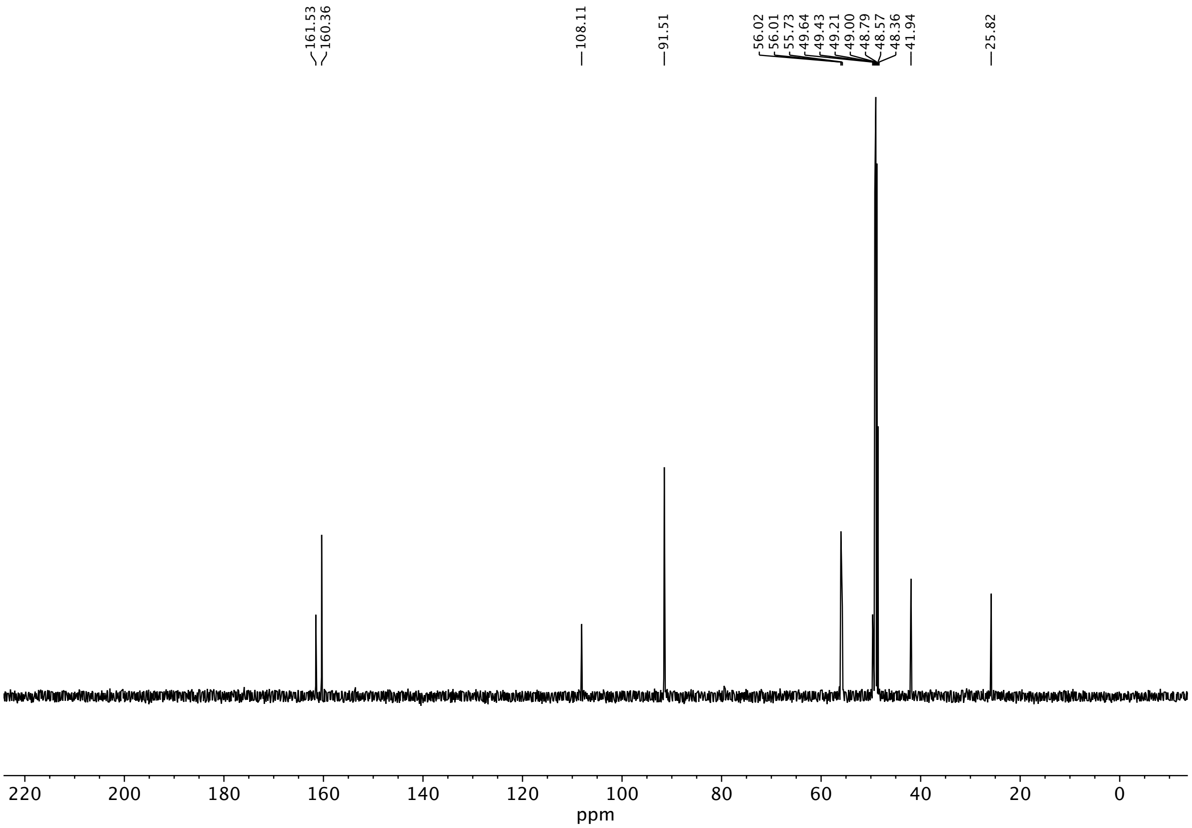
161.53
160.36
108.11
91.51
56.02
56.01
55.73
49.64
49.43
49.21
49.00
48.79
48.57
48.36
41.94
25.82
220
200
180
160
140
120
100
80
60
40
20
0
ppm

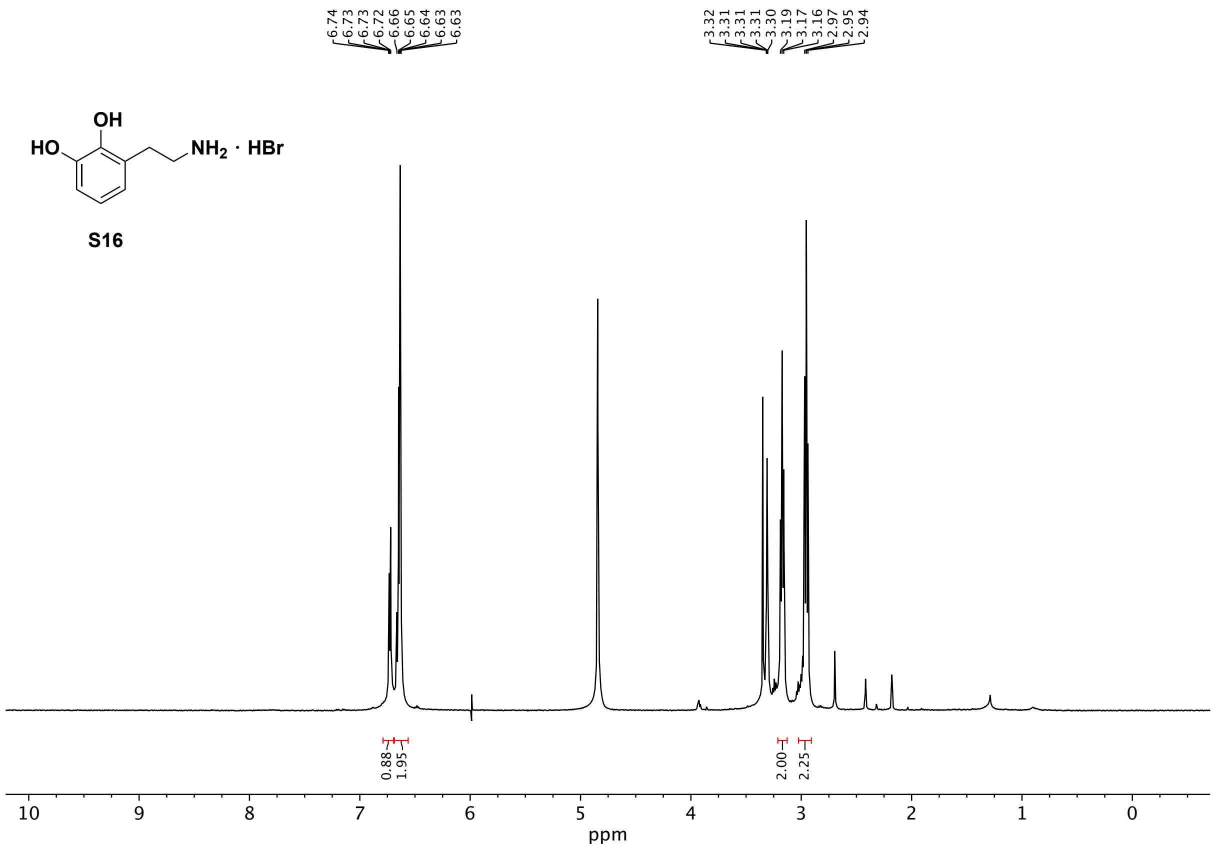
6.74
6.73
6.73
6.72
6.66
6.65
6.64
6.63
6.63
3.32
3.31
3.31
3.31
3.30
3.19
3.17
3.16
2.97
2.95
2.94
OH
HO
NH2 · HBr
S16
0.88
1.95
2.00
2.25
10
9
8
7
6
5
4
3
2
1
0
ppm

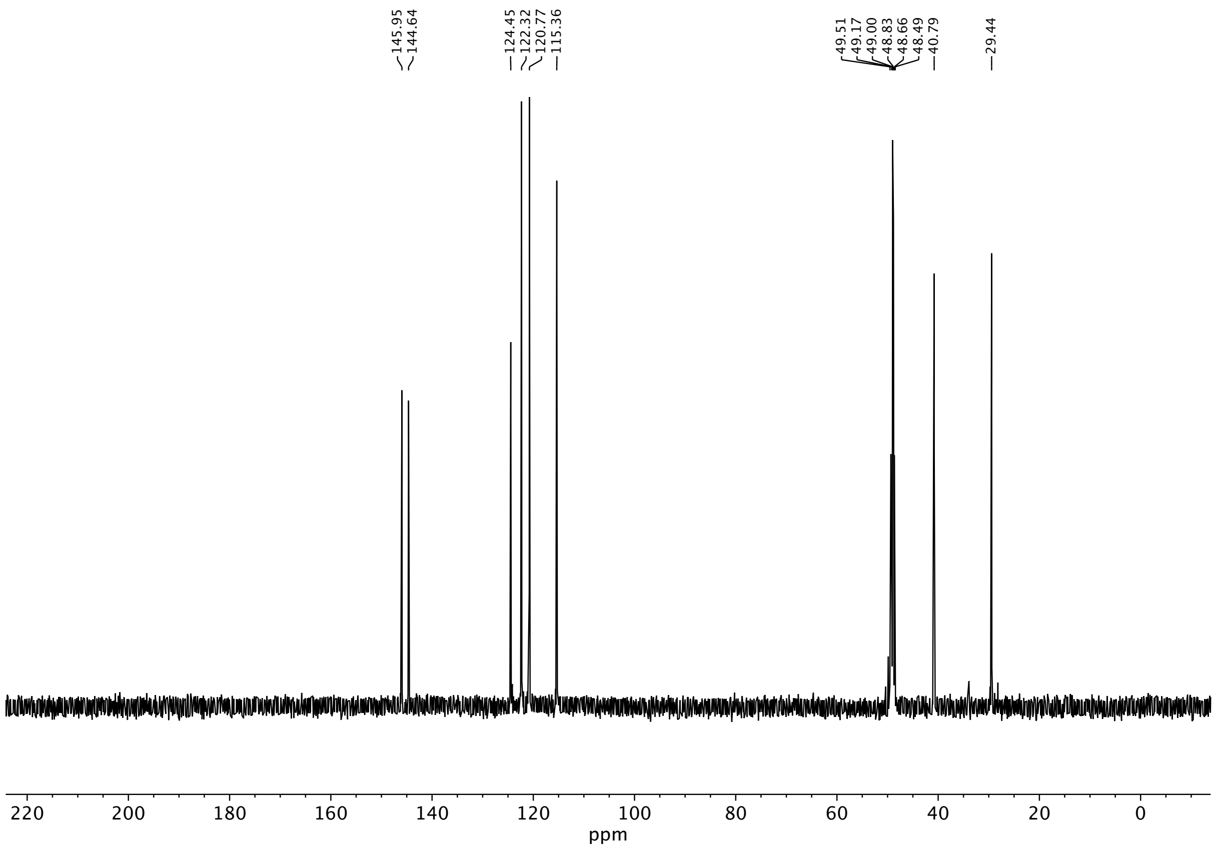
145.95
144.64
124.45
122.32
120.77
115.36
49.51
49.17
49.00
48.83
48.66
48.49
40.79
29.44
220
200
180
160
140
120
100
80
60
40
20
0
ppm

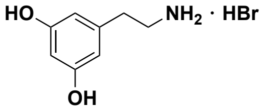
HO
NH2 · HBr
OH

**S17**

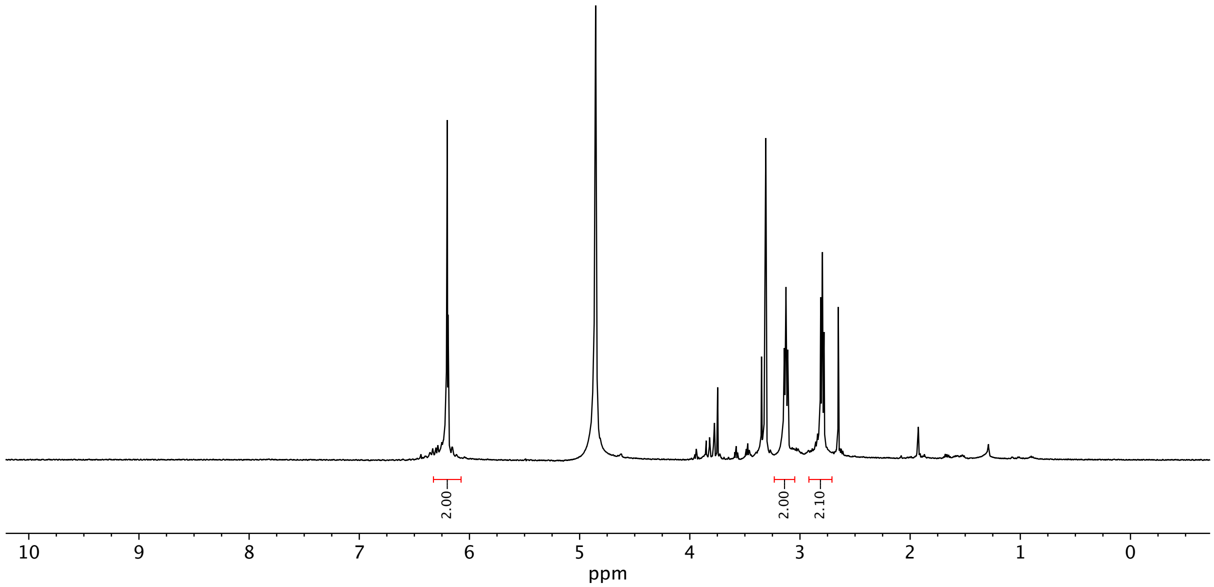
6.21
6.20
6.19
6.19
3.32
3.31
3.31
3.31
3.30
3.14
3.13
3.11
2.81
2.80
2.78
2.00
2.00
2.10
10
9
8
7
6
5
4
3
2
1
0
ppm

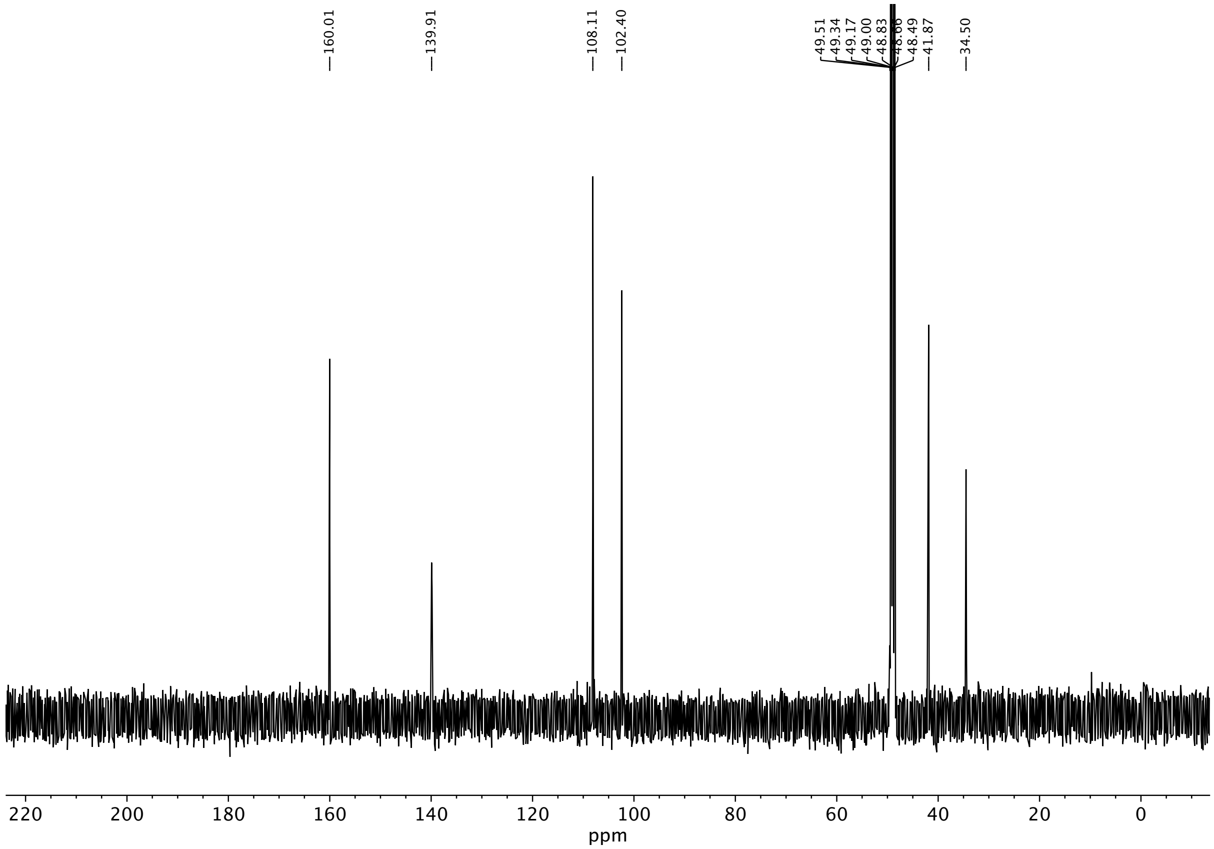
160.01
139.91
108.11
102.40
49.51
49.34
49.17
49.00
48.83
48.66
48.49
41.87
34.50
220 200 180 160 140 120 100 80 60 40 20 0
ppm

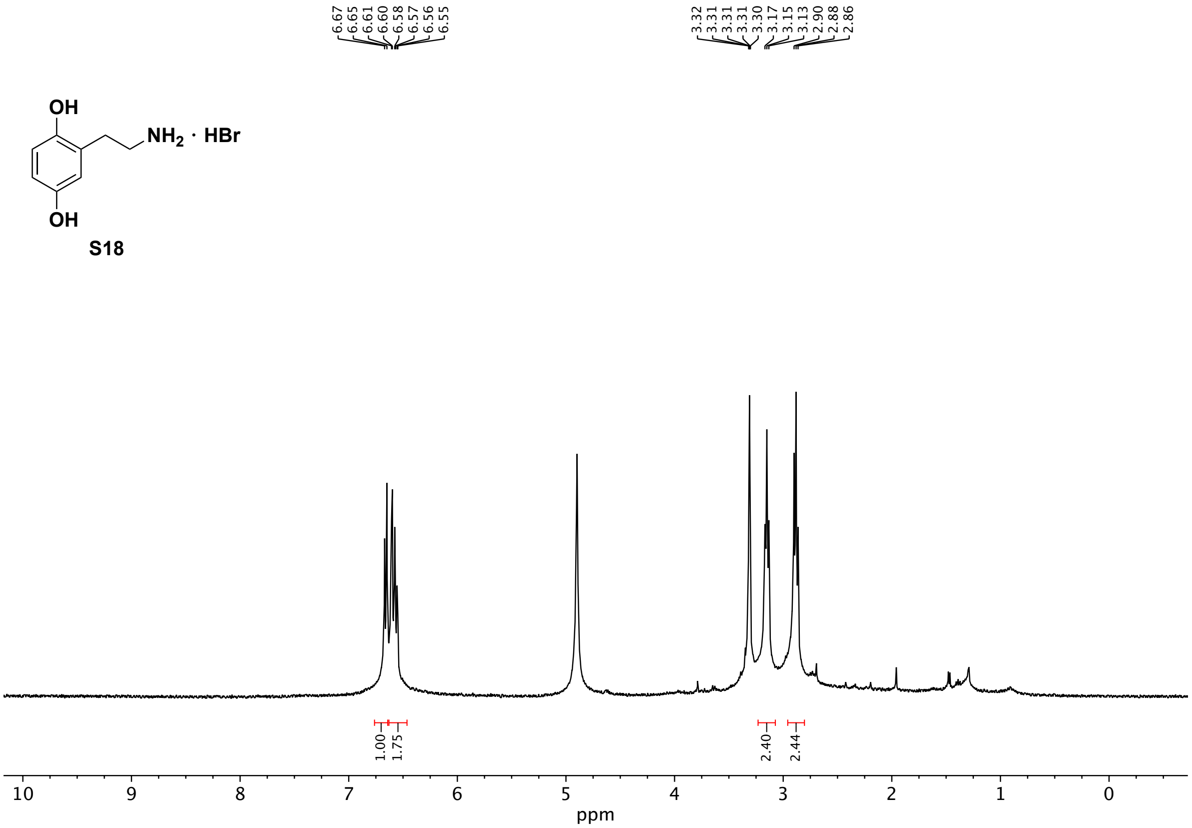

6.67
6.65
6.61
6.60
6.58
6.57
6.56
6.55
3.32
3.31
3.31
3.31
3.30
3.17
3.15
3.13
2.90
2.88
2.86
OH
NH2 · HBr
OH
S18
1.00
1.75
2.40
2.44
10
9
8
7
6
5
4
3
2
1
0
ppm

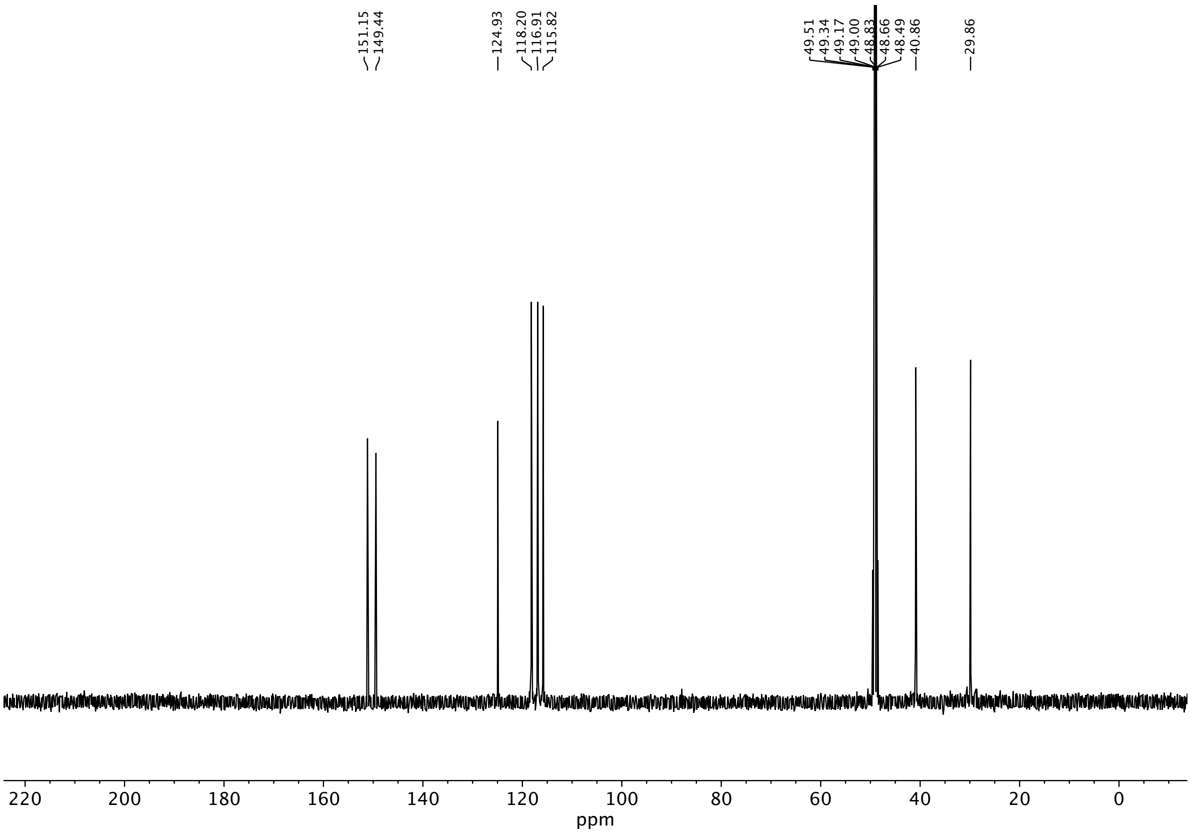

151.15
149.44
124.93
118.20
116.91
115.82
49.51
49.34
49.17
49.00
48.83
48.66
48.49
40.86
29.86
220 200 180 160 140 120 100 80 60 40 20 0
ppm

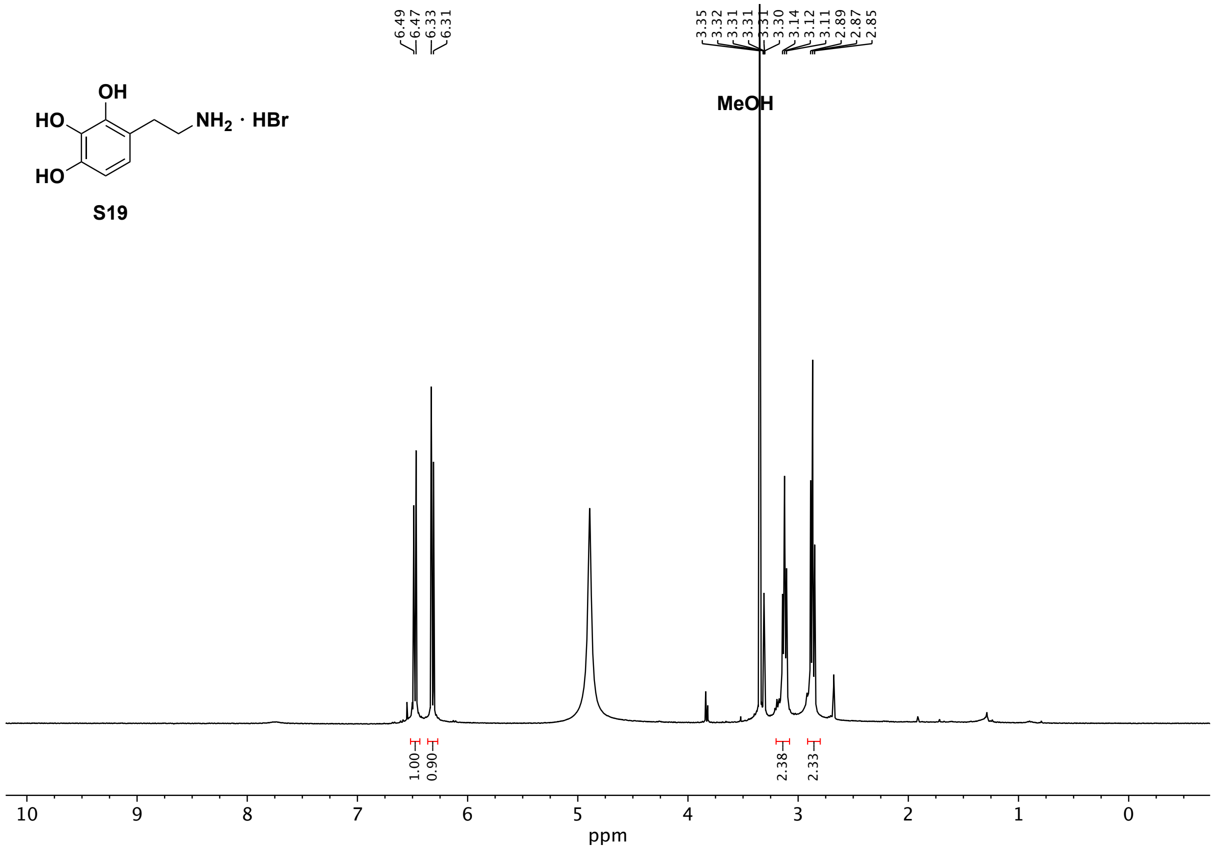

OH
HO
NH2 · HBr
HO
S19
6.49
6.47
6.33
6.31
3.35
3.32
3.31
3.31
3.31
3.30
3.14
3.12
3.11
2.89
2.87
2.85
MeOH
1.00
0.90
2.38
2.33
10
9
8
7
6
5
4
3
2
1
0
ppm

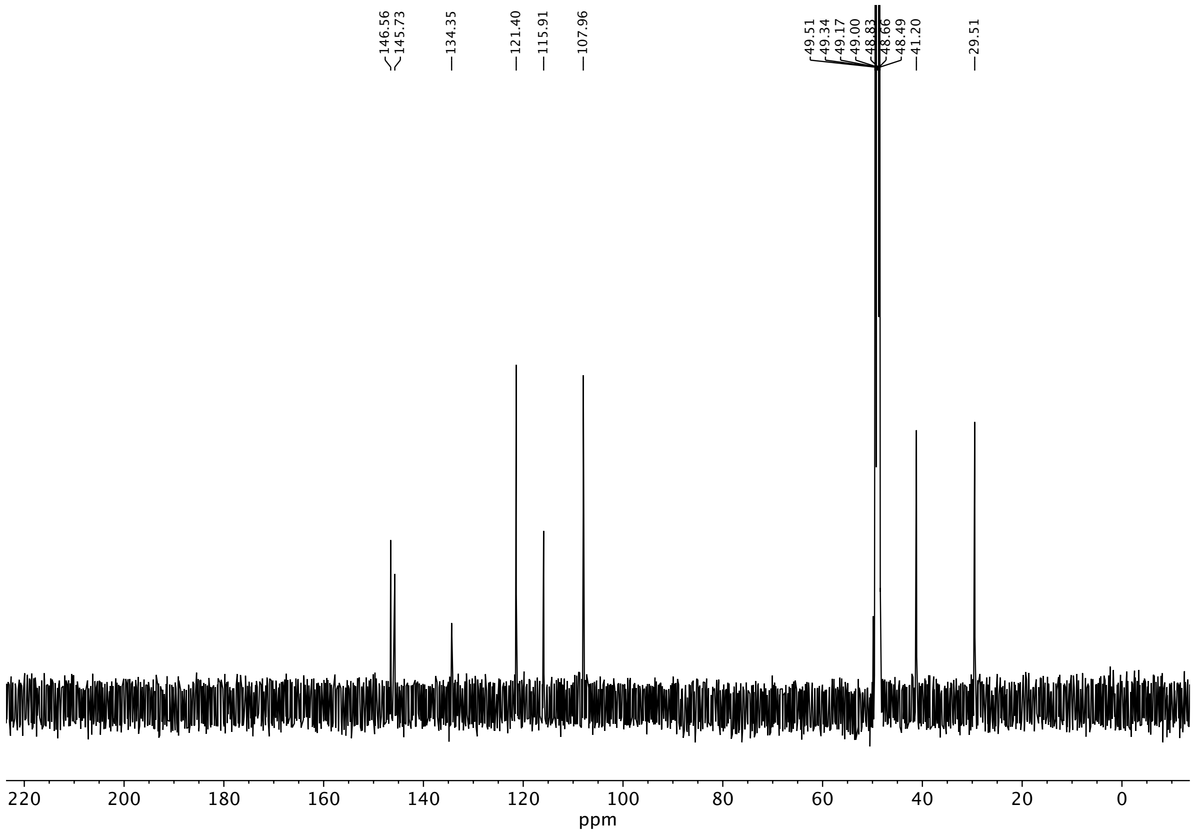
146.56
145.73
134.35
121.40
115.91
107.96
49.51
49.34
49.17
49.00
48.83
48.66
48.49
41.20
29.51
220
200
180
160
140
120
100
80
60
40
20
0
ppm

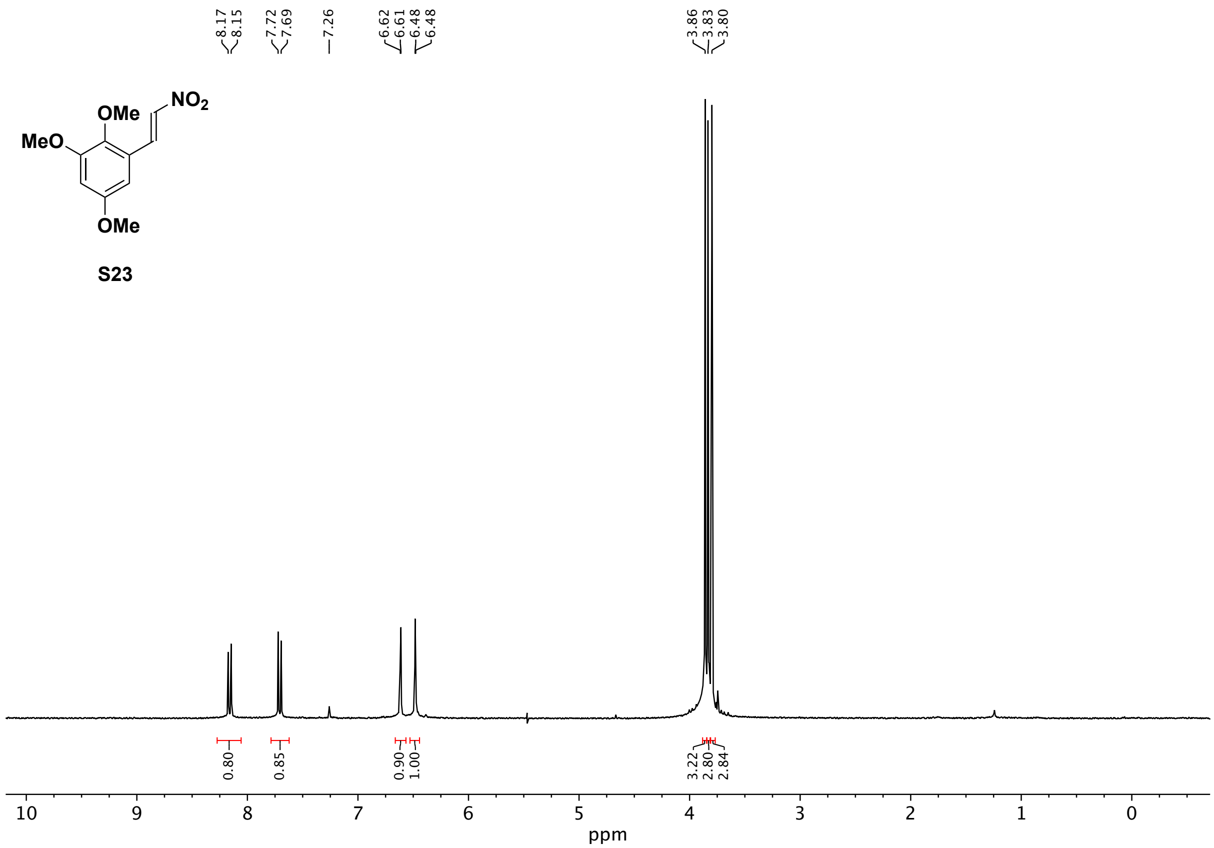
8.17
8.15
7.72
7.69
7.26
6.62
6.61
6.48
6.48
3.86
3.83
3.80
NO2
OMe
MeO
OMe
0.80
0.85
0.90
1.00
3.22
2.80
2.84
10
9
8
7
6
5
4
3
2
1
0
ppm

S23

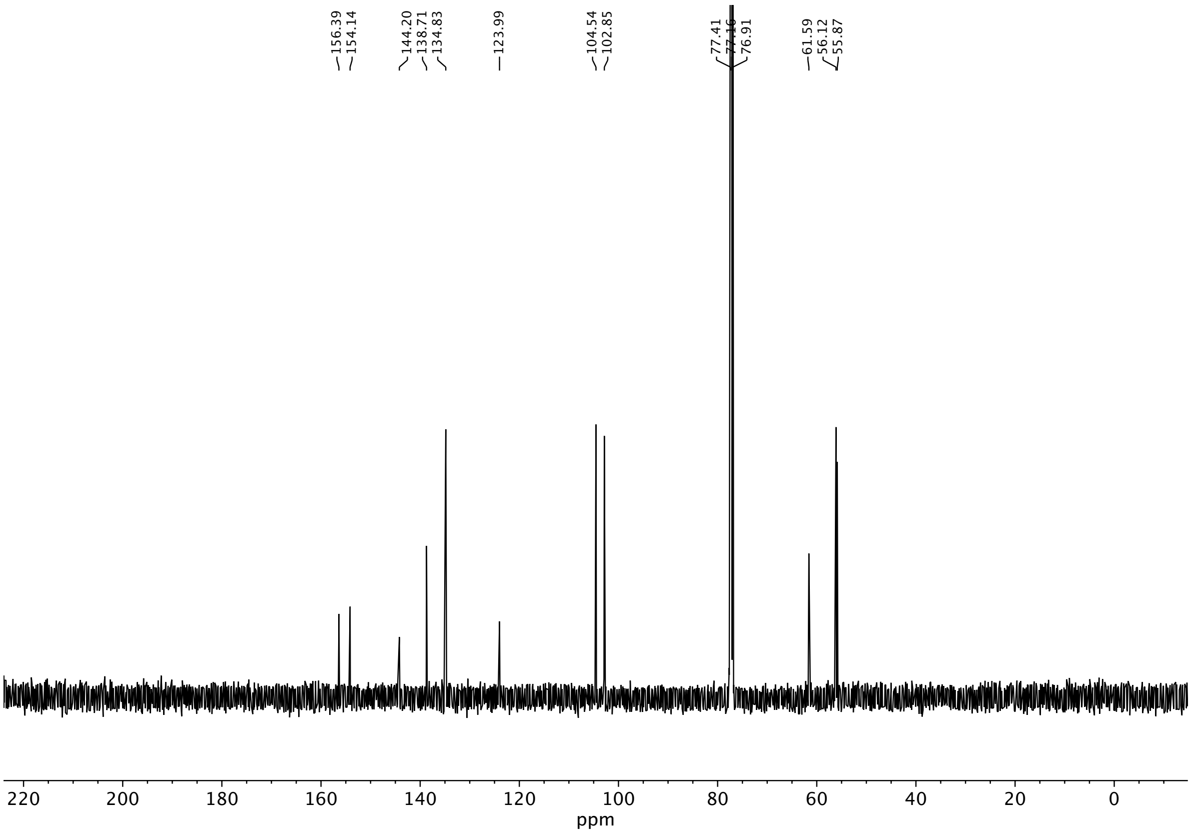
156.39
154.14
144.20
138.71
134.83
123.99
104.54
102.85
77.41
77.16
76.91
61.59
56.12
55.87
220
200
180
160
140
120
100
80
60
40
20
0
ppm

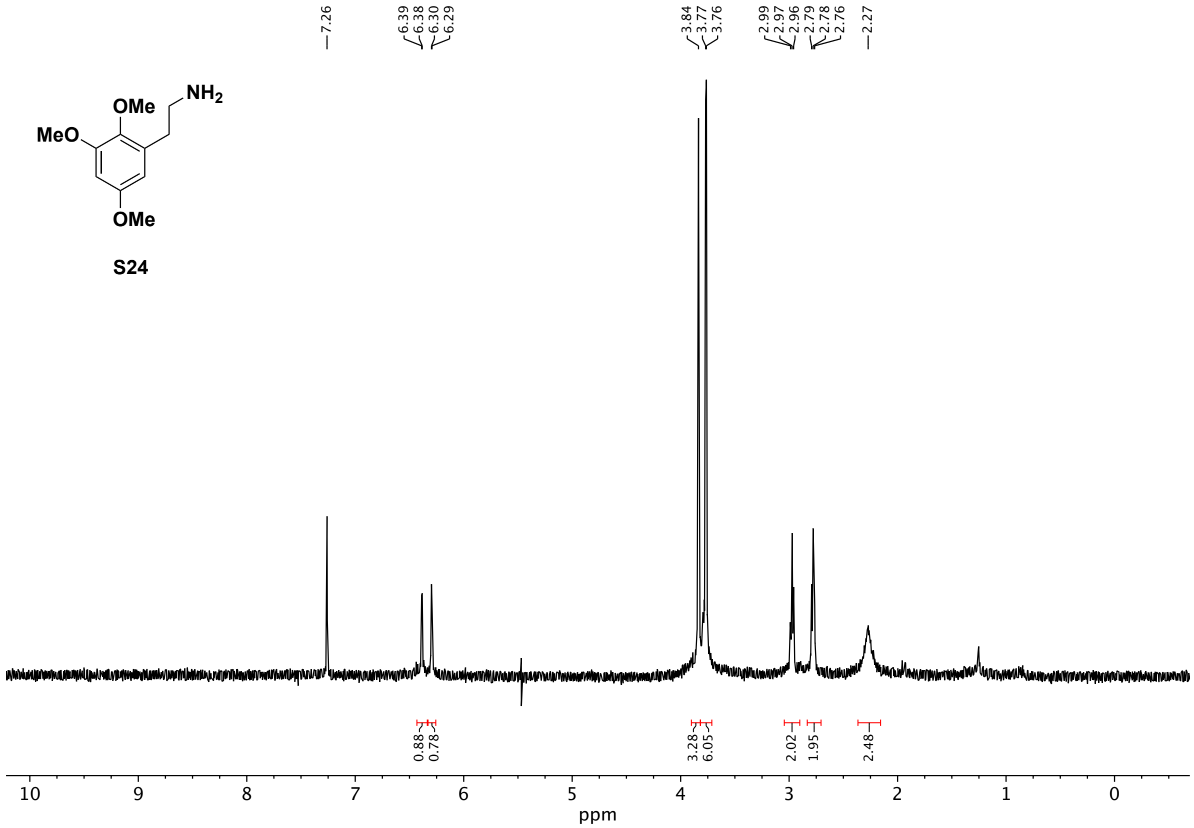
OMe
NH2
MeO
OMe
7.26
6.39
6.38
6.30
6.29
3.84
3.77
3.76
2.99
2.97
2.96
2.79
2.78
2.76
2.27
0.88
0.78
3.28
6.05
2.02
1.95
2.48
10
9
8
7
6
5
4
3
2
1
0
ppm

S24

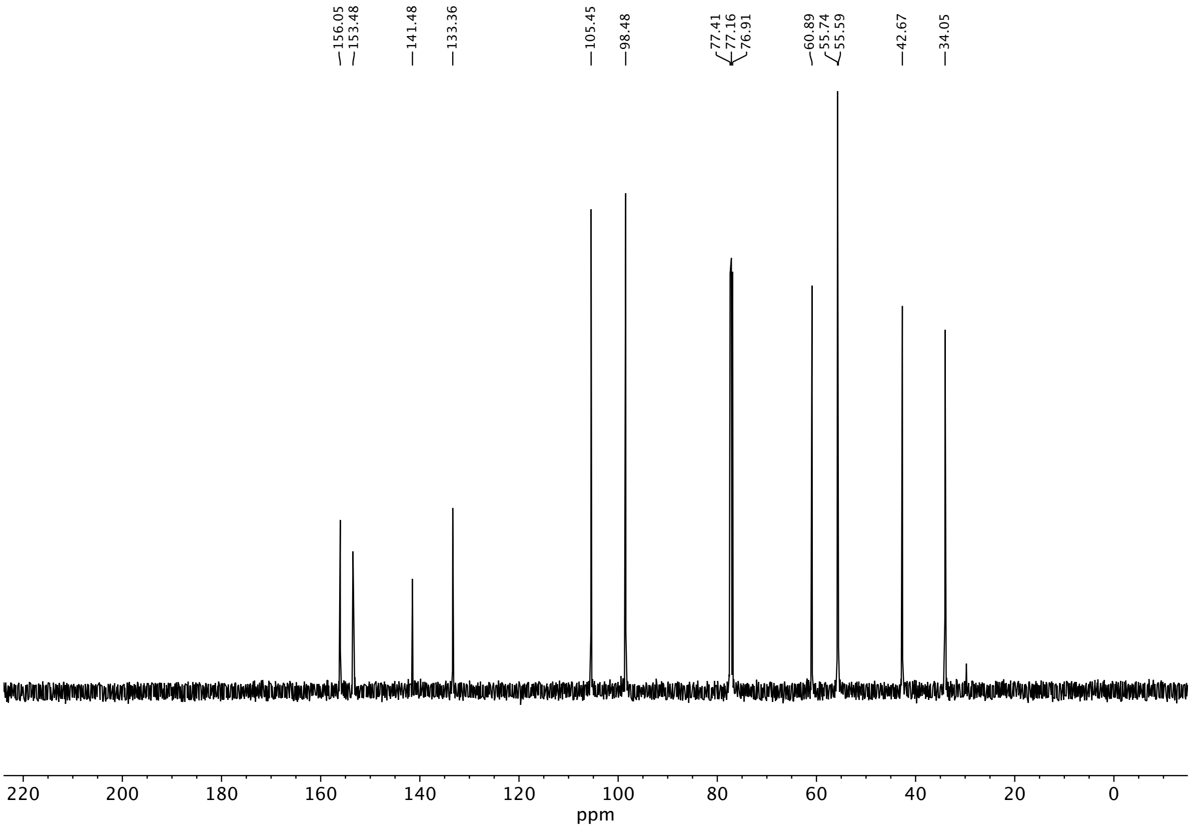
156.05
153.48
141.48
133.36
105.45
98.48
77.41
77.16
76.91
60.89
55.74
55.59
42.67
34.05
220
200
180
160
140
120
100
80
60
40
20
0
ppm

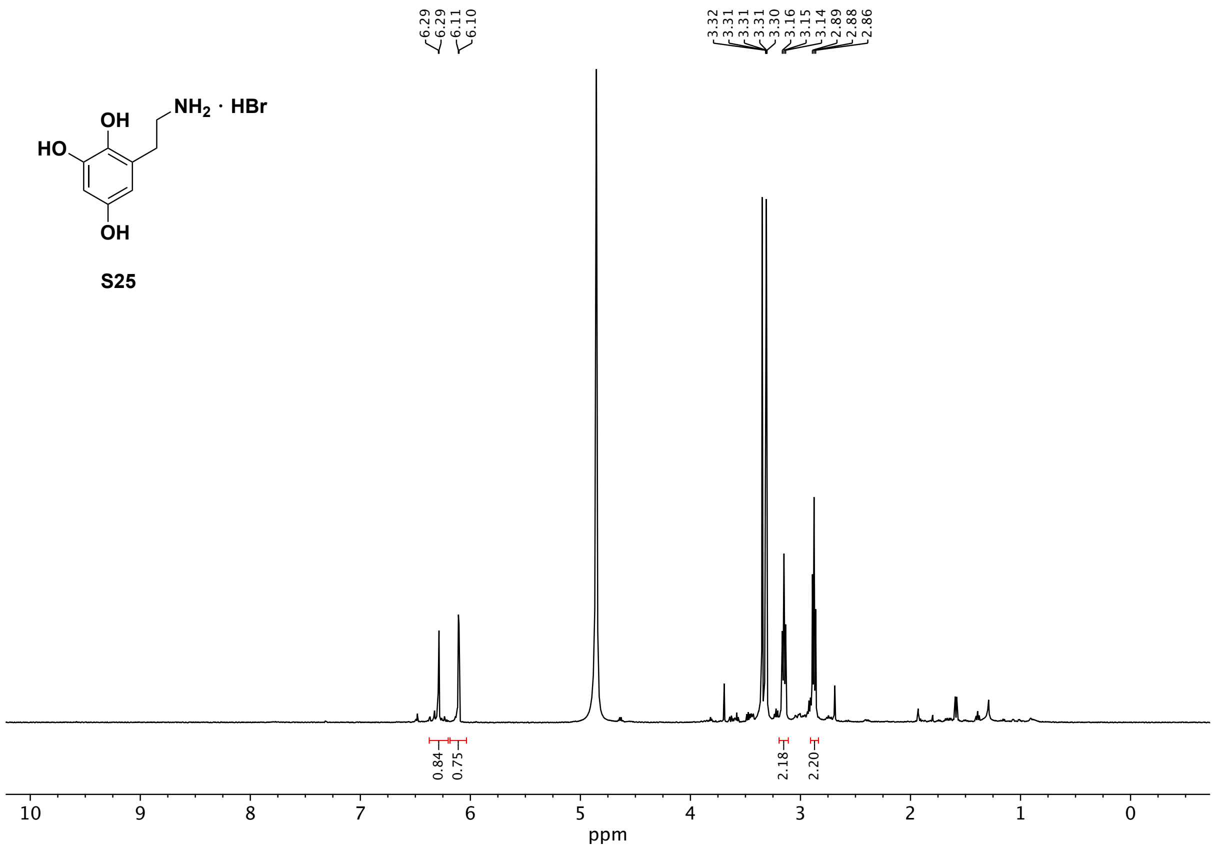

NH2 · HBr
OH
HO
OH
S25
6.29
6.29
6.11
6.10
3.32
3.31
3.31
3.31
3.30
3.16
3.15
3.14
2.89
2.88
2.86
0.84
0.75
2.18
2.20
10
9
8
7
6
5
4
3
2
1
0
ppm

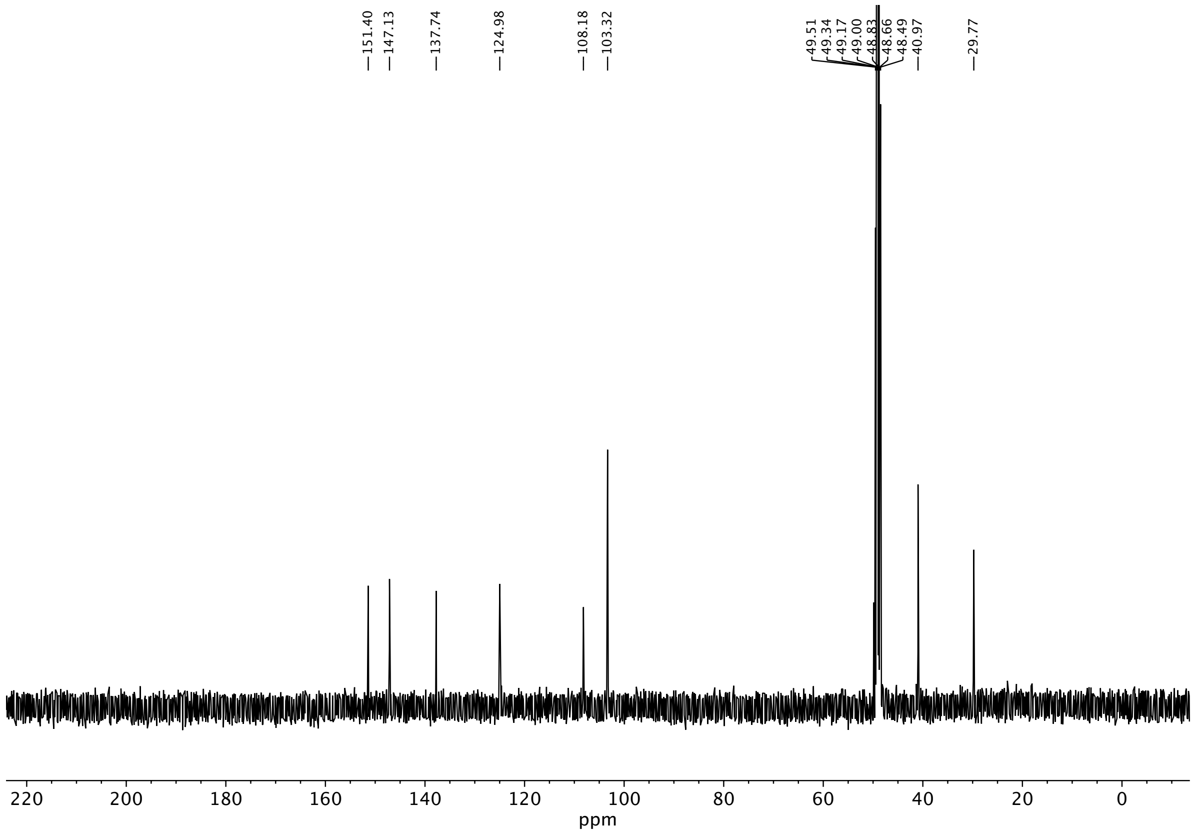
151.40
147.13
137.74
124.98
108.18
103.32
49.51
49.34
49.17
49.00
48.83
48.66
48.49
40.97
29.77
220 200 180 160 140 120 100 80 60 40 20 0
ppm

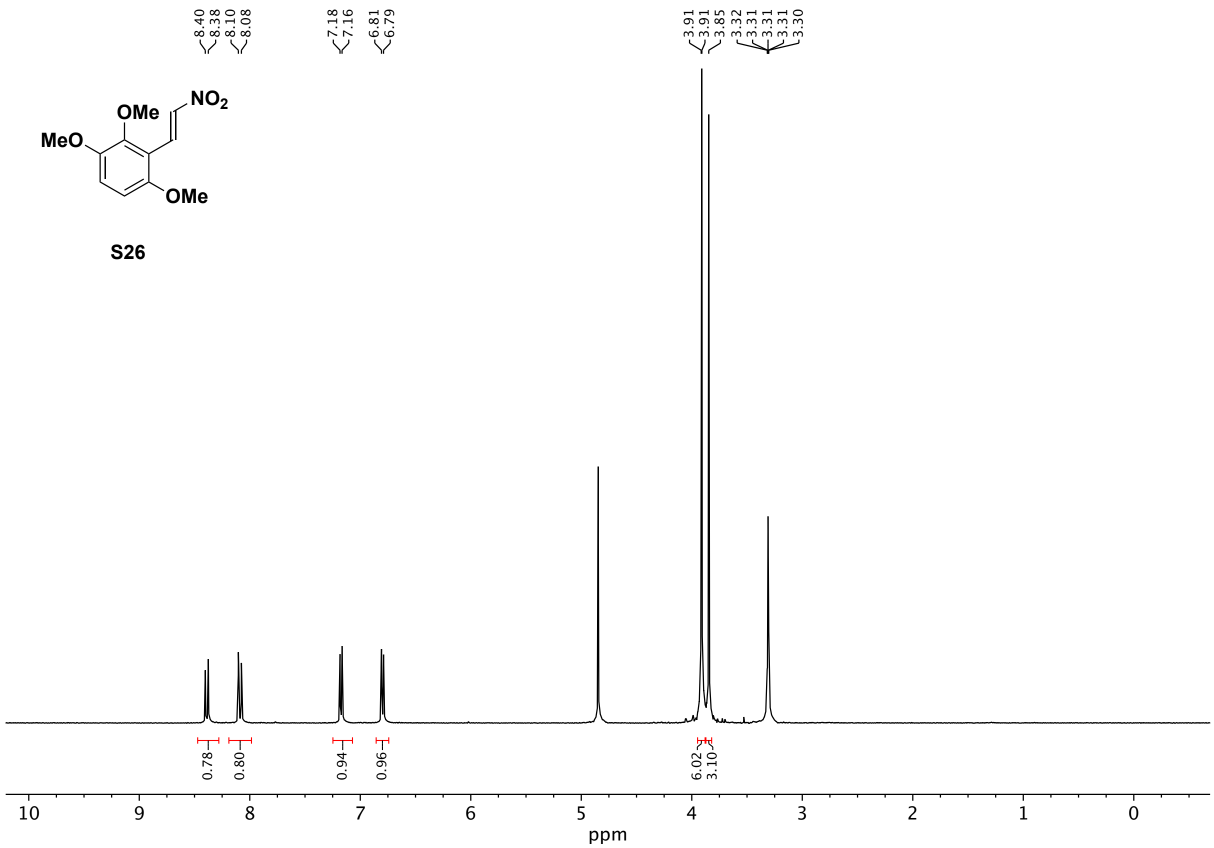
8.40
8.38
8.10
8.08
7.18
7.16
6.81
6.79
3.91
3.91
3.85
3.32
3.31
3.31
3.31
3.30
NO2
OMe
MeO
OMe
S26
0.78
0.80
0.94
0.96
6.02
3.10
10
9
8
7
6
5
4
3
2
1
0
ppm

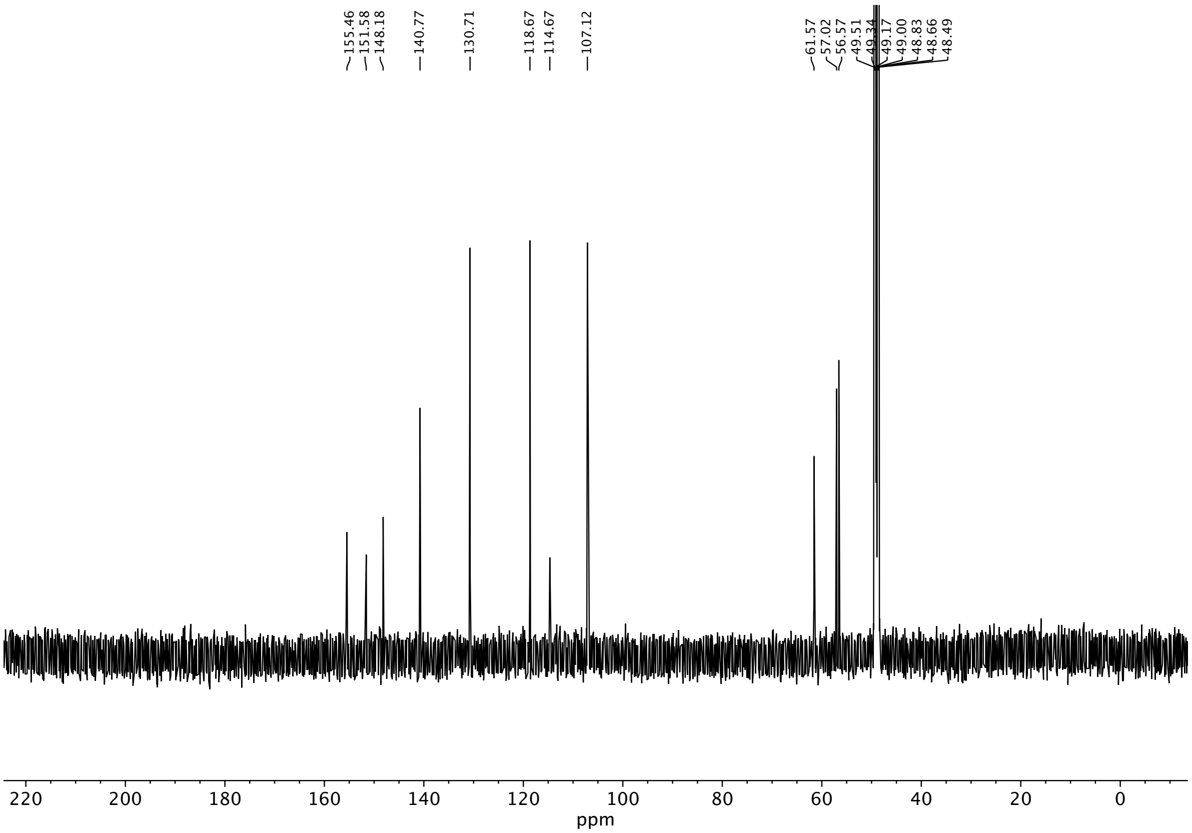

155.46
151.58
148.18
140.77
130.71
118.67
114.67
107.12
61.57
57.02
56.57
49.51
49.34
49.17
49.00
48.83
48.66
48.49
220
200
180
160
140
120
100
80
60
40
20
0
ppm

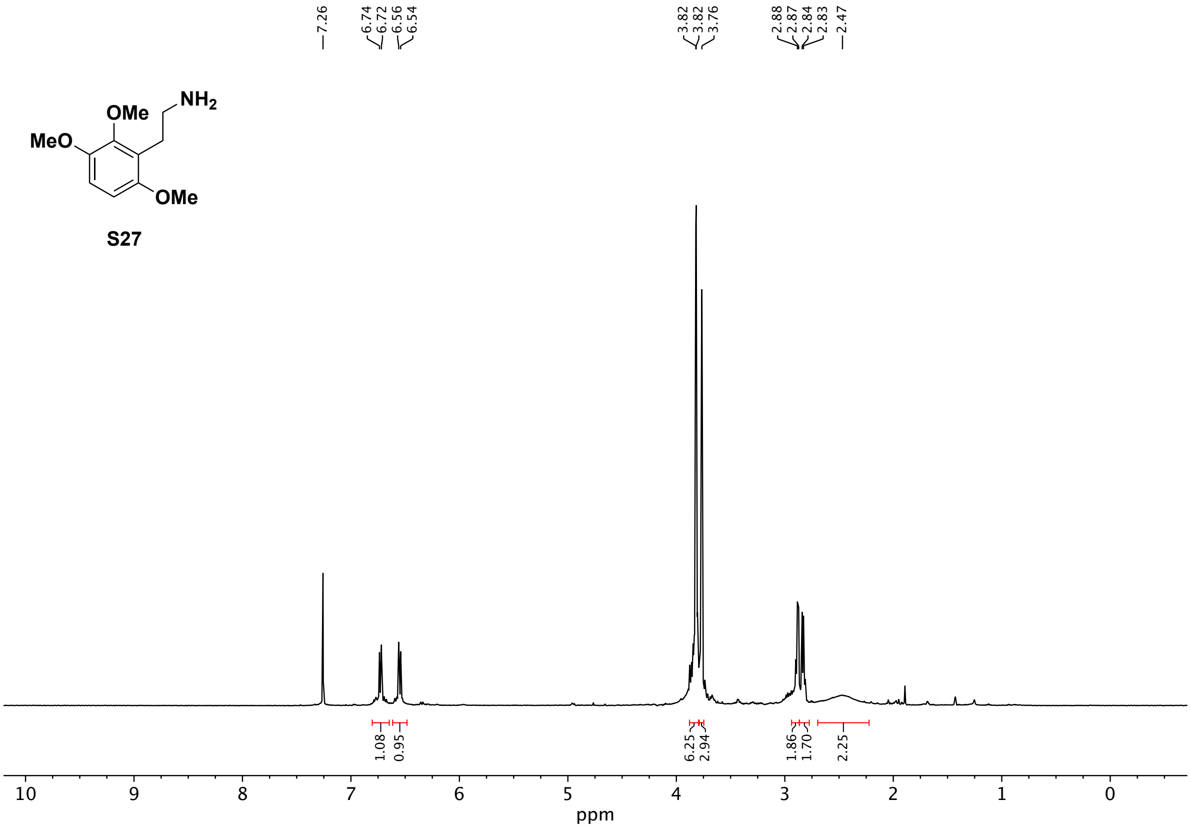
OMe
NH2
MeO
OMe
7.26
6.74
6.72
6.56
6.54
3.82
3.82
3.76
2.88
2.87
2.84
2.83
2.47
1.08
0.95
6.25
2.94
1.86
1.70
2.25
10
9
8
7
6
5
4
3
2
1
0
ppm

S27

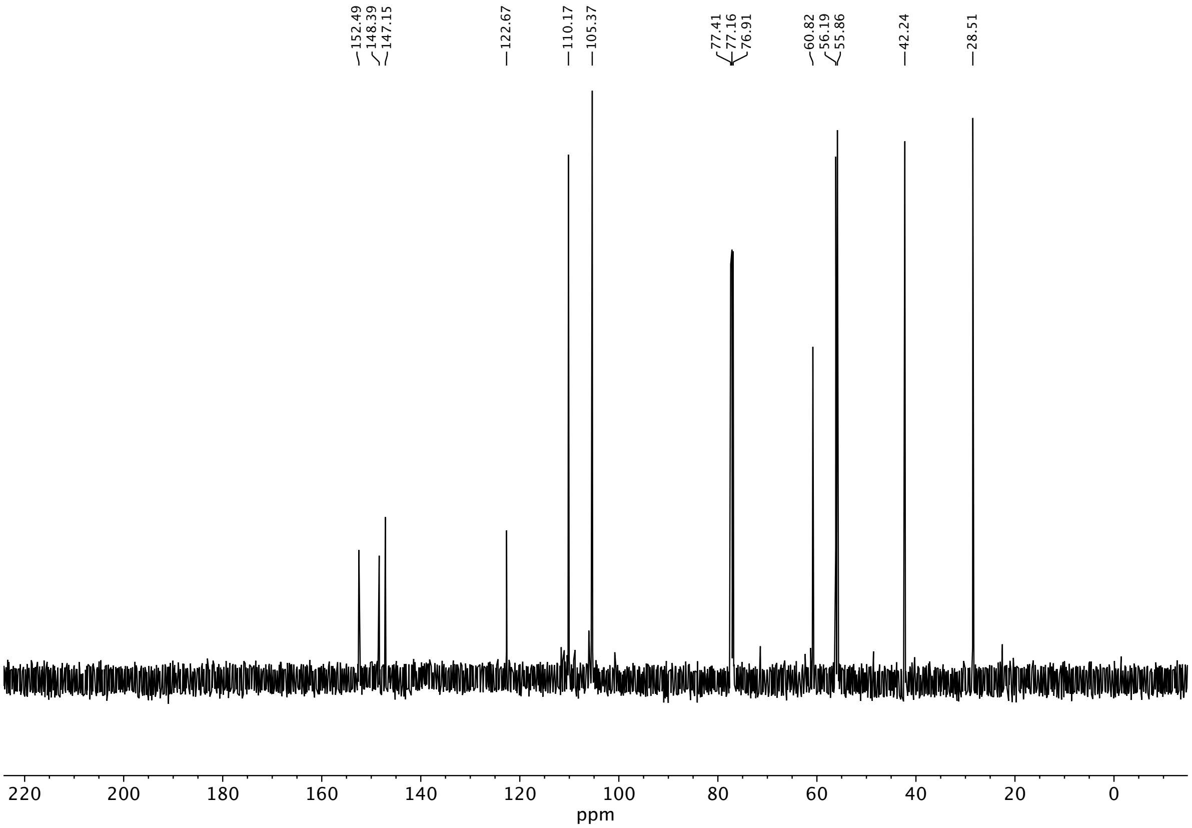
152.49
148.39
147.15
122.67
110.17
105.37
77.41
77.16
76.91
60.82
56.19
55.86
42.24
28.51
220
200
180
160
140
120
100
80
60
40
20
0
ppm

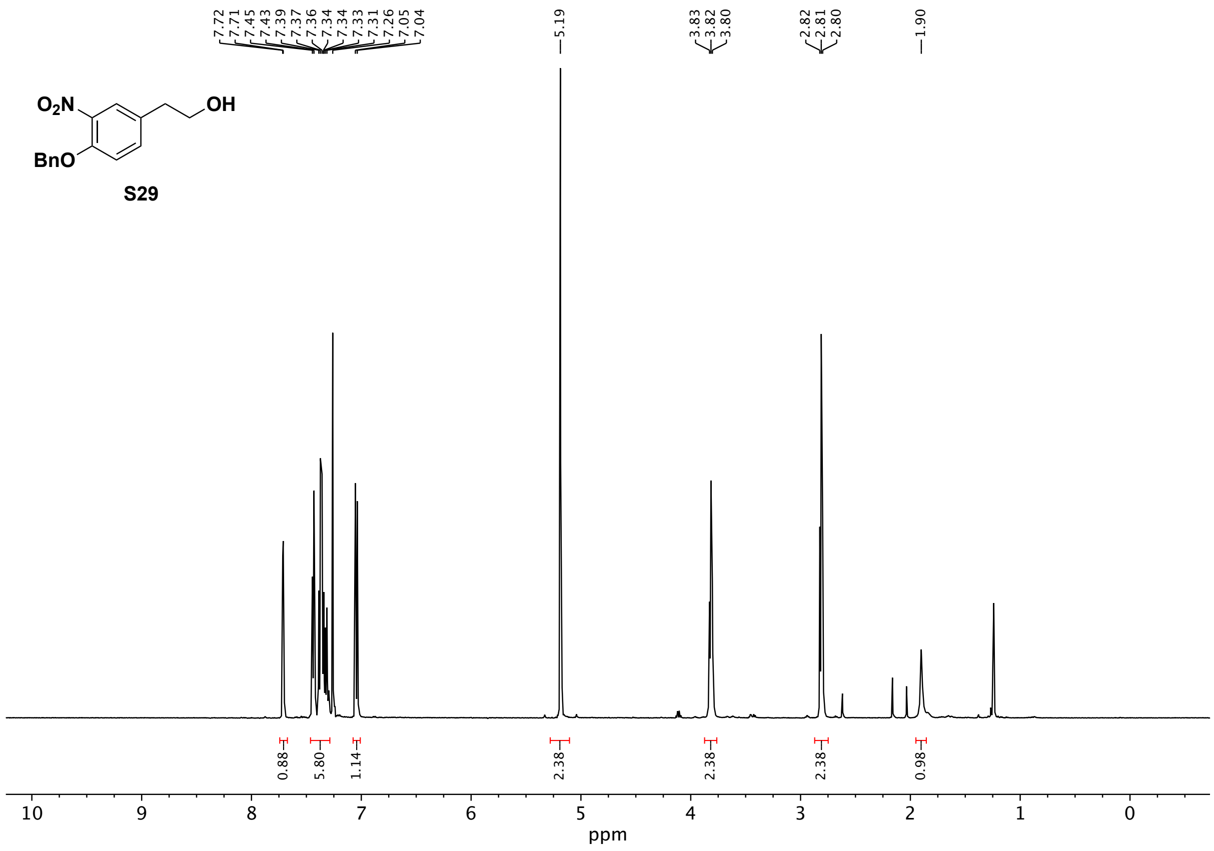
O2N
OH
BnO
S29
7.72
7.71
7.45
7.43
7.39
7.37
7.36
7.34
7.34
7.33
7.31
7.26
7.05
7.04
5.19
3.83
3.82
3.80
2.82
2.81
2.80
1.90
0.88
5.80
1.14
2.38
2.38
2.38
0.98
10
9
8
7
6
5
4
3
2
1
0
ppm

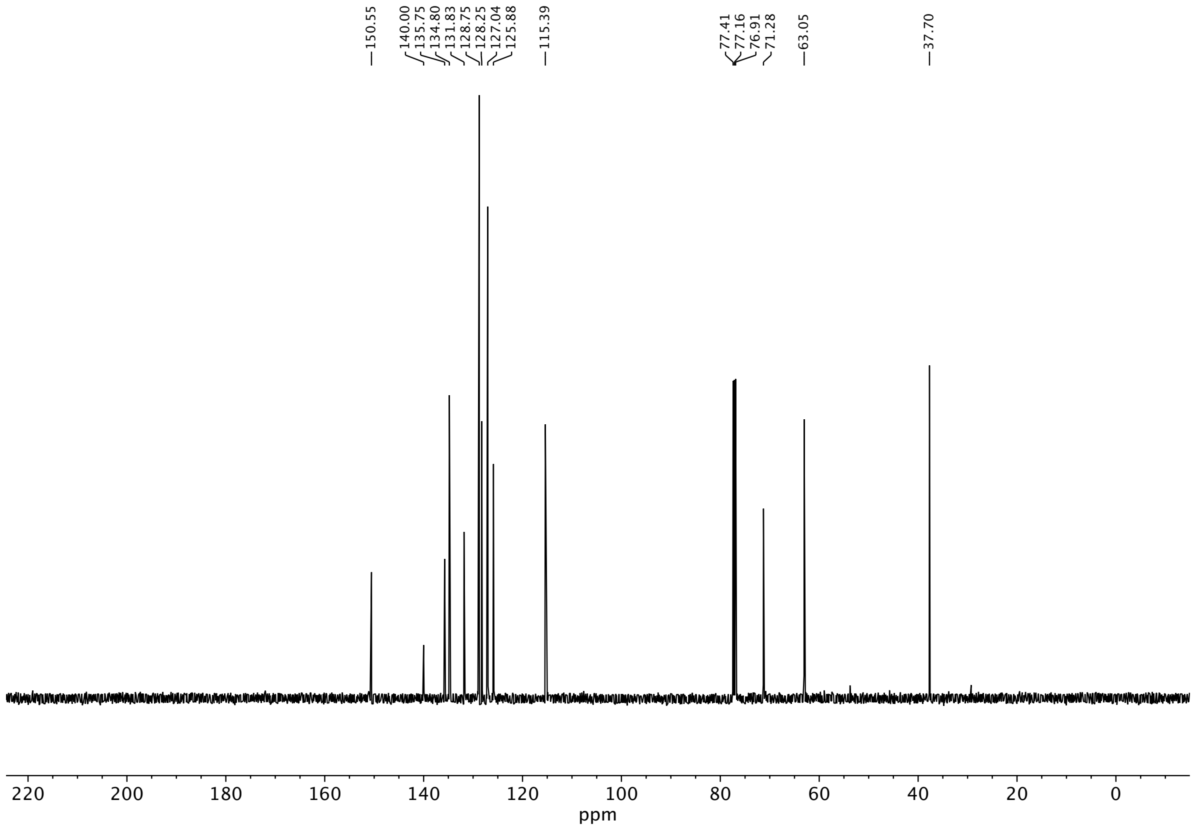
150.55
140.00
135.75
134.80
131.83
128.75
128.25
127.04
125.88
115.39
77.41
77.16
76.91
71.28
63.05
37.70
220
200
180
160
140
120
100
80
60
40
20
0
ppm

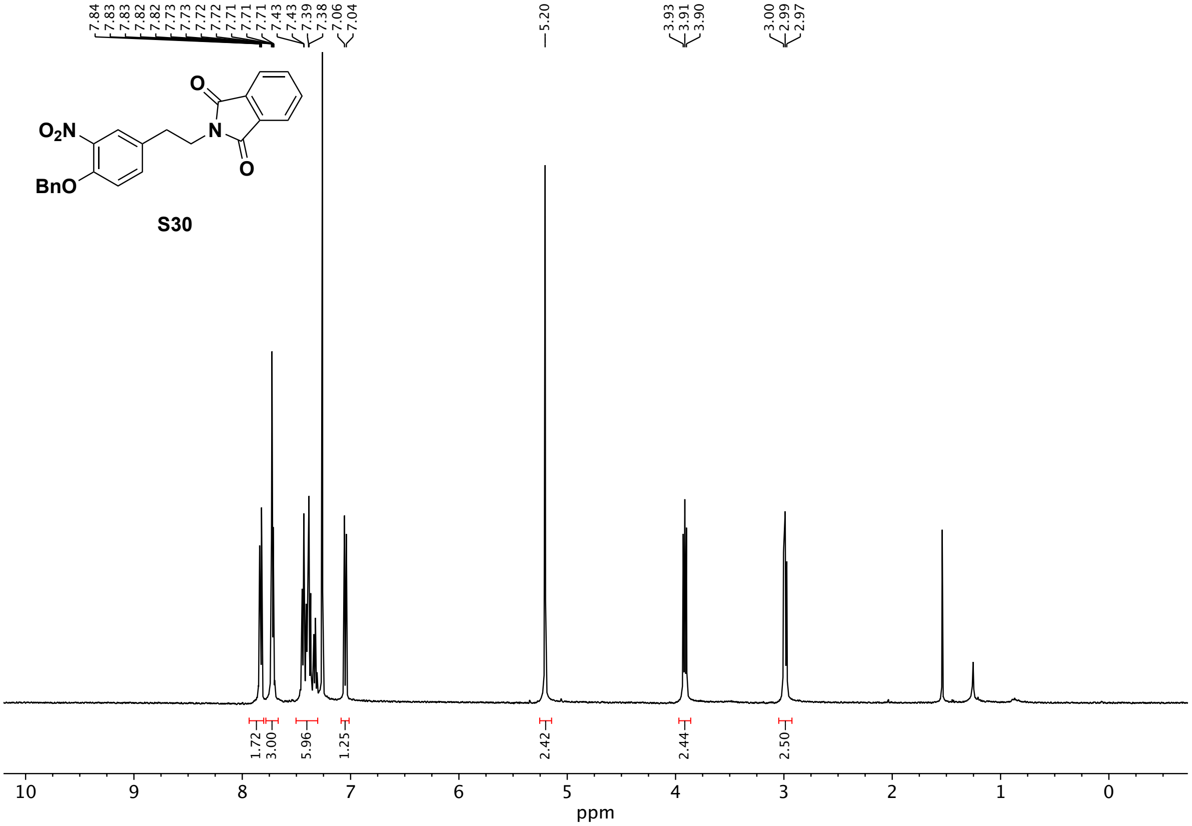
7.84
7.83
7.83
7.82
7.82
7.73
7.73
7.72
7.72
7.71
7.71
7.71
7.43
7.43
7.39
7.38
7.06
7.04
5.20
3.93
3.91
3.90
3.00
2.99
2.97
O
O
O₂N
N
BnO
1.72
3.00
5.96
1.25
2.42
2.44
2.50
10
9
8
7
6
5
4
3
2
1
0
ppm

S30

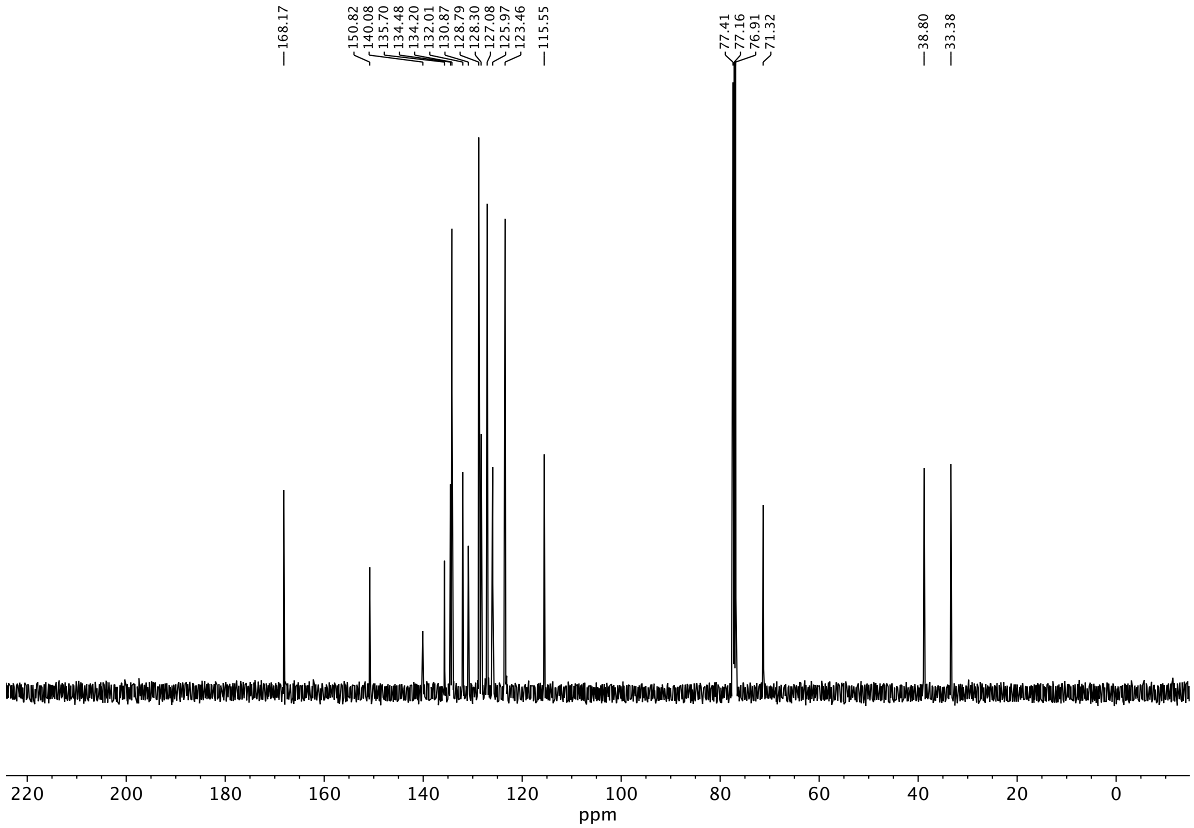
168.17
150.82
140.08
135.70
134.48
134.20
132.01
130.87
128.79
128.30
127.08
125.97
123.46
115.55
77.41
77.16
76.91
71.32
38.80
33.38
220 200 180 160 140 120 100 80 60 40 20 0
ppm

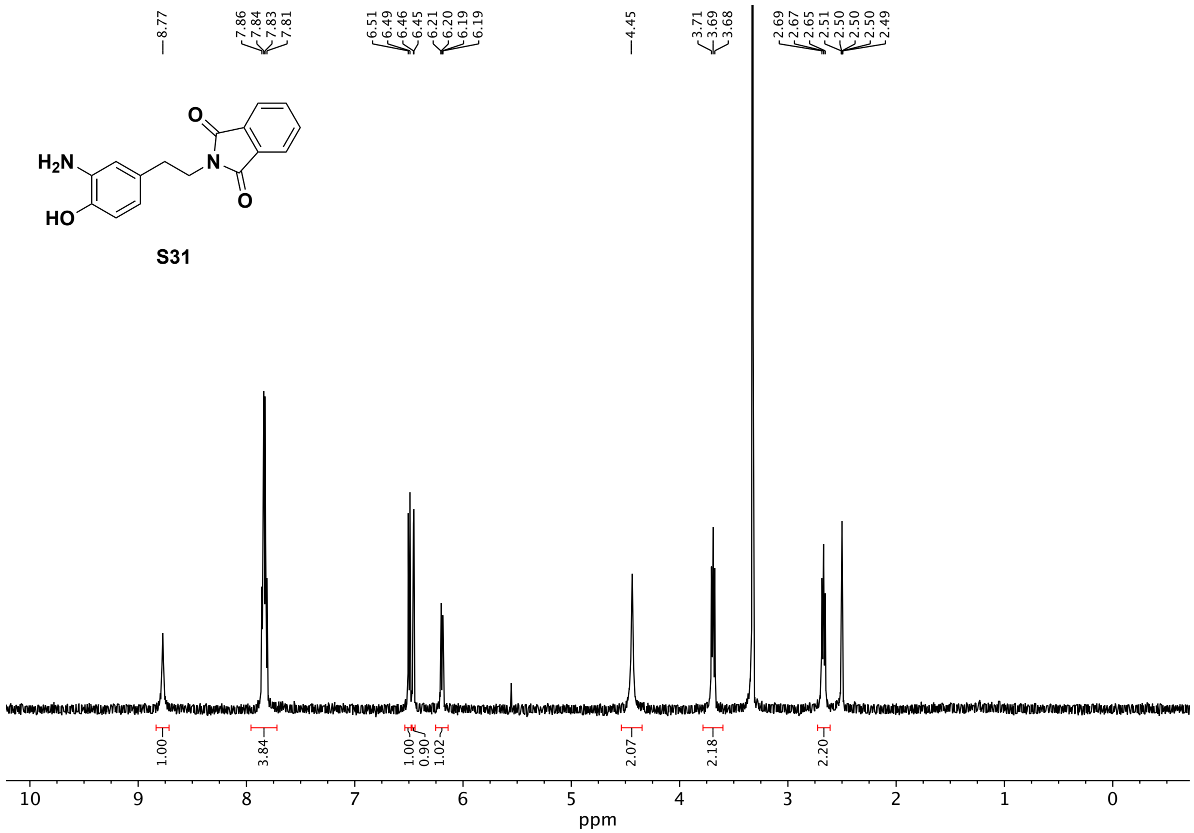
8.77
7.86
7.84
7.83
7.81
6.51
6.49
6.46
6.45
6.21
6.20
6.19
6.19
4.45
3.71
3.69
3.68
2.69
2.67
2.65
2.51
2.50
2.50
2.50
2.49
H2N
HO
O
N
O
S31
1.00
3.84
1.00
0.90
1.02
2.07
2.18
2.20
10
9
8
7
6
5
4
3
2
1
0
ppm

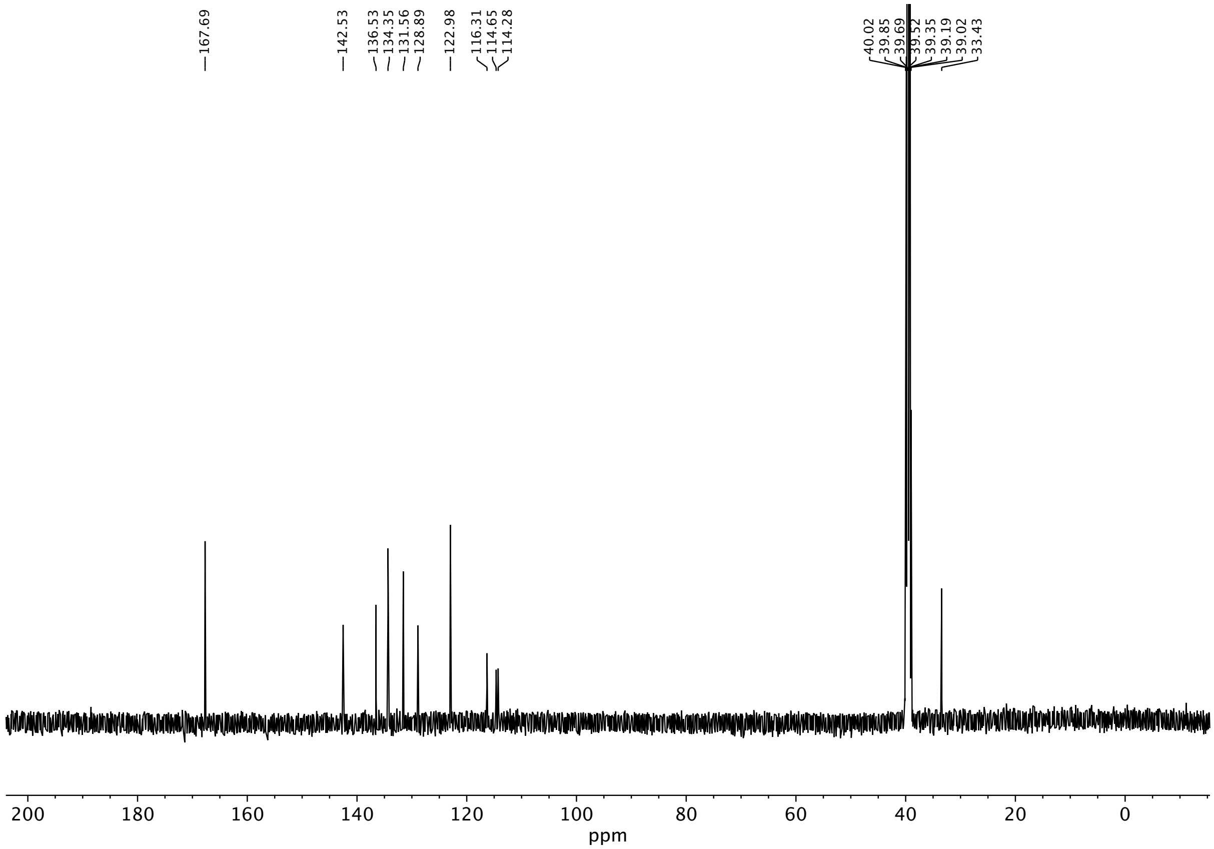

167.69
142.53
136.53
134.35
131.56
128.89
122.98
116.31
114.65
114.28
40.02
39.85
39.69
39.52
39.35
39.19
39.02
33.43
200
180
160
140
120
100
80
60
40
20
0
ppm

**S32**

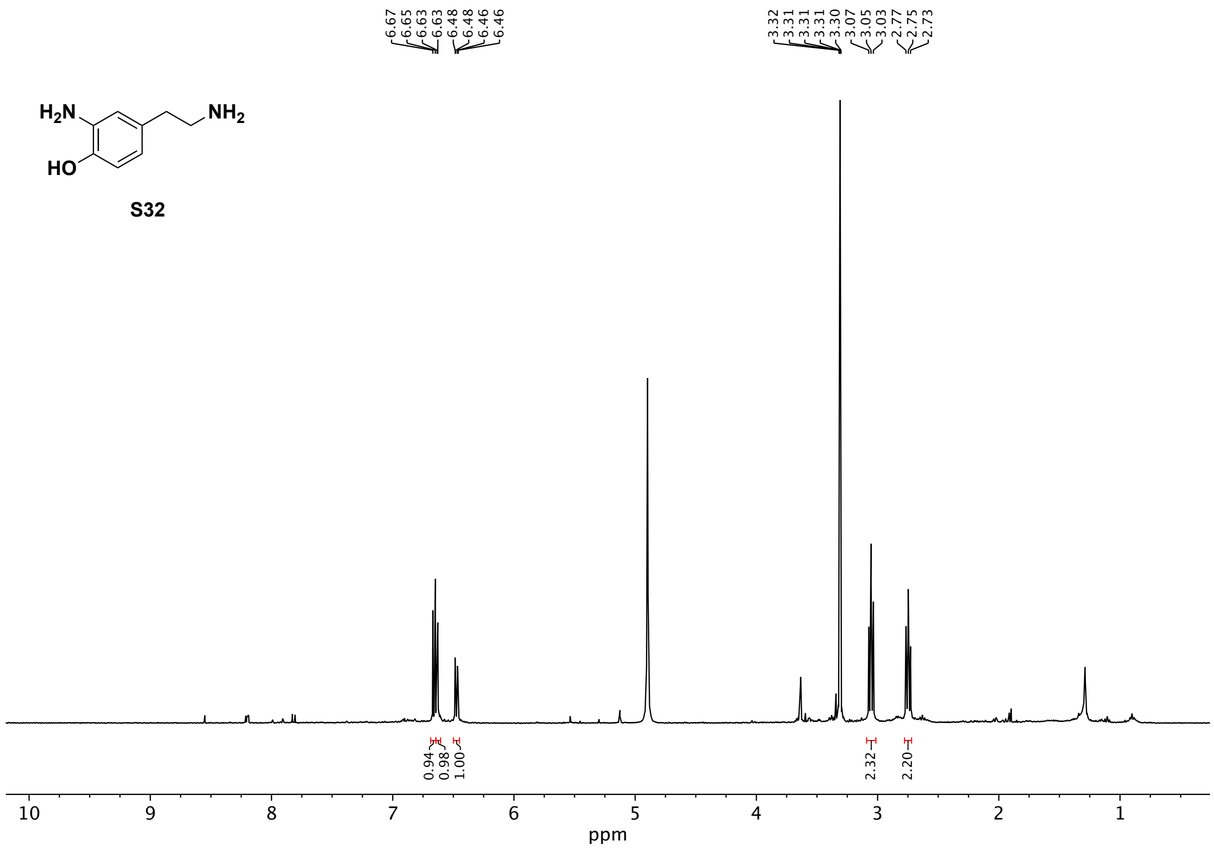
6.67
6.65
6.63
6.63
6.48
6.48
6.46
6.46
3.32
3.31
3.31
3.31
3.30
3.07
3.05
3.03
2.77
2.75
2.73
0.94
0.98
1.00
2.32
2.20
10
9
8
7
6
5
4
3
2
1
ppm

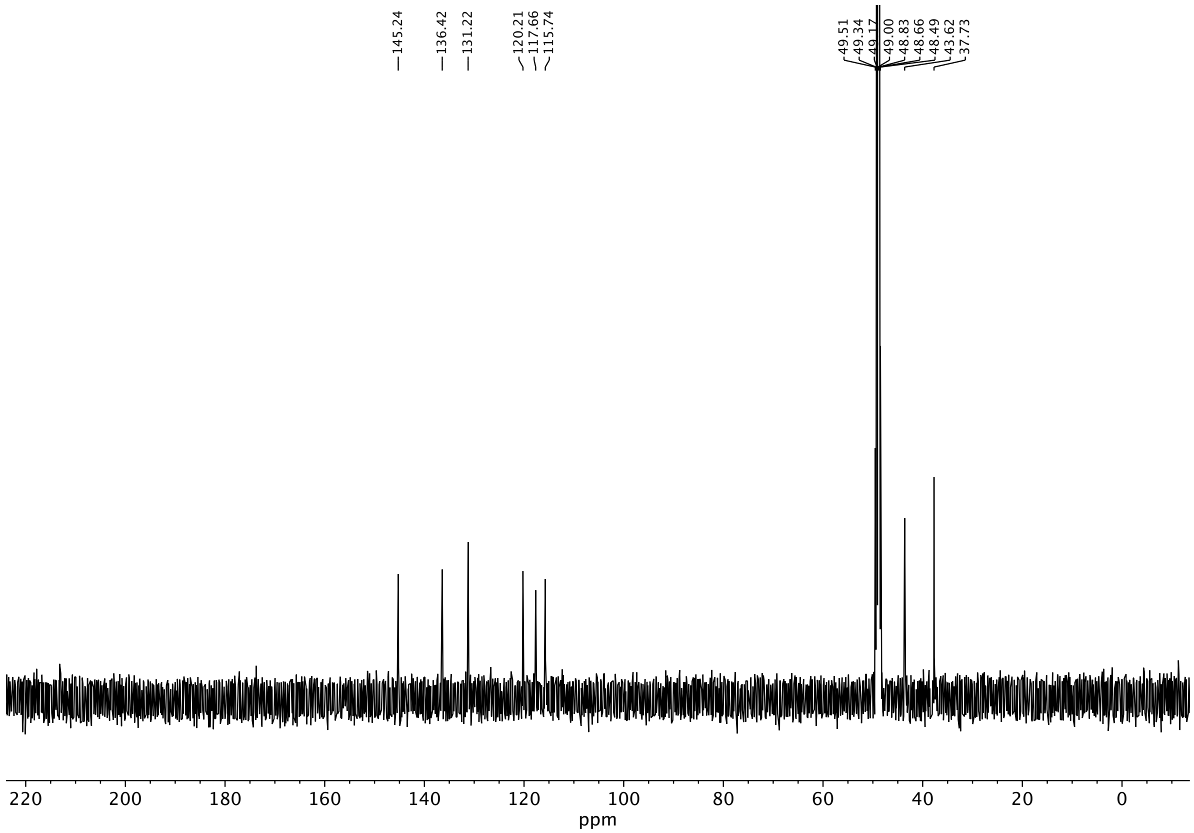
145.24
136.42
131.22
120.21
117.66
115.74
49.51
49.34
49.17
49.00
48.83
48.66
48.49
43.62
37.73
220
200
180
160
140
120
100
80
60
40
20
0
ppm
